# Genome-wide association study of plasma triglycerides, phospholipids and relation to cardio-metabolic risk factors

**DOI:** 10.1101/621334

**Authors:** Ayşe Demirkan, Rene Pool, Joris Deelen, Marian Beekman, Jun Liu, Amy C. Harms, Anika Vaarhorst, Fiona A Hagenbeek, Gonneke Willemsen, Aswin Verhoeven, Najaf Amin, Ko Willems van Dijk, Thomas Hankemeier, Dorret I. Boomsma, Eline Slagboom, Cornelia M. van Duijn

**Affiliations:** Department of Epidemiology, Erasmus Medical Center, Rotterdam, the Netherlands; Department of Clinical and Experimental Research, Faculty of Bioscience and Medicine, Univeristy of Surrey, Guildford, United Kingdom; Department of Genetics, Univeristy Medical Center Groningen, Groningen, The Netherlands; Netherlands Twin Register, Department of Biological Psychology, Vrije Universiteit Amsterdam, Van der Boechorststraat 7, 1081 BT Amsterdam, The Netherlands; Molecular Epidemiology, Department of Biomedical Data Sciences, Leiden University Medical Center, Leiden, the Netherlands; Max Planck Institute for Biology of Ageing, Cologne, Germany; Division of Analytical Biosciences, Leiden Academic Centre for Drug Research, Leiden University, Leiden, the Netherlands; Department of Human Genetics, Leiden University Medical Center, Leiden, the Netherlands; Department of Internal Medicine, division Endocrinology, Leiden University Medical Center, Leiden, the Netherlands; Netherlands Metabolomics Centre, Leiden University, Leiden, the Netherlands; Nuffield Department of Population Health, Oxford University, Oxford, United Kingdom

**Author notes:** The authors contributed equally to this work (Ayşe Demirkan, Rene Pool, Joris Deelen). **Corresponding author**: Cornelia van Duijn, Nuffield Department of Population Health, Oxford University, Oxford, United Kingdom.

## Abstract

There is continuous interest in the genetic determinants of plasma triglycerides (TGs) and phospholipids and their role in the etiology of cardiovascular disease (CVD). Here, we report the results of a Dutch genome wide association study (GWAS) of an in-house developed lipidomics platform, focusing on 90 plasma lipids. Lipids were assessed by liquid chromatography mass spectrometry in participants from the Leiden Longevity Study, the Netherlands Twin Register and the Erasmus Rucphen Family (ERF) study and meta-analysed, resulting in a sample size of 5537 participants. In addition, we performed genetic correlation analyses between the 90 plasma lipids and markers of metabolic health, as well as vascular pathology and CVD combining our GWAS results with publicly available GWAS outputs. We replicated previously known associations between 34 lipids and 10 lipid quantitative trait loci (lipQTL) (*GCKR, APOA1, FADS1, SGPP1,TMEM229B, LIPC, PDXDC1, CETP, CERS4* and *SPTLC3*) with metabolome-wide (P < 1.61 × 10^−9^) significance. Moreover, we report 6 novel phospholipid-related and 5 triglyceride (TG)-related loci: *SGGP1* (SM21:0), *SPTLC3* (SM21:0 and SM25:1), *FADS1* (LPCO16:1, PC38:2, PEO36:5, PEO38:5, TG56:5, TG56:6, and TG56:7), *TMEM229* (LPCO16:1), *GCKR* (TG50:2), and *APOA1* (TG54:4). In addition, we report suggestively significant (P < 5 × 10^−8^) associations mapping to eleven novel lipid quantitative trait loci (lipQTLs), three of which are supported by mining previous GWAS data: *MAU* (PC34:4), *LDLR* (SM16:0), and *MLXIPL* (TG48:1 and TG50:1)). Genetic correlation analysis indicates that one specific specific sphingomyelin, SM22:0, shares common genetic background with CVD. Levels of SM22:0 also positively associate with carotid artery intima-media thickness in the ERF study, and this observation is independent of LDL-C level. Our findings yield higher resolution of plasma lipid species and new insights in the biology of circulating phosholipids and their relation to CVD risk.

## Introduction

Although numerous genetic loci have been associated with metabolic diseases^1,2^ and disease markers^3,4^, functional interpretation of these loci is lagging discovery. Plasma metabolites are hypothesized to function as markers and mediators of cardiovascular disease (CVD) and extensive efforts have sought to both refine and expand our understanding of the causal determinants of circulating metabolite levels^5–7^. Notably, loci have been identified that encode enzymes or transport proteins directly involved in a given metabolite’s turnover^5–8^. Many of these loci have shown relatively large effect sizes on metabolite levels, as compared to effect sizes in genome wide association studies (GWAS) for common diseases, and explain a relatively large proportion of the heritability of these metabolites ^9,10^.

Total circulating triglycerides (TGs) and lipoproteins are established risk factors for both type 2 diabetes (T2D) and CVD^4^. The genetic determinants of circulating total TGs have been partially uncovered by the Global Lipids Genetics Consortium (GLGC)^3^. However, the genetics underlying the individual TG species are largely unknown. Rhee and colleagues have previously addressed this question in 2076 participants of the Framingham Heart Study (FHS)^11^. They found 23 novel genetic loci associated with plasma metabolites, including a set of lipid species that were not investigated in prior GWAS^11^. We aimed to expand upon these findings using a larger sample size and a denser population specific genotype imputation panel. In addition, we exploited a liquid chromatography mass spectrometry (LC-MS)-based lipidomics platform that measures lipids in plasma and serum^12^. This platform additionally includes 4 ether phospholipids (PCO16:1, PCO36:6, PEO36:5, PCO38:5), 3 sphingomyelins (SM17:0, SM 21:0, SM25:1) and 5 TGs (TG50:0, TG51:1, TG51:2, TG51:4 and TG53:1) which have not yet been previously analyzed by other platforms. We investigated the genetics underlying the plasma lipidome in 5537 Dutch individuals whose genotypes were imputed using the Genome of Netherlands (GoNL) reference panel. We further investigated the genetic overlap of identified lipids with CVD-related traits and addressed the dynamics of plasma TGs by comparing the genetic determinants of total plasma TGs to the genetic determinants of individual TG species.

## Methods

### Study populations

The study sample consisted of 5537 participants from 3 Dutch population-based cohorts; the Leiden Longevity Study (LLS)^13^, the Erasmus Rucphen Family (ERF) study^14^ and the Netherlands Twin Register (NTR)^15^. An overview of all samples is provided in Table 1. More detailed information on the design of the cohorts can be found in the **Supplementary text**.

**Table 1.**
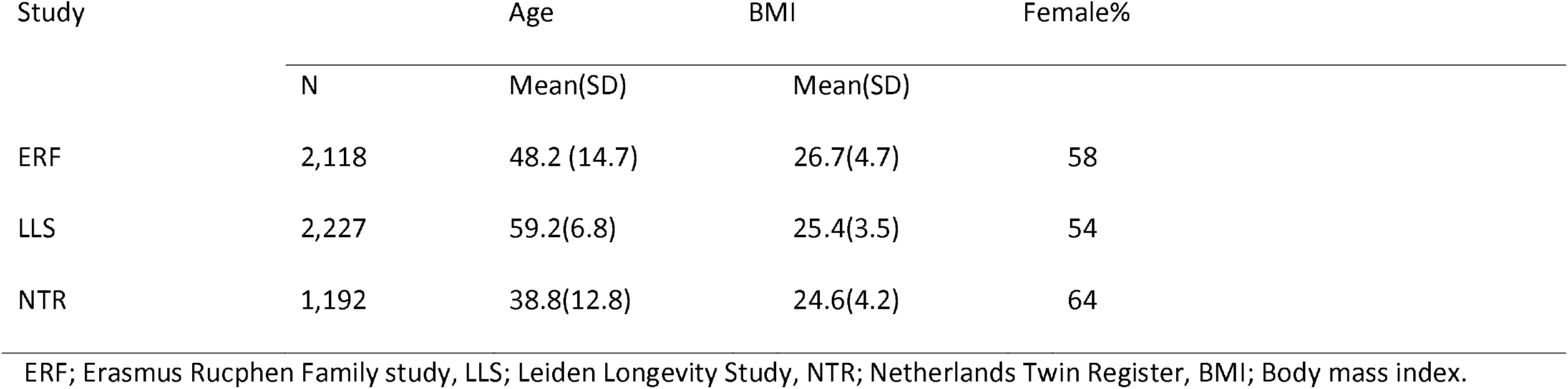
Description of the samples that were used in the meta-analyses

### Metabolomics measurements

Measurements of the three cohorts on a single metabolomics platform were performed as part of the BBMRI-NL initiative (www.bbmri.nl). The plasma lipids were measured by LC-MS using the method described in Gonzalez-Covarrubias et al^12^. For ERF and NTR, the samples were collected after overnight fasting and for the LLS we used non-fasted samples. The 90 lipid species that were successfully measured in all three cohorts and were eligible for meta-analysis comprised 30 TGs, 39 phosphatidylcholines, 4 phosphatidylethanolamines, and 17 sphingolipids. Lipid names and abbreviations were assigned according to the Lipid Maps nomenclature (http://www.lipidmaps.org). The following abbreviations are used: triglyceride, TG; acyl-acyl phosphatidylcholine, PC; alkyl-acyl phosphatidylcholine, PCO lysophosphatidylcholine, LPC; sphingomyelin, SM; acyl-acyl phosphatidylethanolamines, PE and alkyl-acyl phosphatidylethanolamines, PEO. Samples were excluded if the participants used lipid lowering medication or if more than 10% of the lipid markers were reported as missing values.

### Gaussian graphical model (GGM) network

The interactions between the metabolic phenotypes were visualized using a Gaussian graphical model (GGM)^16^. In concordance with the GWAS analyses (c.f. below), the phenotypes were natural log(Ln)-transformed and standardized. The partial correlation network was based on the residual values for each metabolic phenotype after regressing out age and sex. We quantified the separation of chemical classes in this network by calculating their modularities. The width of the edges scales with the absolute value of the partial correlation between nodes.

### GWAS

Genotype data from all three Dutch cohorts were imputed according to the custom-built Genome of the Netherlands project reference panel (GoNL, http://www.nlgenome.nl/), which is based on the genomes of 250 parent-offspring trios that were sequenced at ∼13 x coverage^17,18^ and has previously been used to detect low frequency variants for plasma cholesterol^19^. For this study, we used version 4 of the reference panel. The association analysis was performed using linear regression with the lipid levels as outcome, adjusted for age, sex, and study specific covariates, such as familial relatedness. The genotyping and imputation QC of each cohort are provided in **Table S1**.

### Meta-analysis

Meta-analysis across studies was performed using an inverse variance weighted fixed-effects model^20^, by two separate analysts in parallel, implemented in the METAL^21^ and GWAMA^22^ software. The genome-wide significant p-value (5 × 10^−8^) was adjusted for the largest number of independent variables (as identified using the method of Li and Ji^23^) found among the three cohorts (N=31). Hence, the metabolome-wide adjusted significance level was set at P < 1.61 × 10 ^−9^ and the suggestive significance level at P < 5 × 10 ^−8^.

### Phewas of the top lipQTLs

We looked up PHEWAS results from Pheweb (pheweb.sph.umich.edu) and GWASATLAS (http://atlas.ctglab.nl/PheWAS) and reported the associations with genome-wide significance (P < 5 × 10^−8^).

### Genetic correlation analysis

Genetic correlation analysis between the lipids and other phenotypes was performed using the protocol on: https://github.com/bulik/ldsc/wiki/Heritability-and-Genetic-Correlation, which is based on a previous study by Bulik-Sullivan et al.^24^. We also tested the partitioning of heritability as proposed in ^25^ (see: https://github.com/bulik/ldsc/wiki/Partitioned-Heritability). For the genetic correlation analyses, we used 21 datasets from published GWAS results of body-mass index (BMI)^26^, fat percentage^27^, waist-to-hip ratio (WHR) ^28^, childhood obesity^29^, leptin^30^, adiponectin^31^, systolic blood pressure (SBP)^32^, diastolic blood pressure (DBP)^32^, heart rate ^33^, cardiovascular disease (CVD)^34^, carotid intima-media thickness (IMT) and plaque,^35^ LDL cholesterol^36^, HDL cholesterol^15^, TGs^3^, glucose^37^, Hb1Ac^38^, proinsulin^39^, 2-hour glucose^40^, insulin^41^ and T2D^36^. A false discovery rate (FDR) of 0.05 per experiment was applied to adjust for multiple testing.

## Results

### Gaussian graphical modelling

The GGM network of the 90 lipids measured in NTR study is shown in Figure 1. All partial correlations were observed to be positive. The network shows complete modularity across the phospholipid, sphingolipid species and TGs. We did not observe a distinguished modularity between the phospholipid subgroups i.e. lysophosphatidylcholine (LPC), phosphatidylethanolamines (PE) and phosphatidylcholines (PCs). The clusters within the phospholipid group were driven by two factors; 1) by the chemical bond of residues as all alkyl-acyl bonding types (marked as an additional-O in the abbreviations) clustered together regardless of their choline or ethanolamine head-groups, e.g. PCO36:2, PCO36:5 and PCO38:5; and 2) by the saturation degree of the fatty acid residues, as polyunsaturated fatty acids carrying the same number of double bonds clustered together, e.g. PC38:6 and PC40:6. We next checked individual lipids in the TG pool and estimated their contribution to the total TG in the circulation in the ERF population. The majority (72%) of the TG pool consists of six TGs: TG52:2 (22%), TG52:3 (21%), TG50:2 (8%), TG52:4 (8%), TG50:1 (7%) and TG54:4 (6%). The contribution of the other species was less than 5% per measured lipid (**Figure S1**). All TG species correlated strongly and positively to the total TG (measured by enzymatic method), including the TGs present in trace amounts.

**Figure 1.**
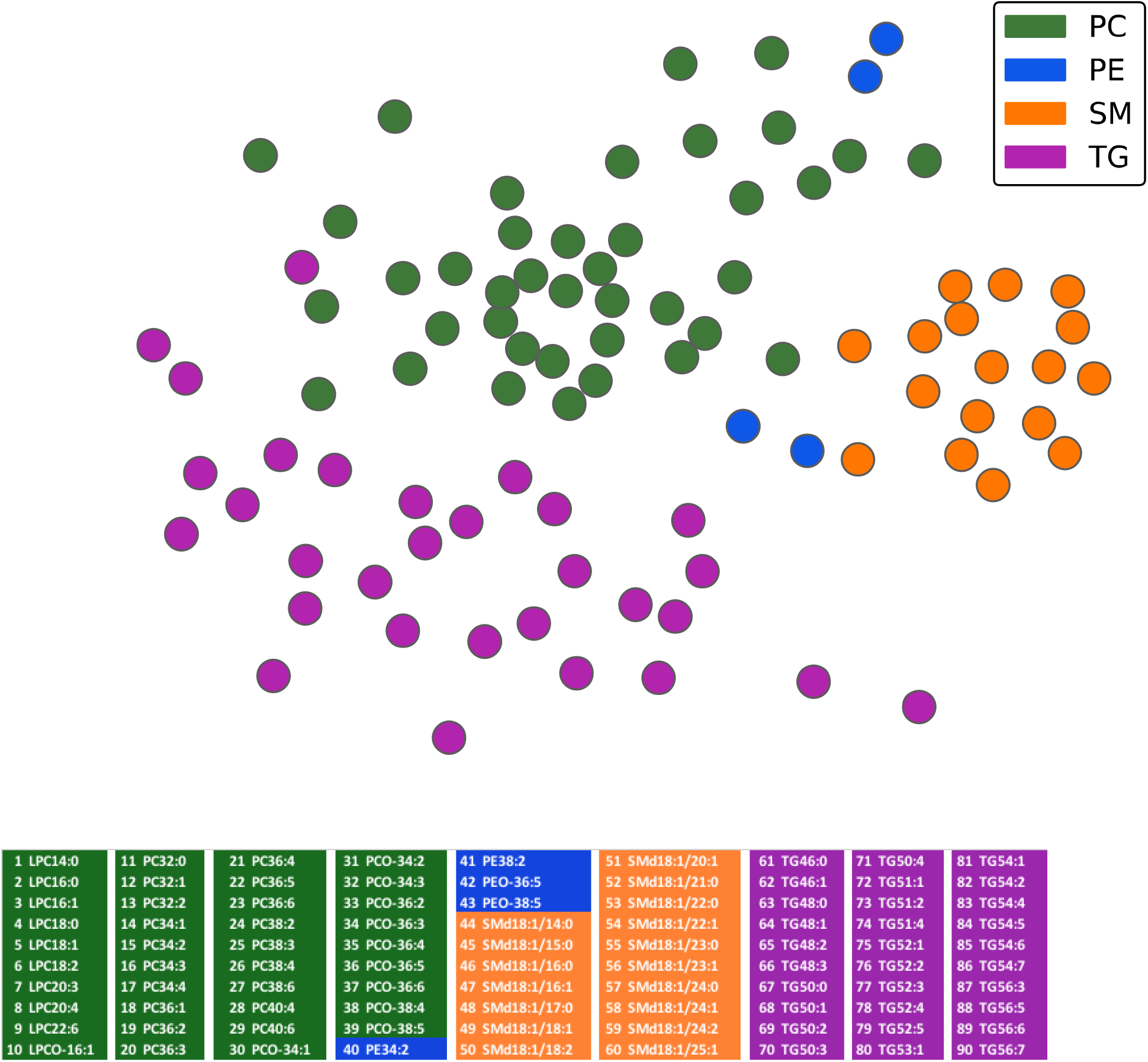
Gaussian graphical model of the plasma lipidome. Figure shows the clustering of individiual lipids captured by the LC-MS platform, in the participants of the Netherland Twin Register.

### GWAS meta-analysis results

The meta-analysis yielded 3521 metabolite/SNP associations passing the predefined metabolome-wide significance threshold (P-value < 1.62 × 10^−9^). These signals were coming from 10 distinct genomic loci (*GCKR, APOA, FADS1, SGPP1,TMEM229B, LIPC, PDXDC1, CETP, CERS4* and *SPTLC3*) shown in Table 2. All of the 10 metabolome-wide significant loci have been previously identified in other studies ^3,5,6,11,42^. In the current study, we show evidence for new lipid traits of which the levels in the circulation were also determined by these loci: i.e we demonstrate significant evidence for the involvement of *GCKR* in TG50:2, *FADS1* in LPCO16:1, PC38:2, PE.O36:5, PE.O38:5, TG56:5, TG56:6 and TG56:7, *APOA1* in TG54:4, *SGPP1* in SM21:0, *TMEM229B* in LPC.O16:1 and *SPTLC3* in SM21:0 and SM25:1. Emerging associations show that various TGs in the circulation are driven by *GCKR, FADS1* and *APOA* but the genes drive different subspecies of TGs. *GCKR* drives TG50:2 which by itself makes up 8% of the TG pool. *APOA1* similarly associates with TG54:4, TG52:3 and TG52:4 effecting up to 35% of the TG pool. On the other hand, *FADS1* associates with TG56:5, TG56:6, and TG56:7 which are relatively underrepresented in the TG pool, summing up to only 3.3 % of the total TG concentration. *FADS1* futher determines a wide range of LPCs, PCs and PEs. *CETP* and *TMEM229B* drive different subspecies of ether phospholipids; *CETP* associates with PCO34:1 and *TMEM229* associates with PCO36:5 and LPCO16:1. *SGPP1*, *CERS4* and *SPTLC3* drive circulating sphingomyelin levels, but the lipids do not overlap across the loci except SM21:0, which is shared between *SPTLC3* and *SGPP1*.

**Table 2.** Top significant LipQTL and their associated lipids which pass the metabolome–wide significance threshold.

| Lipids | SNP | Chr | Position | Locus | #SNPs | NEA/EA | EAF | $\beta$ | SE$_{\beta}$ | P-value |
| --- | --- | --- | --- | --- | --- | --- | --- | --- | --- | --- |
| **TG50.2** shown in the current study | rs11127048 | 2 | 27598097 | *GCKR* | 1 | A/G | 0.63 | -0.13 | 0.02 | $1.27 \times 10^{-9}$ |
| LPC.O16.1, PC38.2, PE.O36.5, PE.O38.5, TG56.5, TG56.6, and TG56.7 shown in the current study; PC.O34.2, PC.O36.2, PC.O36.3, PC.O36.4, PC.O36.5, PC.O38.4, PC.O38.5, LPC20.3, **LPC20.4**, PC32.2, PC34.2, PC34.3, PC34.4, PC36.2, PC36.3, PC36.4, PC36.5, PC38.3, PC38.4, PC40.4, PE34.2, and PE38.2 shown previously<sup>1,2</sup> | rs174547 | 11 | 61570783 | *FADS1* | 228 | T/C | 0.67 | 0.44 | 0.02 | $1.02 \times 10^{-107}$ |
| TG54.4 shown in the current study; **TG52.3**, TG52.4 shown previously<sup>3</sup> | rs12366015 | 11 | 116990851 | *APOA* | 2 | G/A | 0.82 | -0.1 | 0.02 | $2.60 \times 10^{-10}$ |
| SM21.0 shown in the current study; **SM14.0** and SM15.0 shown previously<sup>1</sup> | rs12878001 | 14 | 64239629 | *SGPP1* | 242 | T/G | 0.86 | -0.35 | 0.03 | $8.51 \times 10^{-32}$ |
| LPC.O16.1 shown in the current study, **PC.O36.5** shown previously<sup>1,2</sup> | rs10873201 | 14 | 67966599 | *TMEM229B* | 49 | C/T | 0.52 | 0.19 | 0.02 | $2.08 \times 10^{-21}$ |
| **PE34.2** shown previously<sup>1</sup> | rs10468017 | 15 | 58678512 | *LIPC* | 114 | C/T | 0.68 | -0.26 | 0.02 | $3.26 \times 10^{-33}$ |
| **LPC20.3**, PC38.3 shown previously<sup>1</sup> | rs12928099 | 16 | 15150505 | *PDXDC1* | 54 | C/A | 0.67 | -0.19 | 0.02 | $6.71 \times 10^{-16}$ |
| **PC.O34.1** shown previously<sup>2</sup>, | rs1532624 | 16 | 57005479 | *CETP* | 6 | C/A | 0.54 | -0.12 | 0.01 | $4.59 \times 10^{-10}$ |
| **SM18.1**, SM20.1 shown previously<sup>1</sup> | rs8105664 | 19 | 8290878 | *CERS4* | 56 | G/A | 0.78 | 0.23 | 0.02 | $3.54 \times 10^{-27}$ |
| SM21.0, **SM25.1** shown in the current study | rs680379 | 20 | 12969400 | *SPTLC3* | 63 | G/A | 0.65 | -0.26 | 0.02 | $9.67 \times 10^{-39}$ |

SNP: lead SNP at each locus. Chr : Chromosome. #SNPs: Number of SNPs that were below the metabolome–wide significance threshold at each locus. NEA/EA: Non-effect allele/effect allele. EAF: Effect allele frequency. $\beta$ : Effect estimate. SE $_{\beta}$ : Standard error of the effect estimate. Previous evidence refers to the same or very closely related phenotypes that have shown a genome –wide significant association with the same locus. ***Italic and bold*** text refer to the strongest association in case multiple phenotypes are associated with the same locus. Novel traits associated with the particular loci have been marked. *SPTLC3* has previously been shown to associate with related traits, i.e. SM22:0 and SM16:1-OH<sup>1</sup>. The following abbreviations are used: triglyceride, TG; acyl-acyl phosphatidylcholine, PC; alkyl-acyl phosphatidylcholine, PCO lysophosphatidylcholine, LPC; sphingomyelin, SM; acyl-acyl phosphatidylethanolamines, PE and alkyl-acyl phosphatidylethanolamines, PEO.

Notably, the common allele (T, major allele frequency = 0.67) of *FADS1* SNP rs174547 leads to an increase (β > 0) in levels of species which typically carry the polyunsaturated fatty acid arachidonic acid (20:4): PC20:4, PC38:4, PC36:4, PCO36:5, PCO38:4, PC36:5, PC34:4, PCO36:4, PC40:4, PCO38:5, PEO38:5, TG56:6, PEO36;5, TG56:5, and TG56:7 and additionally with PCO16.1. On the other hand, the same allele leads to a decrease (β < 0) in levels of species whith lower saturation status: PC34:2, PC38:2, PC36:3, PC36:2, PE34:2, PCO36:2, PE38:2, PC34:3, PC32:2, PCO36:3, and PCO34:2.

Additionally, 867 metabolite /SNP pairs showed suggestive association with 1.61 × 10^−9^ < P < 5 ×10^−8^ and were located in 18 distinct genomic loci. Seven of these suggestively significant signals were coming from already established lipQTLs. These included association of *GCKR* with TG48:2, TG48:3, TG50:3, TG50:4, TG48:1, TG50:1 and TG52:2, *FADS1* with TG56:3 and TG54:6, *APOA1* with TG54:5 and TG56:3*, SGPP1* with SM22:1 *PDXDC1* with PC34:2, *CETP* with PCO34:3 and finally *PKD2L1* with LPC16:1. We set up in-silico replication for the remaining 11 loci using published GWAS summary statistics from earlier lipid QTL studies^5,11^. Three signals were already known loci for cholesterol and triglyceride metabolism from GWAS^3^, i.e. *LDLR* (SM16:0), *MAU2* (PC34:4)^3^ and *MLXIPL* (TG48:1 and TG50:1) (Figure 2). The first locus with supporting evidence is the LDL receptor (*LDLR*), harboring rs11668477. This SNP has a P-value of 0.0002 for association in the GWAS of the EUROSPAN consortium, an earlier report^5^ of the same lipid trait; SM16:0. The second locus is rs73001065 near the *MAU2* gene, which is associated with PC34:4. We did not find rs73001065 in the published earlier GWAS dataset. However, rs3794991, in strong LD with rs73001065 (R^2^=0.81), significantly associated with PC34:4 (P-value = 9.3 ×10 ^−7^) in the EUROSPAN dataset. The EUROSPAN dataset included persons from the ERF population. To avoid bias due to duplicated samples, we excluded the 912 ERF participants that participated in EUROSPAN from the analysis. As a result of this, the P-value for the association between SM16:0 and rs11668477 attenuated to 0.002, whereas the P-value for the association between rs3794991 and PC34:4 became 7.4 ×10 ^−5^. We did not find the third locus (rs10245965 from *MLXIPL*) among the top findings of the study by Rhee et al^11^, indicating that the SNPs in this loci have P > 0.001. For the remaining eight novel lipQTL we failed to find any additional evidence supporting their involvement (replication results are given in **Table S2**. Quantile-quantile plots for the GWAS of the 90 metabolites are displayed in **Figures S2** and Manhattan plots are shown in **Figure S3.** The full GWAS summary statistics have been uploaded to the BBMRI-NL public repository and can be downloaded.

**Figure 2.**
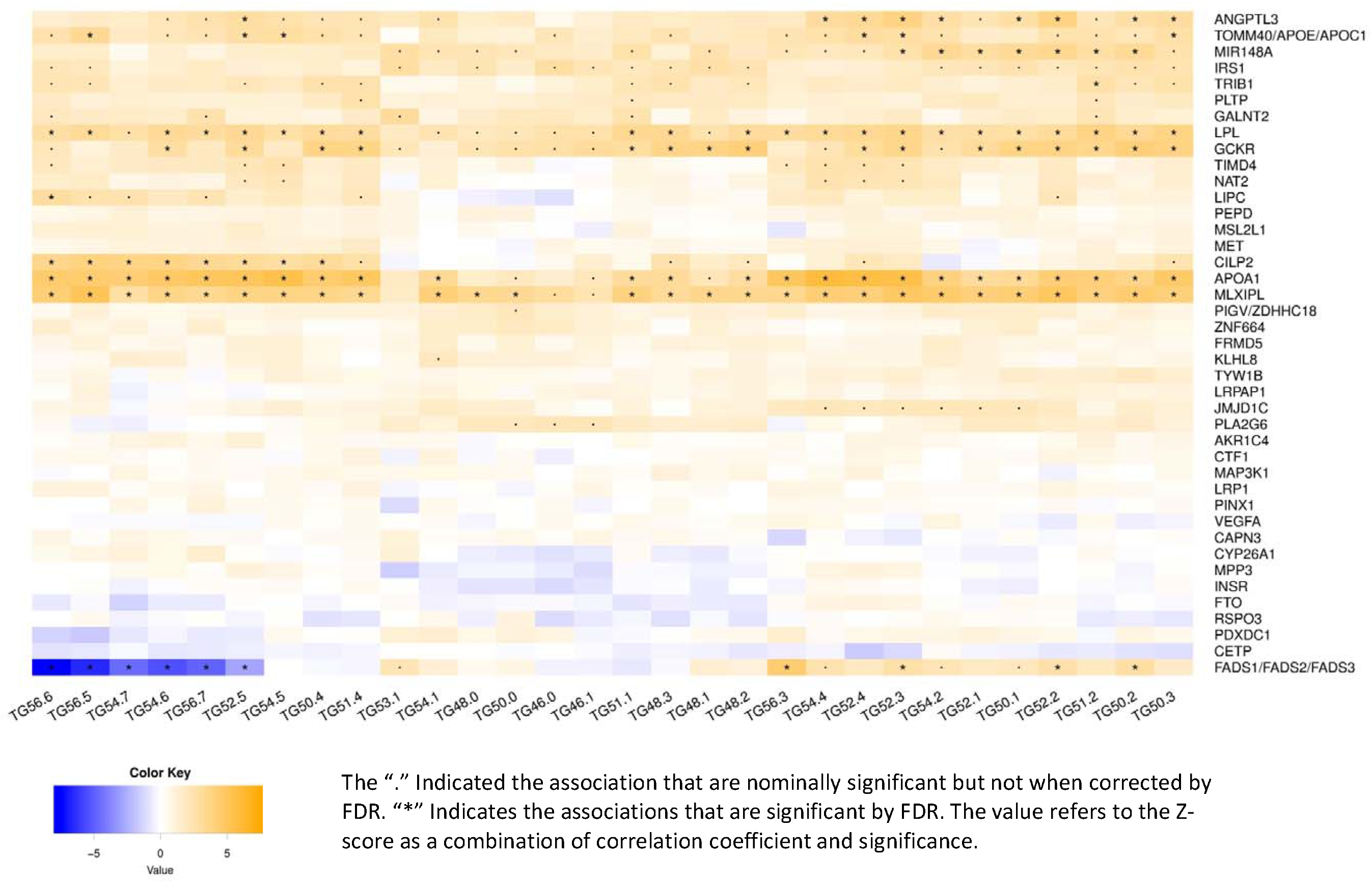
Effect of previously discovered triglyceride loci

### Phenome-wide association study (Phewas)

Top SNPs from lipQTLs *APOA1*, *FADS1*, *LIPC* and *GCKR* are associated with several International Classification of Diseases codes and clinical measurements. rs174547 located at the *FADS1* locus associates with pulse rate, cholelithiasis, asthma, standing height, wine intake and naps during the day. rs12366015 located at the *APOA1* locus associates with hyperlipidemia and lipid lowering medication intake. The *GCKR*-related SNP rs780094, a proxy for rs11127048, associates with 46 UK Biobank phenotypes, including cholelithiasis, height, weight, impedance, diabetes, gout, hypercholesterolemia, daytime dozing, sodium in urine, basal metabolic rate and alcohol intake frequency. The *LIPC*-related SNP, rs10468017, associates with hypercholesterolemia, while the *CETP*-related SNP, rs1532624, and *LIPC* - related SNP, rs10468017, associates with age-related macular degeneration in earlier GWAS. **Table S3** shows the Phewas lookup results.

### Effect of the known TG loci on the individual TG species

Subsequently, we tested whether the 41 loci that have been previously associated with total TGs as primary or secondary trait in the published GLGC study^3^ also associate with the TG subspecies captured by our lipdomics platform. We uncovered 11 loci which, after false discovery rate (FDR) correction, significantly associated (FDR < 0.05) with at least one or more of the TGs (Figure 2). Four loci, i.e. *LPL*, *GCKR*, *APOA1*, and *MLXIPL*, showed a global association to several TGs, at the experiment-wide P-value level. The *LPL* polymorphism associates with 22, *GCKR* with 16, *APOA1* with 24 and *MLXIPL* with 27 out of the 30 studied TGs. Other loci that associate with multiple TGs are *FADS1-2-3* (10), *ANGPTL3* (9), *CILP2* (8), *MIR148* (7) and *TOMM40/APOE/APOC1* (6). The *TIRB1* and *LIPC* loci associated with only one species. Across the known total TG loci, the direction of effect for the individual TG species was consistent, with the exception of the *FADS1-2-3* locus. For the leading SNP rs174546 at the *FADS1* locus, the common allele, which associates with increased level of total TGs, associates with decreased levels of TGs with high fatty acid polyunsaturation number (e.g. TG56:6) and increased levels of TGs with low polyunsaturation number (e.g. TG52:2). rs174546 did not associate with saturated TGs.

### Genetic correlations with cardio-metabolic health

The SNP-based genetic correlations between the 90 lipids and 21 measurements focusing on CVD-related traits are depicted in Figure 3. As expected, we observed a strong positive correlation between all the different TG species and total TG. Total TG also positively correlates with phospholipid species of PC38:3, PE34:2, PC36:3, PC16:0, PC36:4 PC40:4, PC38:2, PC18:0, PC34:3, PC34:1, PC36:1, PC36:2, PC34:2, and PC40:6. The genetic correlations with total TG were negative for the ether phosphatidylcholines; PCO34:3, PCO36:3, PCO34:1 and PC36:2, as well as the sphingomyelin; SM15:0. As expected, the majority of TG species also show a negative correlation with HDL-C and positive correlations with LDL-C. In addition, both LDL-C and HDL-C significantly correlate with a cluster of sphingomyelins (SM18:1, SM16:1, SM20:1, SM23:1, SM15:0, SM18:2, SM24:1, SM16:0, SM14:0) and ether phospholipids (PCO34:3, PCO36:3, PCO36:2, PCO16:1, PCO38:5, PCO36:5, PEO38:5), a well as PC32:0, whereas eight phospholipids specifically share a genetic background with LDL-C (PCO36:4, PCO38:4, PEO36:5, SM25:1, SM21:0, SM23:0,SM24:0 and SM22:0) and three to HDL-C (PCO34:1, PCO34:2 and PCO36:6). One remarkable observation is the significant genetic correlation with two of the LDL-C specific lipids and CVD; (P-value = 8.3 × 10^−6^ for SM22:0 and P-value = 0.03 for PCO38:4). SM22:0 also shares a genetic background with IMT, however this was not significant after FDR correction. There was no statistical evidence for genetic correlation of any of the lipids studies with either heart rate, blood pressure, or WHR. Sixteen of the TG species, as well as PC38:3, correlate positively with total body fat percentage, whereas PCO34:3 and PCO36:3 correlate negatively. PC38:3, SM18:1 and SM16:1 correlate positively with BMI as well, whereas PC18:2 show a negative genetic correlation to BMI.

**Figure 3.**
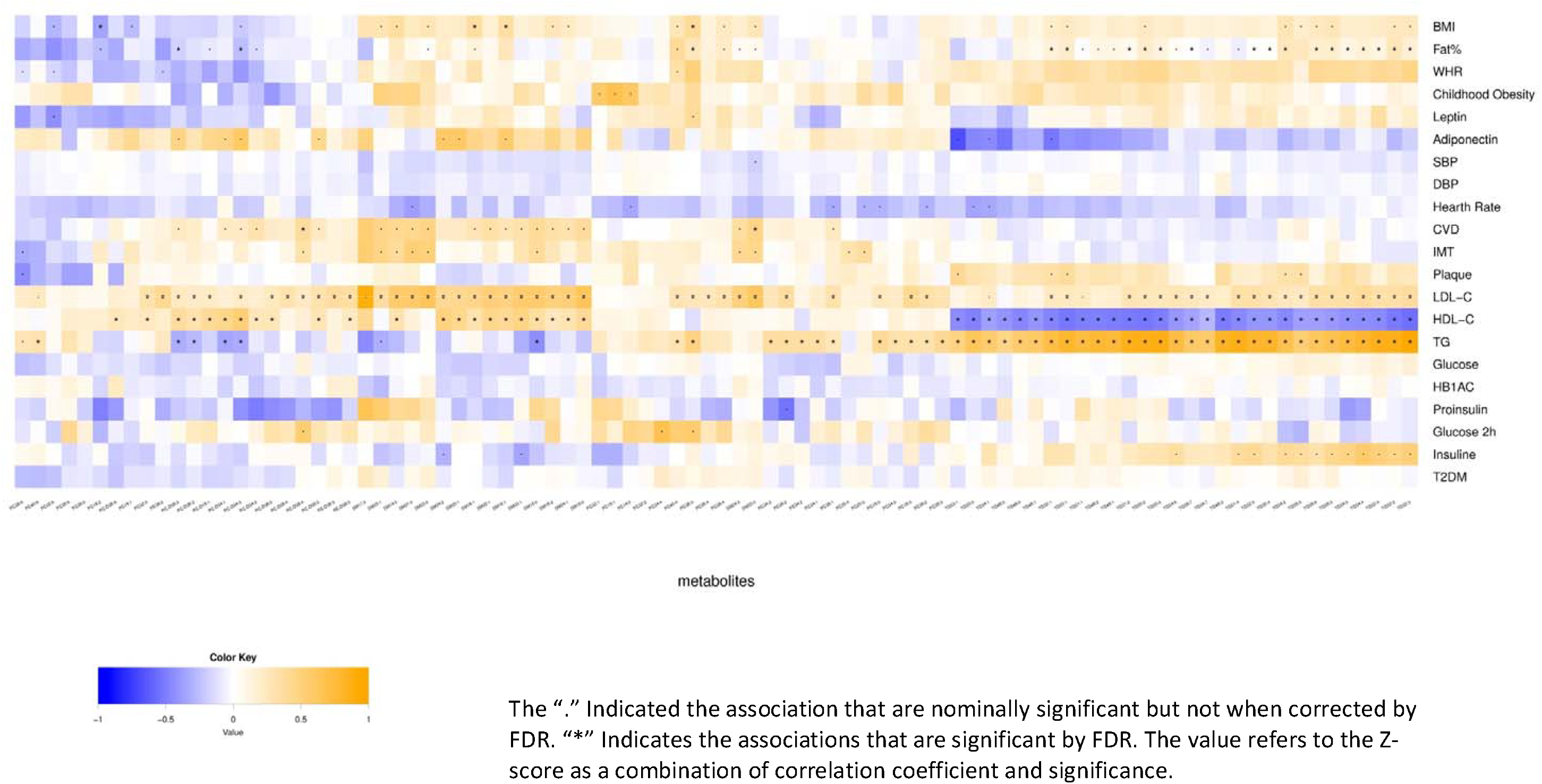
Genetic correlation between the circulating lipids and metabolic health-related phenotypes

### Lipids and risk of CVD

In the dataset of the ERF population we tested whether SM22:0 and PCO38:4 have a phenotypic correlation to carotid intima media thickness (IMT) and whether they predict CVD diagnosis. We estimated that one unit increase in plasma SM22:0 associates with an increase in IMT (Beta = 0.031, P = 0.023) independent of age, sex, BMI, blood pressure, LDL-C and lipid lowering medication therapy. However the level of SM22:0 did not associate to the risk of CVD in 15 year follow –up. We found no evidence supporting a role of PCO38:4 in IMT nor CVD.

## Discussion

In the current study, we focused on 30 different TG and 60 phospholipids species measured by mass spectrometry in 3 Dutch population-based cohorts. Overall, there was high clustering of the lipid species within their overall chemical groups, TGs, phospholipids and sphingolipids, suggesting distinct molecular pathways responsible for each group. The GWAS meta-analysis replicated 10 previously identified major associations for lipid species including the major determinants such as *GCKR, APOA, FADS1, SGPP1,TMEM229B, LIPC, PDXDC1, CETP, CERS4* and *SPTLC3* and reported eleven novel lipids associated with one or more of these loci. In addition, by screening across the suggestively significant hits we identified three novel loci; *MAU* (in close proximity with the earlier identified TG locus CILP2) for PC34:4, *MLXIPL* for TG48:1 and TG50:1 and *LDLR* for SM16:0. The lipidomics data included 30 TG species with different relative abundances in the TG pool of the plasma, two of the main components, TG52:2 and TG52:3, make up some 43 % of the circulating TGs we detected, wheres the remaining 57% was divided among 28 species. We further tested whether distinct TG species were associated with genes that were identified in the total TG GWASs. We showed that the genes which determine the total TG levels mostly have an overall effect on individual TGs, whereas some are species specific. Of the 41 genetic variants we tested, the *CILP2* locus showed obvious different effect sizes across different species and is mainly involved in the unsaturated TGs, whereas the *FADS1-2-3* locus regulated the TGs in a fatty acid saturation level specific manner. Finally, we identified genetic correlations between particular lipids and determinants of CVD: a total of eight sphingomyelines and ether phospholipids genetically correlated with LDL-C, and three ether lipids correlated with HDL-C. SM22:0 shares a genetic background with both IMT and CVD. Our study replicates previously identified genes implicated in plasma TG and phospholipid levels including *GCKR, APOA1, FADS1, SGPP1,TMEM229B, LIPC, PDXDC1, CETP, CERS4* and *SPTLC3^5,6^*. Among them *GCKR, APOA1, FADS1, LIPC* and *CETP* have been under investigation for their effects on metabolic disorders. Look-ups in UK Biobank datasets and earlier sources indicate various outcomes related to *FADS1* and *GCKR*, whereas *APOA1, CETP* and *LIPC* are more restcricted to dyslipidemia. Here, we provide potential mediators for these genes through which their global effects could be manifested. We also identified a series of SNPs that are borderline genome-wide significant but located in interesting candidate genes. Among the newly discovered phospholipid loci is the *MAU2* gene, which codes for the Cohesin Loading Complex Subunit SCC4 Homolog protein that is involved in rare neuronal disease Cornelia de Lange syndrome. The top SNP itself in this locus, rs73001065, has a strong *cis*-effect on the expression of *GATAD2A* gene (P-value = 5.6 × 10^−33^ in Westra et al.^43^), which is a nuclear protein involved in transcriptional repression. We were not able to link the *GATAD2A* molecular function to lipid metabolism directly. The second locus of interest was the *LDLR* locus involved in SM16:0. This variant has been previously identified as a strong determinant of LDL-C, total cholesterol, as well as for waist-to hip ratio. The third locus, *MLXIP*, is a major determinant of total TG level. We have not found evidence for this locus in the previous report of Rhee et al. However, this could be due to lack of power as the effect sizes are particularly small.

The TG pool in the circulation does not consist of a homogenous set of individual TG species but is dominated by six species of TGs: TG52:2, TG52:3, TG50:2, TG52:4, TG50:1 and TG54:4. These show high similarity in terms of their top genetic determinants, clustering on seven different loci: *ANGPTL3*, *APOE*, *MIR148A, LPL*, *GCKR*, *APOA1*, *MLXLPL* and *FADS*. Except the *FADS* genetic variant, the effects of these loci are also similar for the rest of the TGs. From our results, the *FADS* locus determines the difference between two clusters of TGs, namely saturated ones and polyunsaturated ones. It is of note that the effects of *FADS* were as expected in opposite direction between these two clusters. In addition, the *CILP2* SNP associates only with TG56:6 (1.1 %), TG60:8 (0.5 %), TG54:7 (0.3%), TG54:6 (1.6), TG 56:7 (1.3 %), TG52:5 (1.5 %). TG54:5 (3.5 %) and TG50:4 (0.6 %), which constitute the less abundant group of TGs.

The genetic correlation matrix confirmed known associations and also uncovered novel associations in line with what has been previously reported for the plasma lipidome. We identified six lipids (four ether PCs and a SM) which correlated negatively with total TGs. From these, PCO36:3 and PCO34:1 also negatively correlated to adiposity measured as fat percentage. Sphingolipids are surface components of serum lipoproteins and are abundant in LDL followed by VLDL > HDL^44^. In line with this, genetic correlation analysis suggest to classify thee lipids in two categories; a group of molecules that genetically correlate with both LDL-C and HDL-C, and a second group of molecules which only correlate specifically with LDL-C. Of interest is also our finding that SM22:0 shares a common genetic background with CVD and atherosclerosis as measured by IMT. Sphingomyelins in general were previously suggested to be involved in atherosclerosis^45^. However, to date, it has not been possible to pin-point a particular molecule. SMs may influence atherosclerosis, either directly or by affecting other risk factors such as cholesterol. It has been shown that sphingomyelin levels affect LDL binding and internalization^46^. Hydrolysis of LDL-SM by an extracellular sphingomyelinase in atherosclerotic lesions alters the aggregation state of the particle and promotes foam cell formation by macrophages^47–49^ The fact that SM 22:0 correlates with IMT as a measure of atherosclerosis independent of other risk factors, including LDL-C, suggests it could be a new molecular candidate for further research for prevention and treatment of CVD.

Focusing on 90 lipid species of the plasma lipidome and a Dutch population specific genotype imputation panel in 5537 samples, we show a total of 9 new locus-lipid associations which were replicated via *in-silico* look-ups. We confirmed previously identified associations and suggest new phenotypes for known loci. Moreover, we identified a new LDL-C and CVD specific lipid; SM 22:0, which is associated with IMT and is a potential target for prevention of CVD. Our findings yield higher resolution of plasma lipid species and provide new insights in the biology of circulating phosholipids and their relation to CVD risk.

