## supplementary text for "Genome-wide association study of plasma triglycerides, phospholipids and relation to cardio-metabolic risk factors"

**Cohort information, acknowledgements and funding**

*Erasmus Rucphen Family study (ERF)* The Erasmus Rucphen Family genetic isolate study (ERF) is a prospective family -based study located in Southwest of the Netherlands. This young genetic isolate was founded in the mid-eighteenth century and minimal immigration and marriages occurred between surrounding settlements due to social and religious reasons. The ERF study population includes 3,465 individuals that are living descendants of 22 couples with at least six children baptized. The study protocol was approved by the medical ethics board of the Erasmus Medical Center Rotterdam, the Netherlands. The baseline demographic data and measurements of the ERF participants were collected around 2002 to 2006. All the participants filled out questionnaires on socio-demographics, disease and medical history and lifestyle factors, and were invited to the research center for an interview and blood collection for biochemistry and physical examinations including blood pressure and anthropometric measurements have been performed. The participants were asked to bring all their current medications for registration during the interview. Venous blood samples were collected after at least eight hours fasting. ERF was supported by the Consortium for Systems Biology (NCSB), both within the framework of the Netherlands Genomics Initiative (NGI)/Netherlands Organization for Scientific Research (NWO). ERF study as a part of EUROSPAN (European Special Populations Research Network) was supported by European Commission FP6 STRP grant number 018947 (LSHG-CT-2006-01947) and also received funding from the European Community's Seventh Framework Program (FP7/2007-2013)/grant agreement HEALTH-F4-2007-201413 by the European Commission under the program “Quality of Life and Management of the Living Resources” of 5th Framework Program (no. QLG2-CT-2002-01254) as well as FP7 project EUROHEADPAIN (nr 602633). High-throughput analysis of the ERF data was supported by joint grant from Netherlands Organization for Scientific Research and the Russian Foundation for Basic Research (NWO-RFBR 047.017.043). High throughput metabolomics measurements of the ERF study has been supported by BBMRI-NL (Biobanking and Biomolecular Resources Research Infrastructure Netherlands). Ayse Demirkan is supported by a Veni grant (2015) from ZonMw. Ayse Demirkan, Jun Liu and Cornelia van Duijn have used exchange grants from PRECEDI. The funders had no role in study design, data collection and analysis, decision to publish, or preparation of the manuscripts. ERF study is grateful to all study participants and their relatives, general practitioners and neurologists for their contributions and to P. Veraart for her help in genealogy, J. Vergeer for the supervision of the laboratory work and P. Snijders for his help in data collection.

***Leiden Longevity Study (LLS)***

In the Leiden Longevity Study (http://www.molepi.nl/research/longevity), nonagenarian sibling pairs were included when aged older than 89 years for men and 91 years for women. Additional inclusion criteria were that participants needed to have at least one sister or a brother fulfilling these age criteria, and who was also willing to participate10. Because proper controls are lacking at very high ages, the offspring of the nonagenarian siblings were asked to be included in the study as well. The partners thereof were included in the study to serve as a control group, representing the general population at an age comparable to the offspring. The total study population, excluding the nonagenarian siblings, consisted of 2,415 participants (1,671 offspring; 744 partners). ^10^The Medical Ethical Committee of the Leiden University Medical Centre approved the study and informed consent was obtained from all participants. Blood samples were taken at baseline for extraction of DNA and the determination of non-fasted serum parameters. For metabolomics measurements, blood was drawn using a safety lock butterfly needle. Within two hours tubes with clotted blood were centrifuged for 15 minutes at 2,800 x g and serum was obtained and divided into subsamples, snap-frozen and stored at –80° C. The LLS has received funding from the European Union's Seventh Framework Programme (FP7/2007-2011) under grant agreement n° 259679. This study was supported by a grant from the Innovation-Oriented Research Program on Genomics (SenterNovem IGE05007), the Centre for Medical Systems Biology, and the Netherlands Consortium for Healthy Ageing (grant 050-060-810), all in the framework of the Netherlands Genomics Initiative, Netherlands Organization for Scientific Research (NWO), Unilever Colworth and by BBMRI-NL, a Research Infrastructure financed by the Dutch government (NWO 184.021.007).

***Netherlands Twin Registry (NTR)***

The Netherlands Twin Register (NTR; http://www.tweelingenregister.org/) recruits twins and their family members to study the causes of individual differences in health, behavior and lifestyle. Participants are followed longitudinally; a subsample of unselected twins and their family members has taken part in the NTR-Biobank project. NTR Research was funded by the Netherlands Organization for Scientific Research (NWO: MagW/ZonMW grants 904-61-090, 985-10-002,904-61-193,480-04-004, 400-05-717, Addiction-31160008 Middelgroot-911-09-032, Spinozapremie 56-464-14192), Center for Medical Systems Biology (CSMB, NWO Genomics), NBIC/BioAssist/RK(2008.024), Biobanking and Biomolecular Resources Research Infrastructure (BBMRI –NL, 184.021.007), the VU University’s Institute for Health and Care Research (EMGO+ ), the European Community's Seventh Framework Program (FP7/2007-2013), ENGAGE (HEALTH-F4-2007-201413) and the European Science Council (ERC - 230374 and ERC-284167). HHMD is supported by an EMGO+ Fellowship as part of the Mental Health research program of the EMGO Institute for Health and Care Research. NTR thanks all participants in the Netherlands Twin Register. This work was carried out on the Dutch national e-infrastructure with the support of SURF Foundation
