## supplementary figures for "Genome-wide association study of plasma triglycerides, phospholipids and relation to cardio-metabolic risk factors"

**Supplementary Figure 1** TG species and their contribution to the circulating TG pool

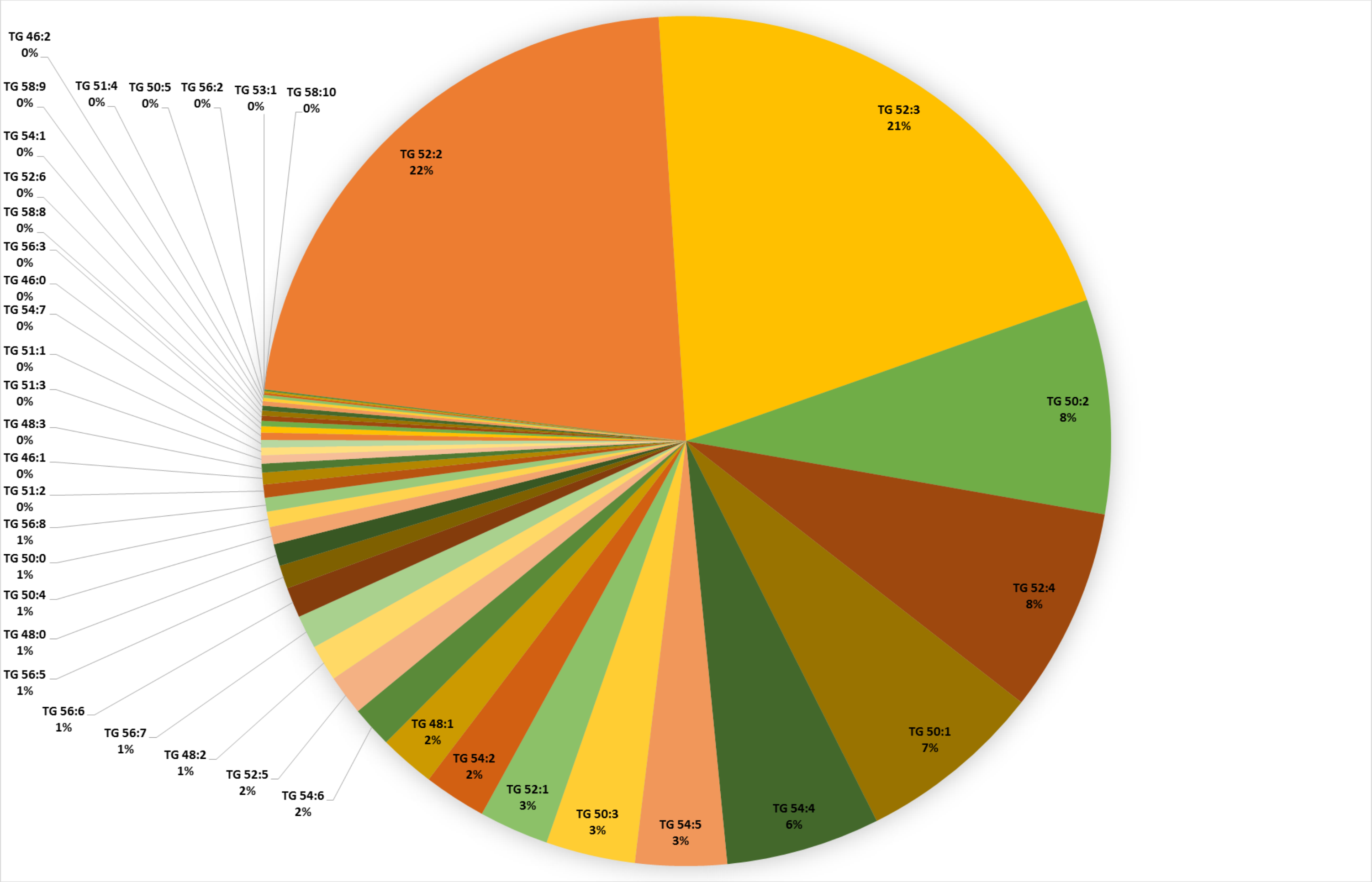

**Supplementary Figure 2** QQ plots of the metabolites studied

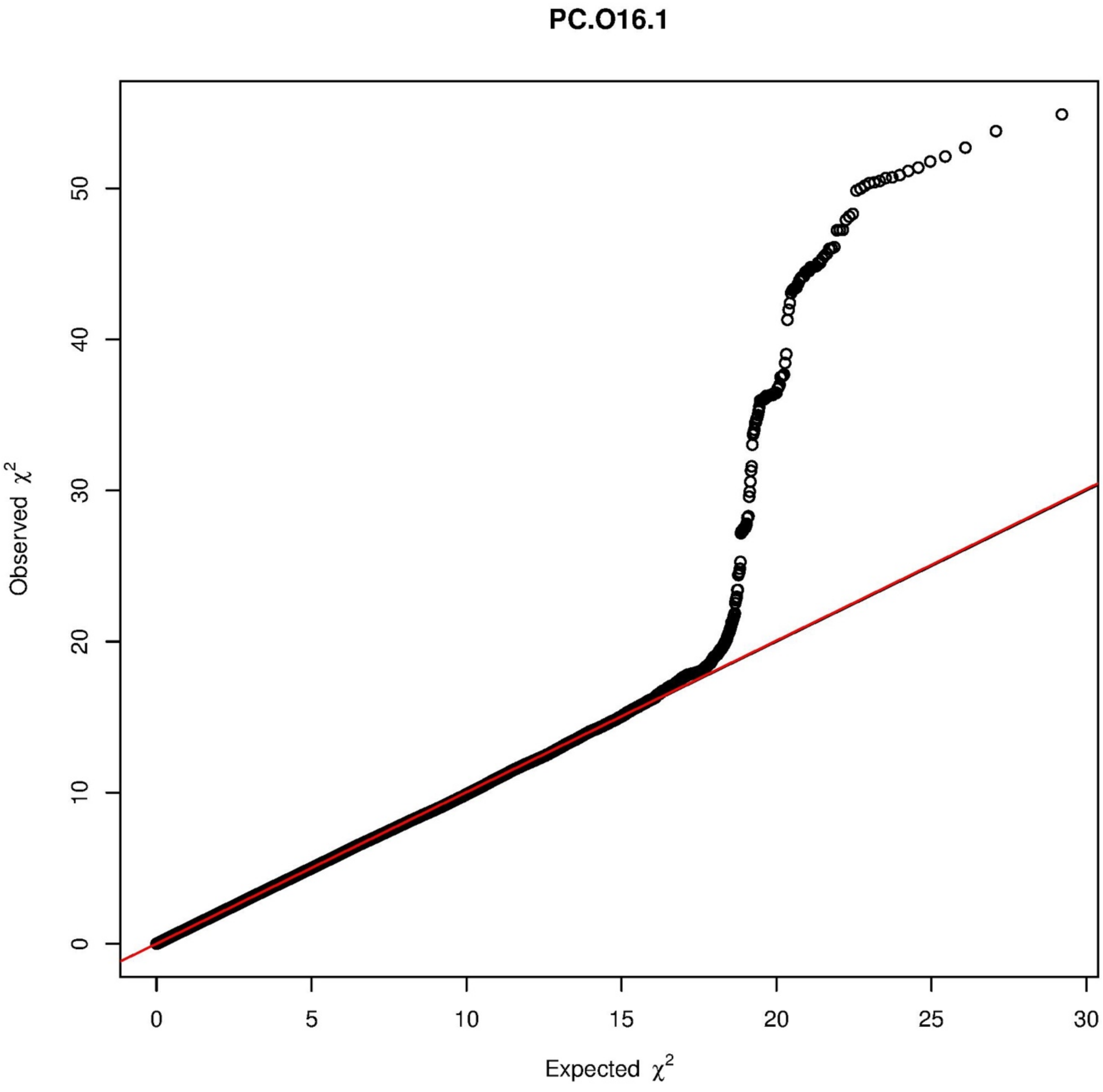

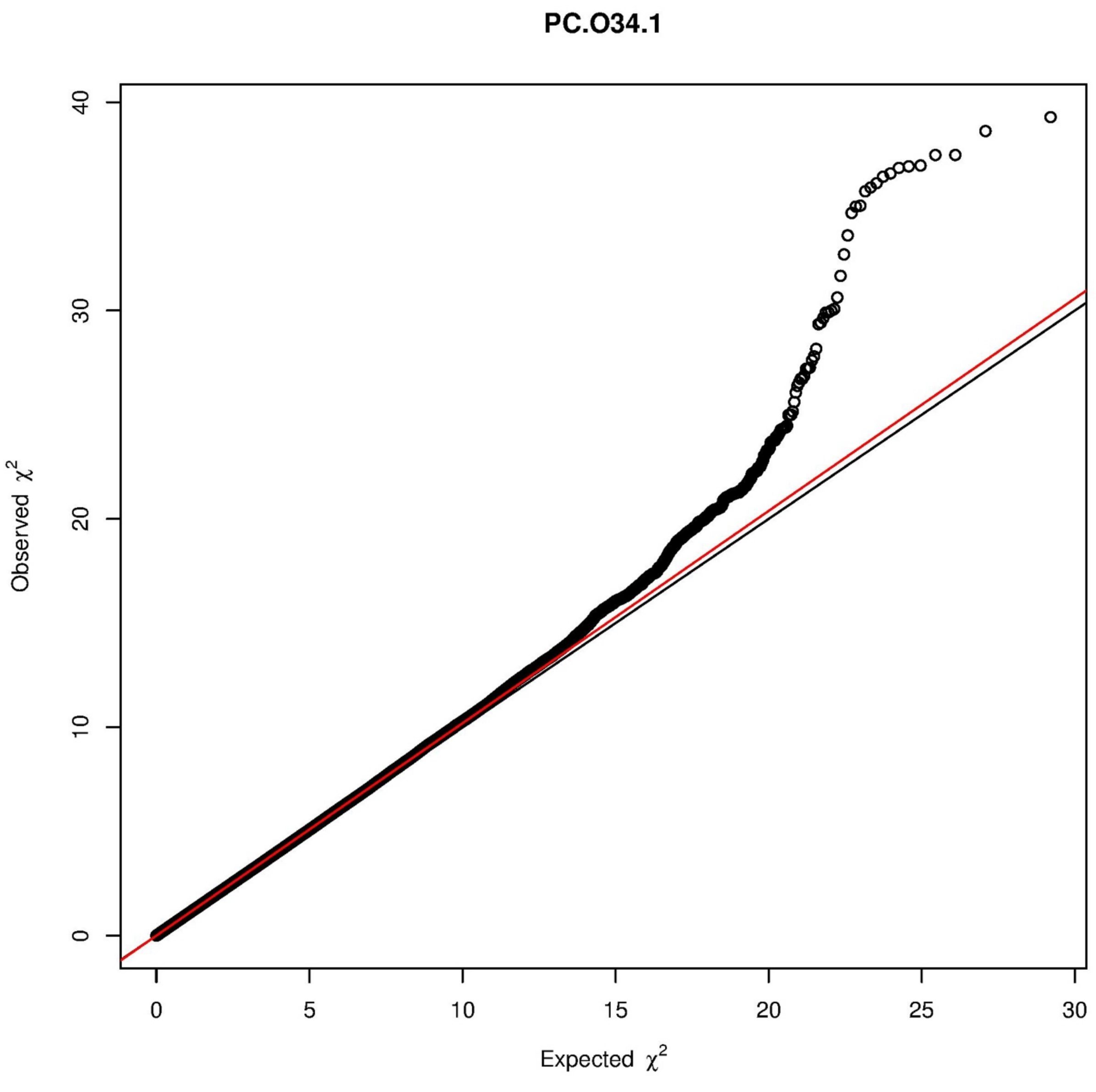
PC.O34.1
Observed $\chi^2$
Expected $\chi^2$
0
10
20
30
40
0
5
10
15
20
25
30

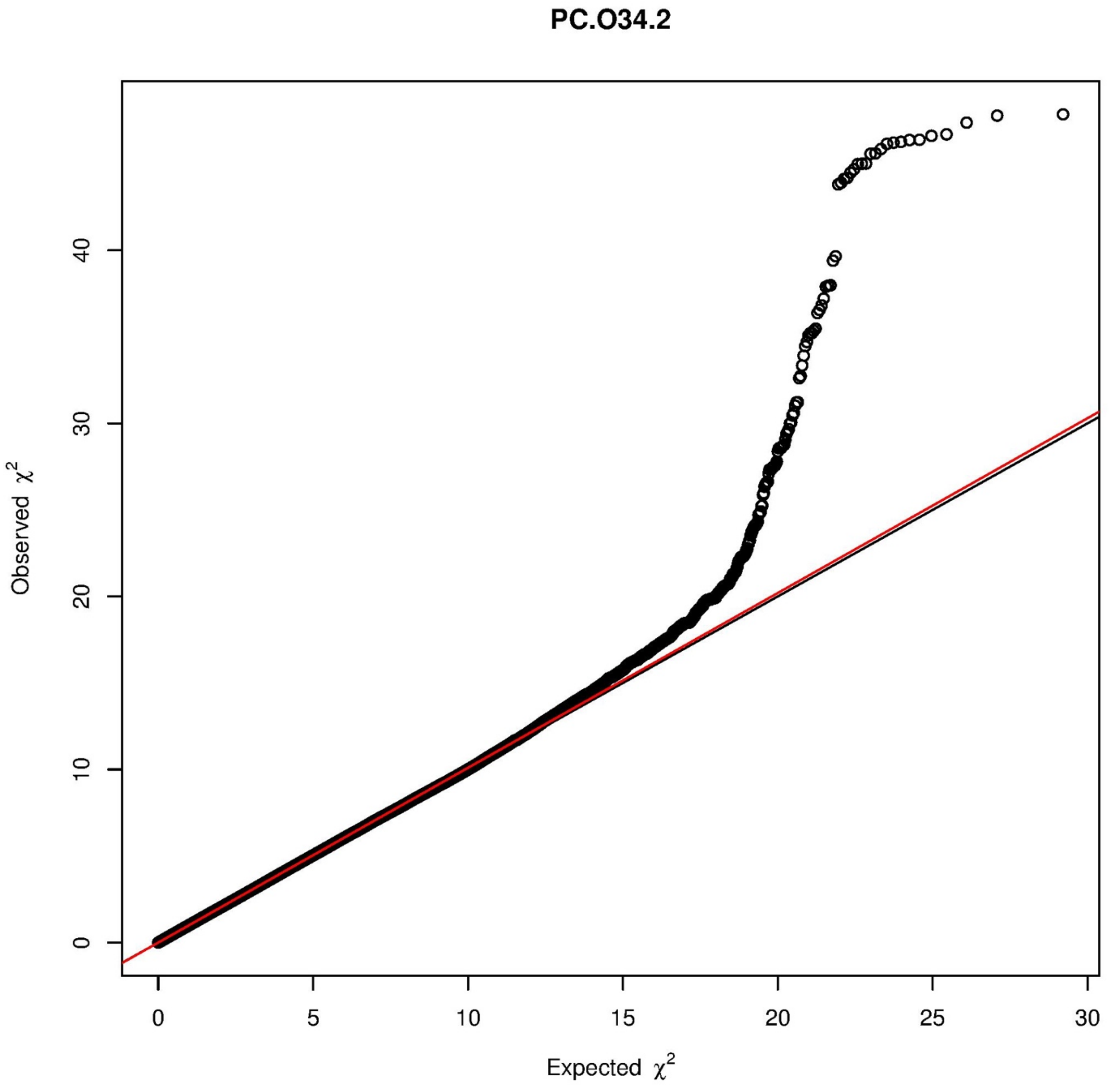
PC.O34.2
Observed $\chi^2$
Expected $\chi^2$
0
10
20
30
40
0
5
10
15
20
25
30

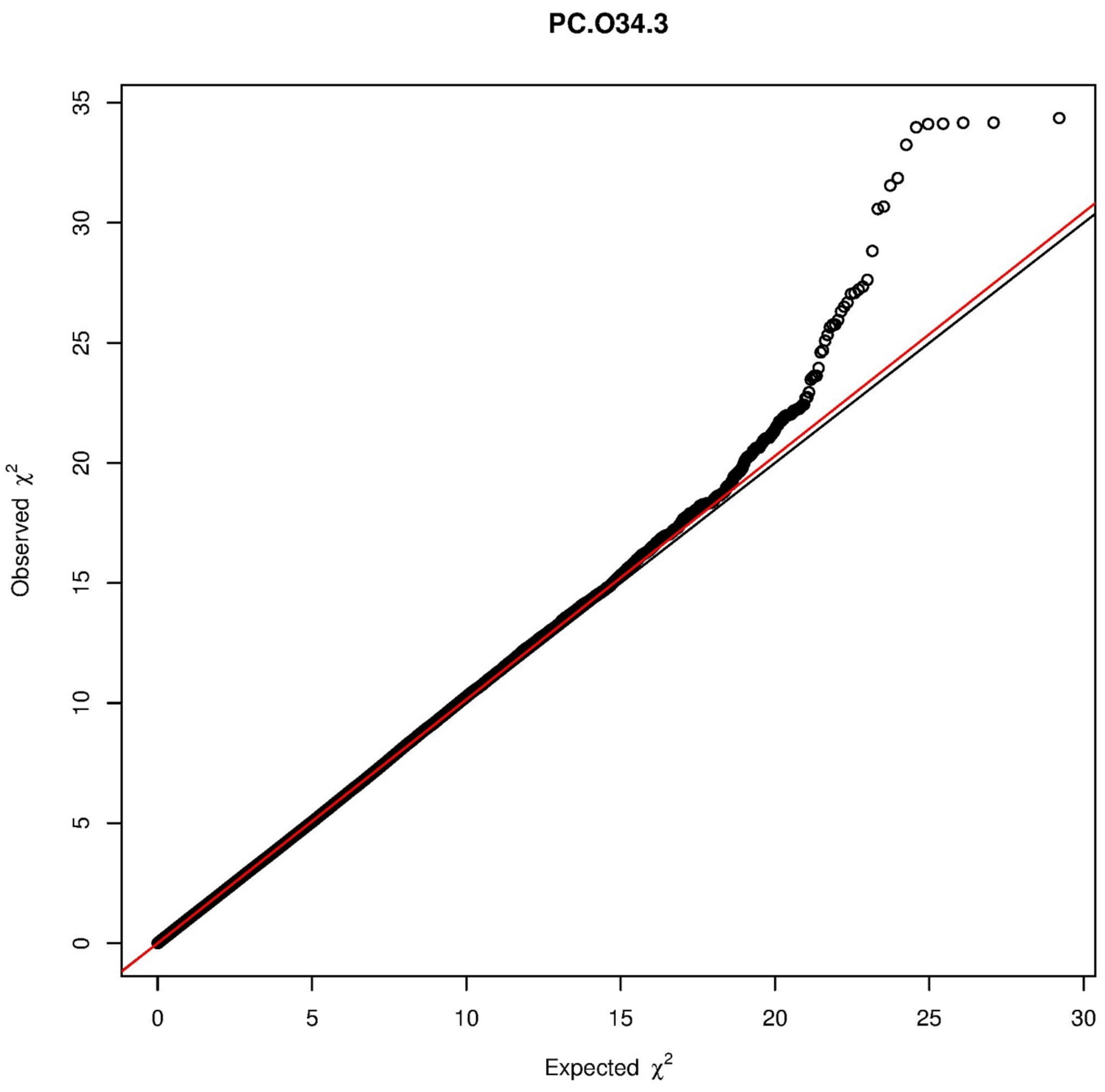

PC.O34.3
Observed $\chi^2$
Expected $\chi^2$
0
5
10
15
20
25
30
35

PC.O36.2

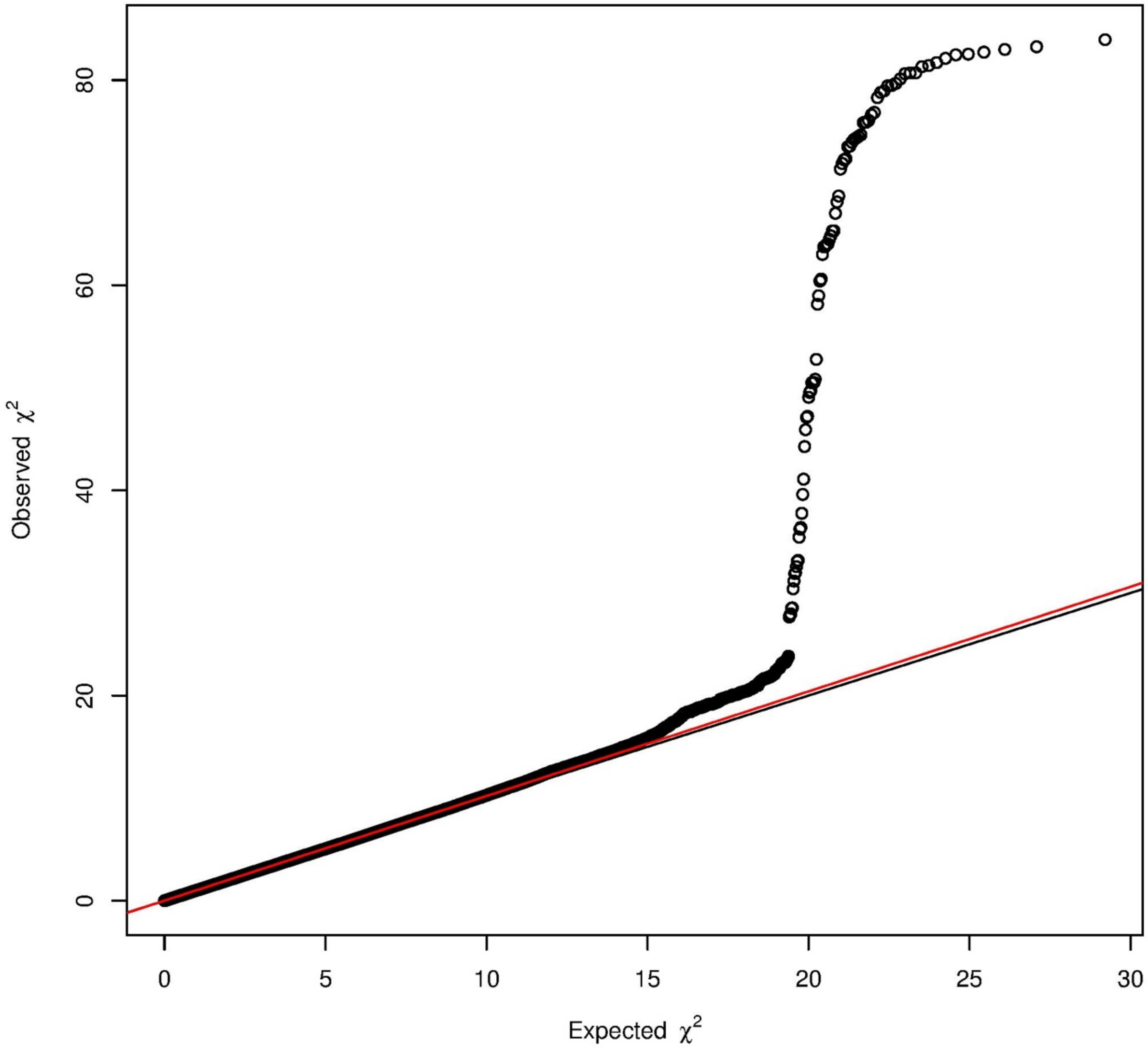

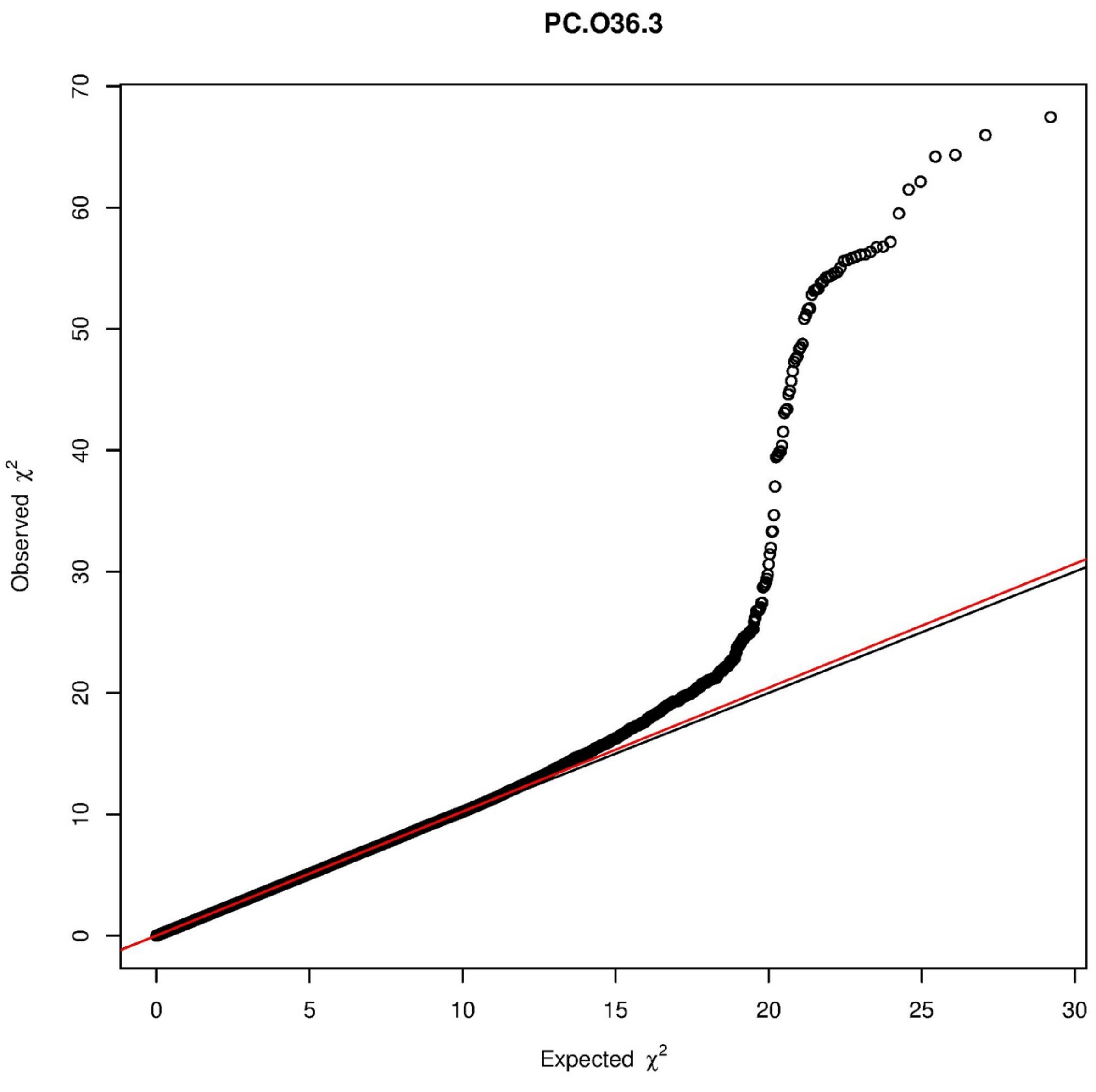
PC.O36.3
Observed $\chi^2$
Expected $\chi^2$
0
10
20
30
40
50
60
70
0
5
10
15
20
25
30

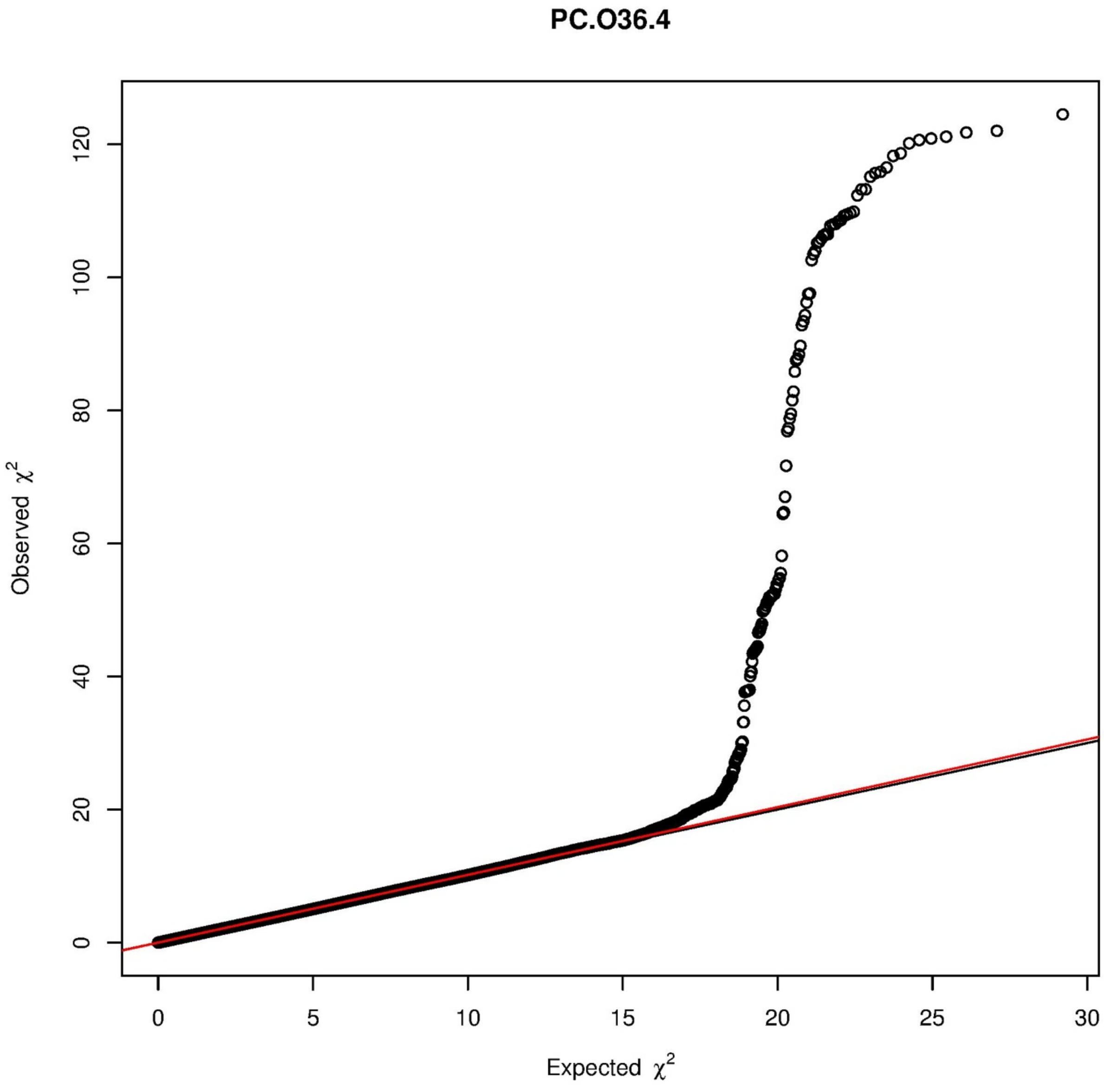
PC.O36.4
Observed χ²
Expected χ²
0
20
40
60
80
100
120
0
5
10
15
20
25
30

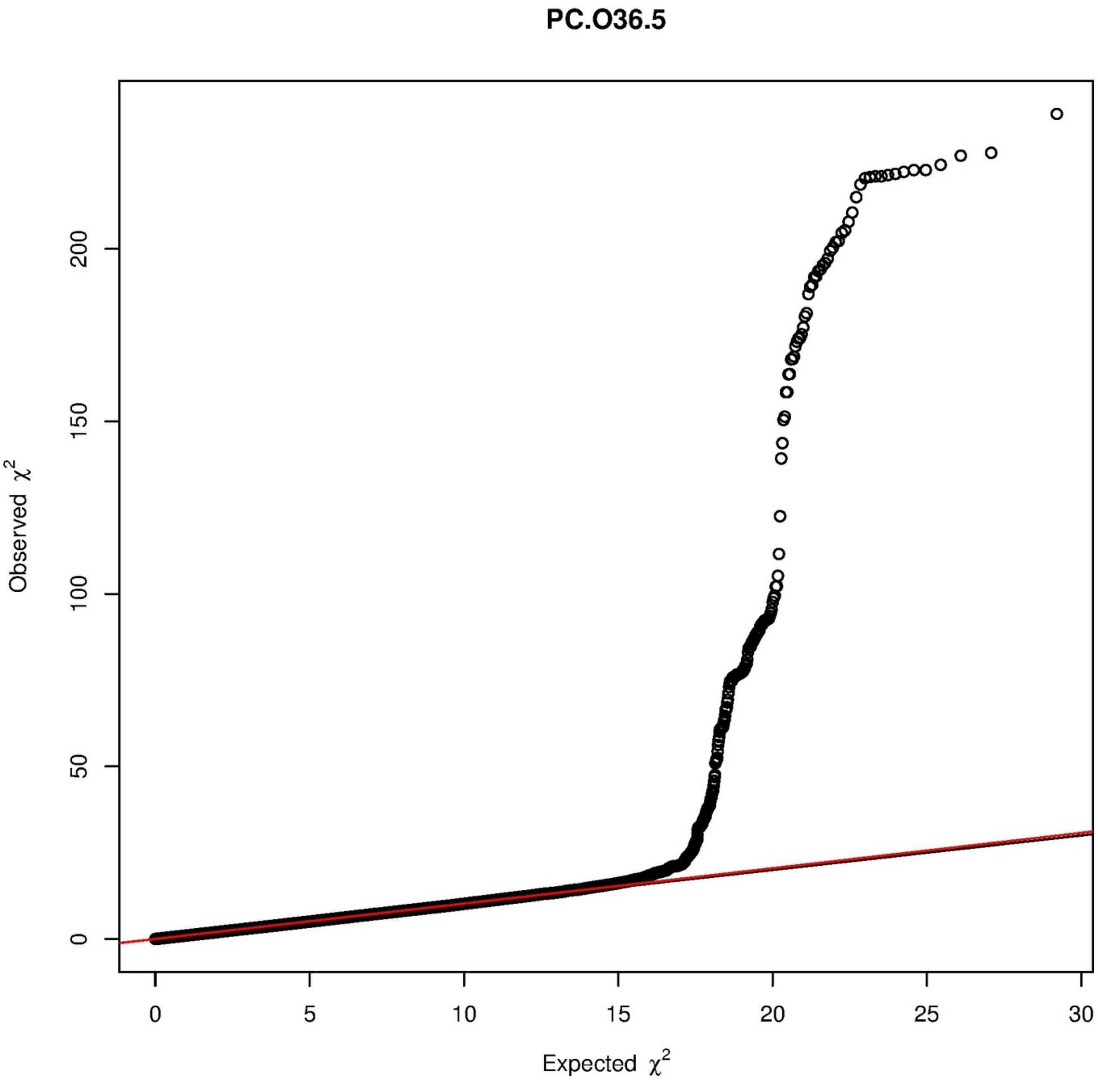

PC.O36.5
Observed $\chi^2$
Expected $\chi^2$
0
50
100
150
200
0
5
10
15
20
25
30

# PC.O36.6

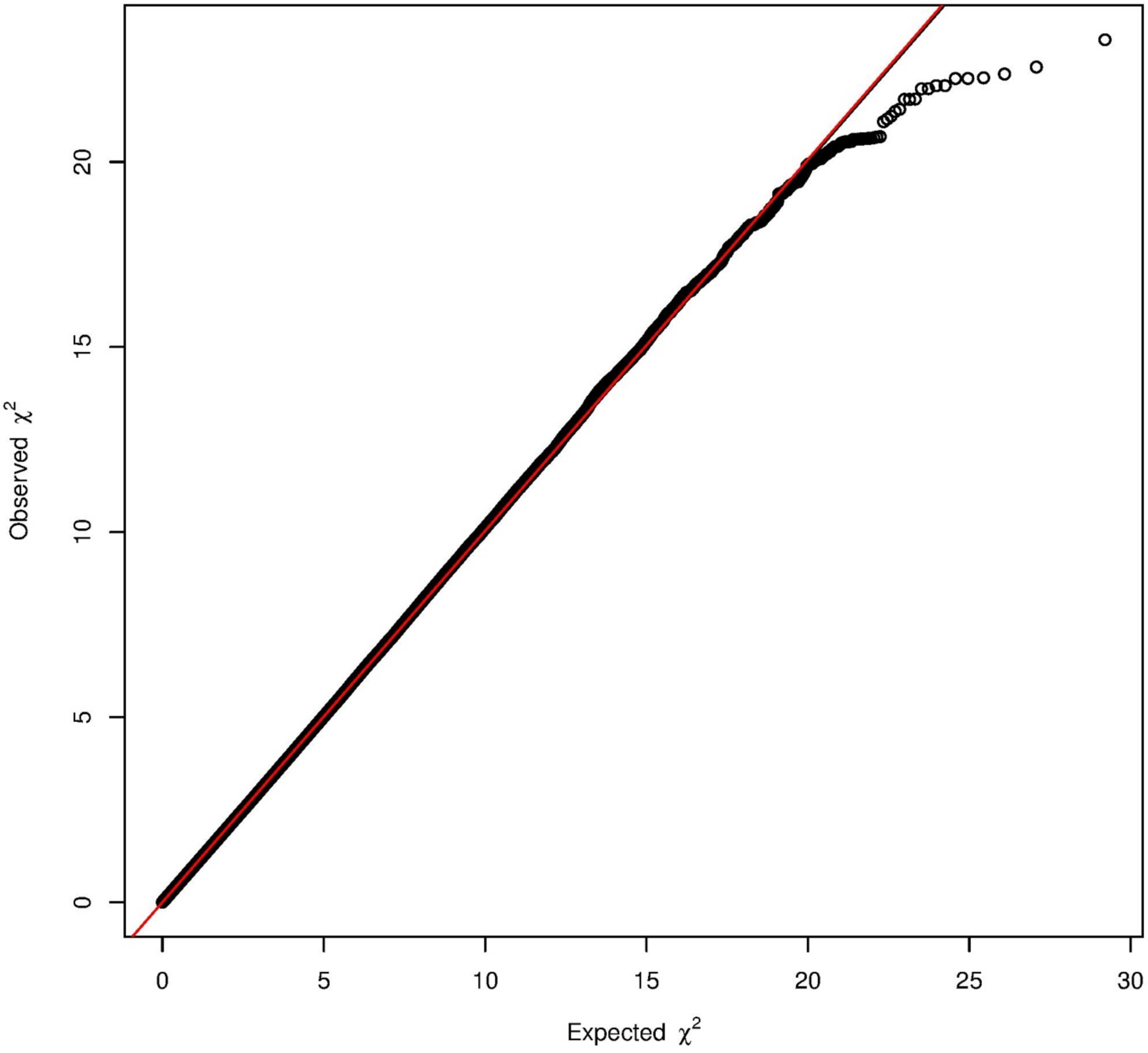

PC.O38.4

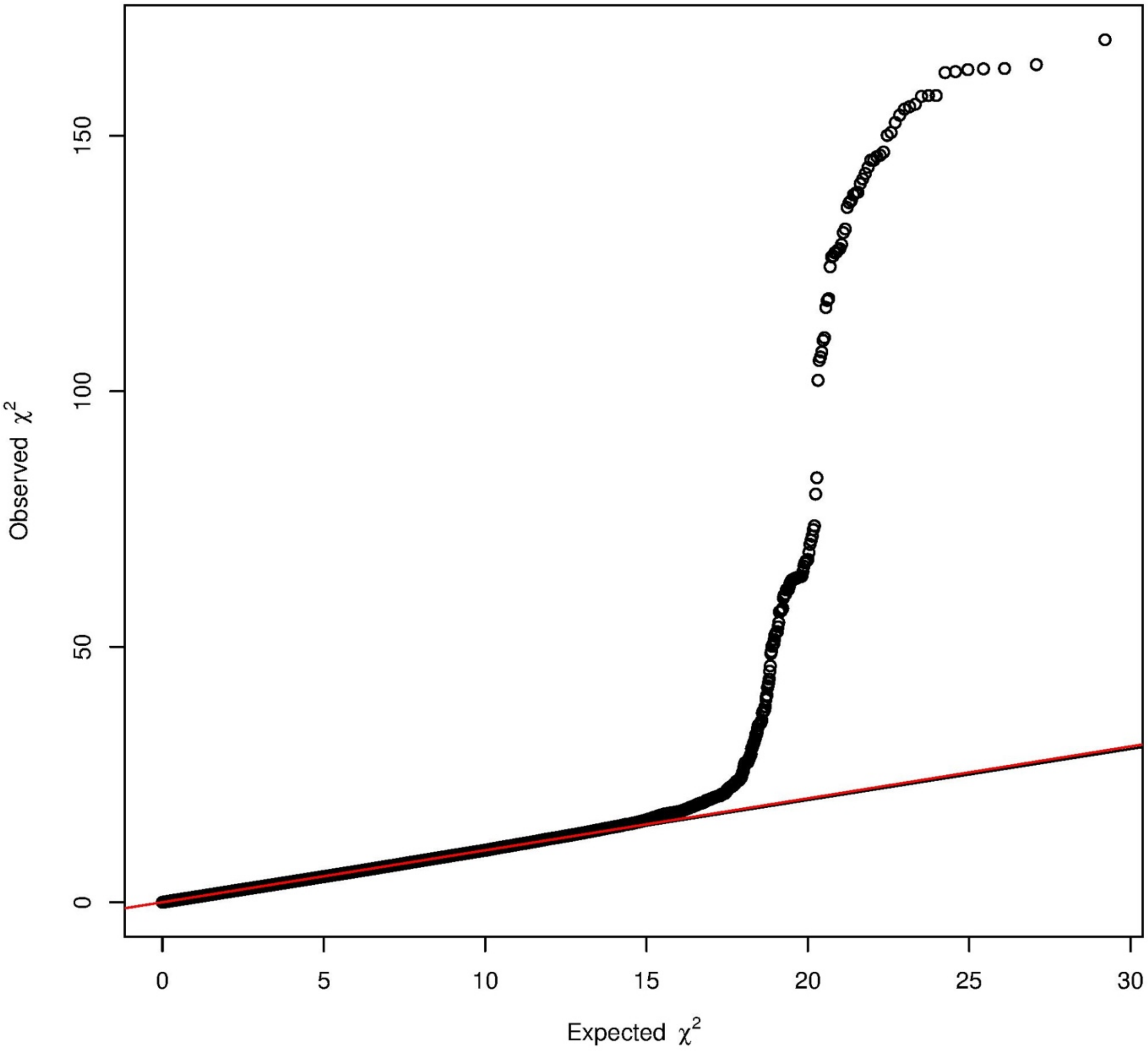

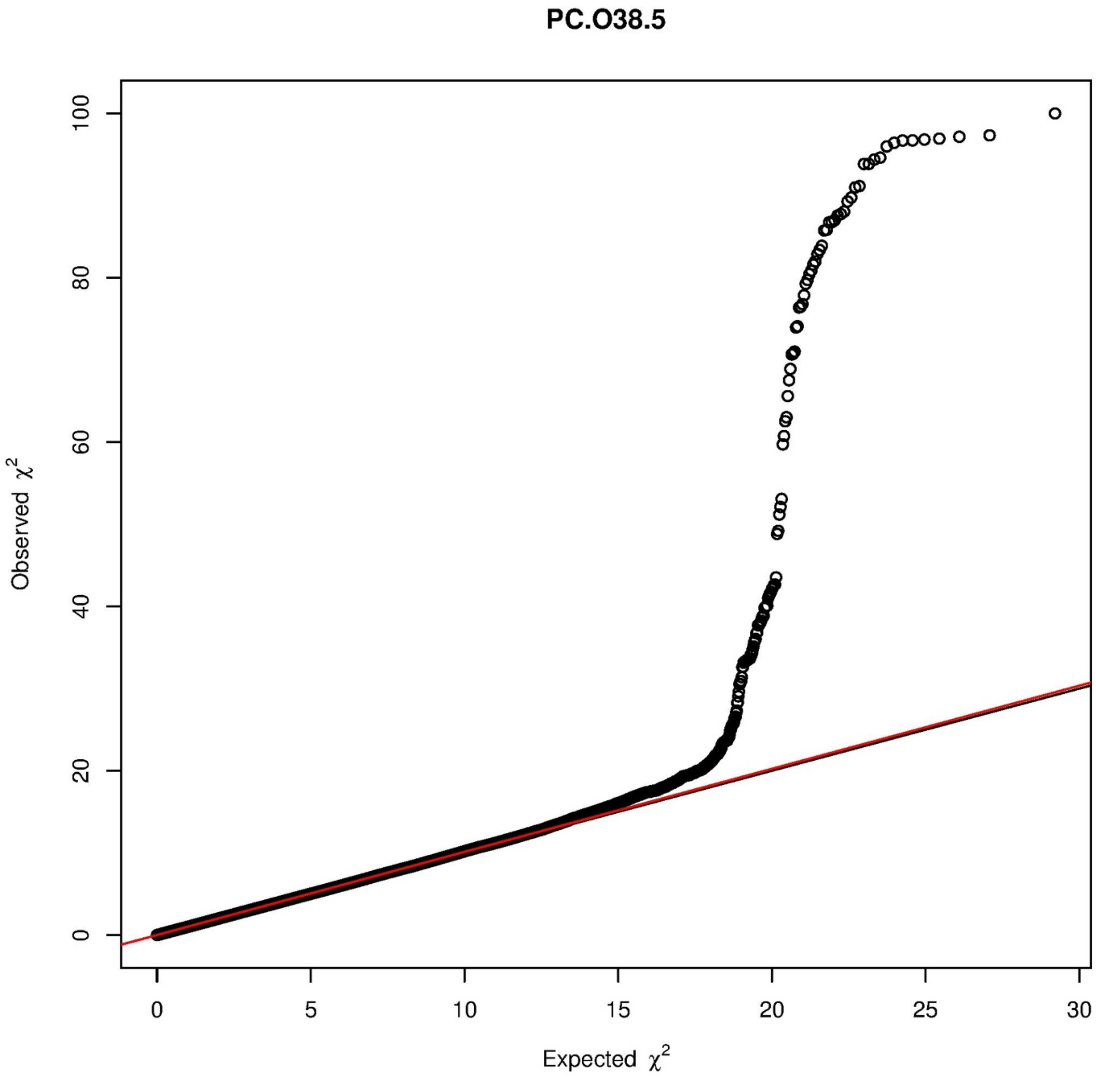

PC.O38.5
Observed χ²
Expected χ²
0
20
40
60
80
100
0
5
10
15
20
25
30

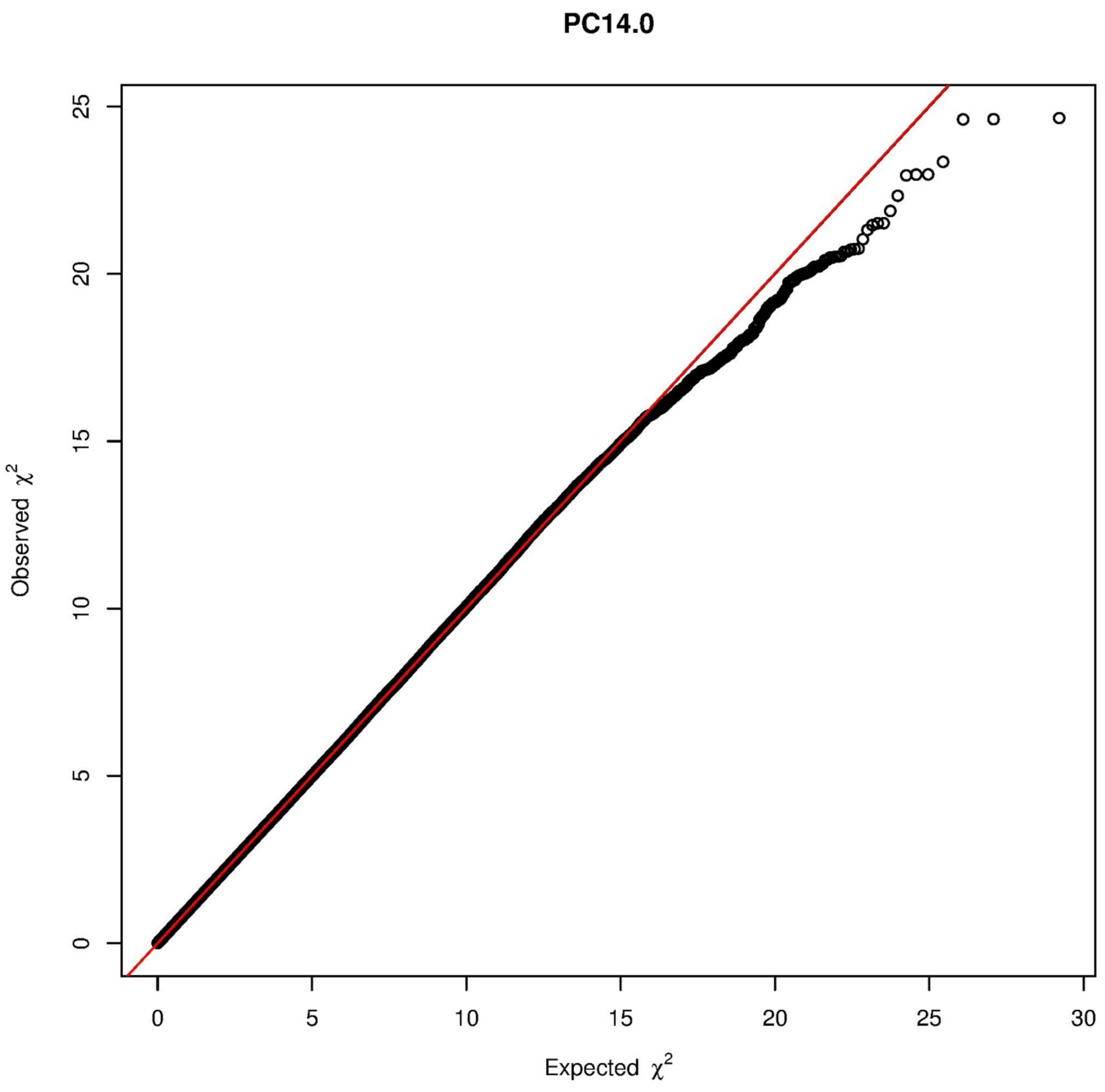

PC14.0
Observed $\chi^2$
Expected $\chi^2$
0
5
10
15
20
25
30

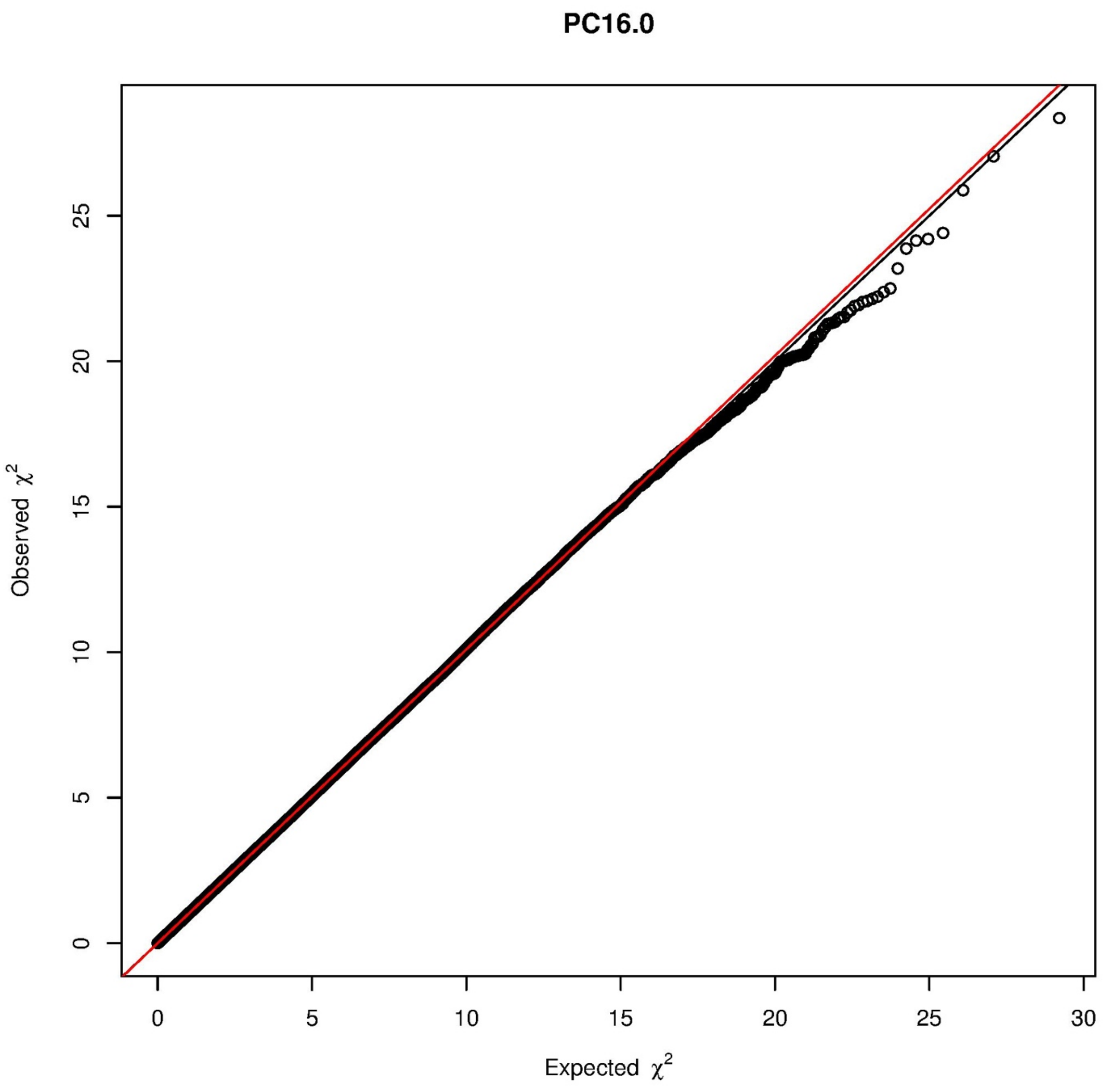

PC16.0
Observed $\chi^2$
Expected $\chi^2$
0
5
10
15
20
25
30

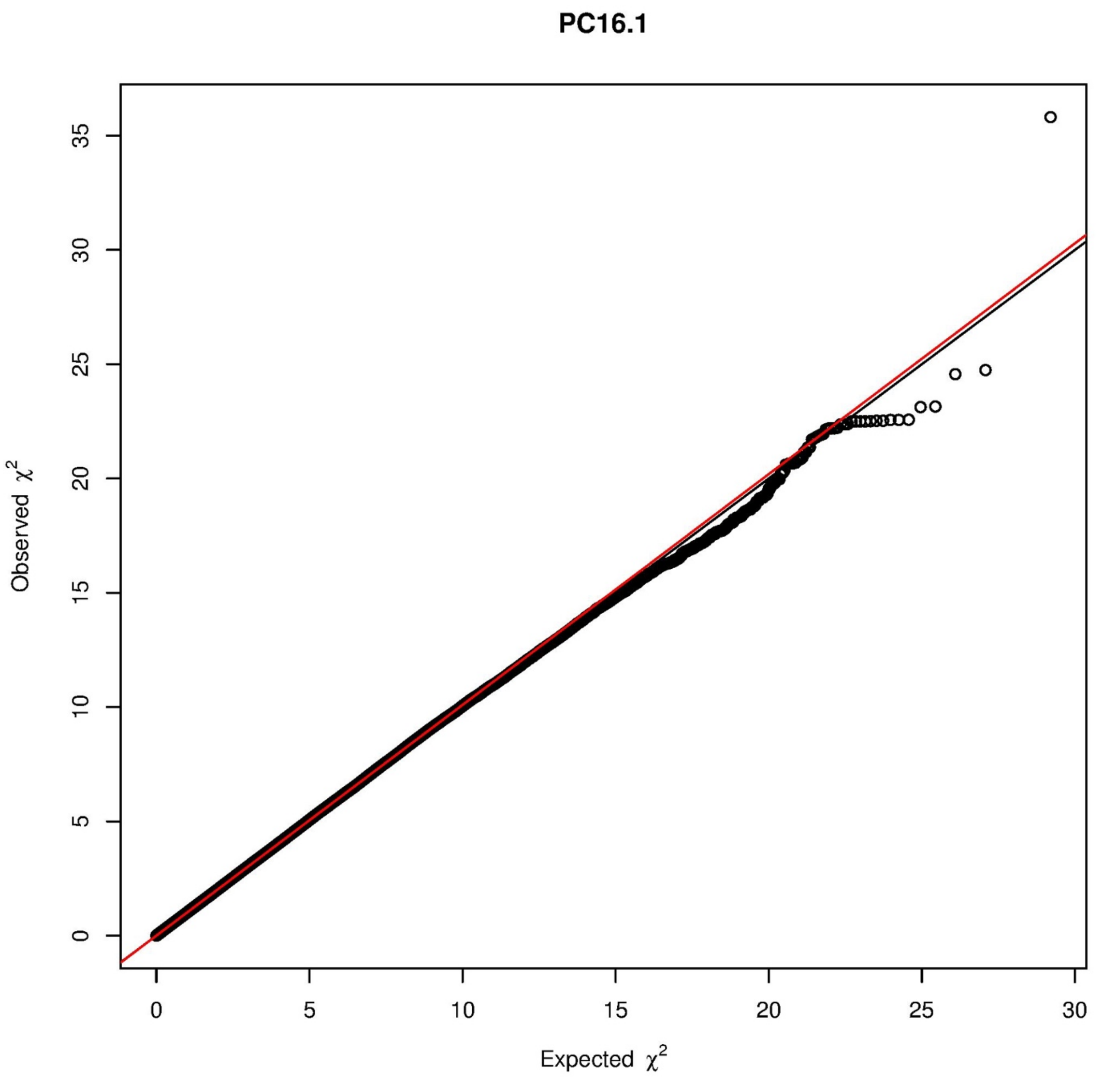

PC16.1
Observed $\chi^2$
Expected $\chi^2$
0
5
10
15
20
25
30
35

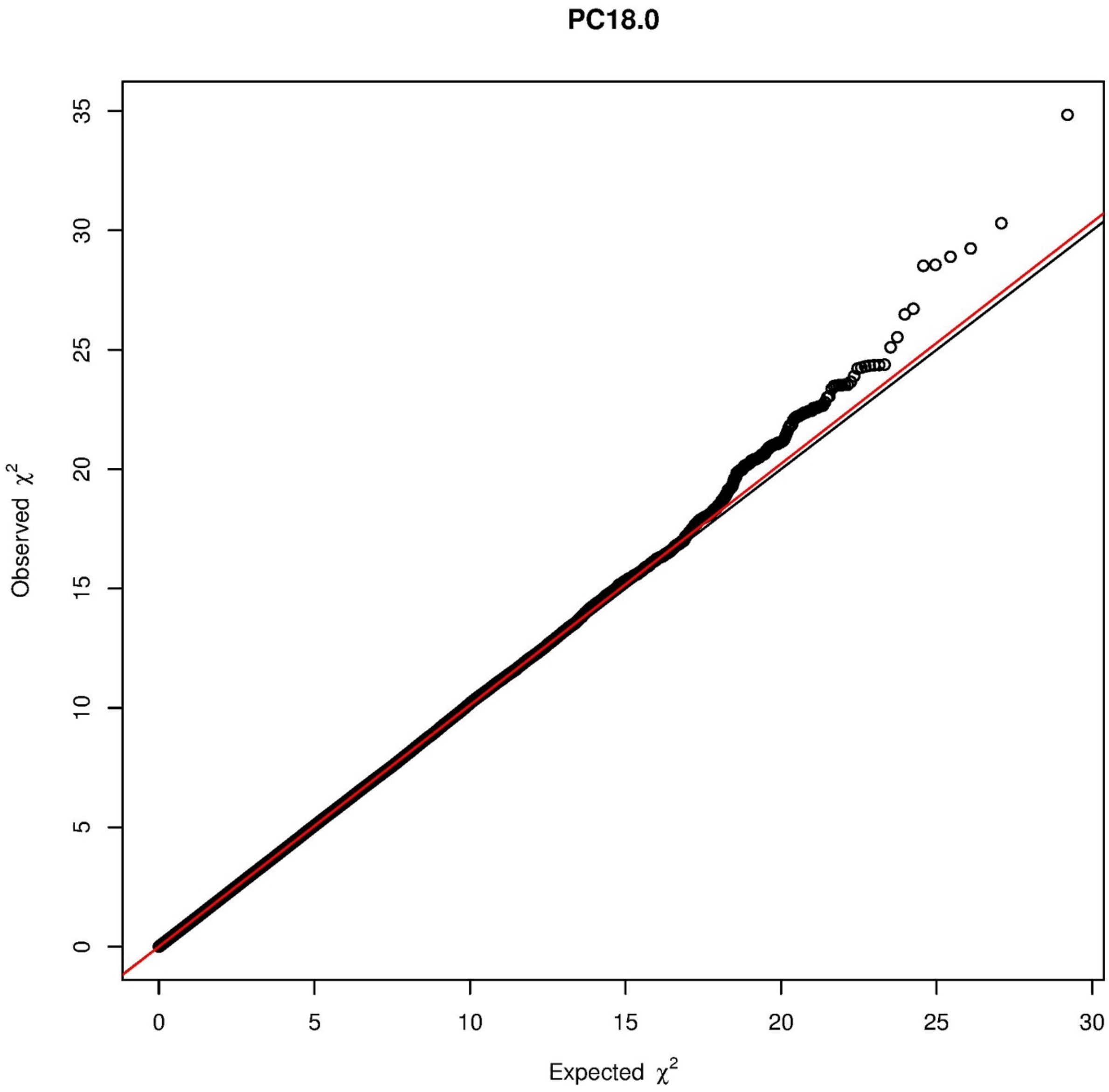

PC18.0
Observed $\chi^2$
Expected $\chi^2$
0
5
10
15
20
25
30
35

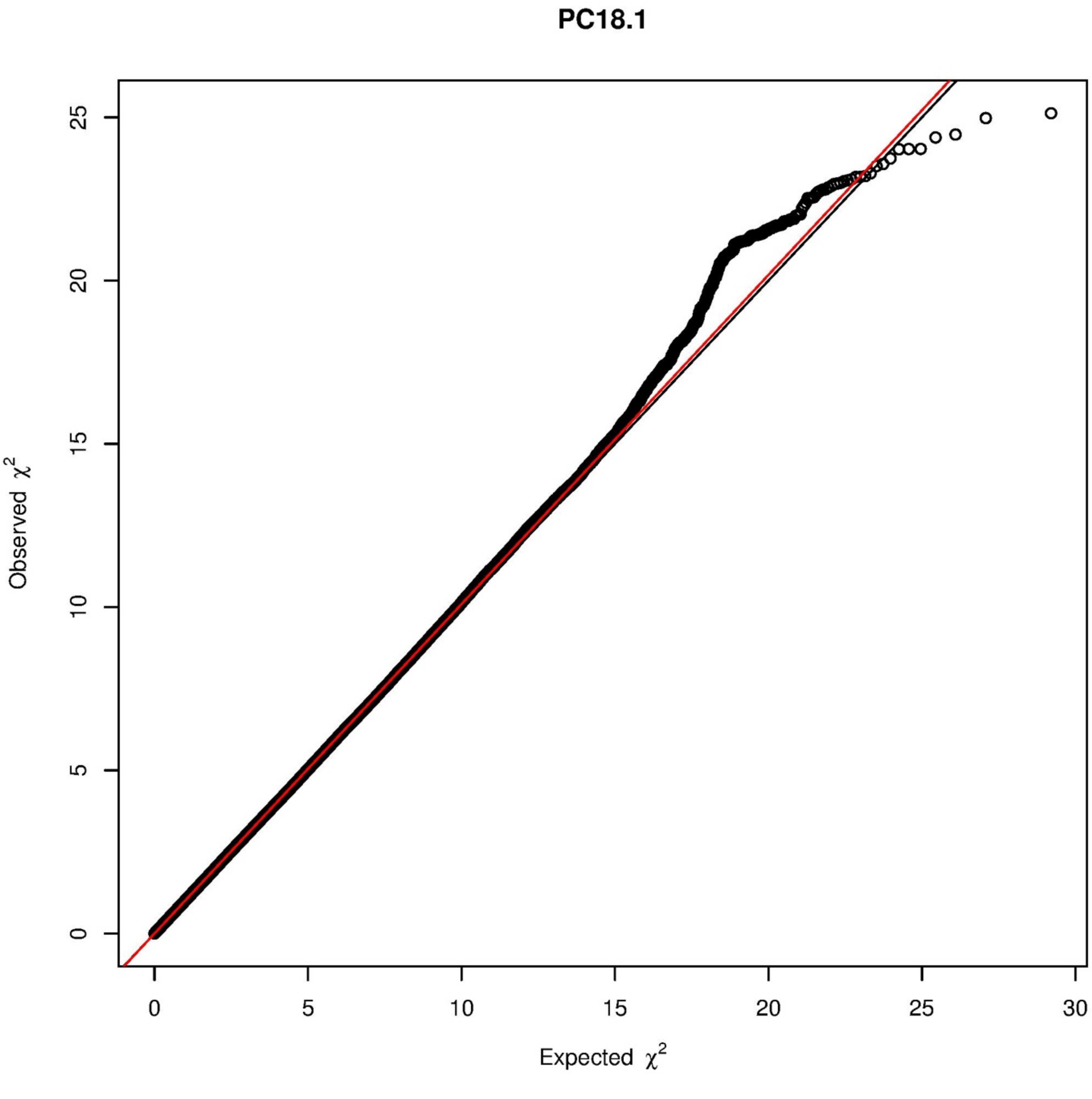

PC18.1
Observed $\chi^2$
Expected $\chi^2$
0
5
10
15
20
25
30

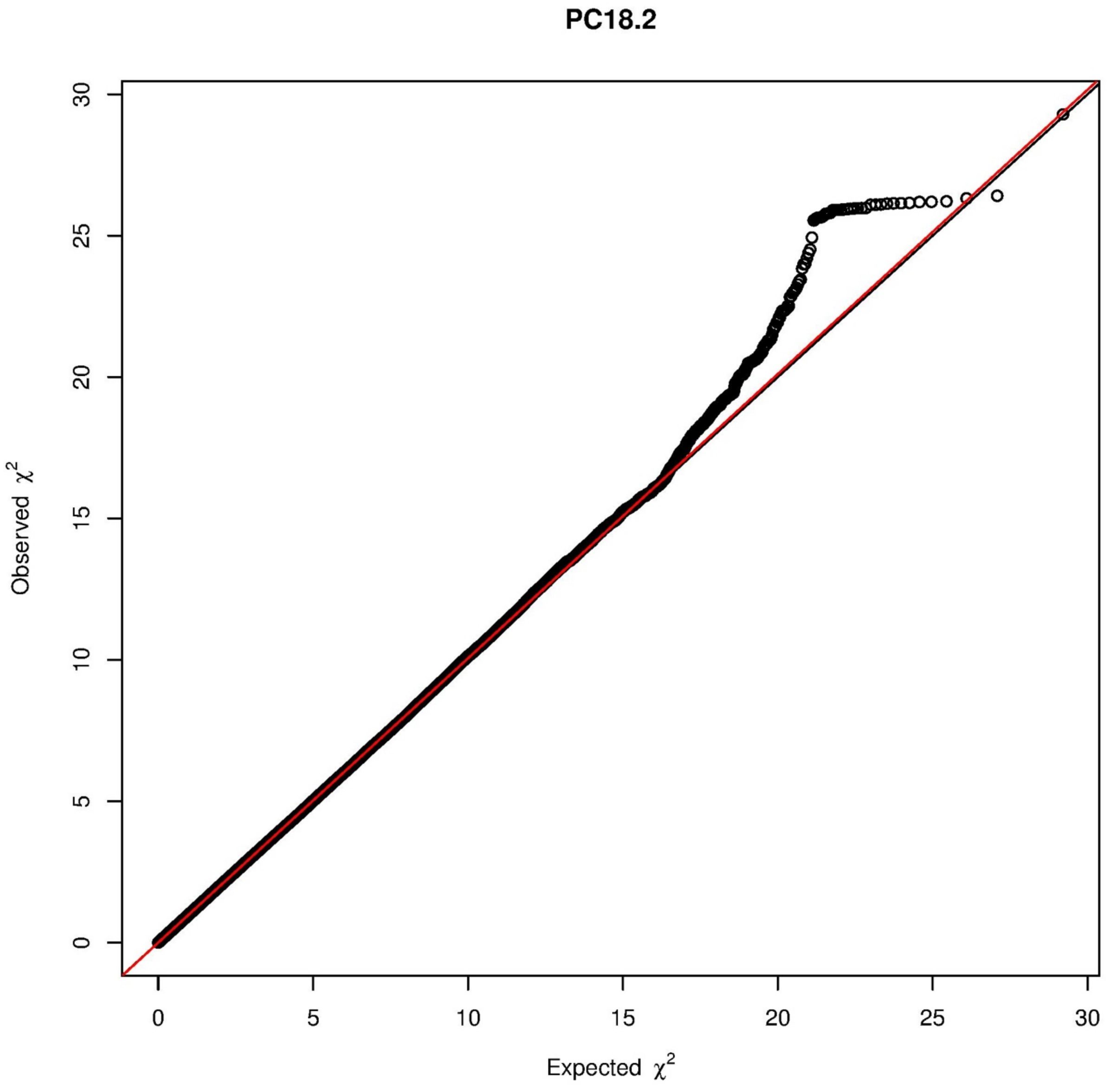
PC18.2
Observed $\chi^2$
Expected $\chi^2$
0
5
10
15
20
25
30

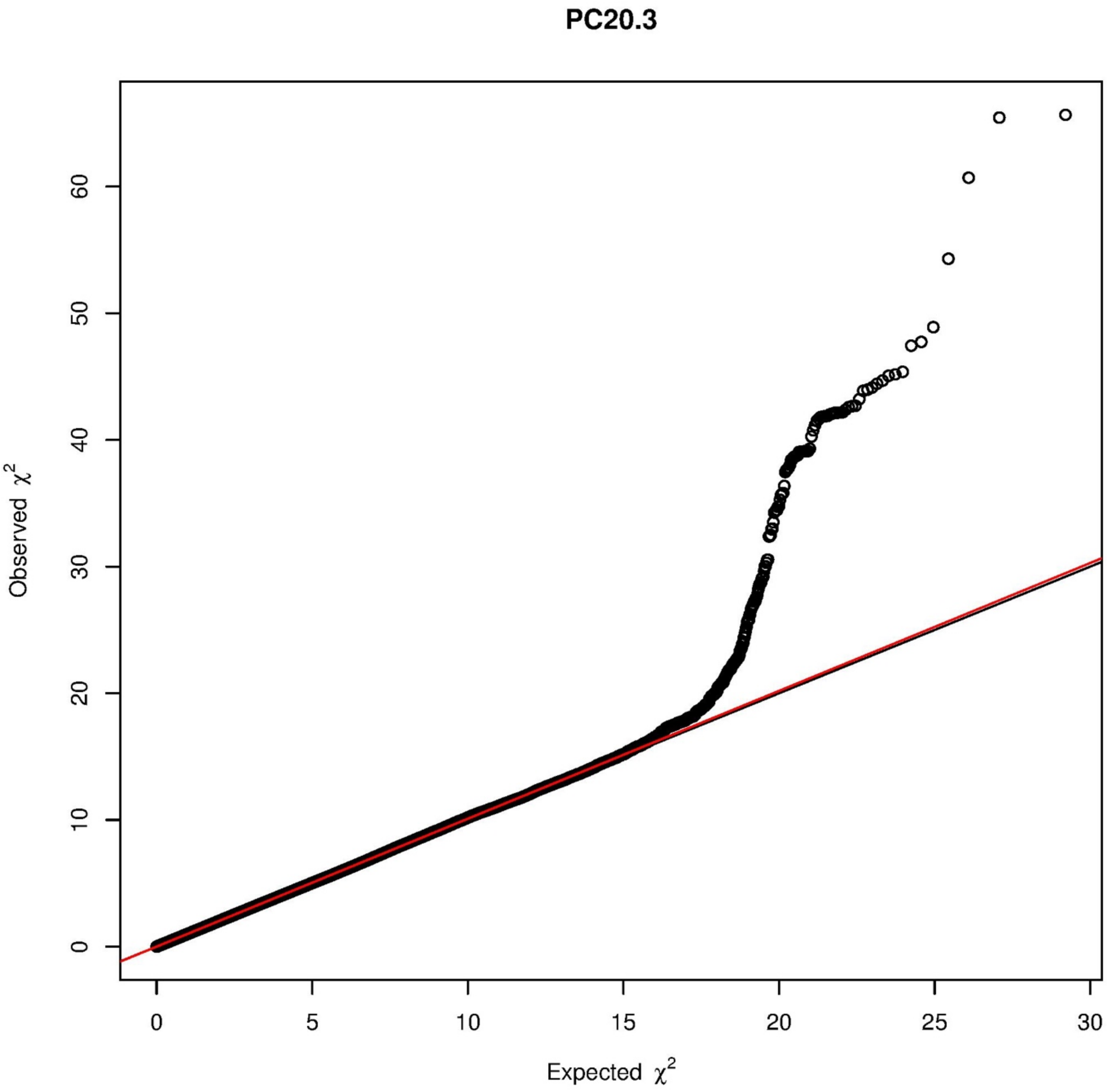
PC20.3
Observed $\chi^2$
Expected $\chi^2$
0
10
20
30
40
50
60
0
5
10
15
20
25
30

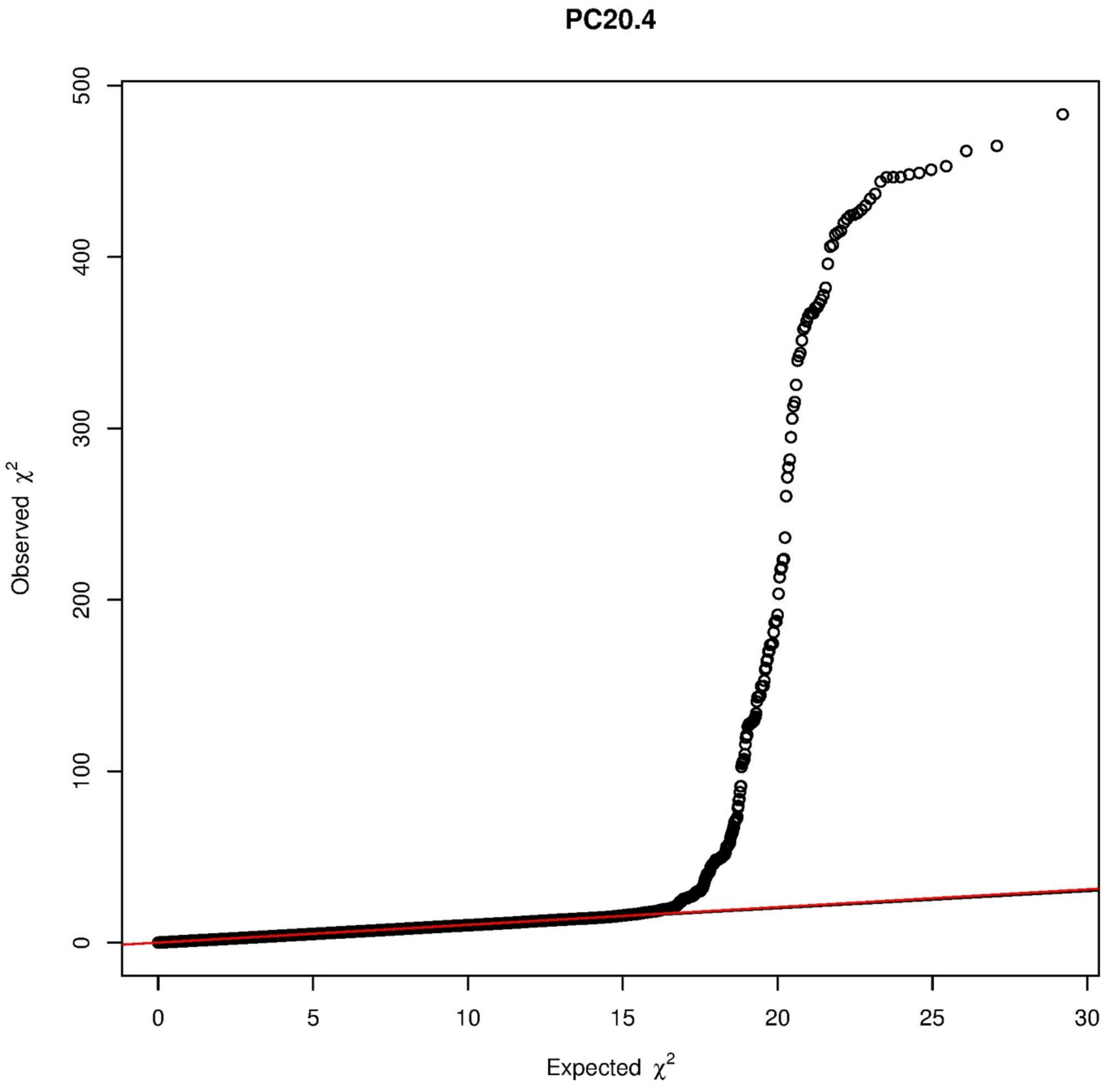
PC20.4
Observed $\chi^2$
Expected $\chi^2$
0
100
200
300
400
500
0
5
10
15
20
25
30

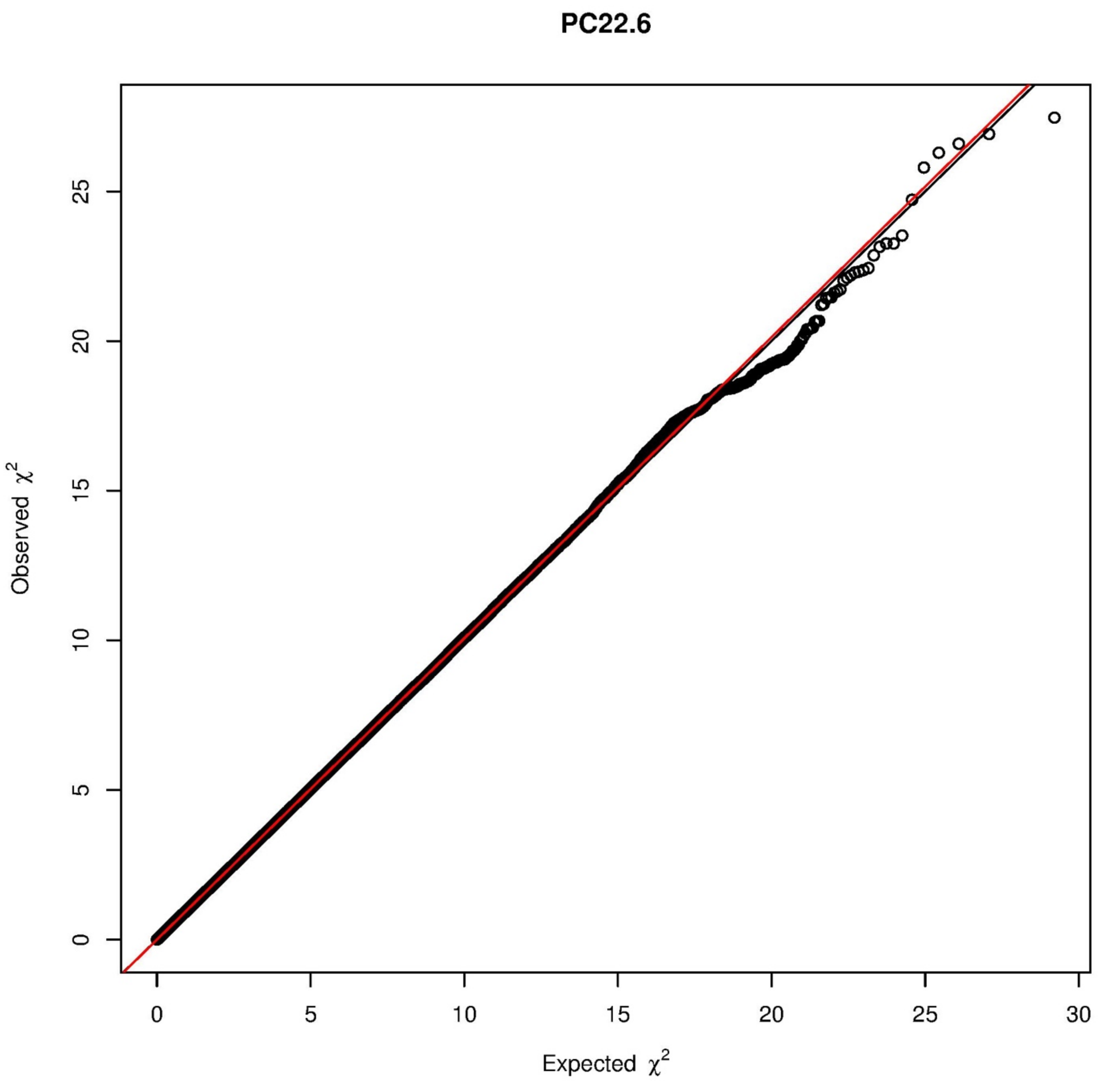
PC22.6
Observed $\chi^2$
Expected $\chi^2$
0
5
10
15
20
25
30

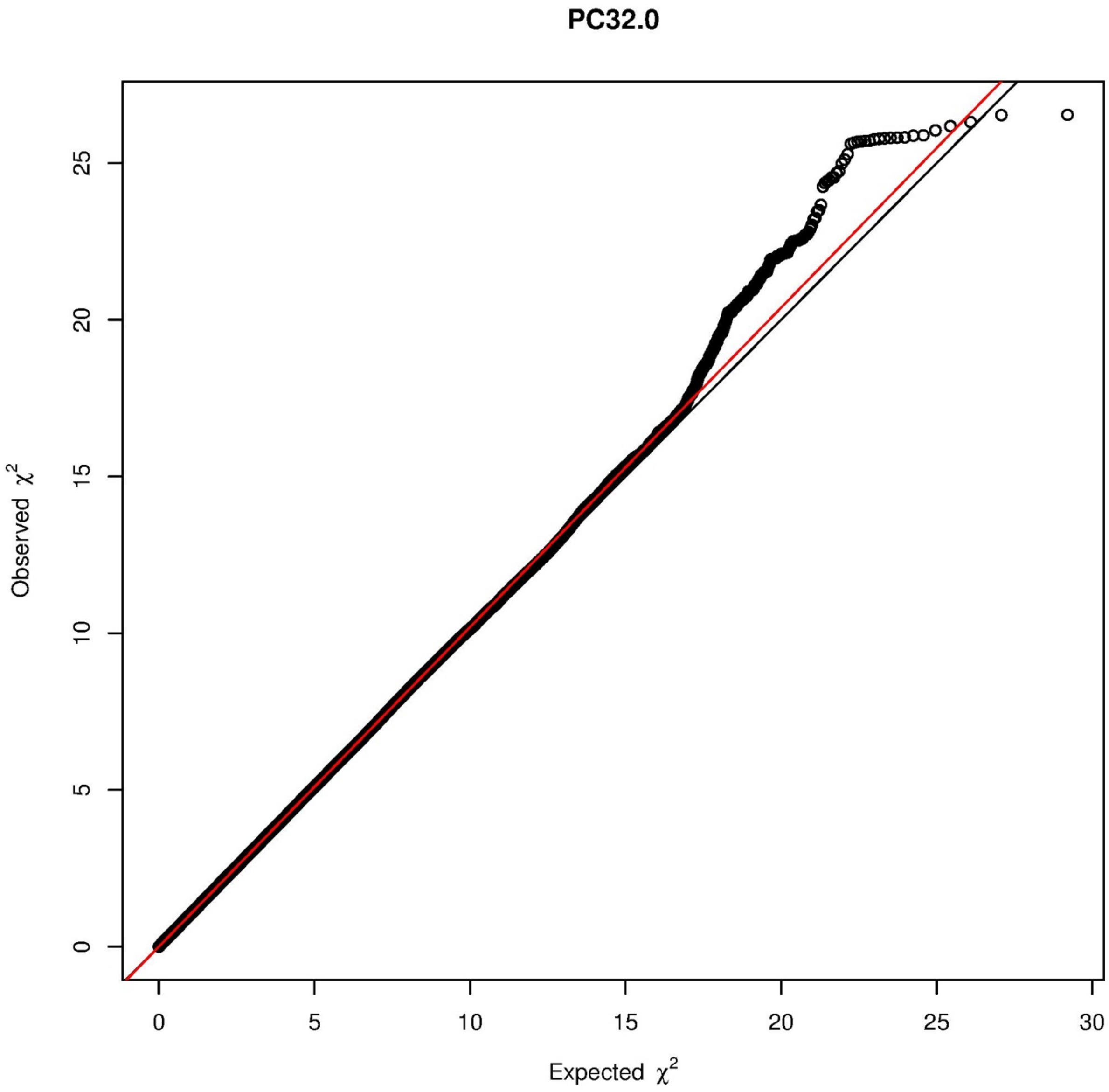
PC32.0
Observed χ2
Expected χ2
0
5
10
15
20
25
30

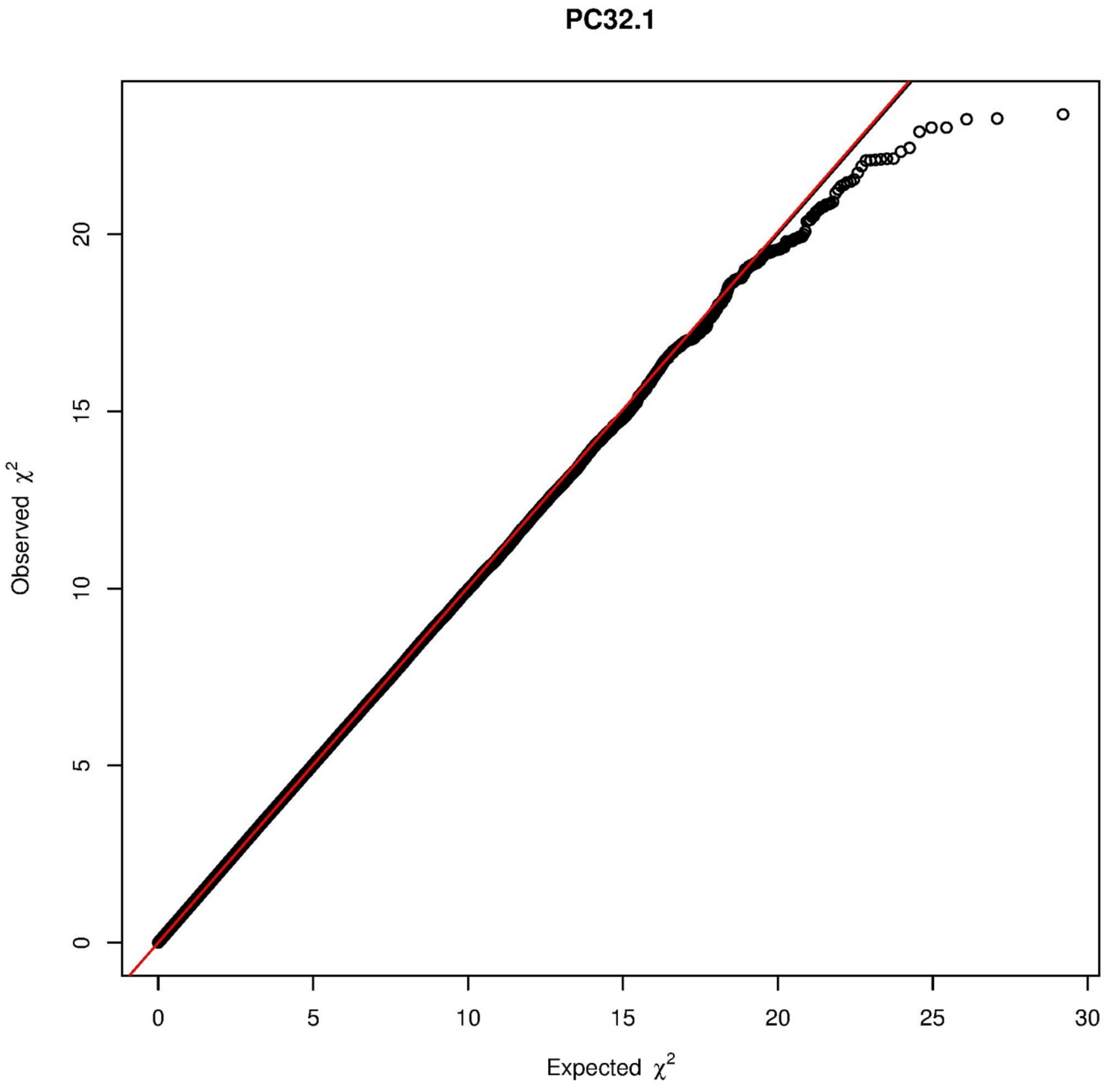
PC32.1
Observed χ2
Expected χ2
0
5
10
15
20
25
30

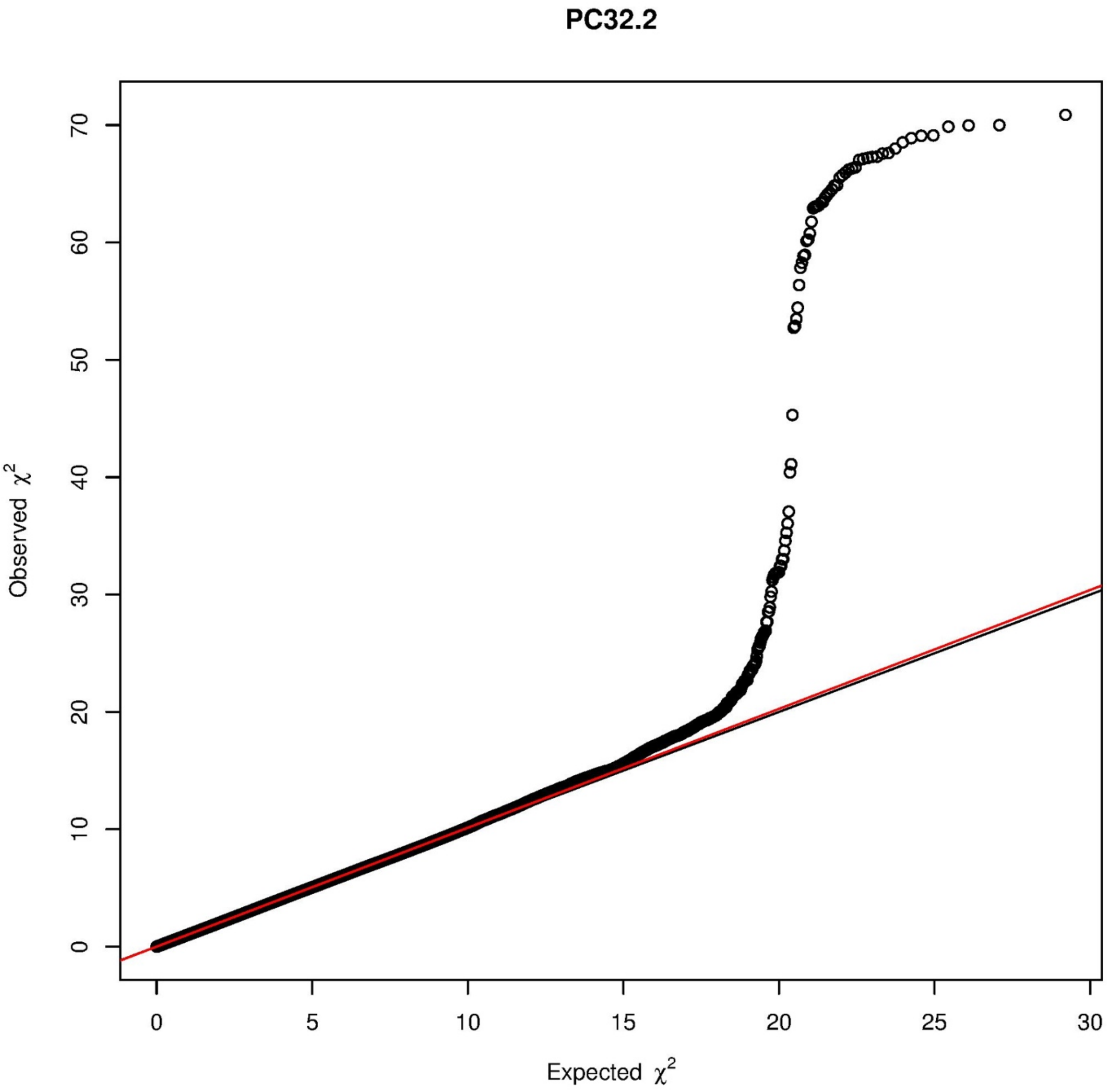
PC32.2
Observed $\chi^2$
Expected $\chi^2$
0
10
20
30
40
50
60
70
0
5
10
15
20
25
30

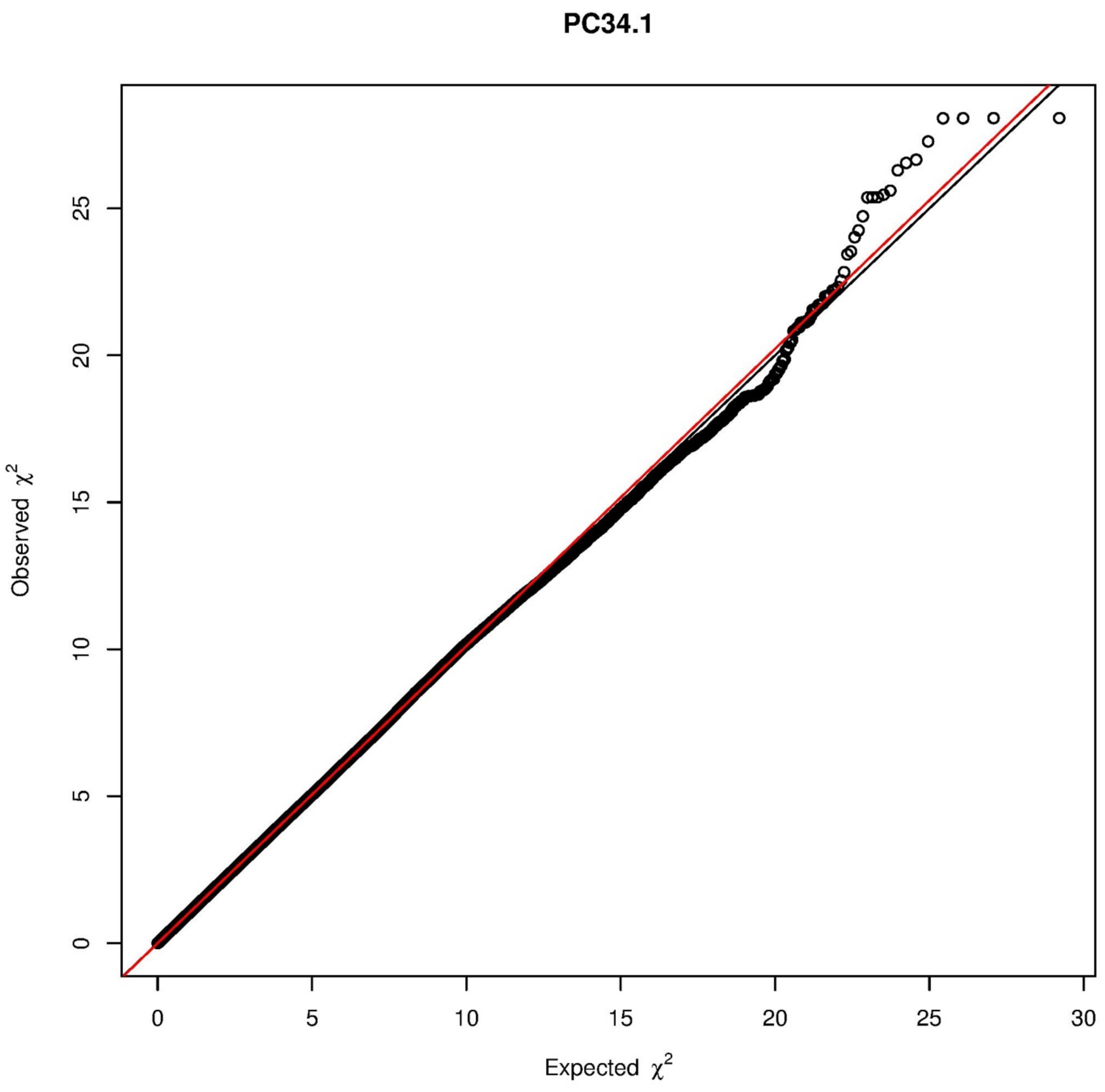
PC34.1
Observed $\chi^2$
Expected $\chi^2$
0
5
10
15
20
25
30

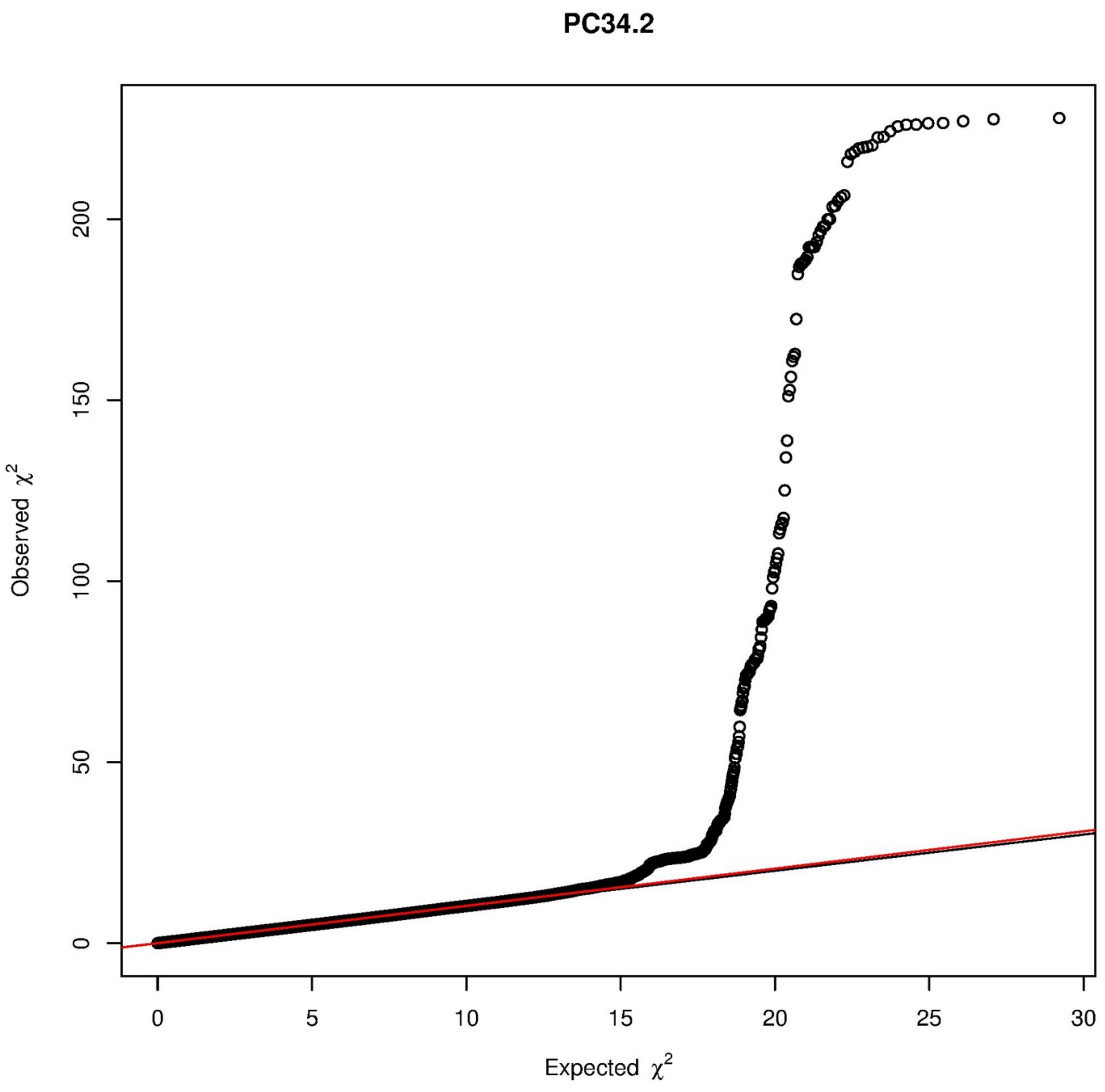

PC34.2
Observed $\chi^2$
Expected $\chi^2$
0
50
100
150
200
0
5
10
15
20
25
30

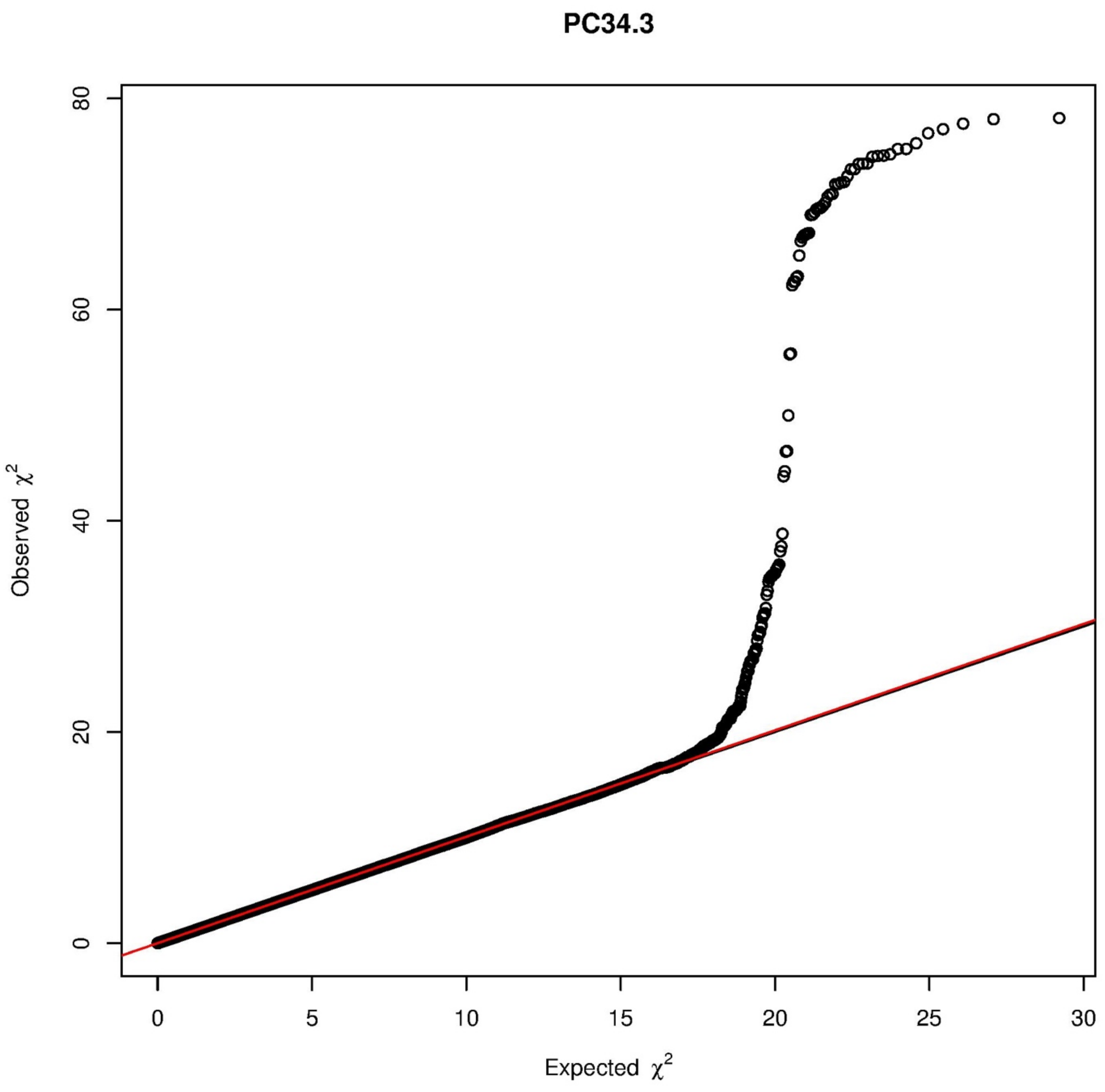

PC34.3
80
60
40
20
0
Observed χ2
0
5
10
15
20
25
30
Expected χ2

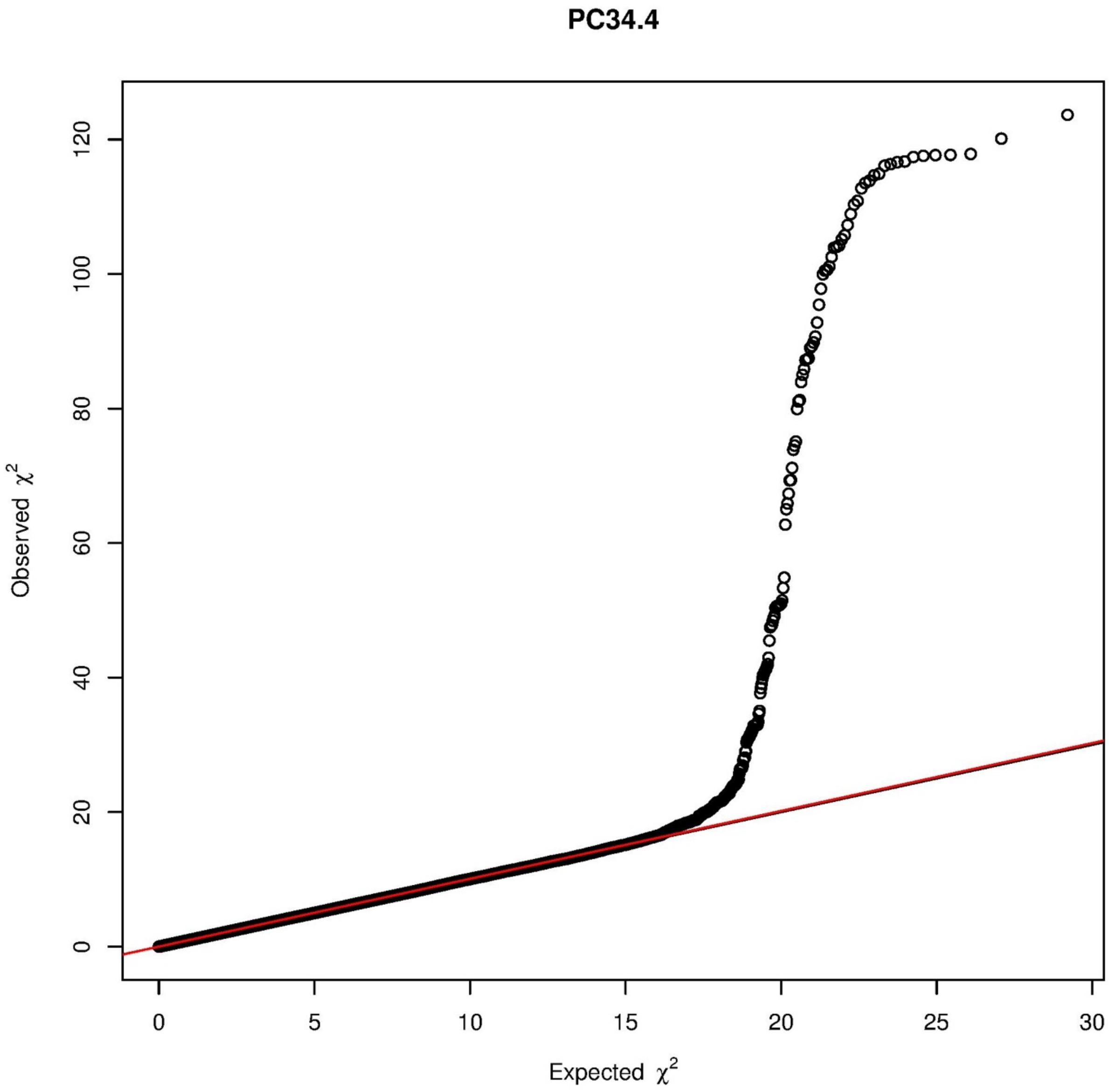

PC34.4
Observed χ²
Expected χ²
0
20
40
60
80
100
120
0
5
10
15
20
25
30

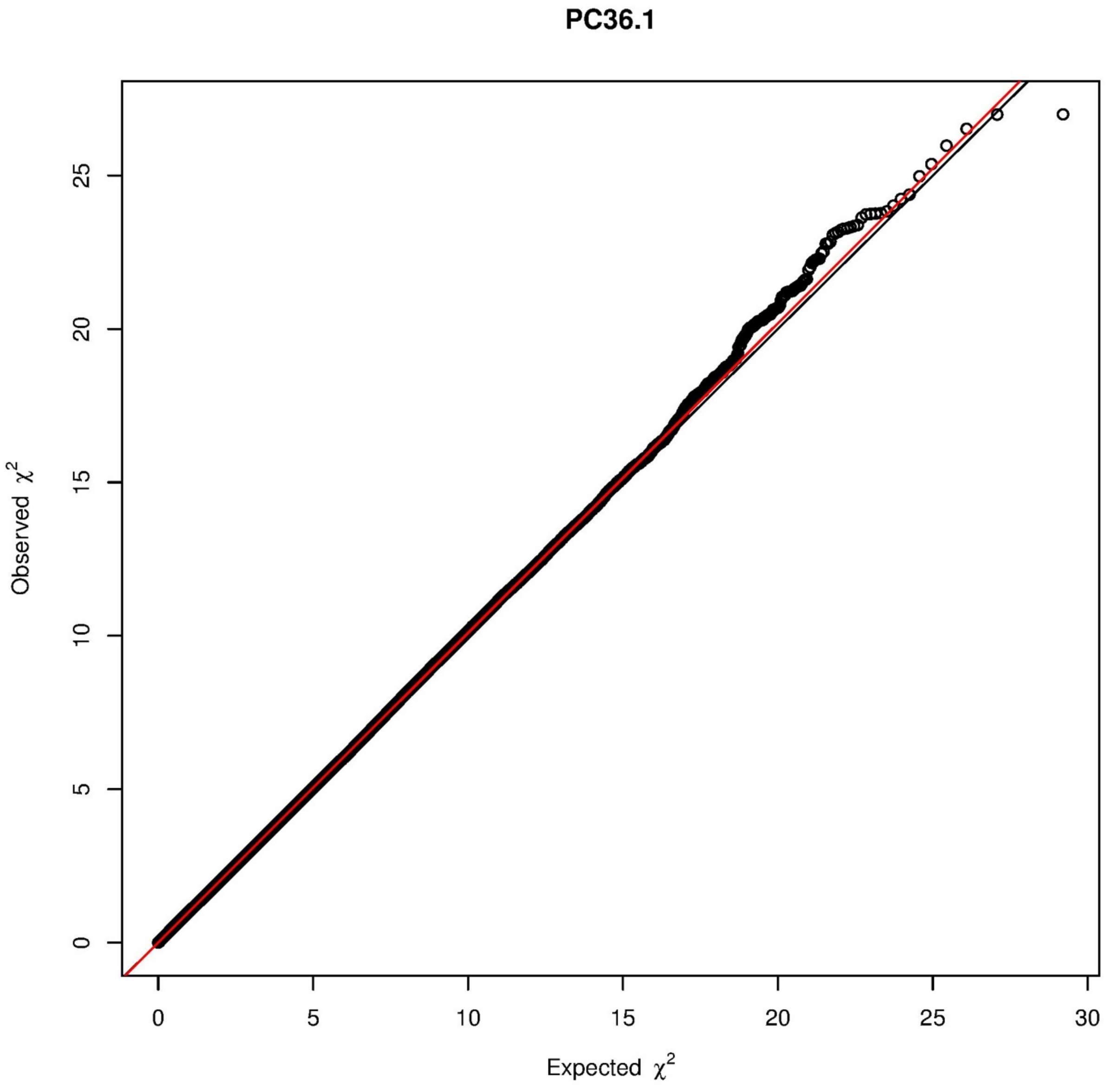
PC36.1
Observed χ2
Expected χ2
0
5
10
15
20
25
30

PC36.2

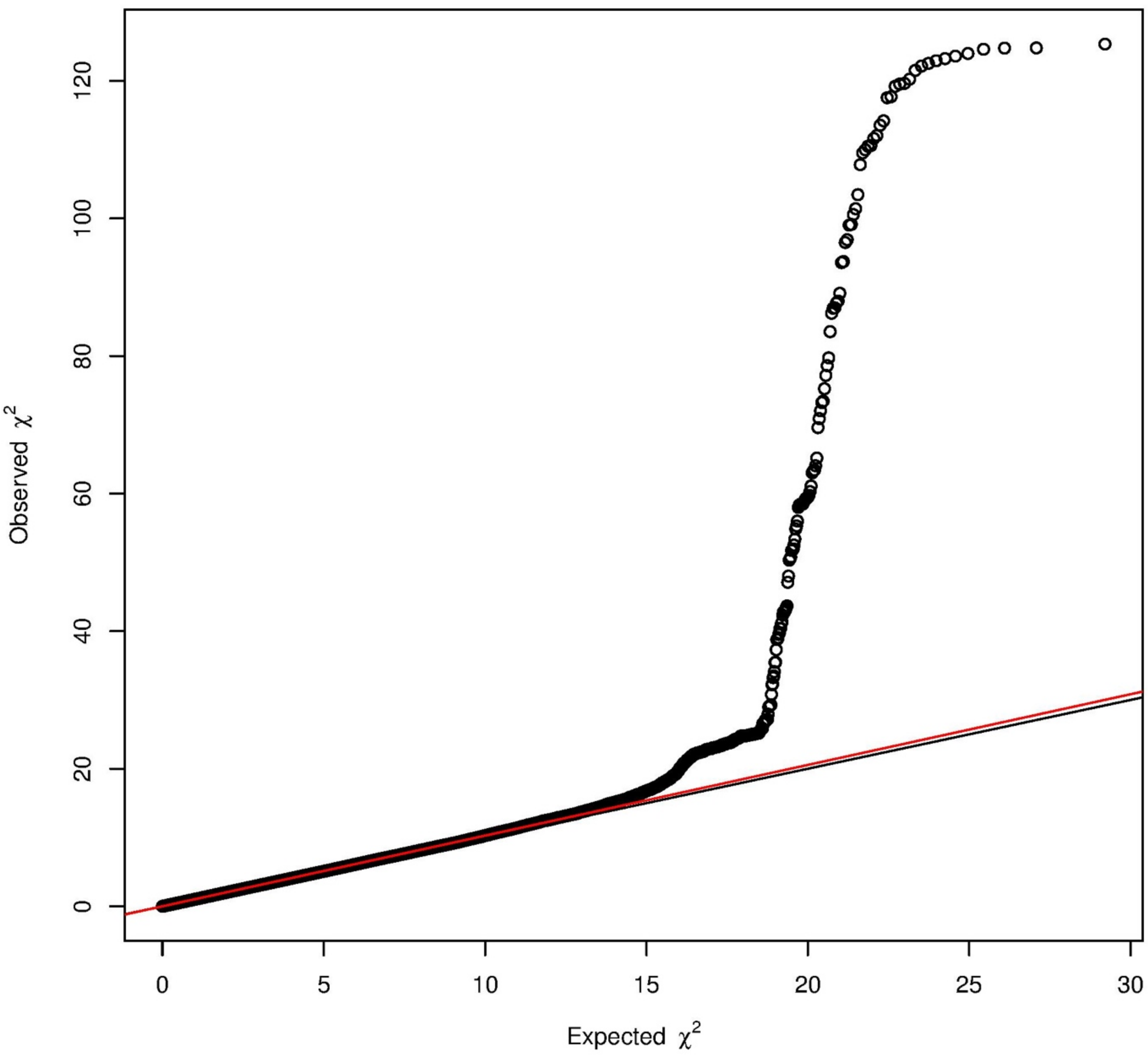

Observed $\chi^2$
Expected $\chi^2$
0
20
40
60
80
100
120
0
5
10
15
20
25
30

## PC36.3

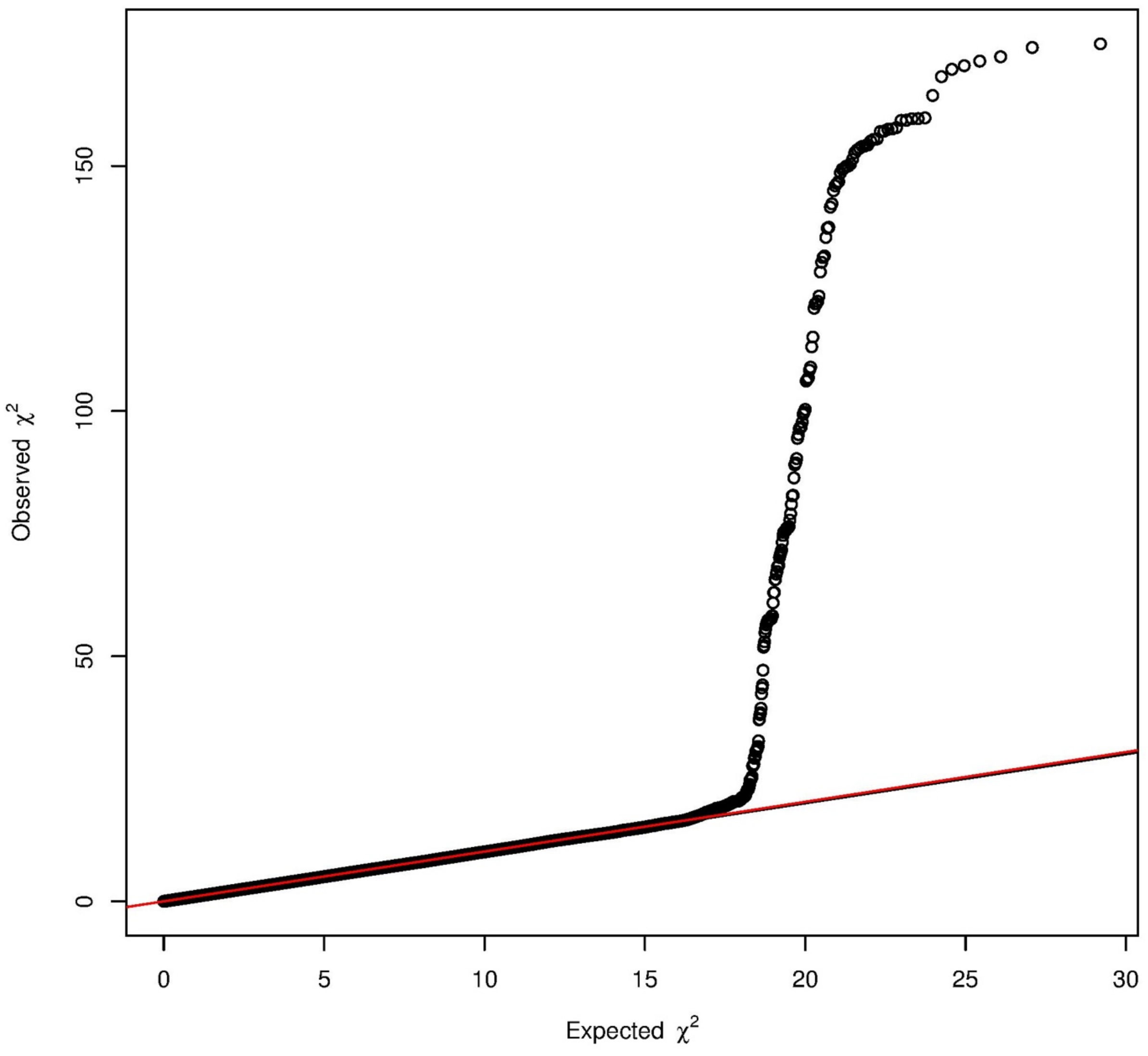

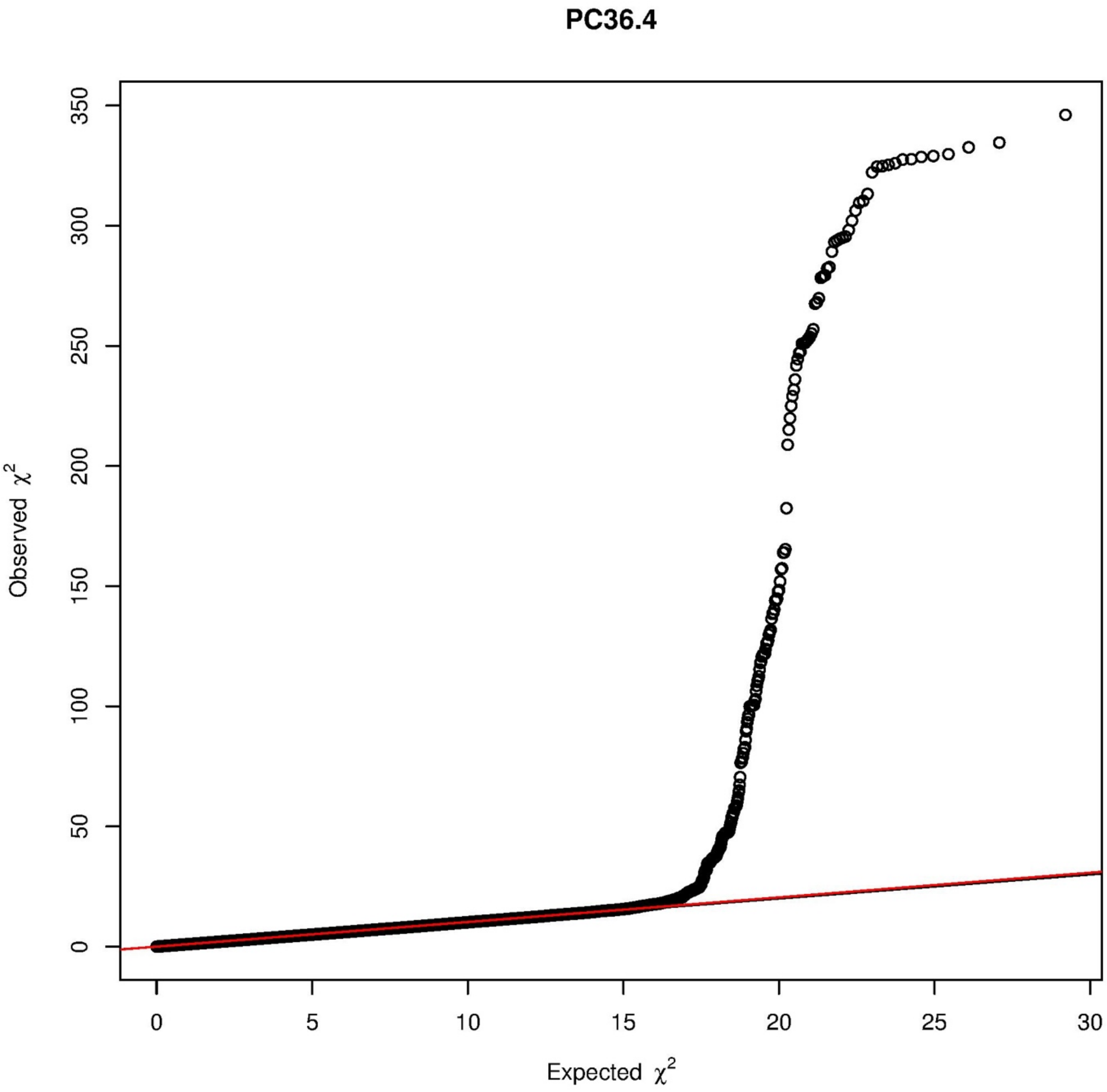
PC36.4
Observed $\chi^2$
Expected $\chi^2$
0
50
100
150
200
250
300
350
0
5
10
15
20
25
30

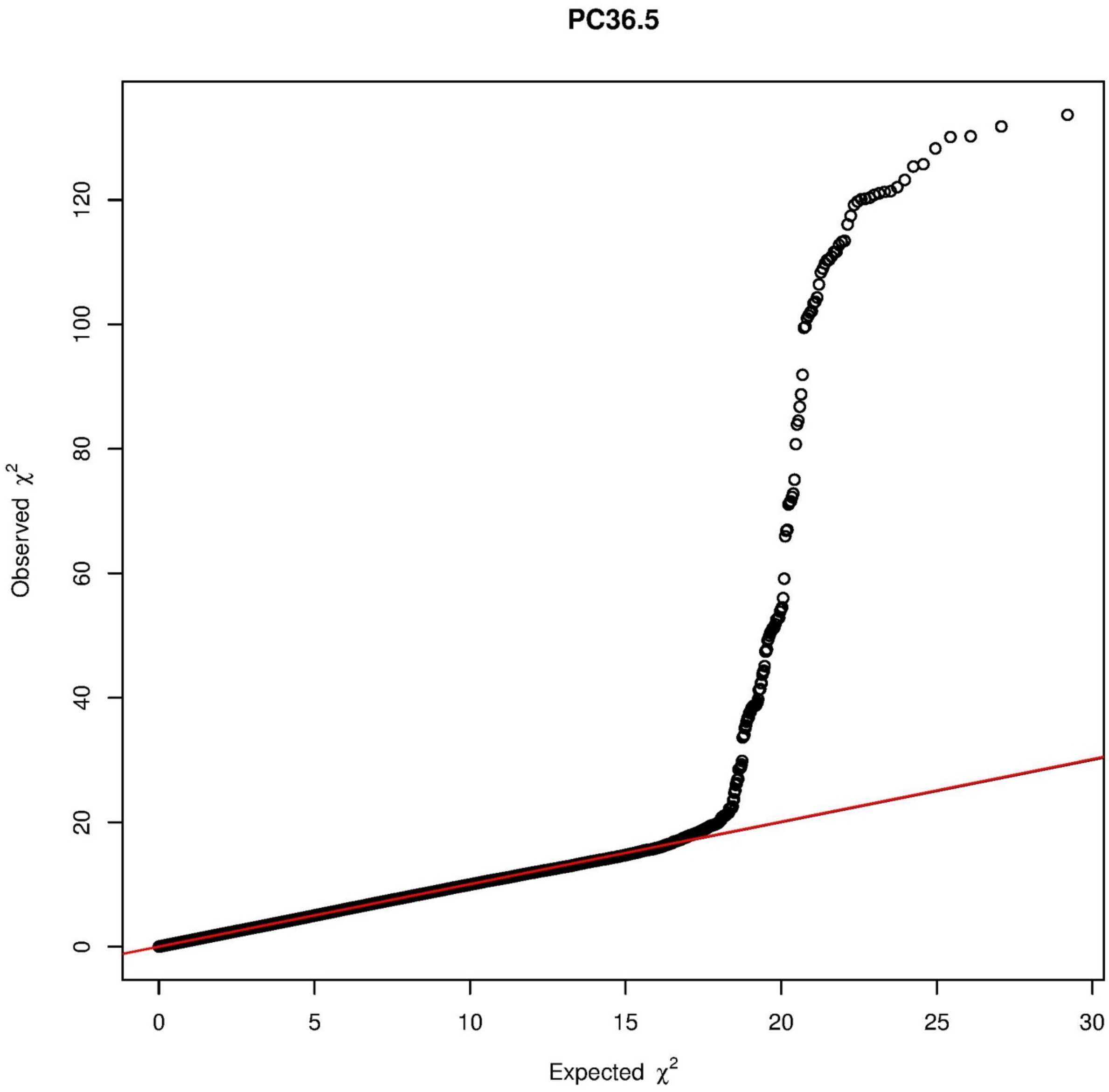
PC36.5
Observed $\chi^2$
0
20
40
60
80
100
120
Expected $\chi^2$
0
5
10
15
20
25
30

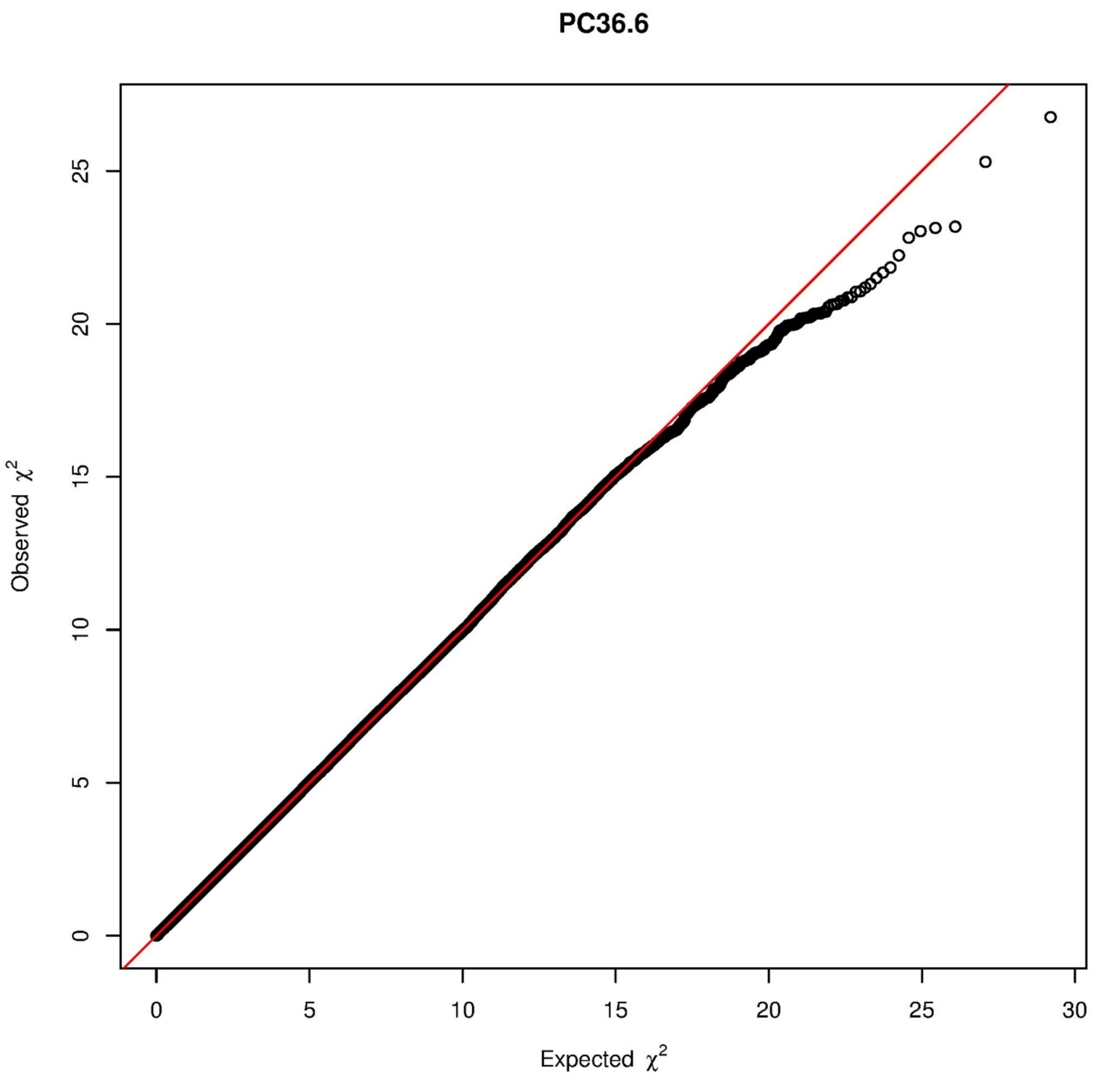

PC36.6
Observed $\chi^2$
Expected $\chi^2$
0
5
10
15
20
25
30

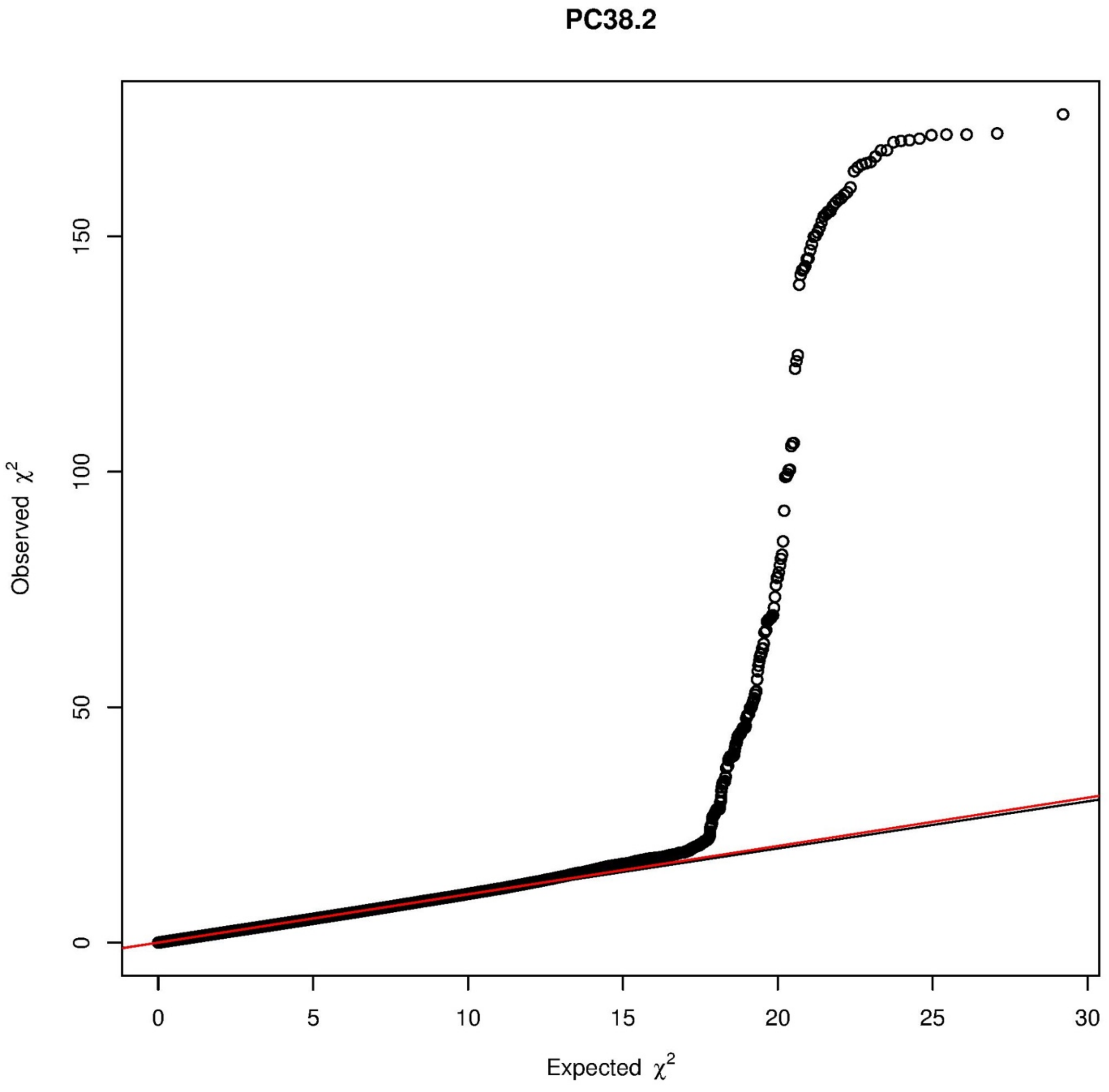
PC38.2
Observed $\chi^2$
Expected $\chi^2$
0
50
100
150
0
5
10
15
20
25
30

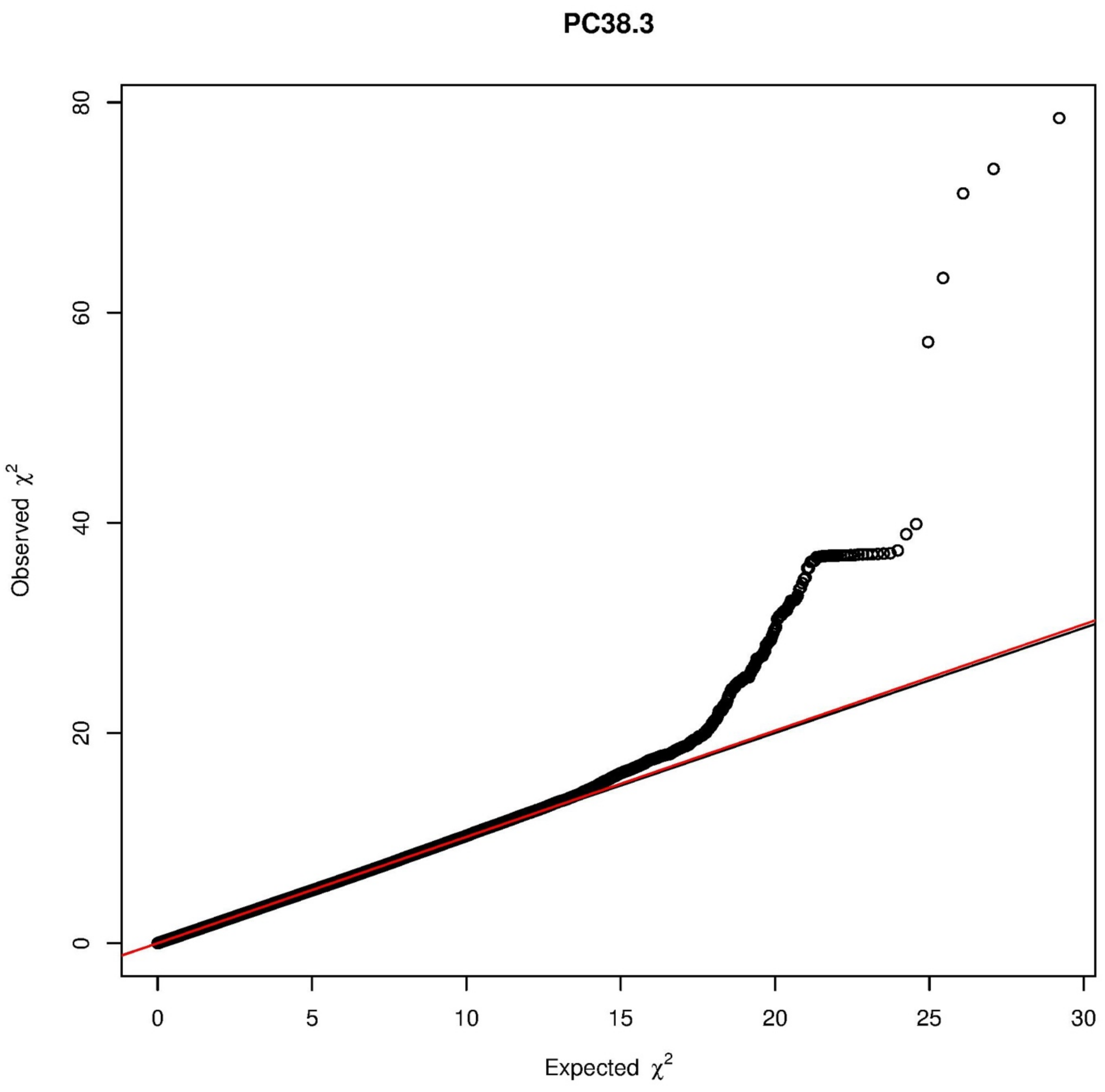

PC38.3
Observed $\chi^2$
Expected $\chi^2$
0
20
40
60
80
0
5
10
15
20
25
30

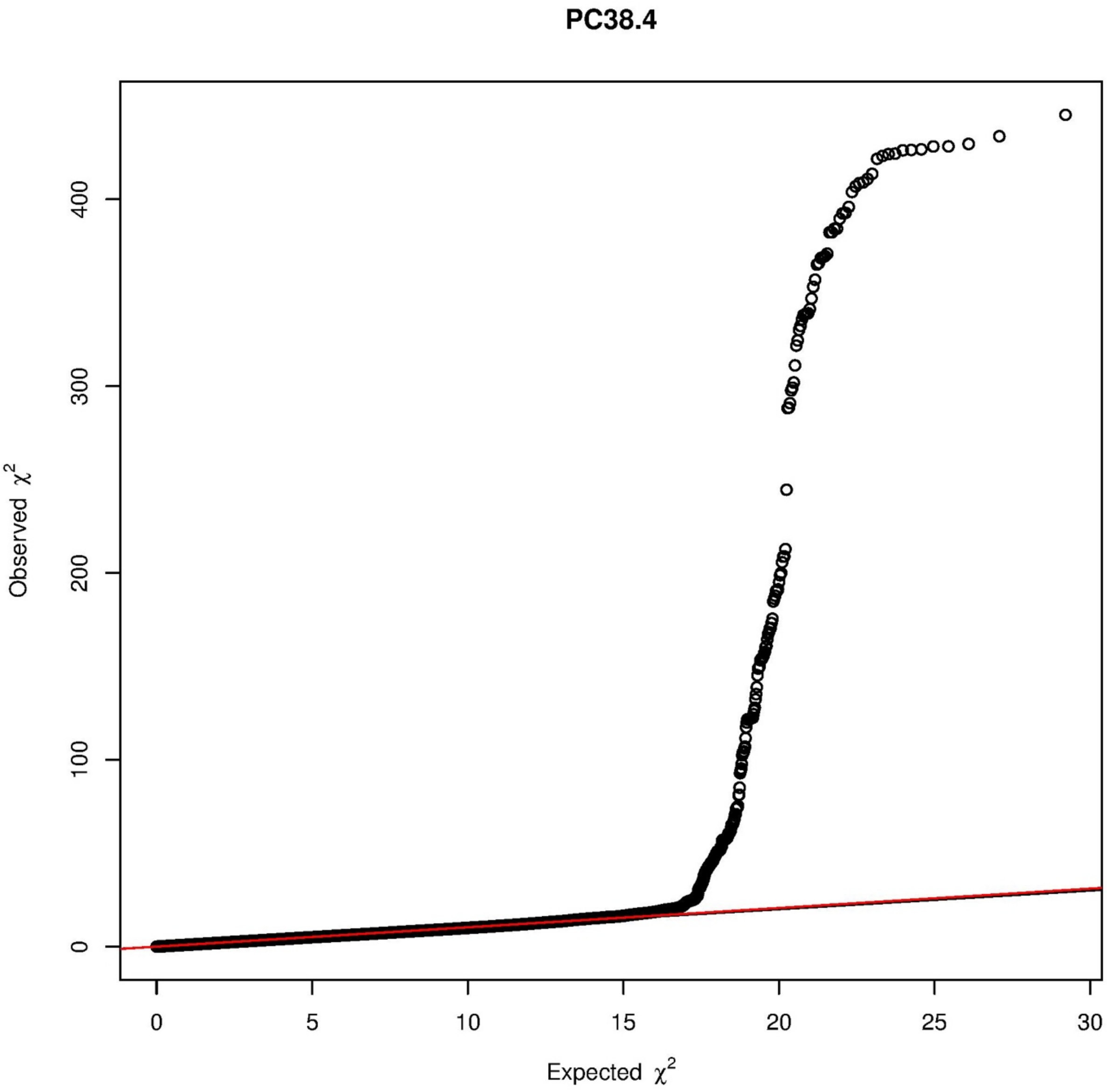

PC38.4
Observed $\chi^2$
Expected $\chi^2$
0
100
200
300
400
0
5
10
15
20
25
30

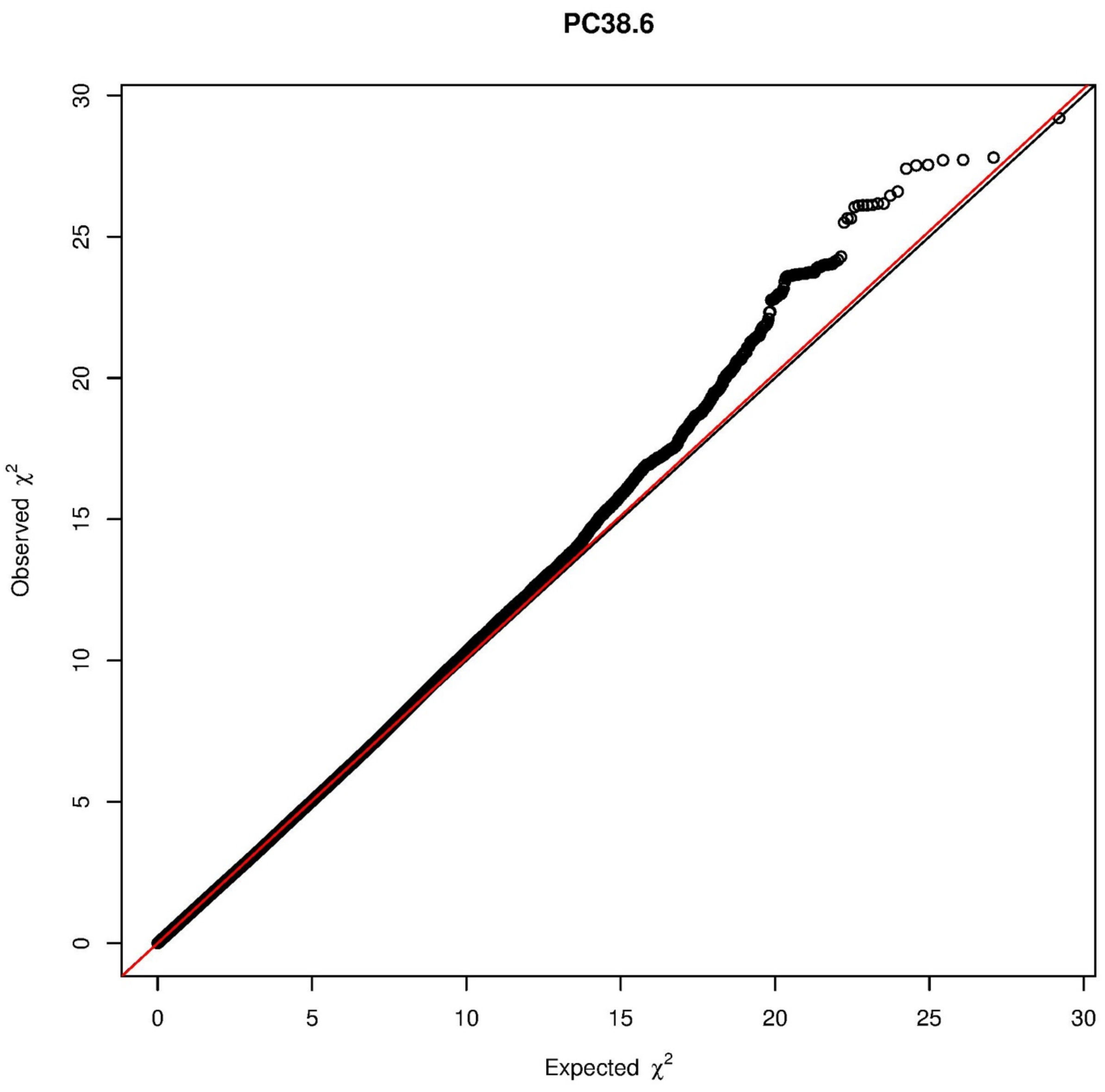
PC38.6
Observed χ2
Expected χ2
0
5
10
15
20
25
30
0
5
10
15
20
25
30

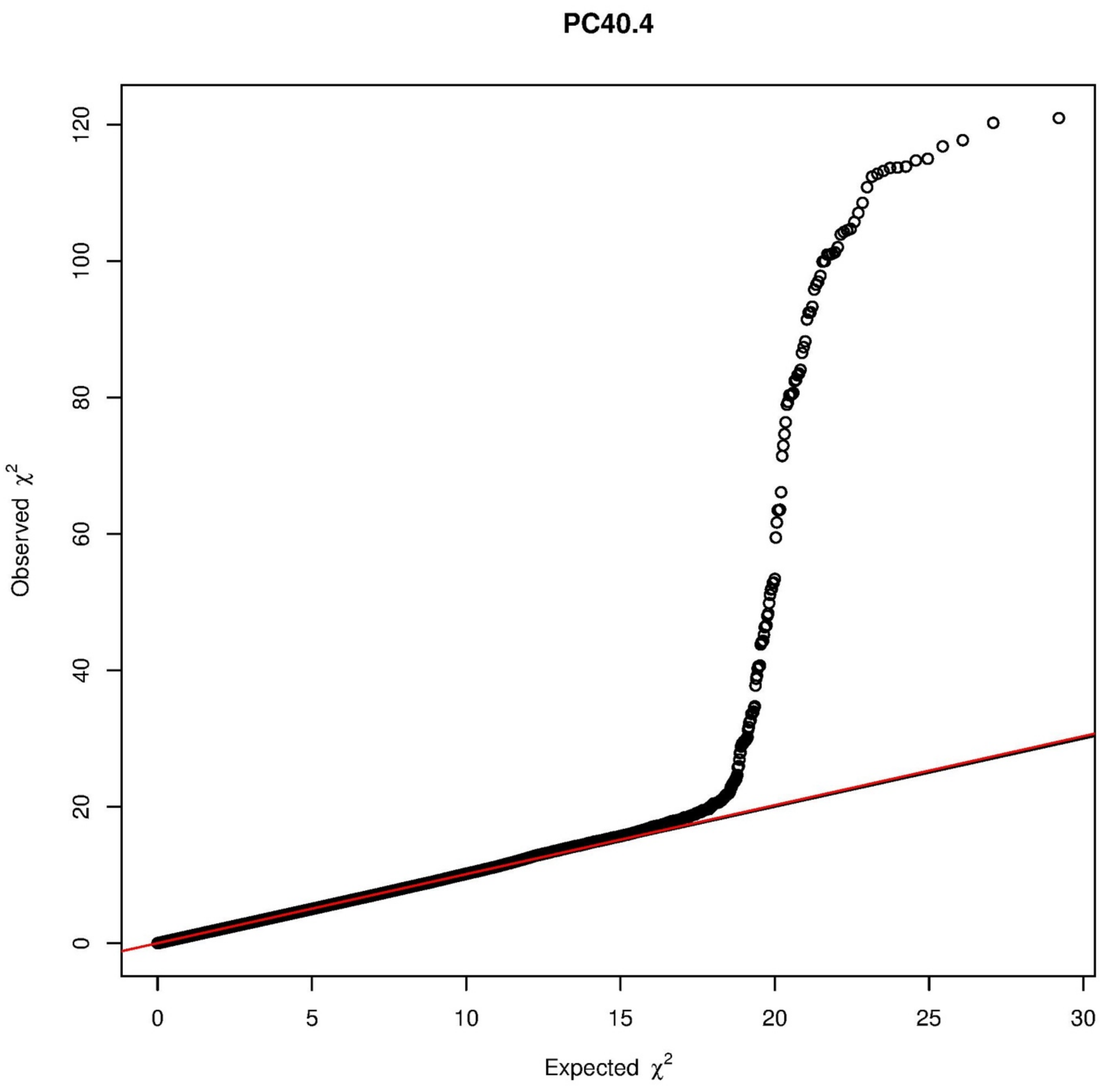
PC40.4
Observed $\chi^2$
Expected $\chi^2$
0
20
40
60
80
100
120
0
5
10
15
20
25
30

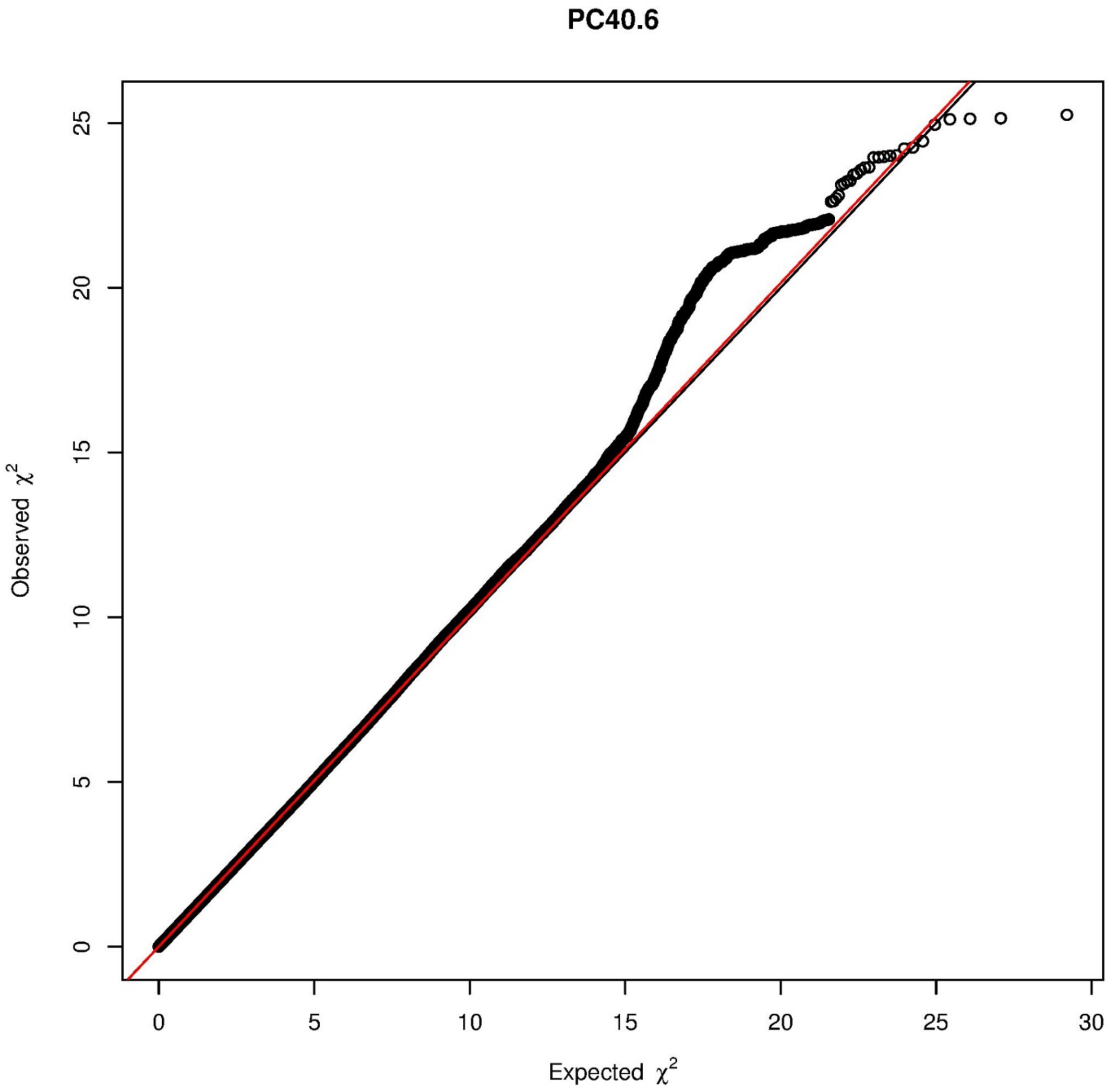
PC40.6
Observed χ2
Expected χ2
0
5
10
15
20
25
30

PE.O36.5

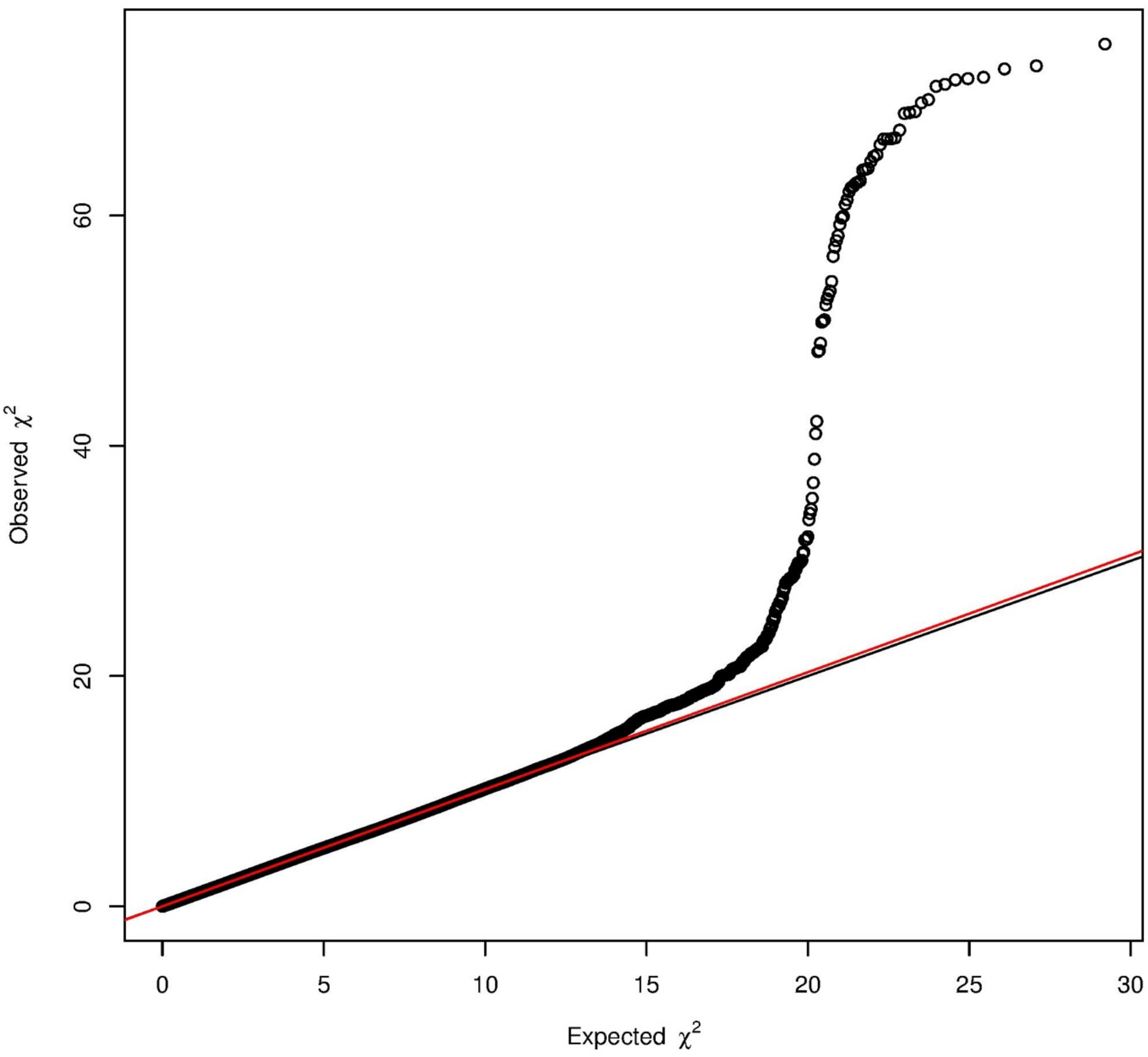

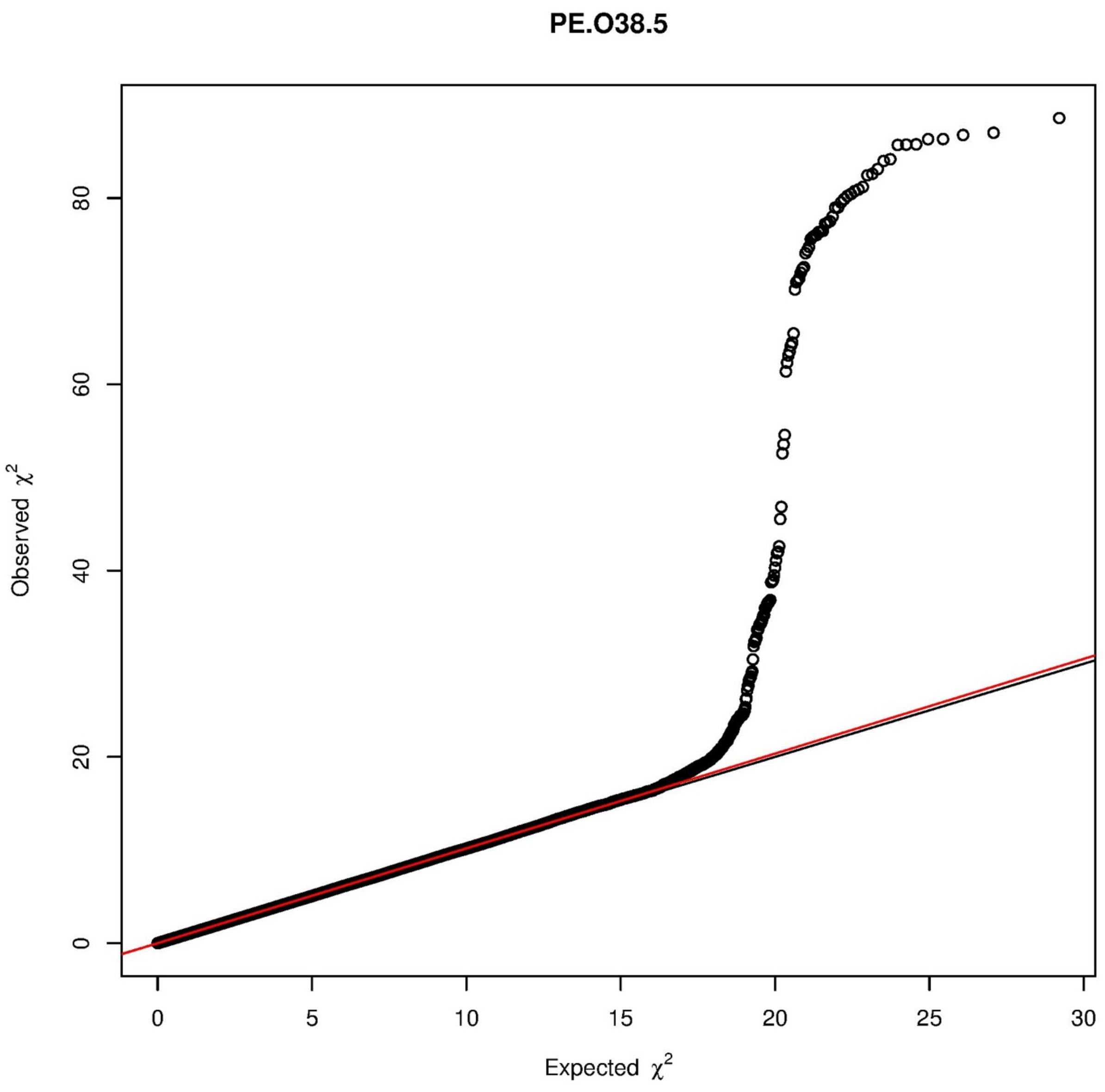

PE.O38.5
Observed $\chi^2$
Expected $\chi^2$
0
20
40
60
80
0
5
10
15
20
25
30

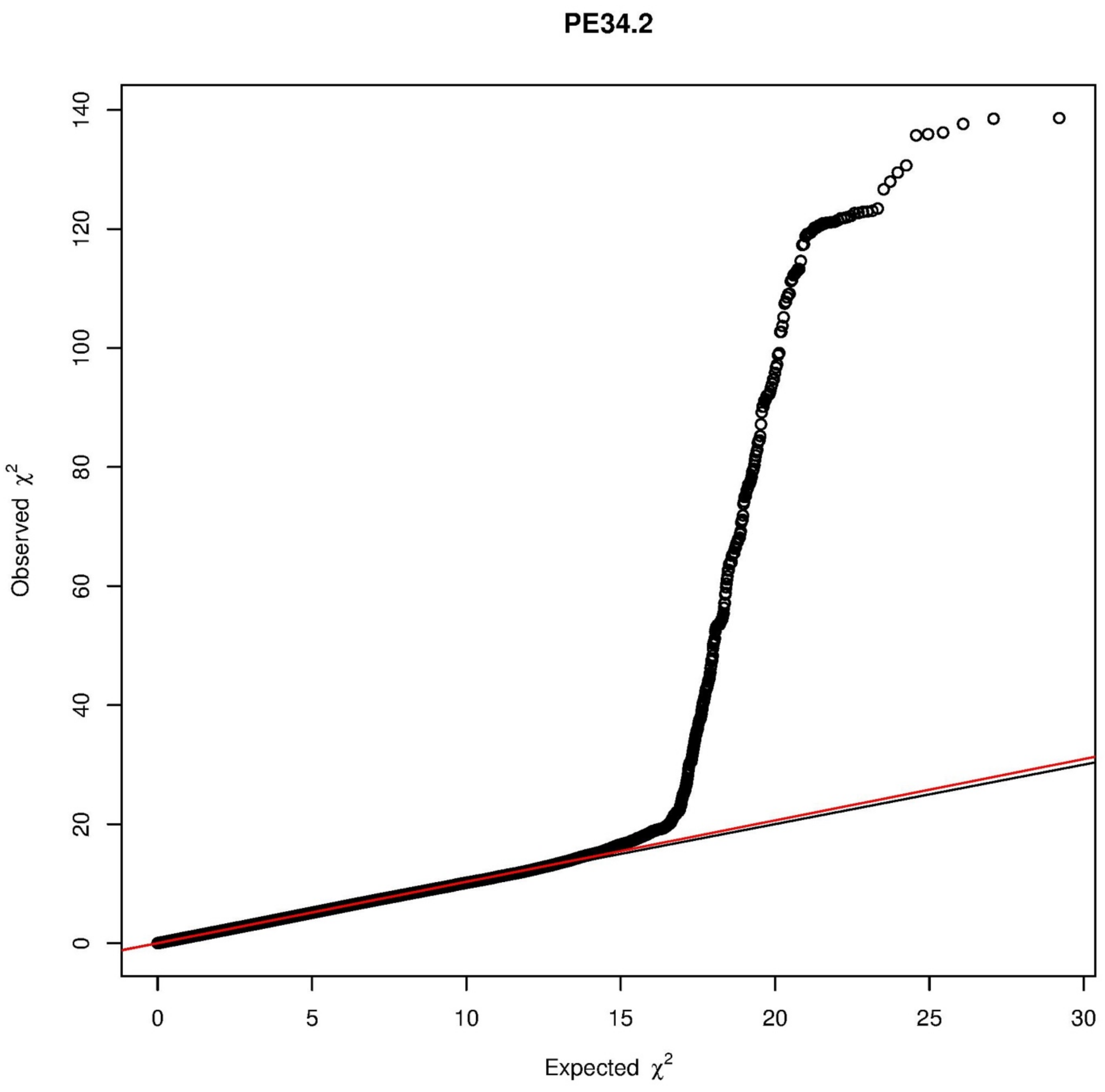
PE34.2
Observed χ2
Expected χ2
0
20
40
60
80
100
120
140
0
5
10
15
20
25
30

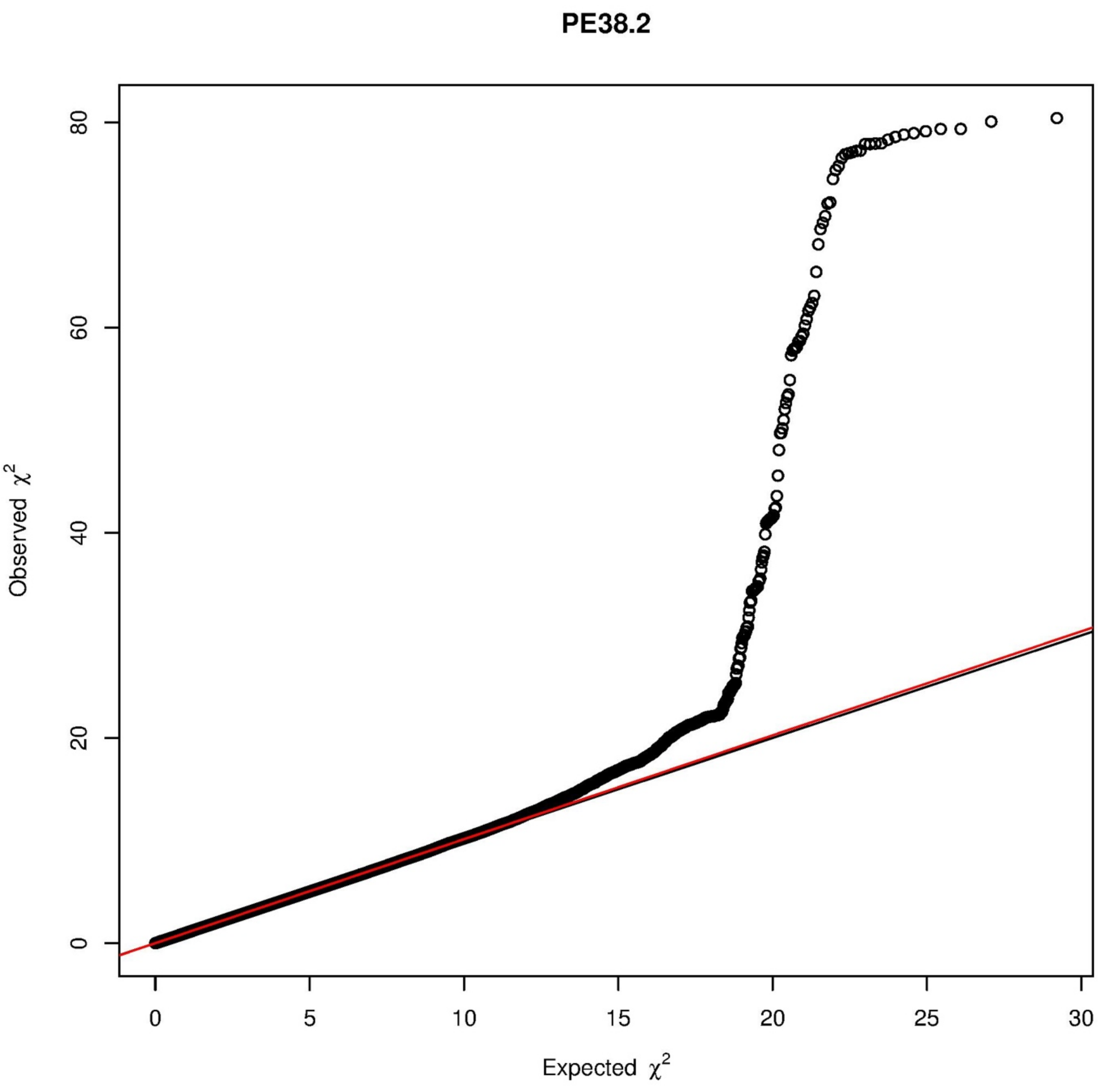

PE38.2
Observed $\chi^2$
0
20
40
60
80
Expected $\chi^2$
0
5
10
15
20
25
30

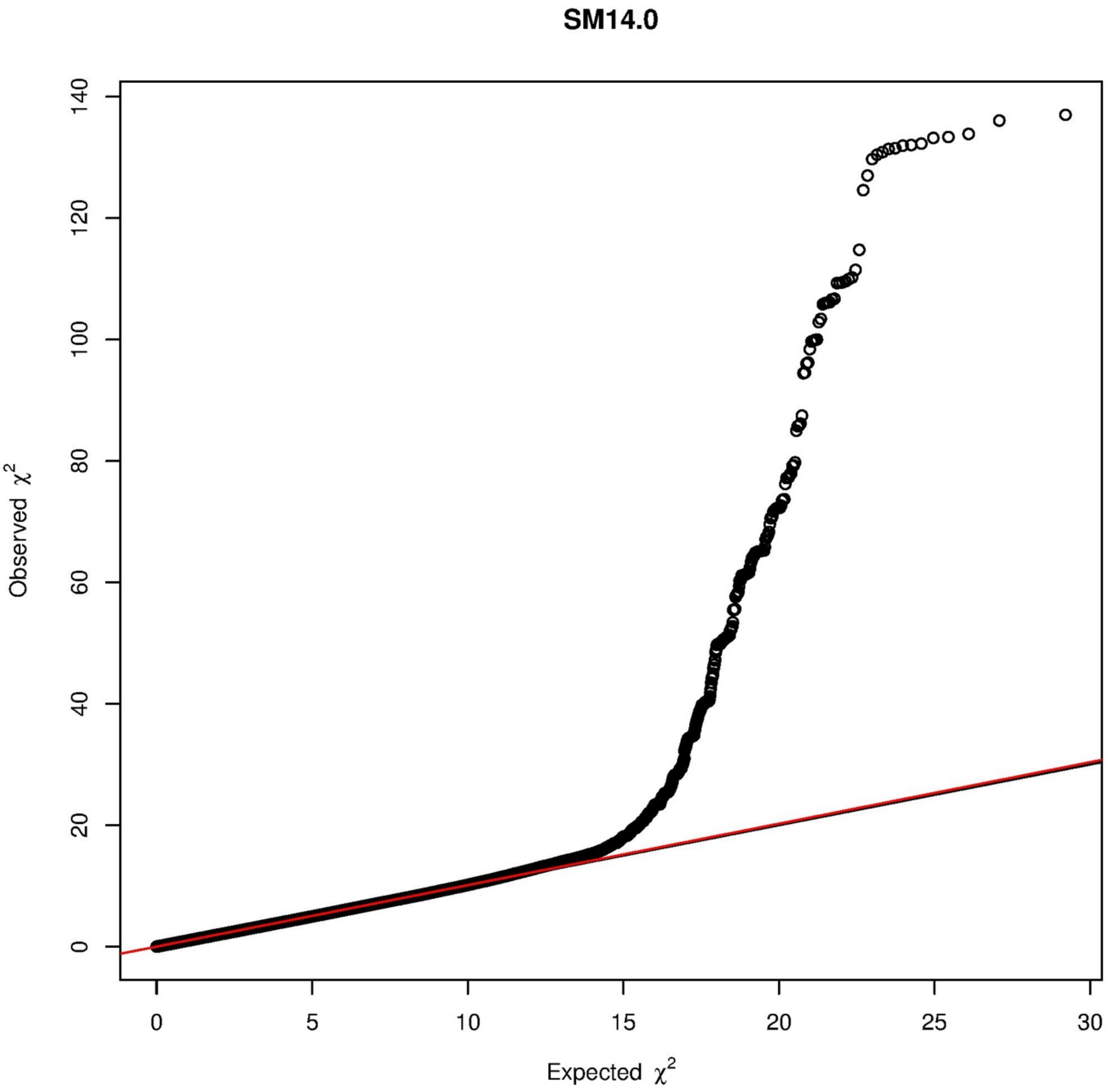

SM14.0
Observed $\chi^2$
Expected $\chi^2$
0
20
40
60
80
100
120
140
0
5
10
15
20
25
30

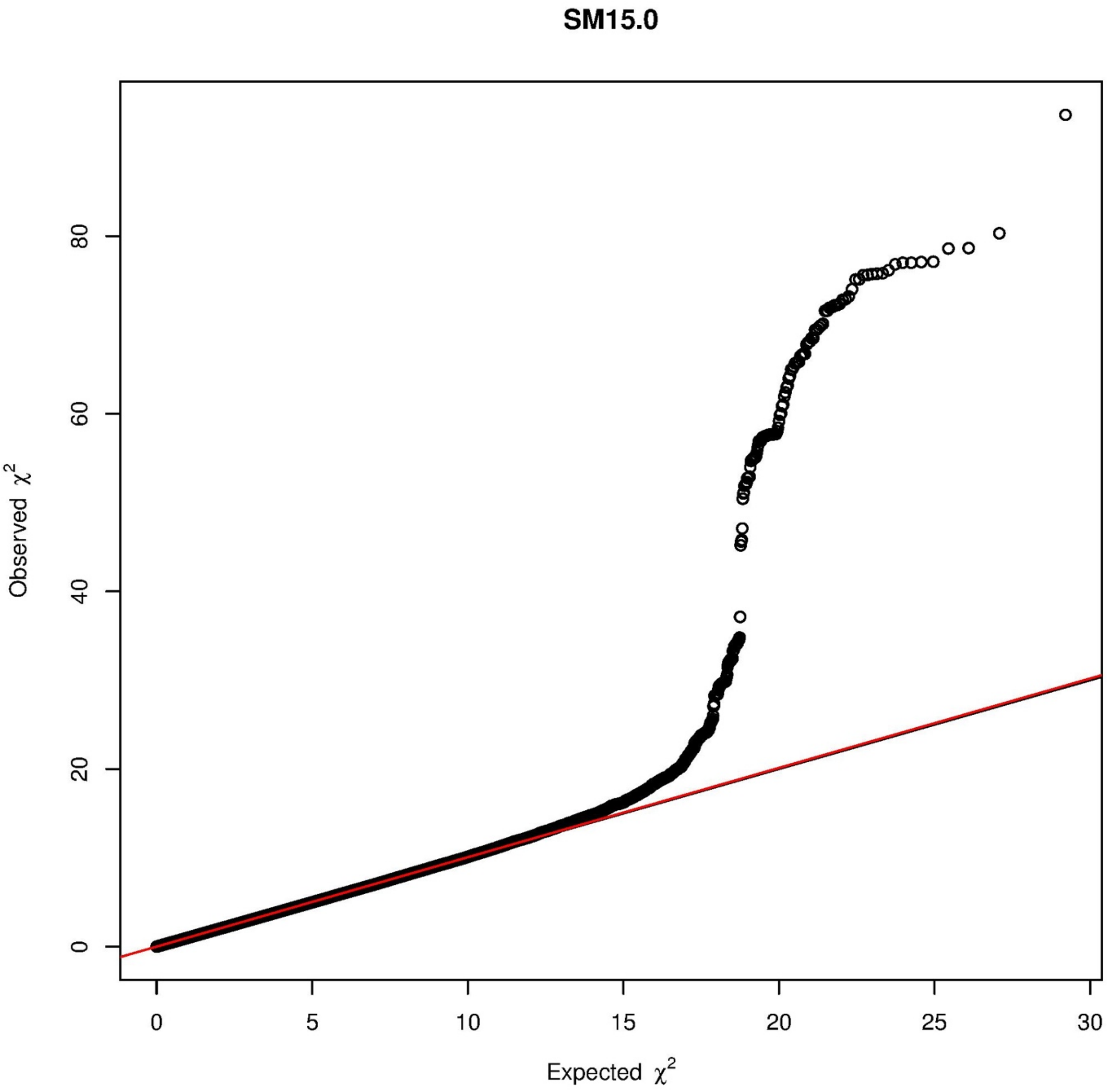

SM15.0
Observed $\chi^2$
Expected $\chi^2$
0
20
40
60
80
0
5
10
15
20
25
30

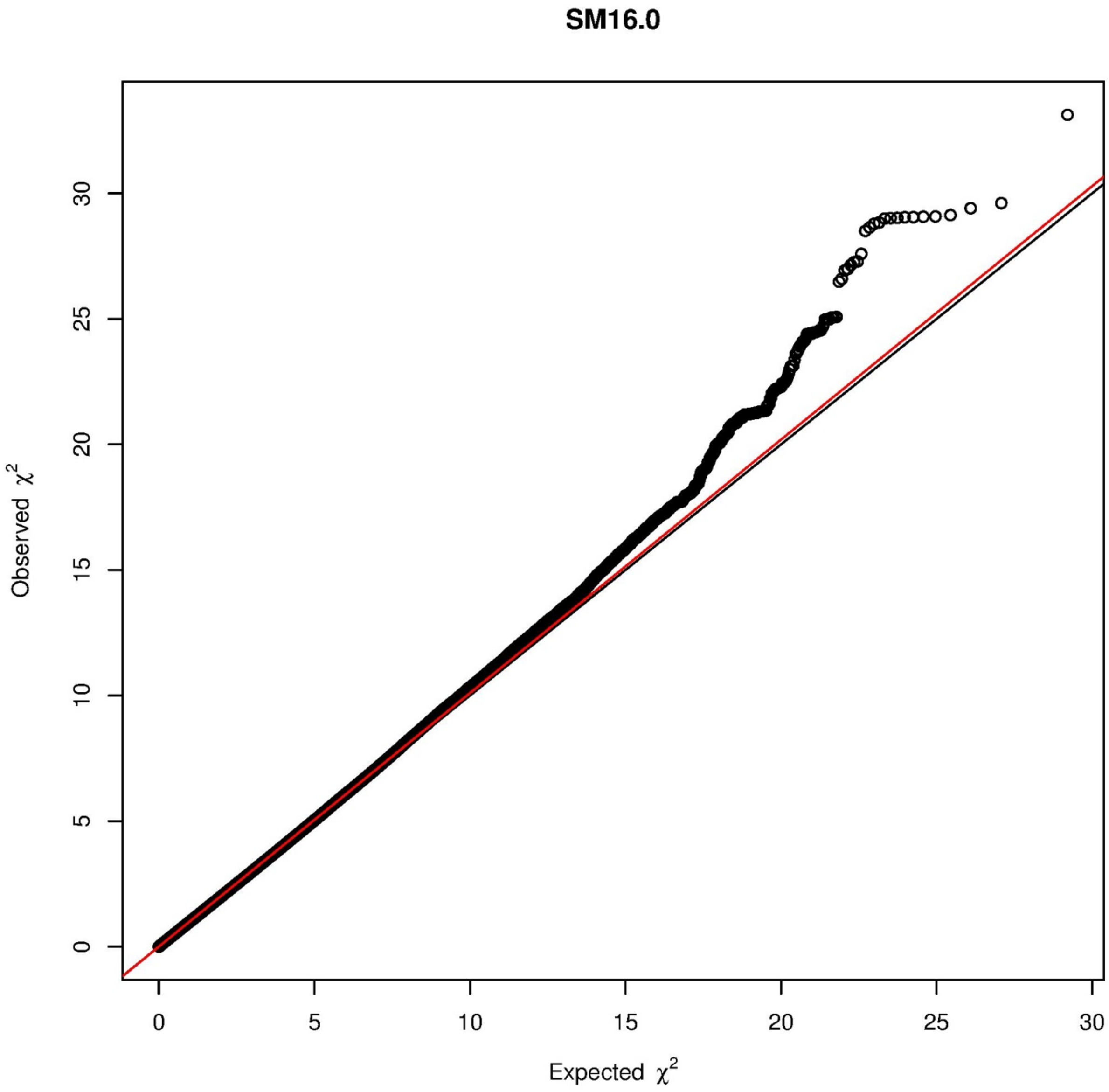
SM16.0
Observed $\chi^2$
Expected $\chi^2$
0
5
10
15
20
25
30

SM16.1

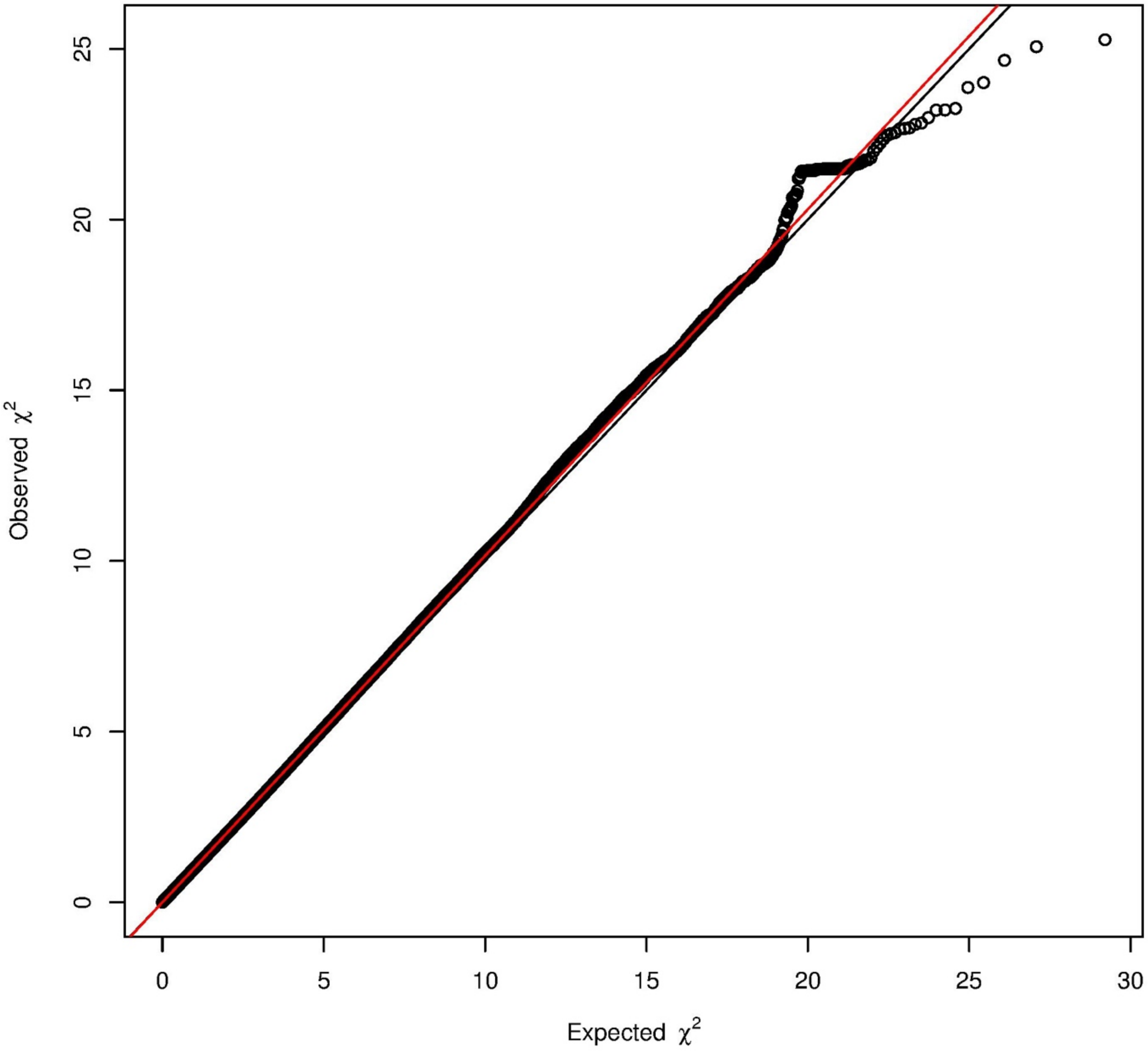

## SM17.0

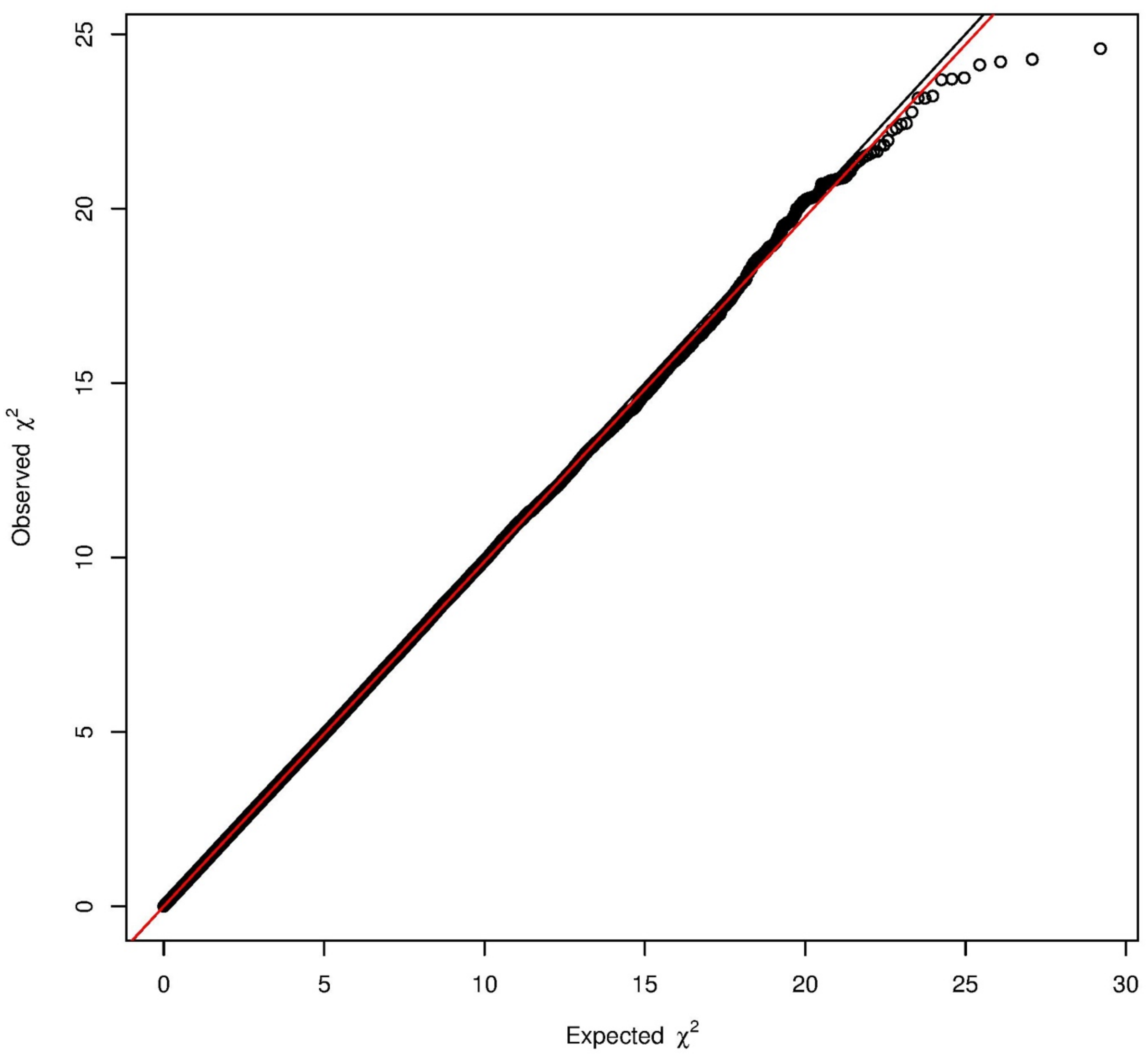

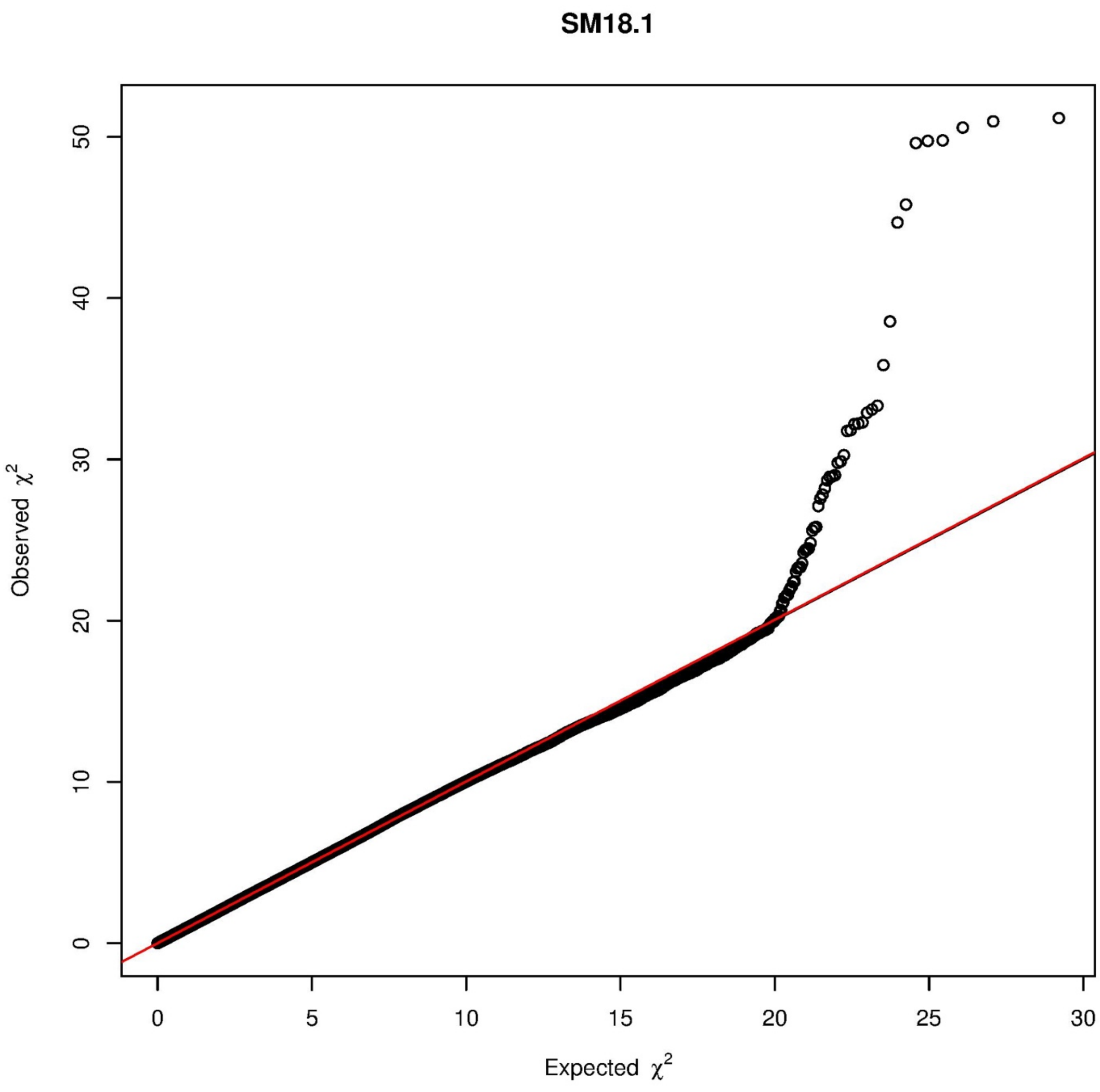

SM18.1
Observed $\chi^2$
Expected $\chi^2$
0
10
20
30
40
50
0
5
10
15
20
25
30

## SM18.2

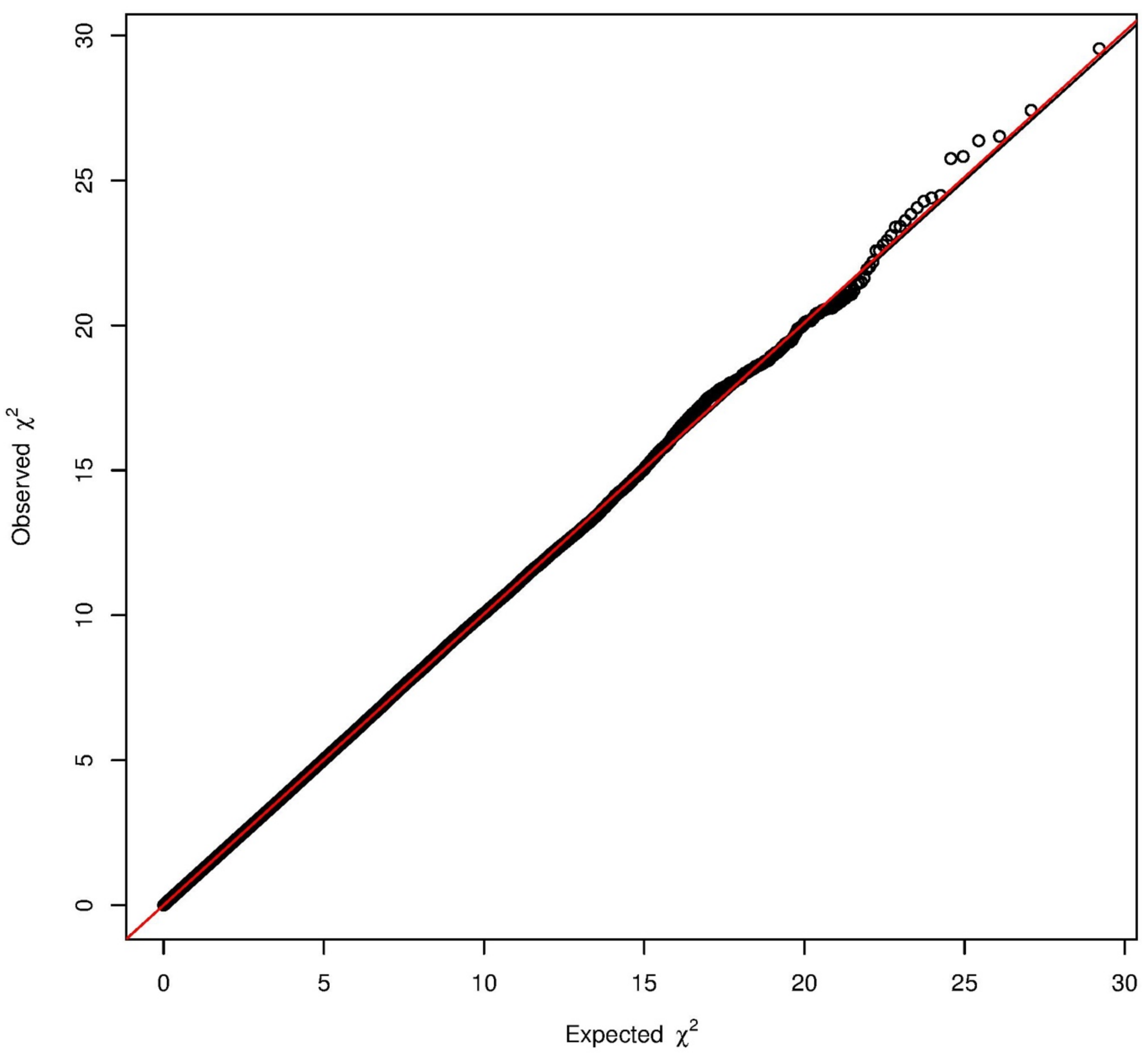

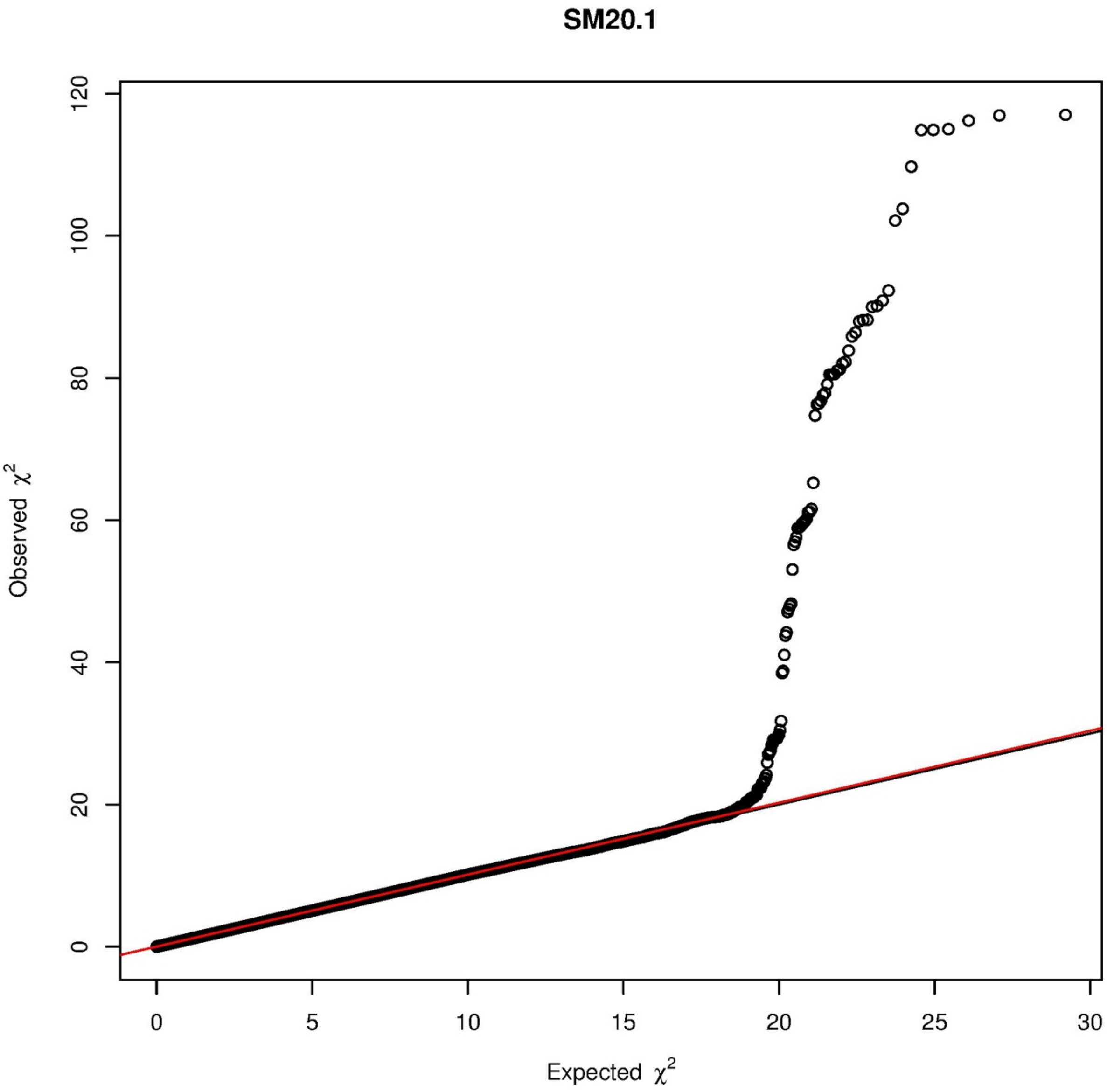

SM20.1
Observed $\chi^2$
Expected $\chi^2$
0
20
40
60
80
100
120
0
5
10
15
20
25
30

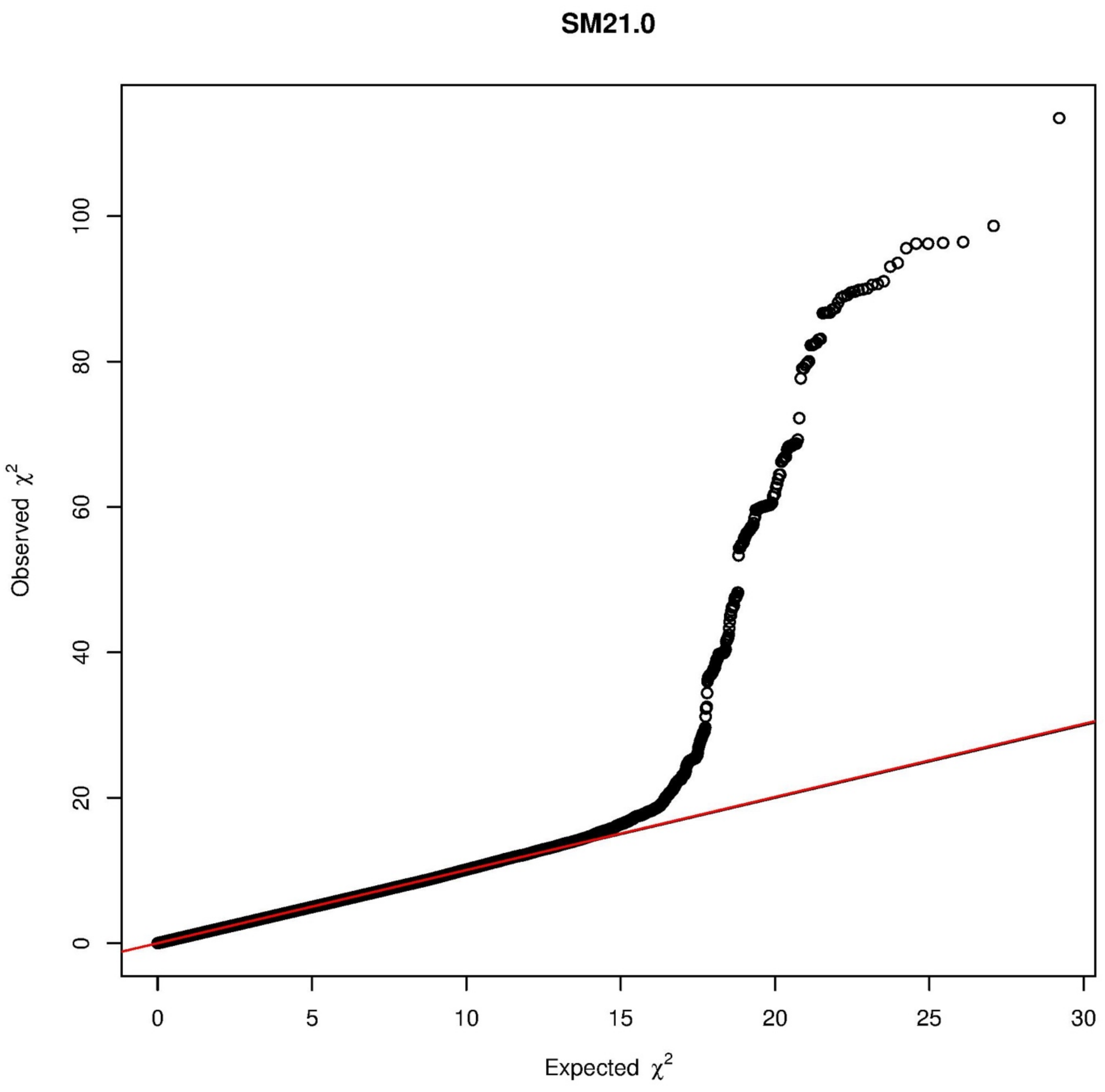
SM21.0
Observed $\chi^2$
Expected $\chi^2$
0
20
40
60
80
100
0
5
10
15
20
25
30

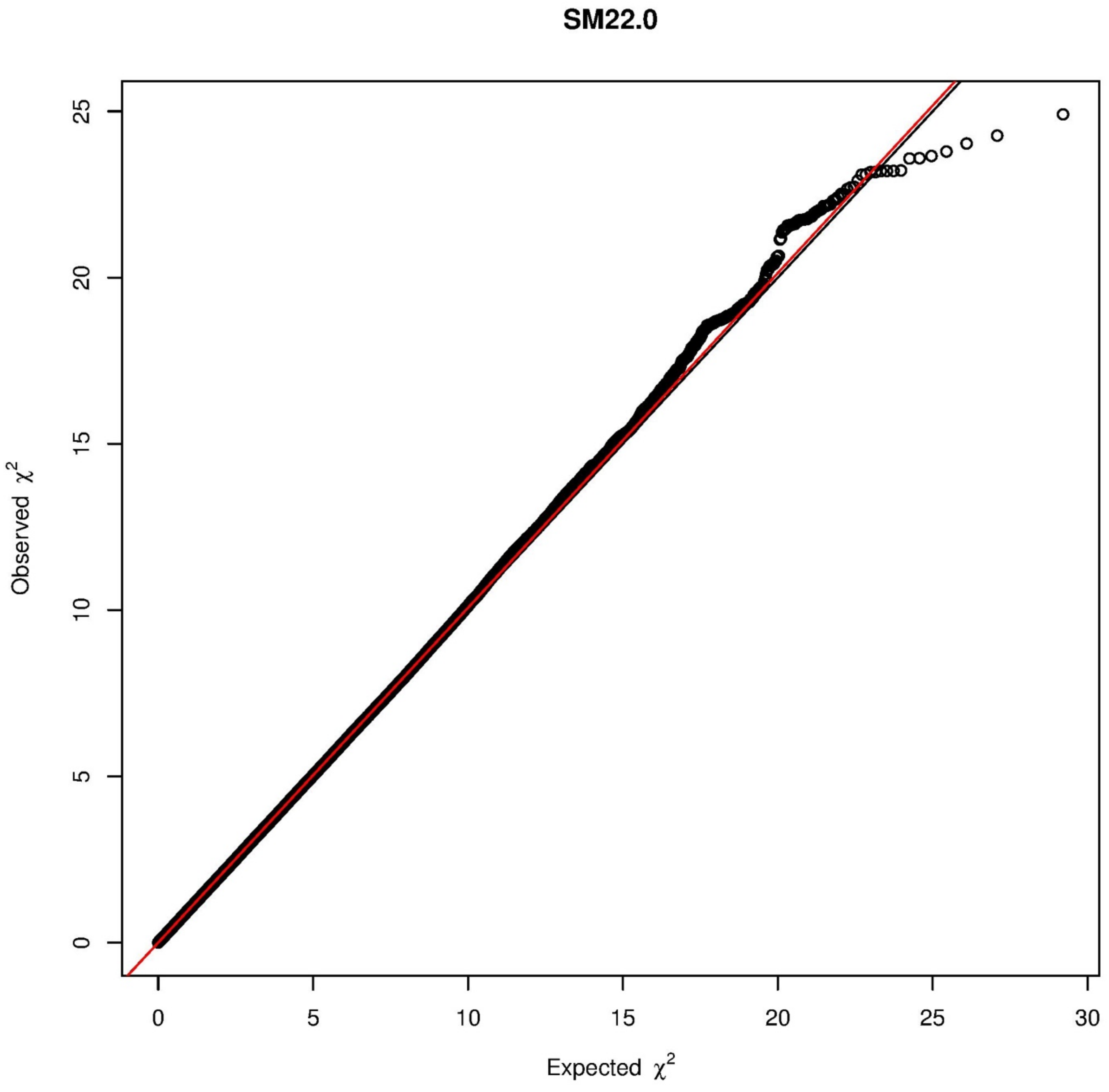

SM22.0
Observed $\chi^2$
Expected $\chi^2$
0
5
10
15
20
25
30

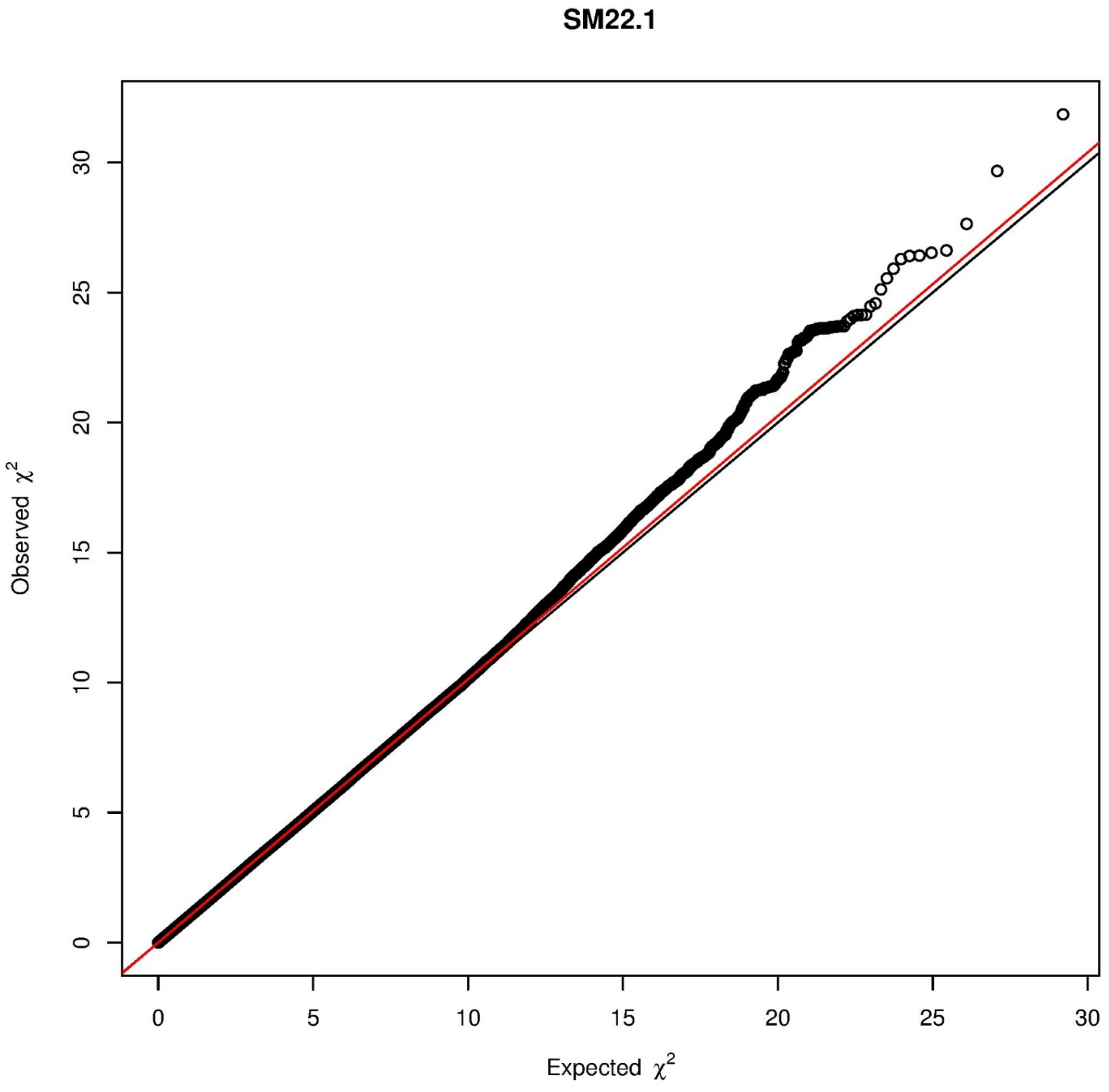
SM22.1
Observed χ2
Expected χ2
0
5
10
15
20
25
30

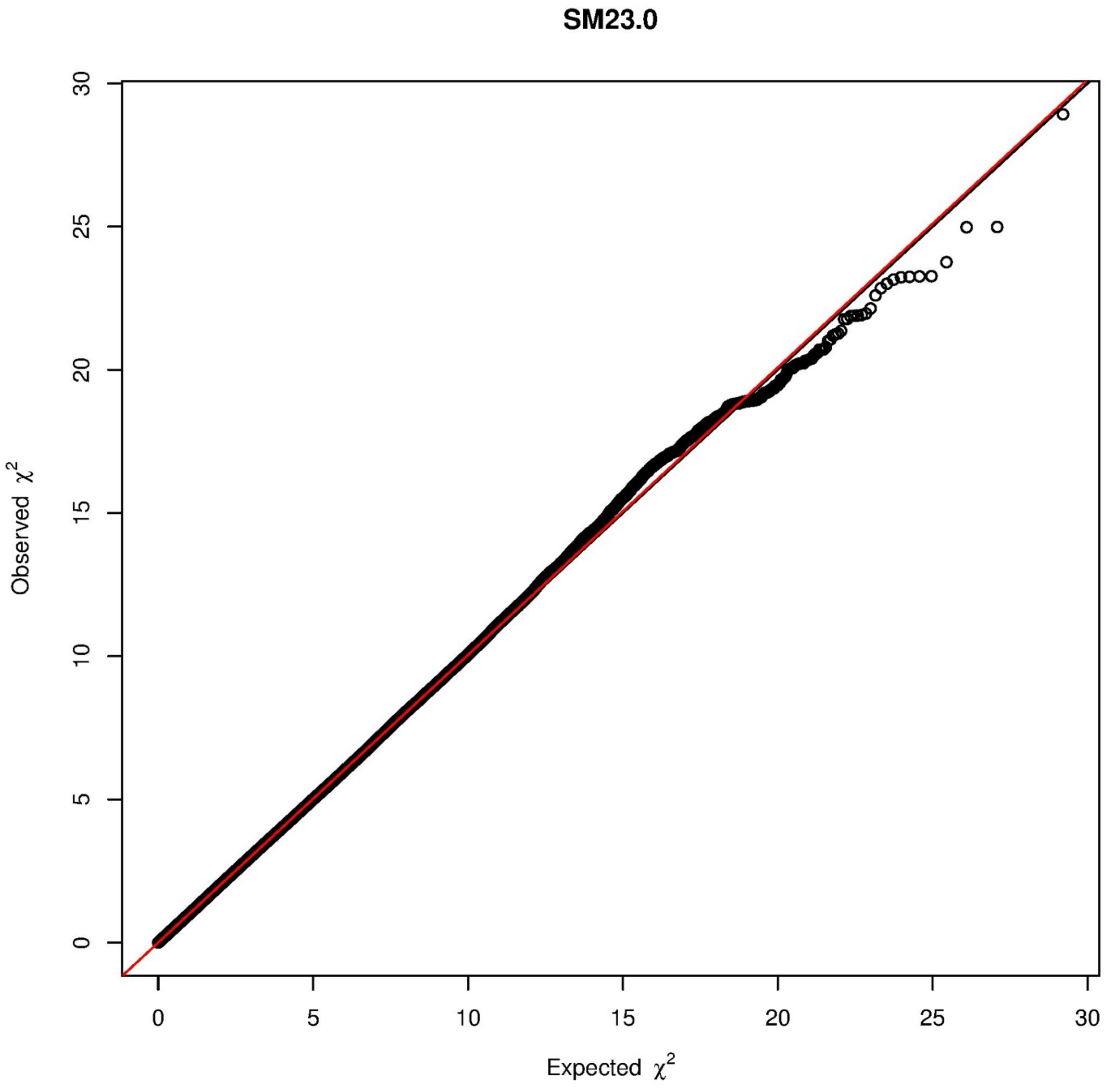
SM23.0
Observed χ2
Expected χ2
0
5
10
15
20
25
30

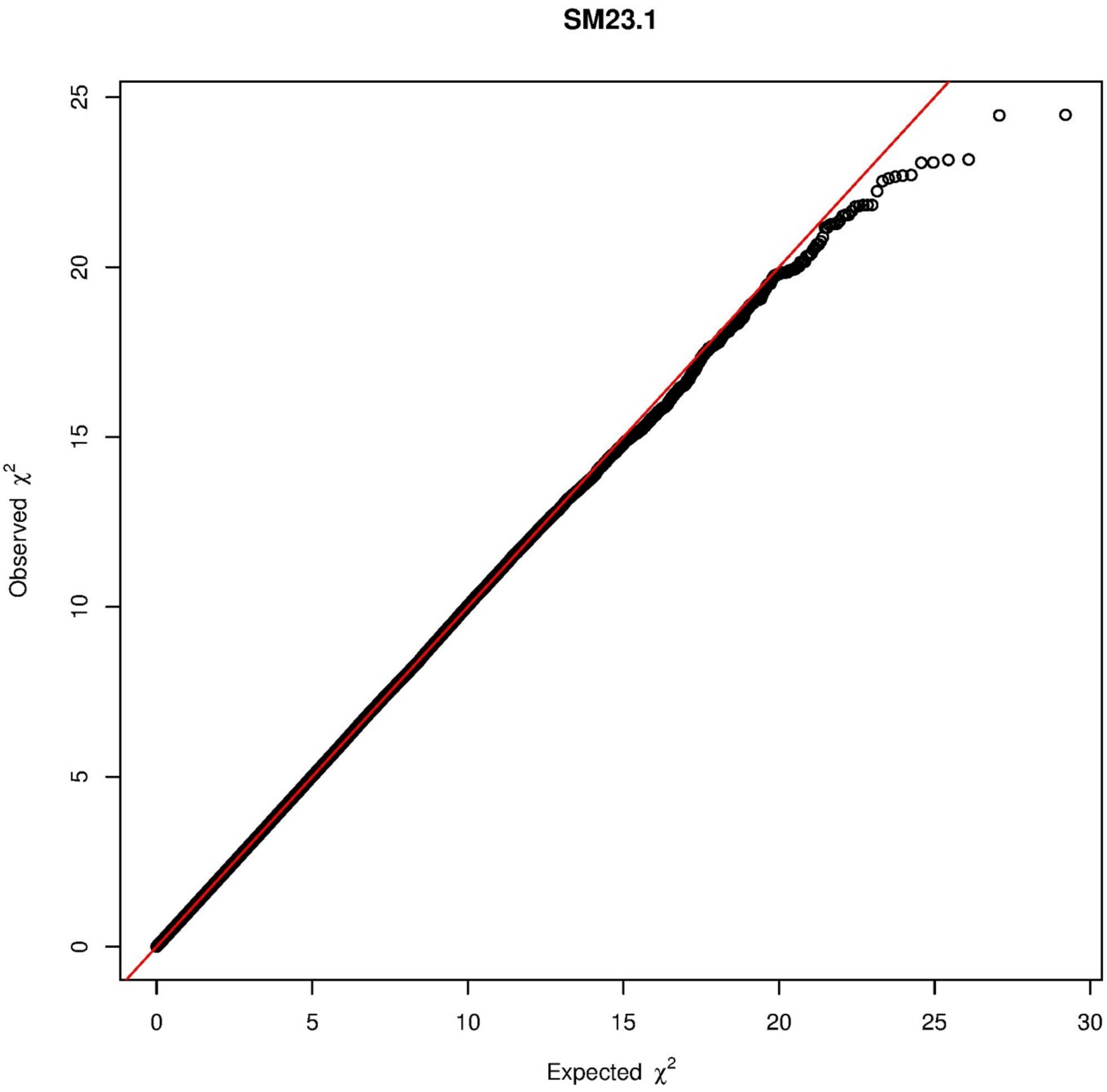

SM23.1
Observed $\chi^2$
Expected $\chi^2$
0
5
10
15
20
25
30

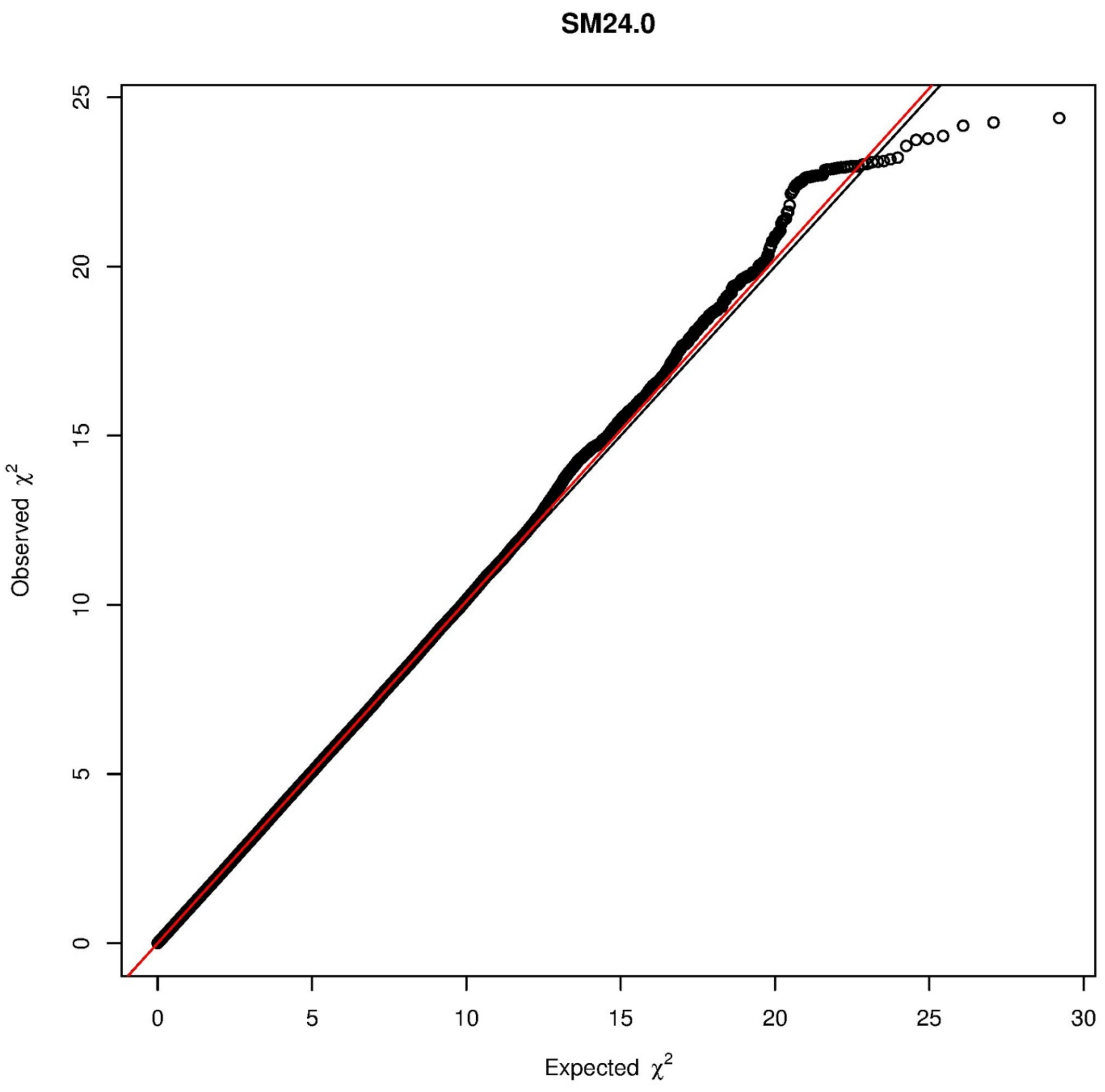

SM24.0
Observed $\chi^2$
Expected $\chi^2$
0
5
10
15
20
25
30

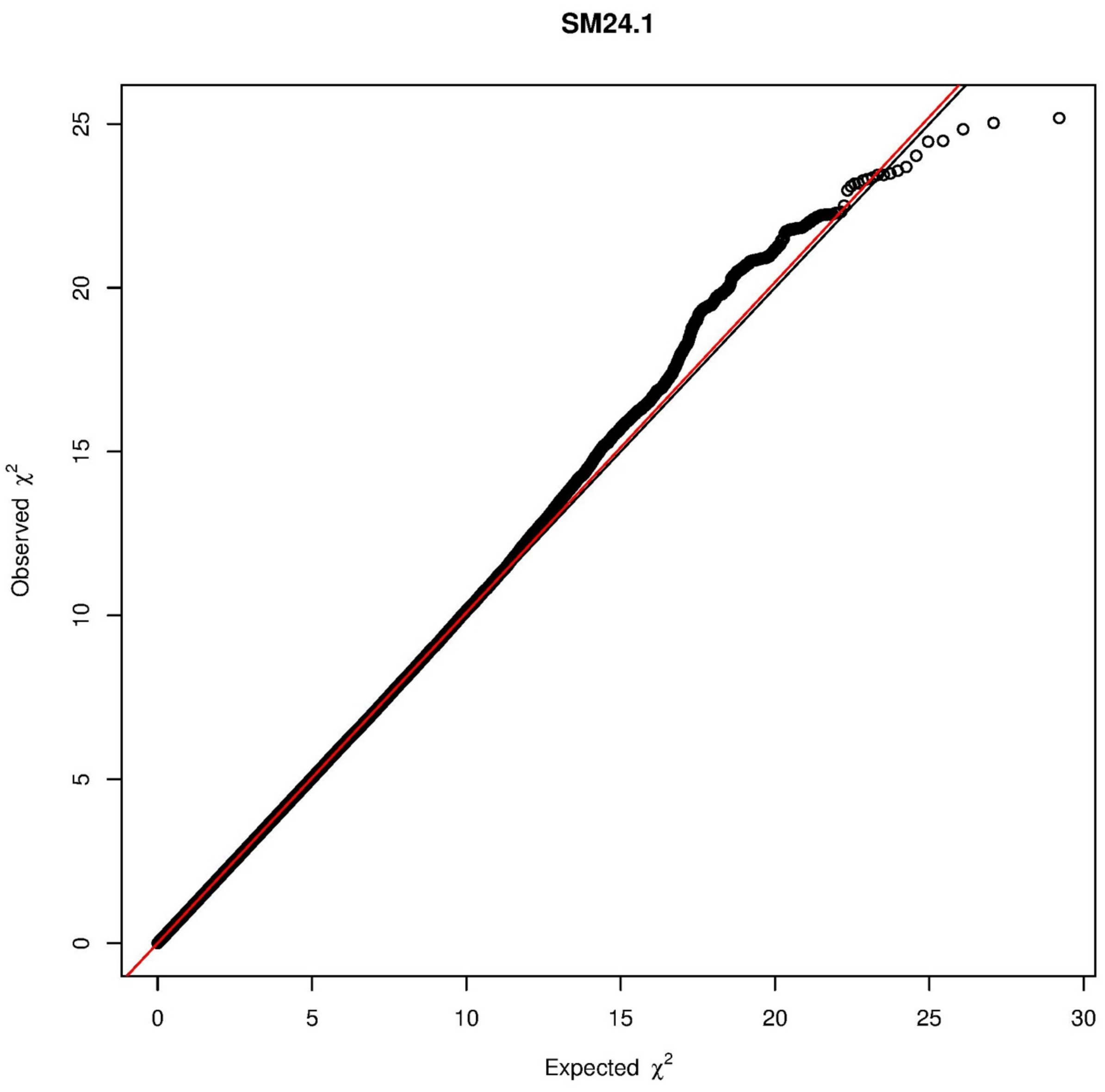

SM24.1
Observed $\chi^2$
Expected $\chi^2$
0
5
10
15
20
25
30

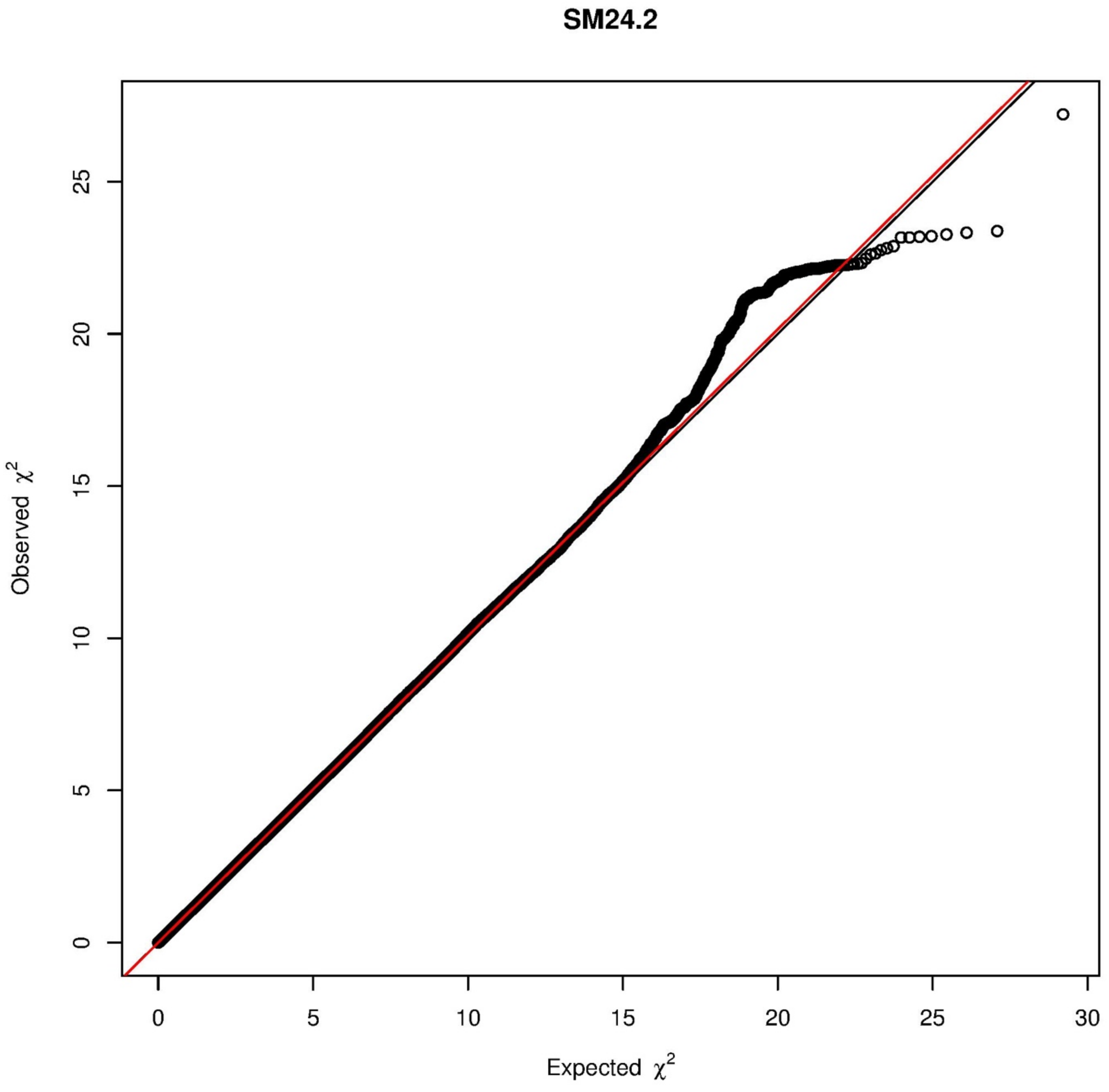
SM24.2
Observed χ²
Expected χ²
0
5
10
15
20
25
30

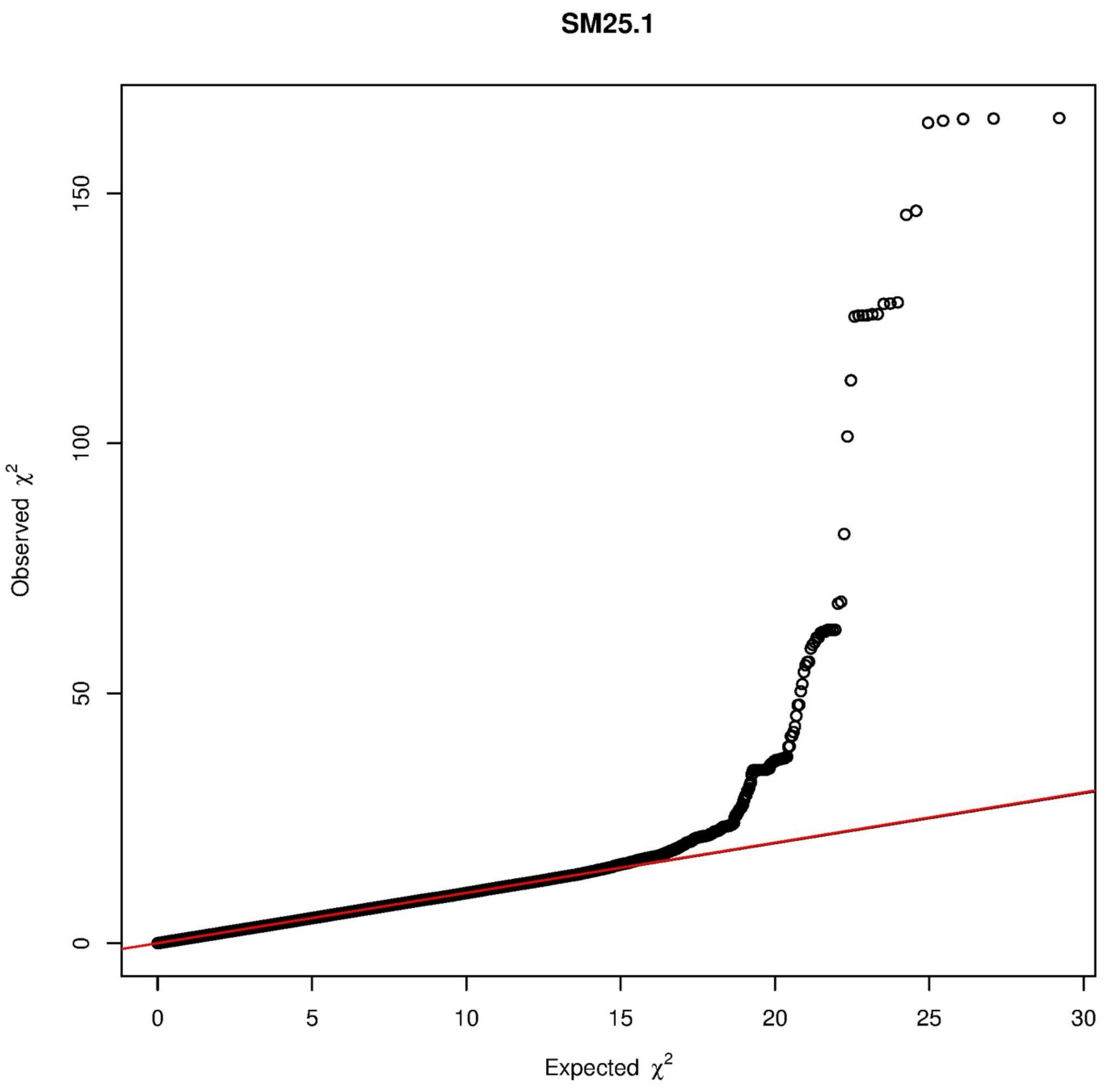

SM25.1
Observed $\chi^2$
Expected $\chi^2$
0
50
100
150
0
5
10
15
20
25
30

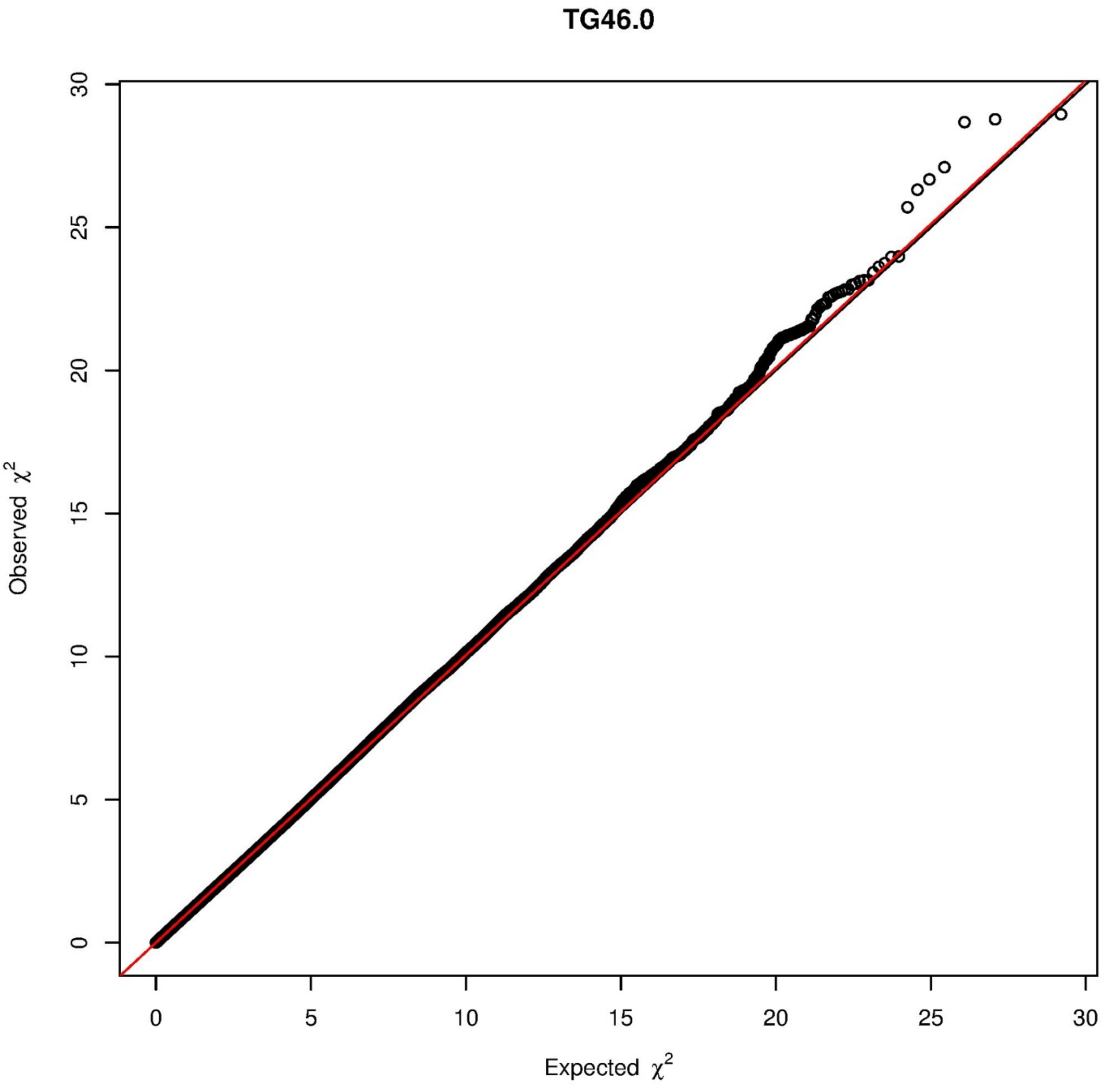

TG46.0
Observed $\chi^2$
Expected $\chi^2$
0
5
10
15
20
25
30
0
5
10
15
20
25
30

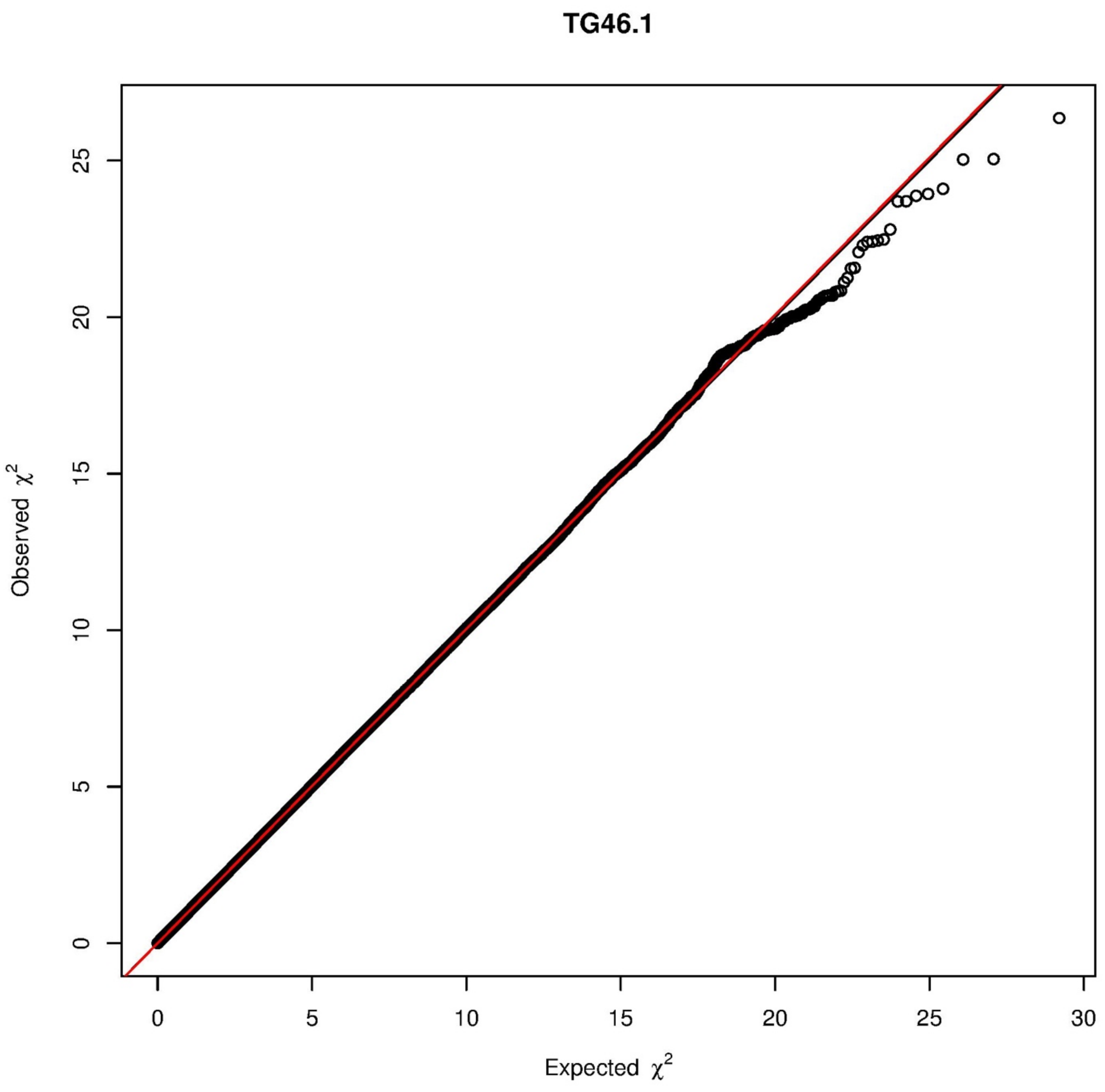

TG46.1
Observed $\chi^2$
Expected $\chi^2$
0
5
10
15
20
25
30

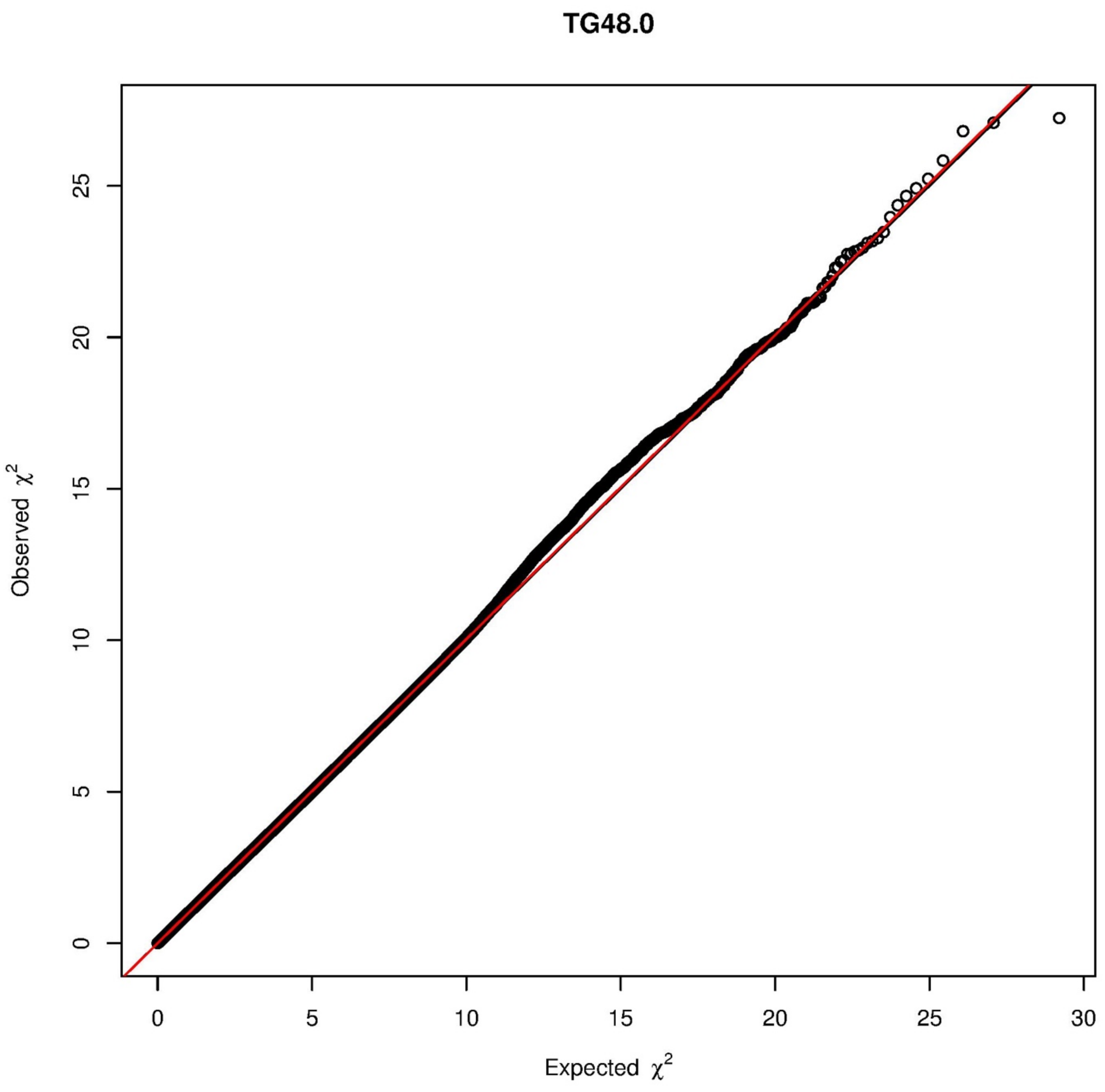
TG48.0
Observed $\chi^2$
Expected $\chi^2$
0
5
10
15
20
25
30

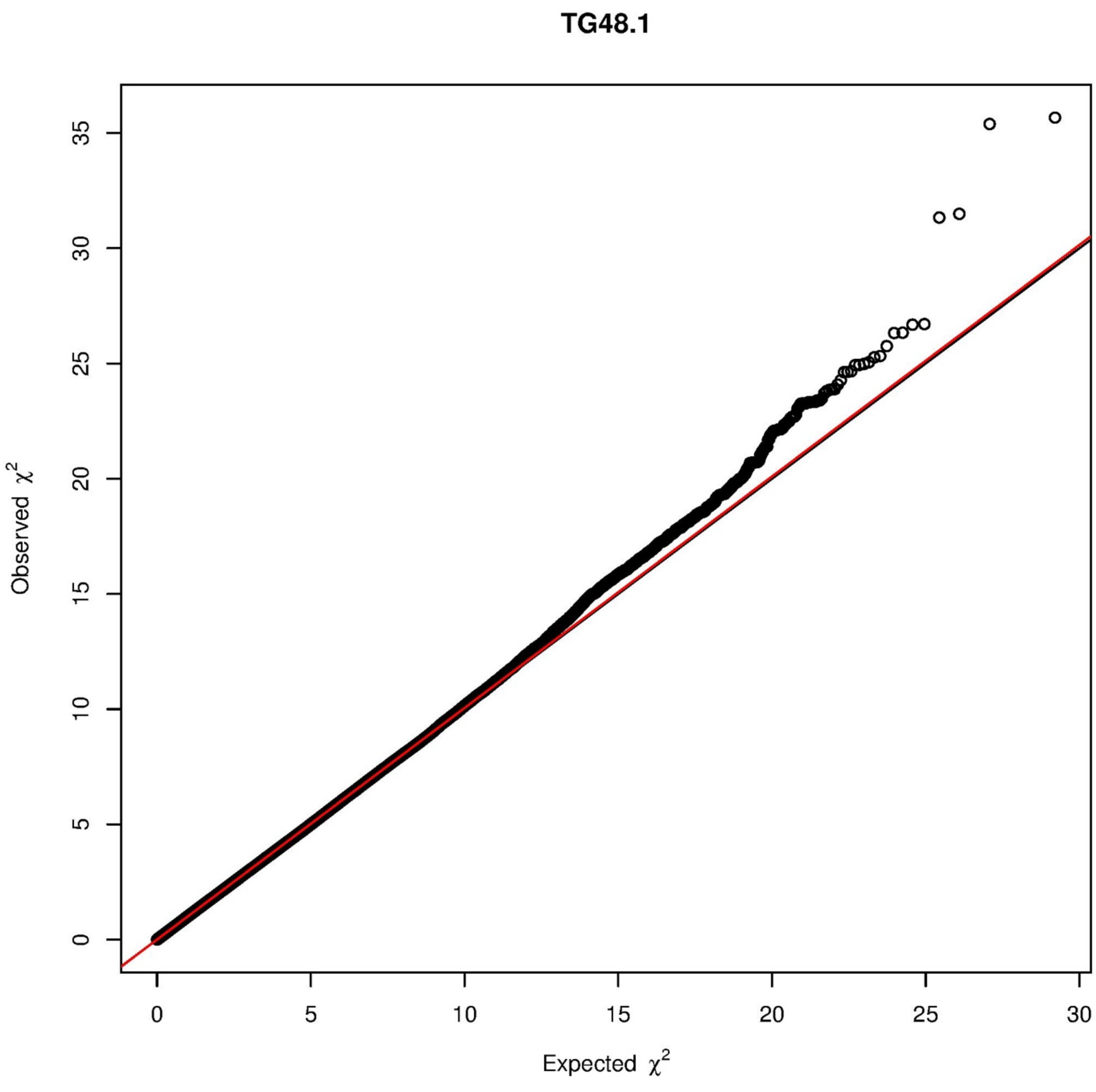

TG48.1
Observed χ2
Expected χ2
0
5
10
15
20
25
30
35

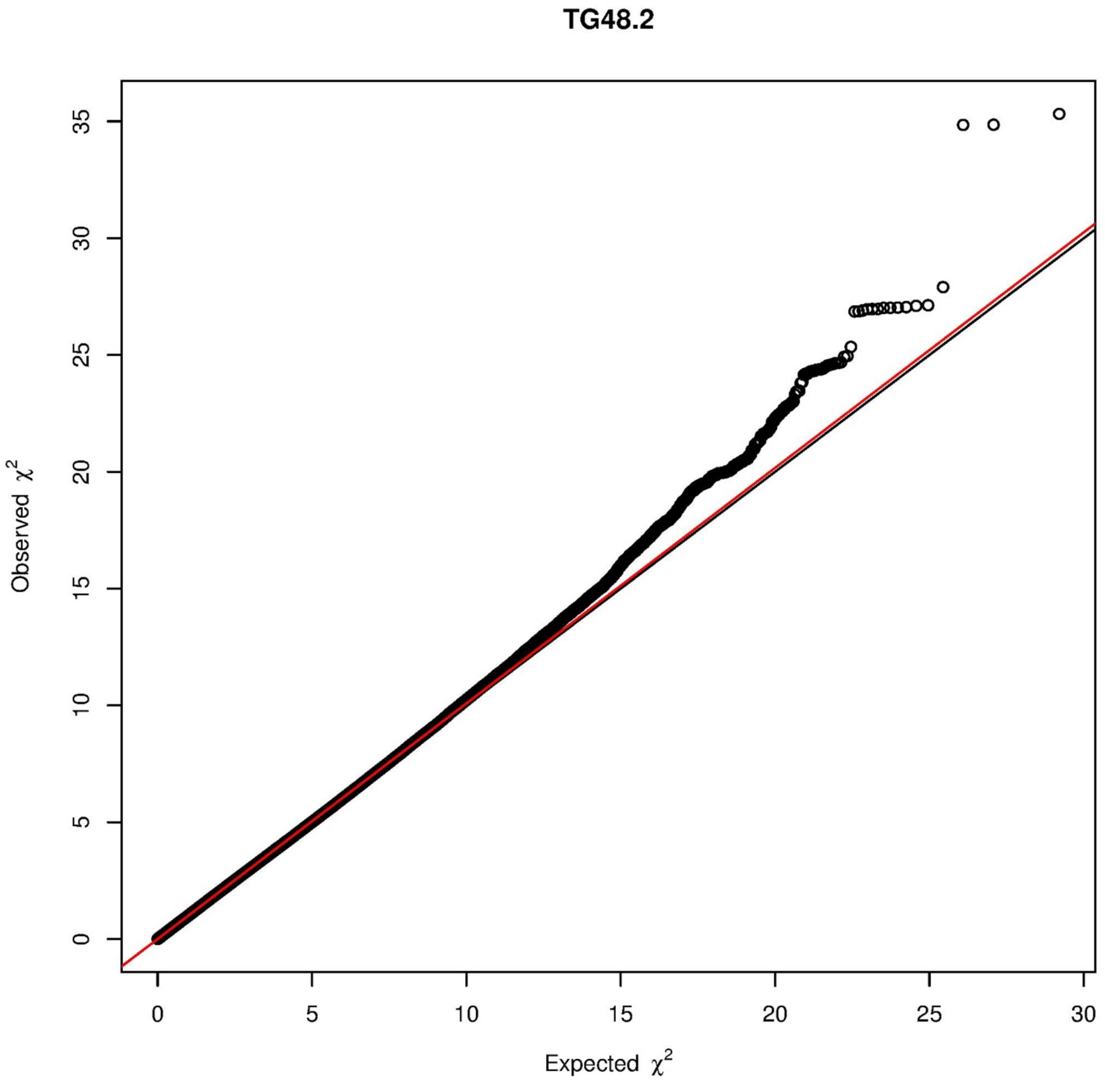

TG48.2
Observed $\chi^2$
Expected $\chi^2$
0
5
10
15
20
25
30
35

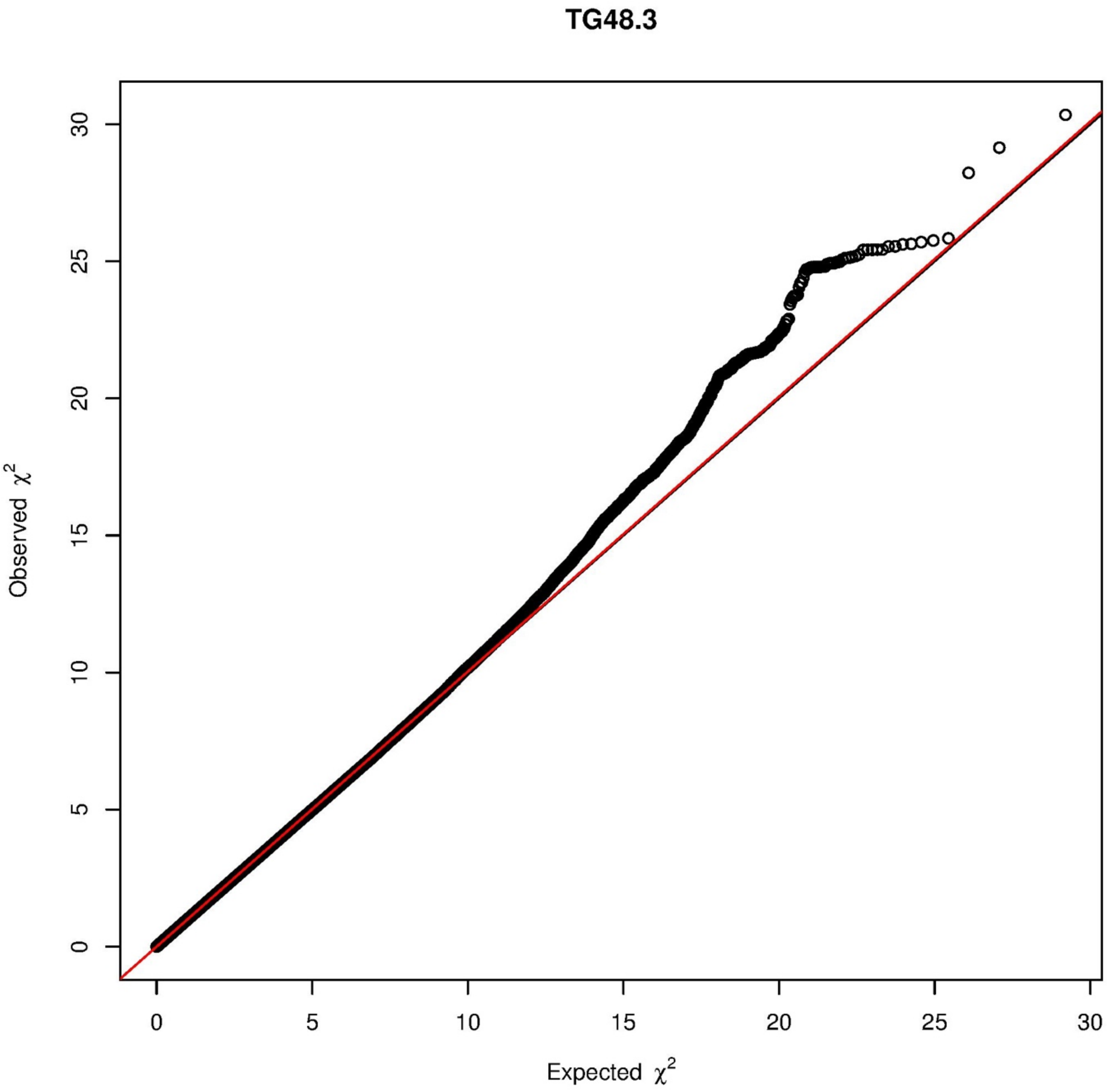

TG48.3
Observed $\chi^2$
Expected $\chi^2$
0
5
10
15
20
25
30

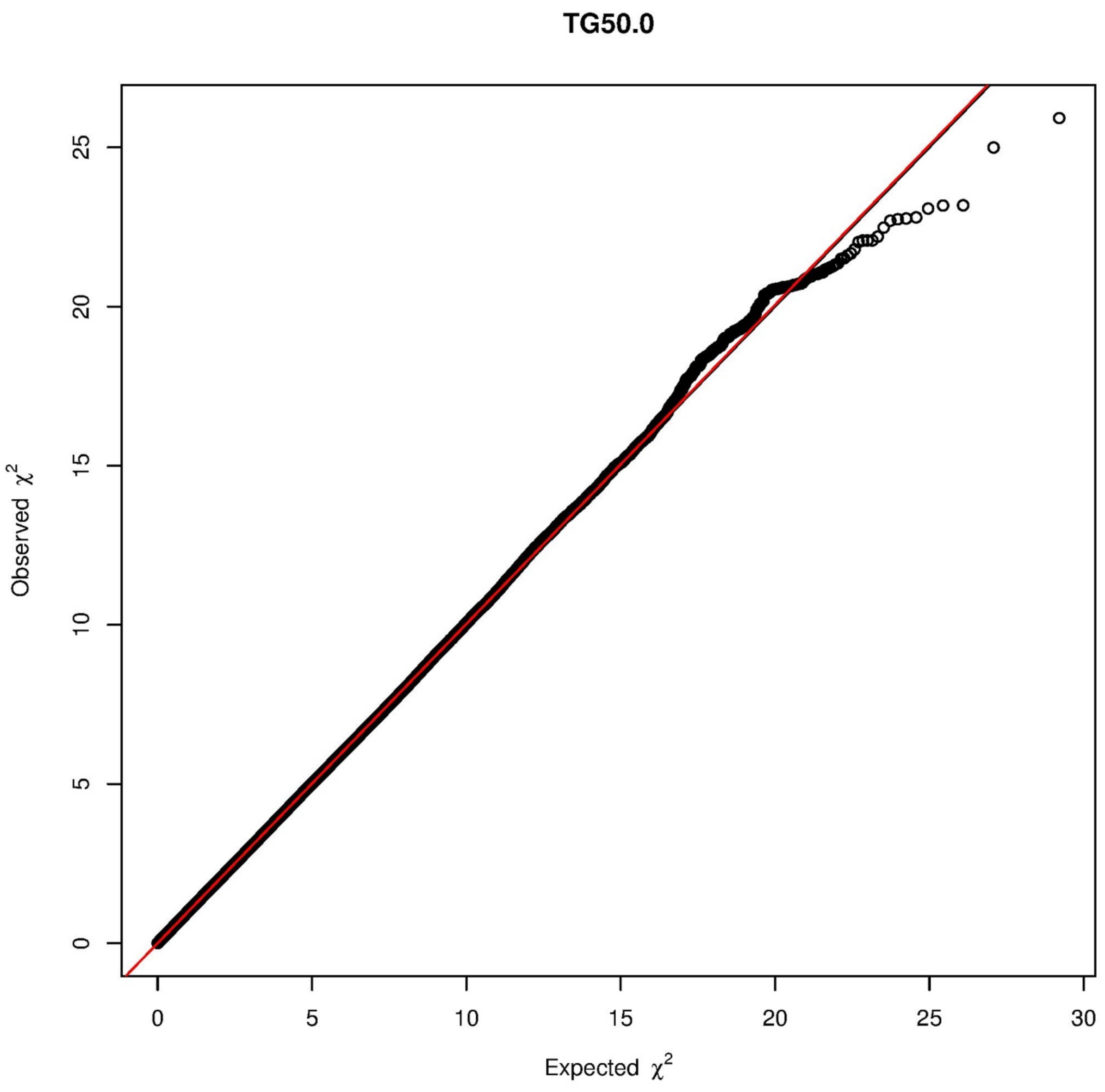

TG50.0
Observed $\chi^2$
Expected $\chi^2$
0
5
10
15
20
25
30

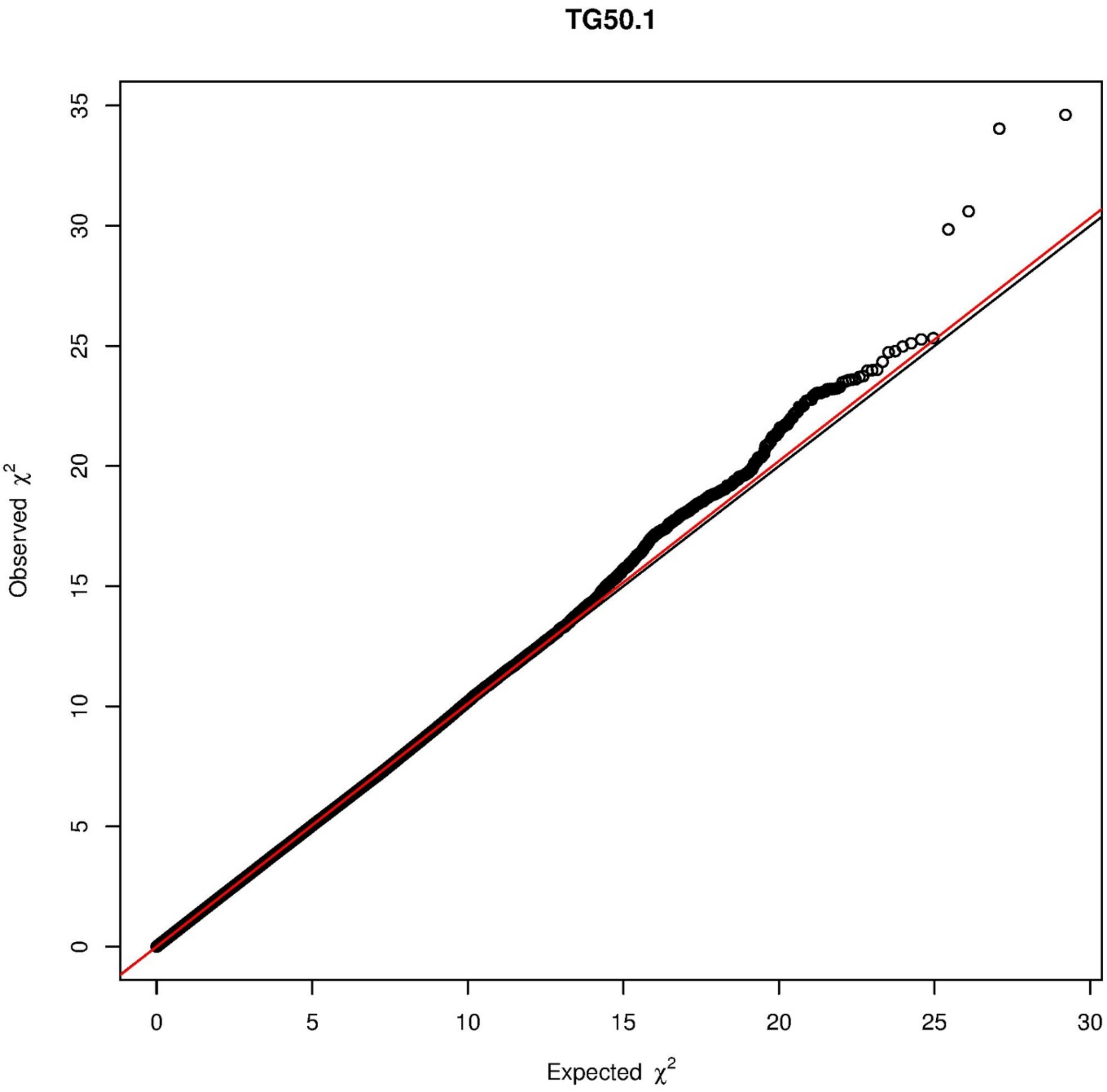

TG50.1
Observed $\chi^2$
Expected $\chi^2$
0
5
10
15
20
25
30
35

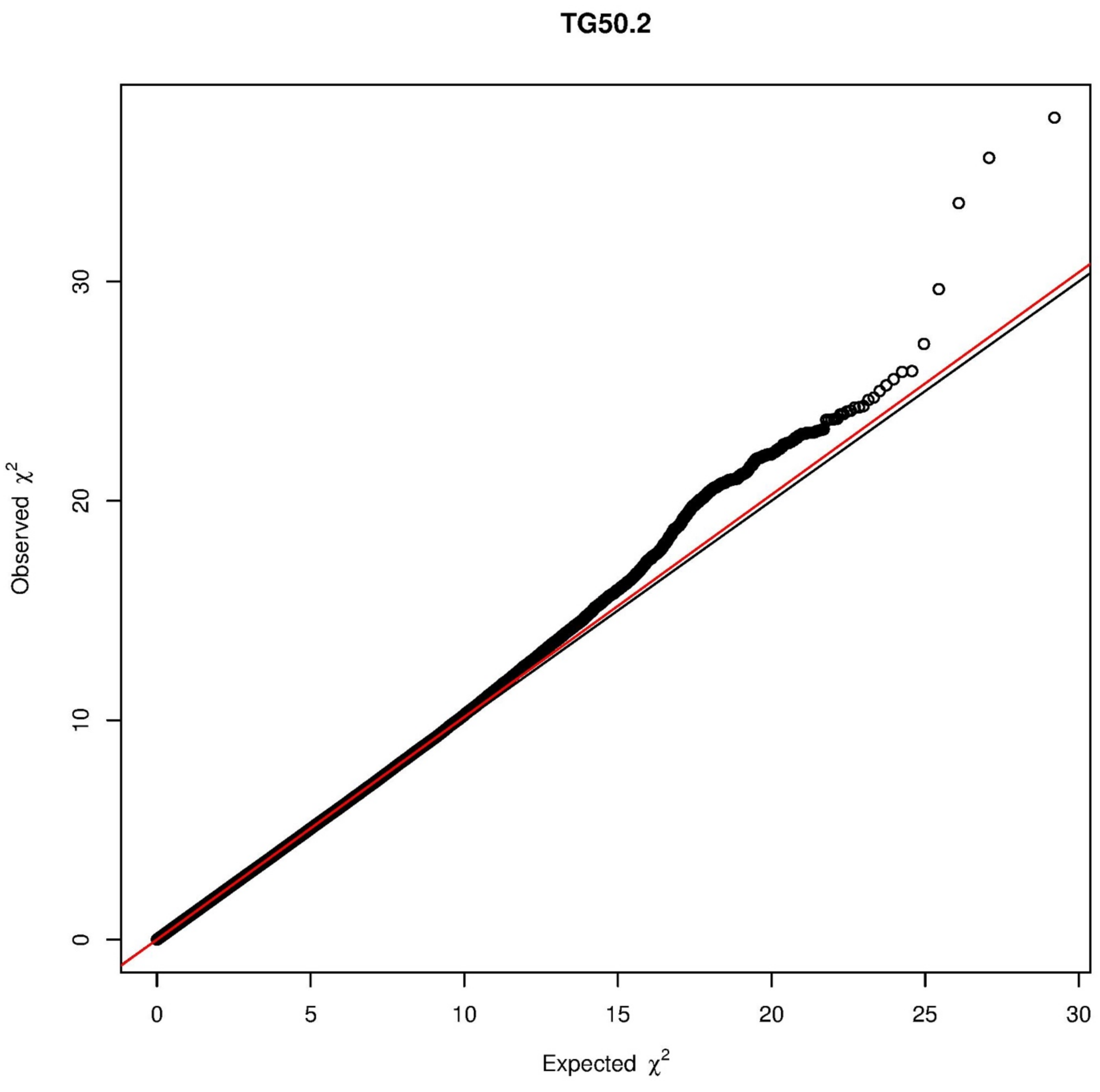

TG50.2
Observed χ2
Expected χ2
0
10
20
30
0
5
10
15
20
25
30

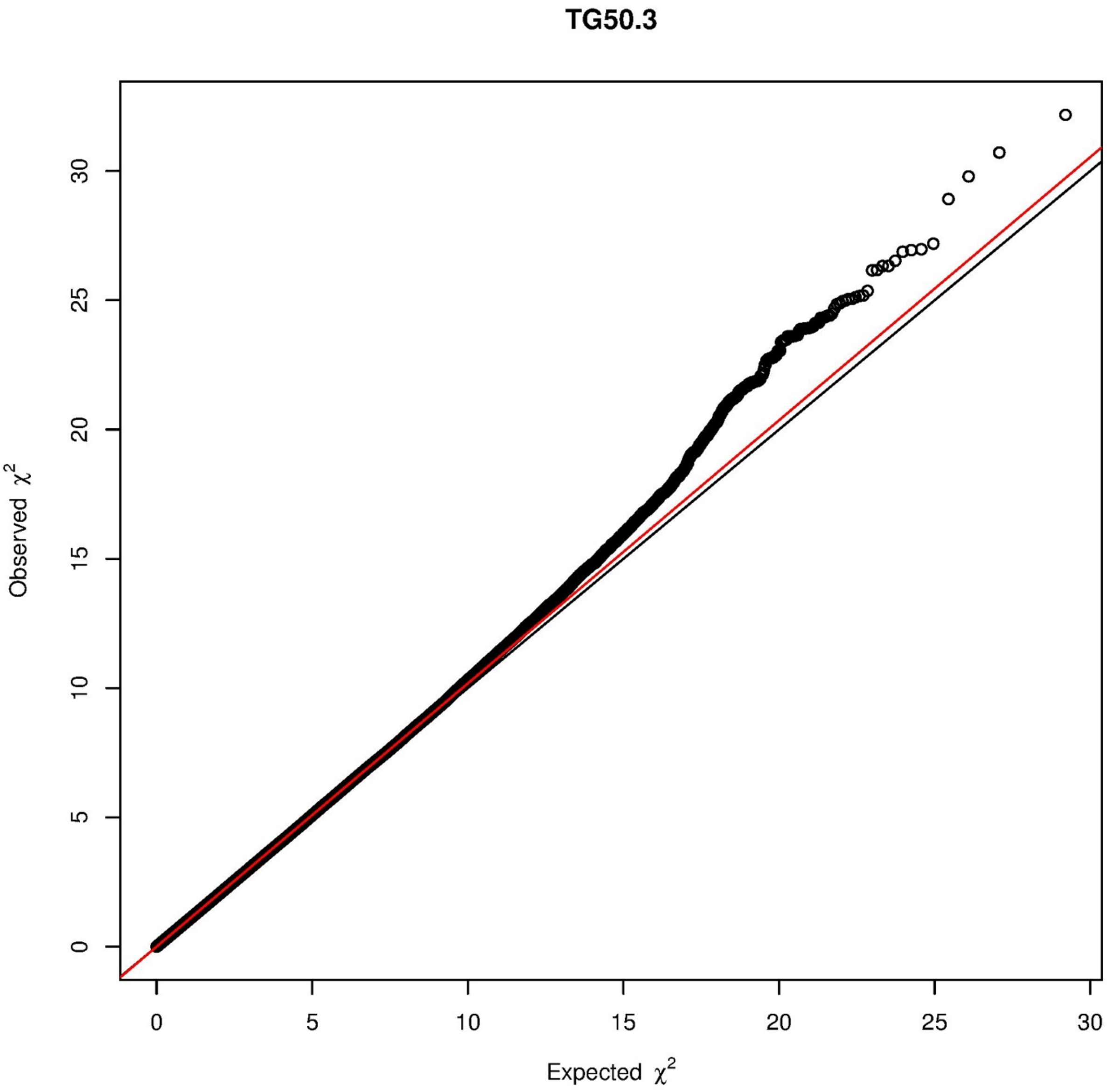

TG50.3
Observed $\chi^2$
Expected $\chi^2$
0
5
10
15
20
25
30

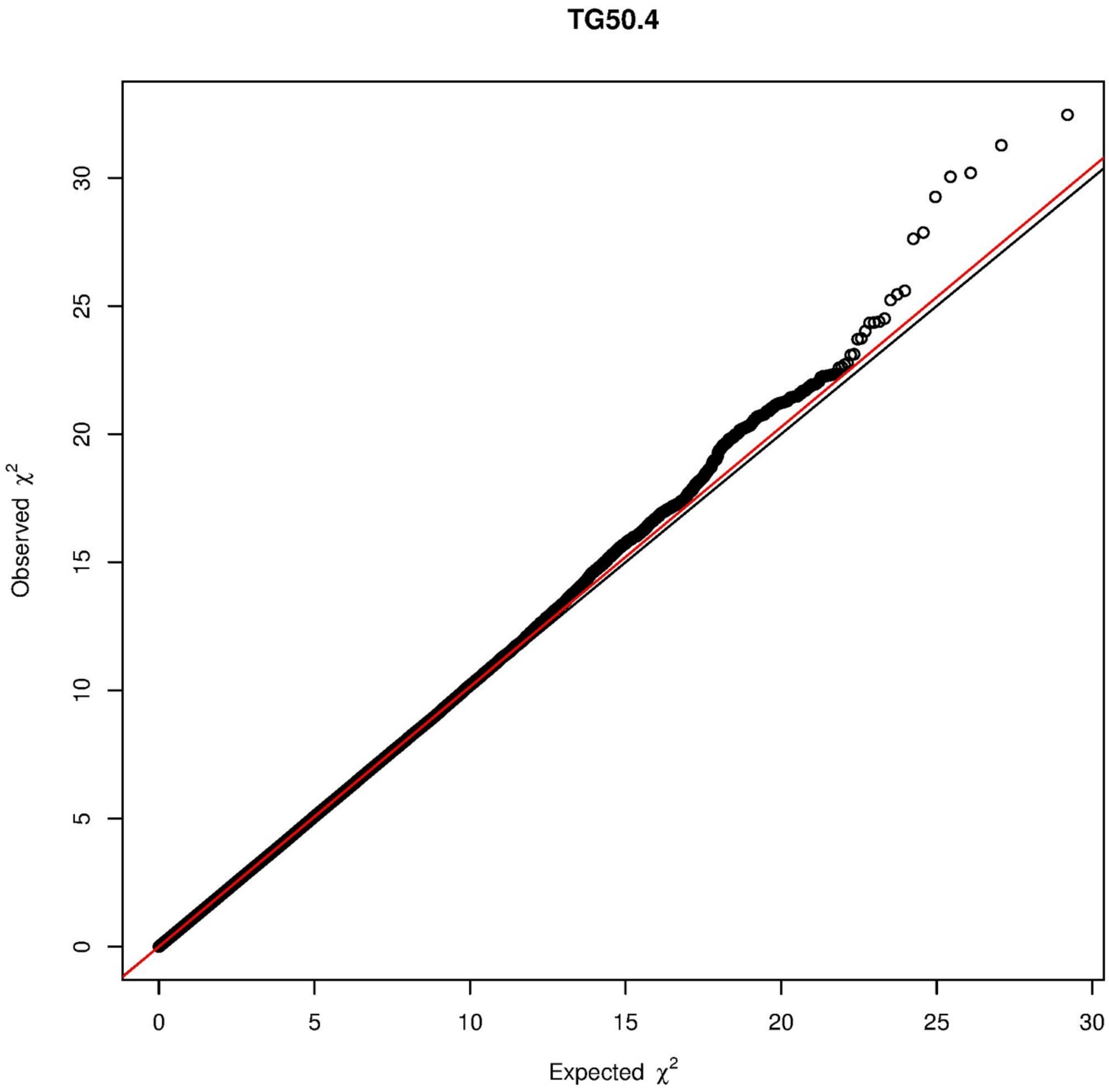

TG50.4
Observed $\chi^2$
Expected $\chi^2$
0
5
10
15
20
25
30

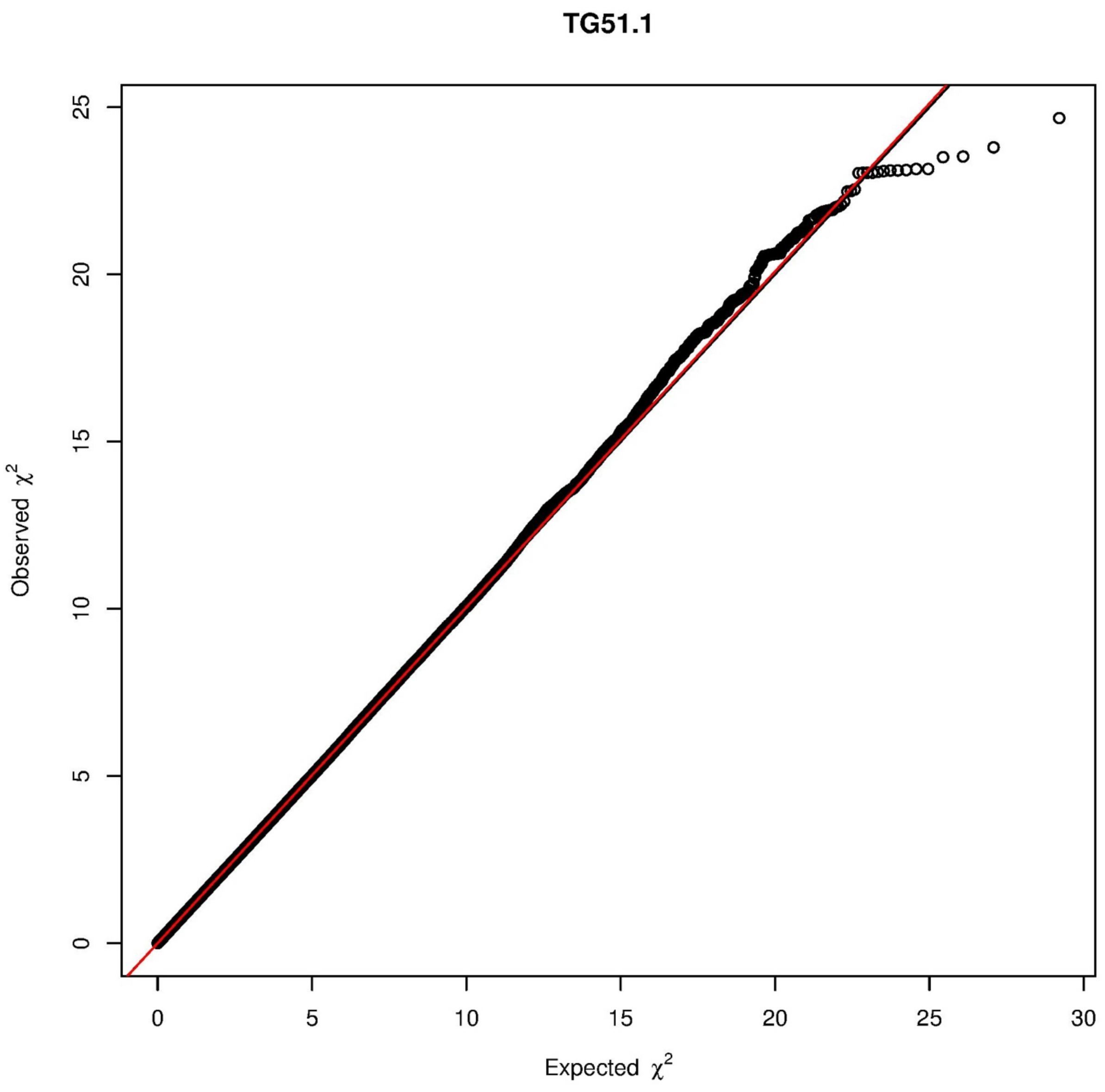
TG51.1
Observed $\chi^2$
Expected $\chi^2$
0
5
10
15
20
25
30

TG51.2

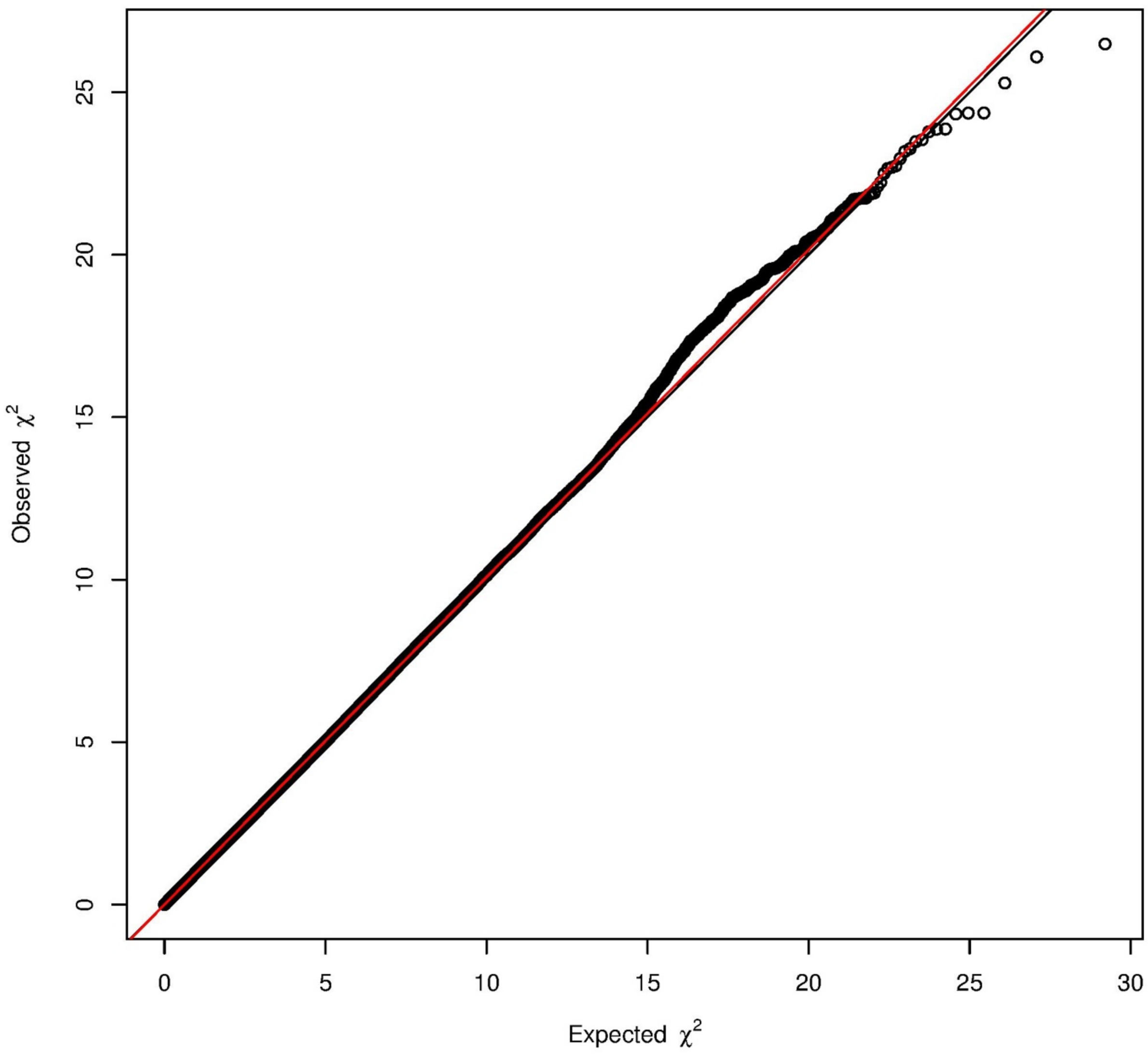

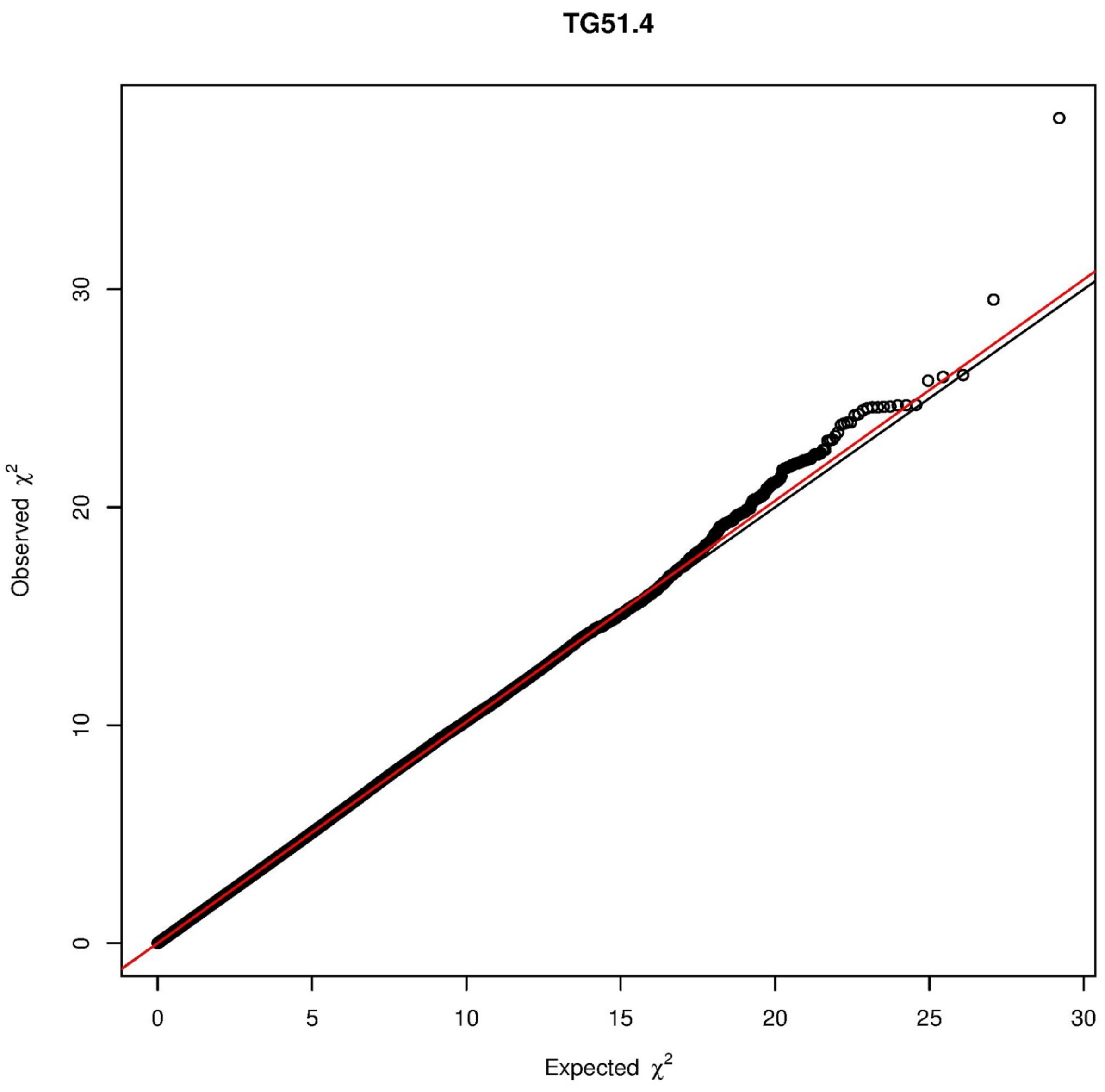

TG51.4
Observed $\chi^2$
Expected $\chi^2$
0
10
20
30
0
5
10
15
20
25
30

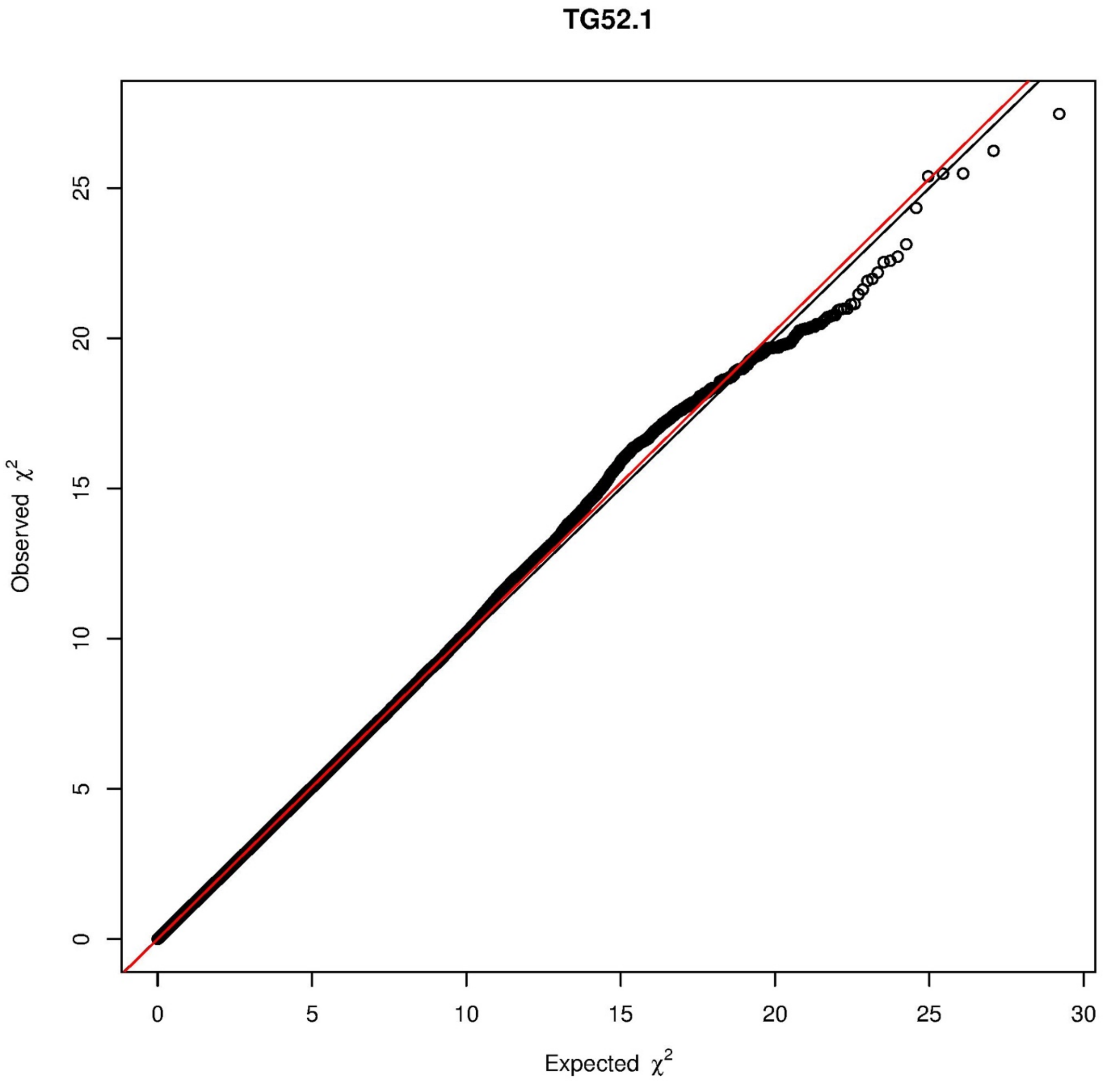
TG52.1
Observed χ2
Expected χ2
0
5
10
15
20
25
30

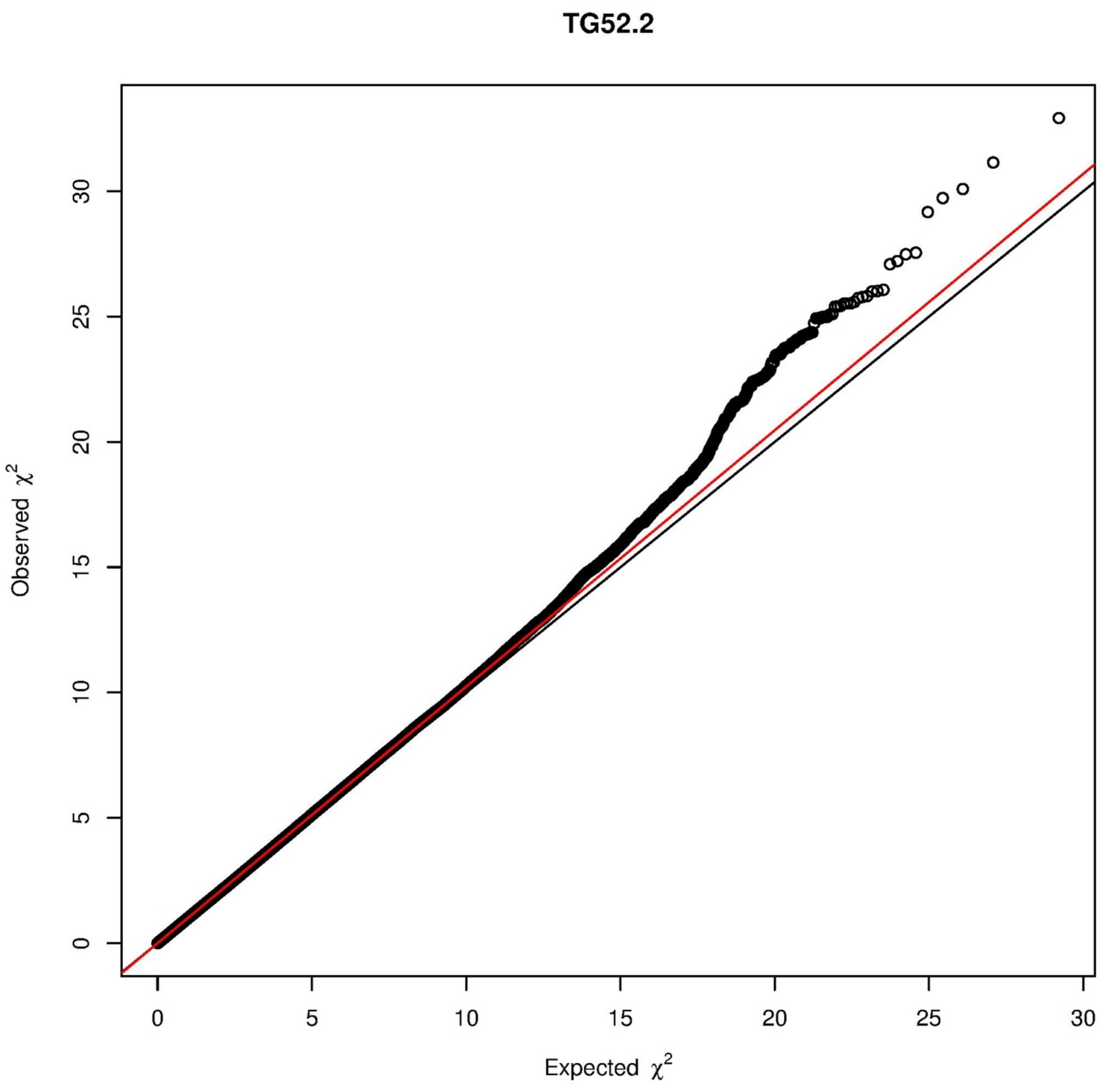
TG52.2
Observed $\chi^2$
Expected $\chi^2$
0
5
10
15
20
25
30
0
5
10
15
20
25
30

## TG52.3

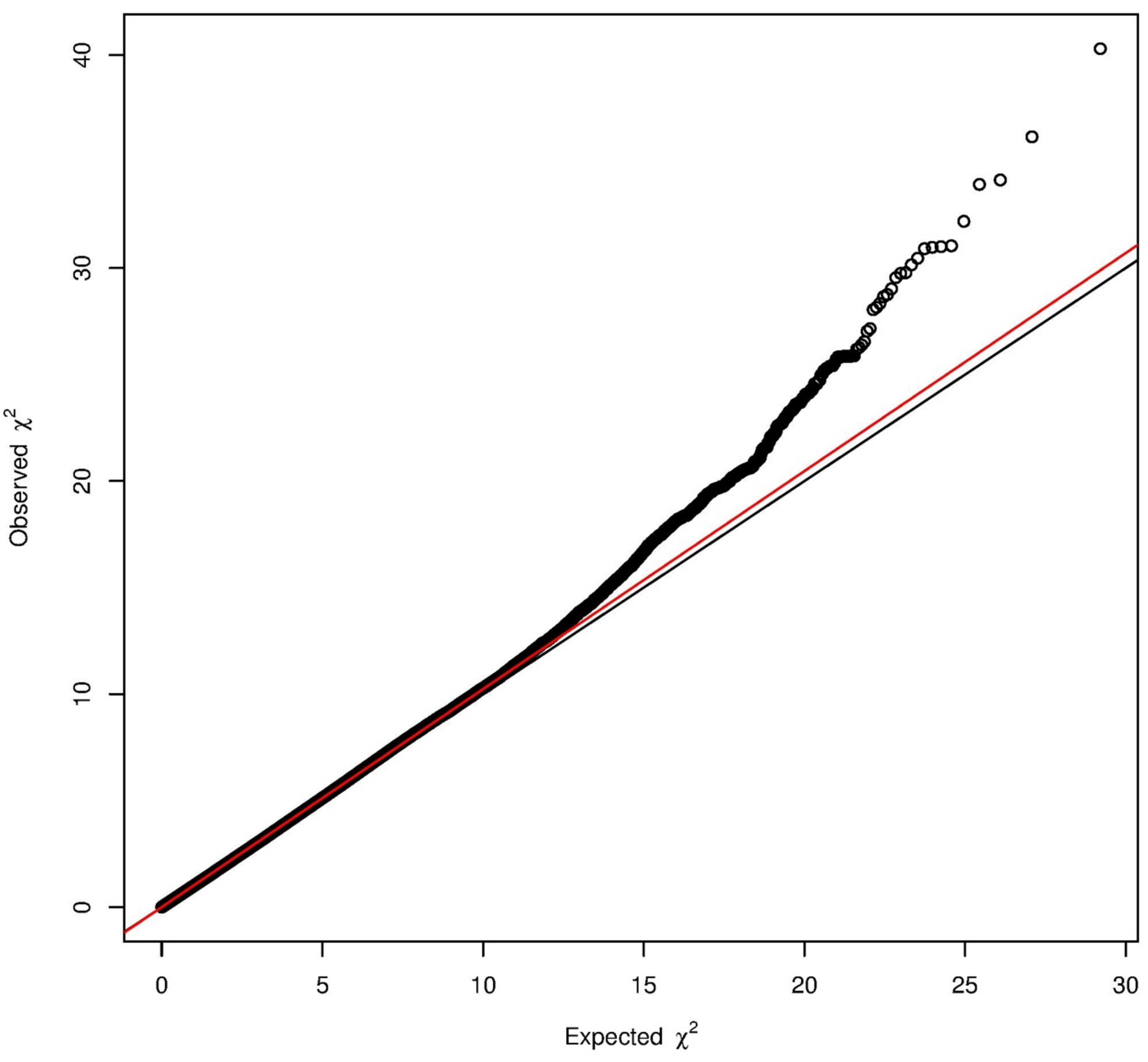

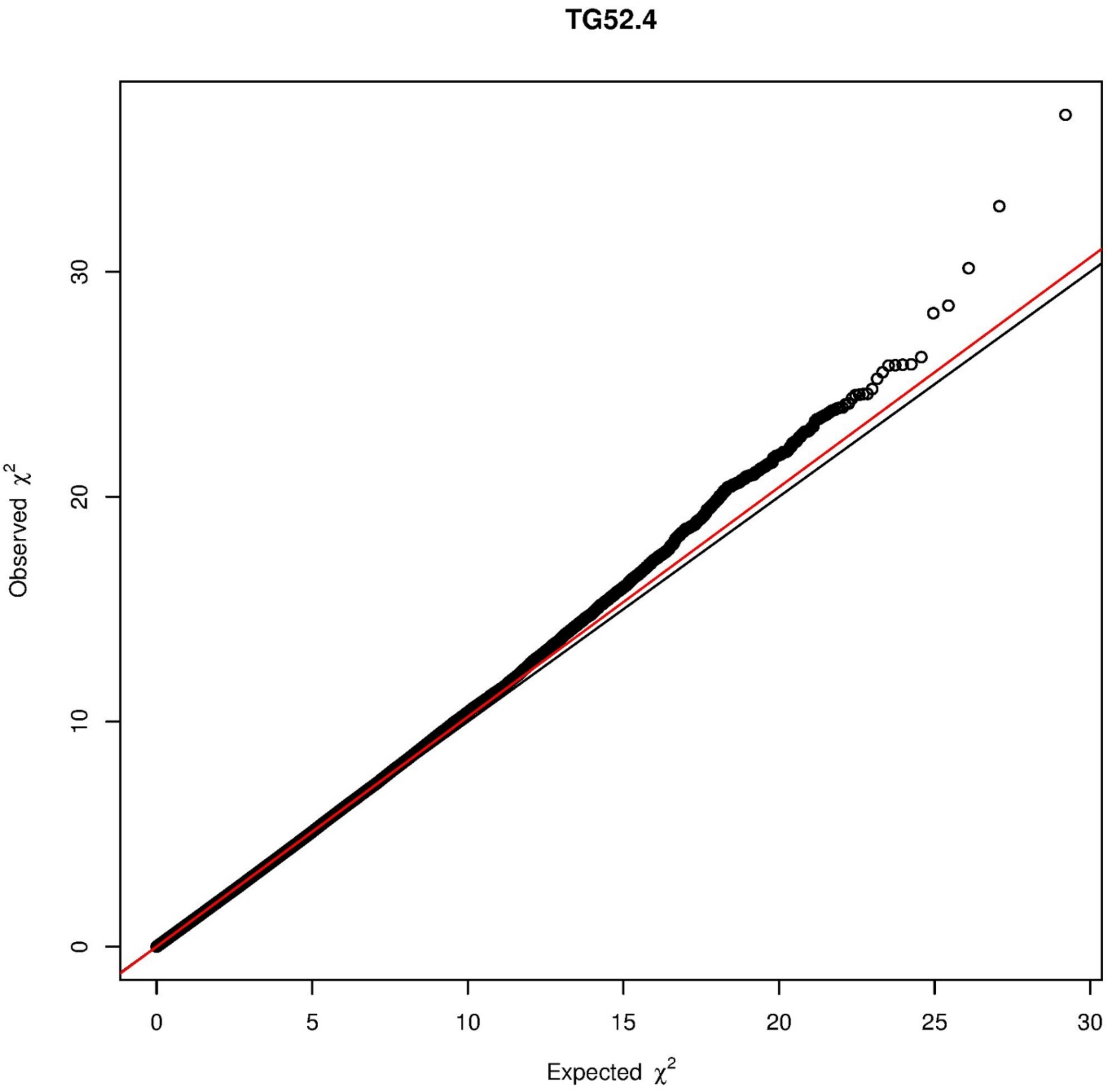

TG52.4
Observed $\chi^2$
Expected $\chi^2$
0
10
20
30
0
5
10
15
20
25
30

# TG52.5

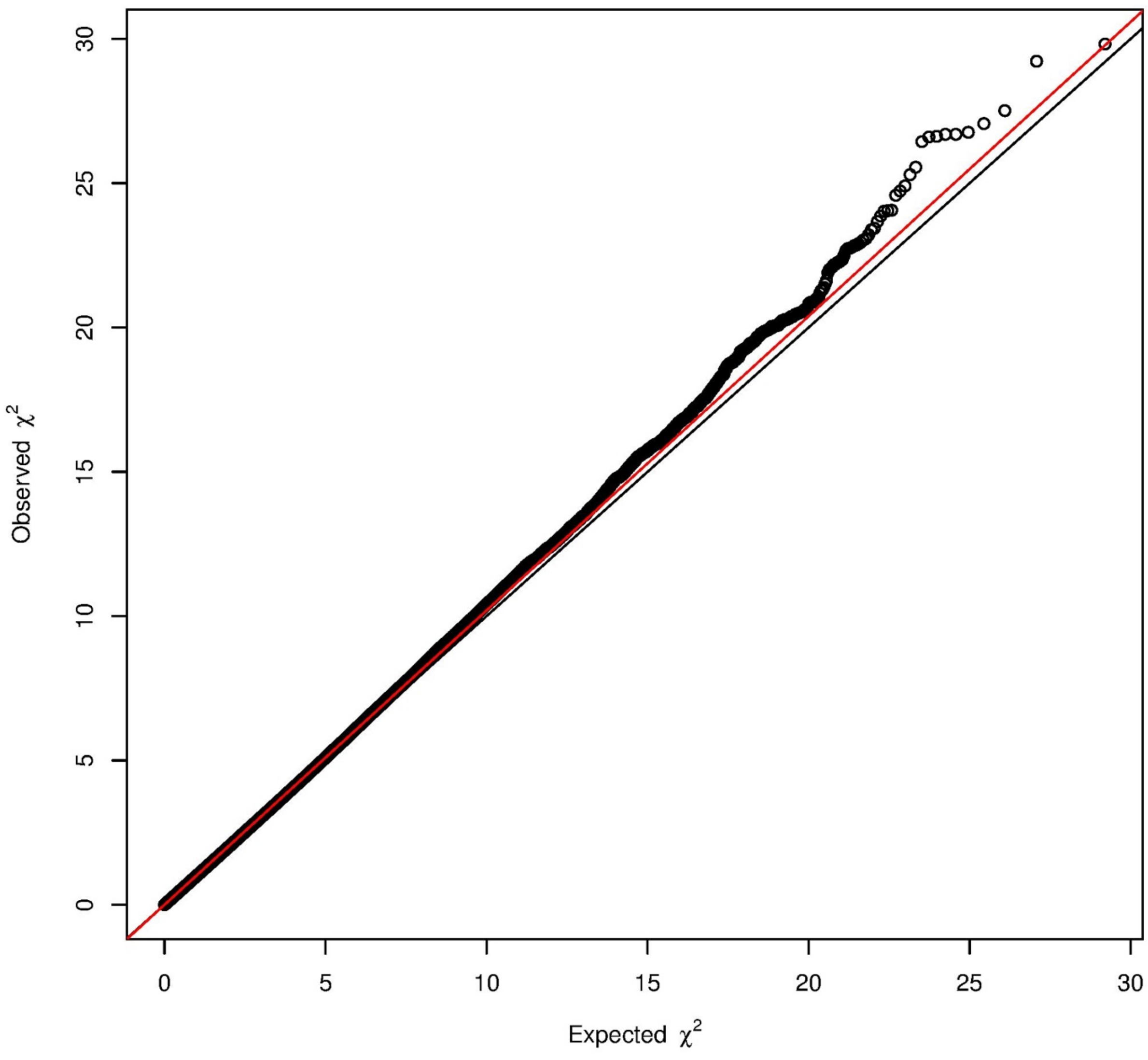

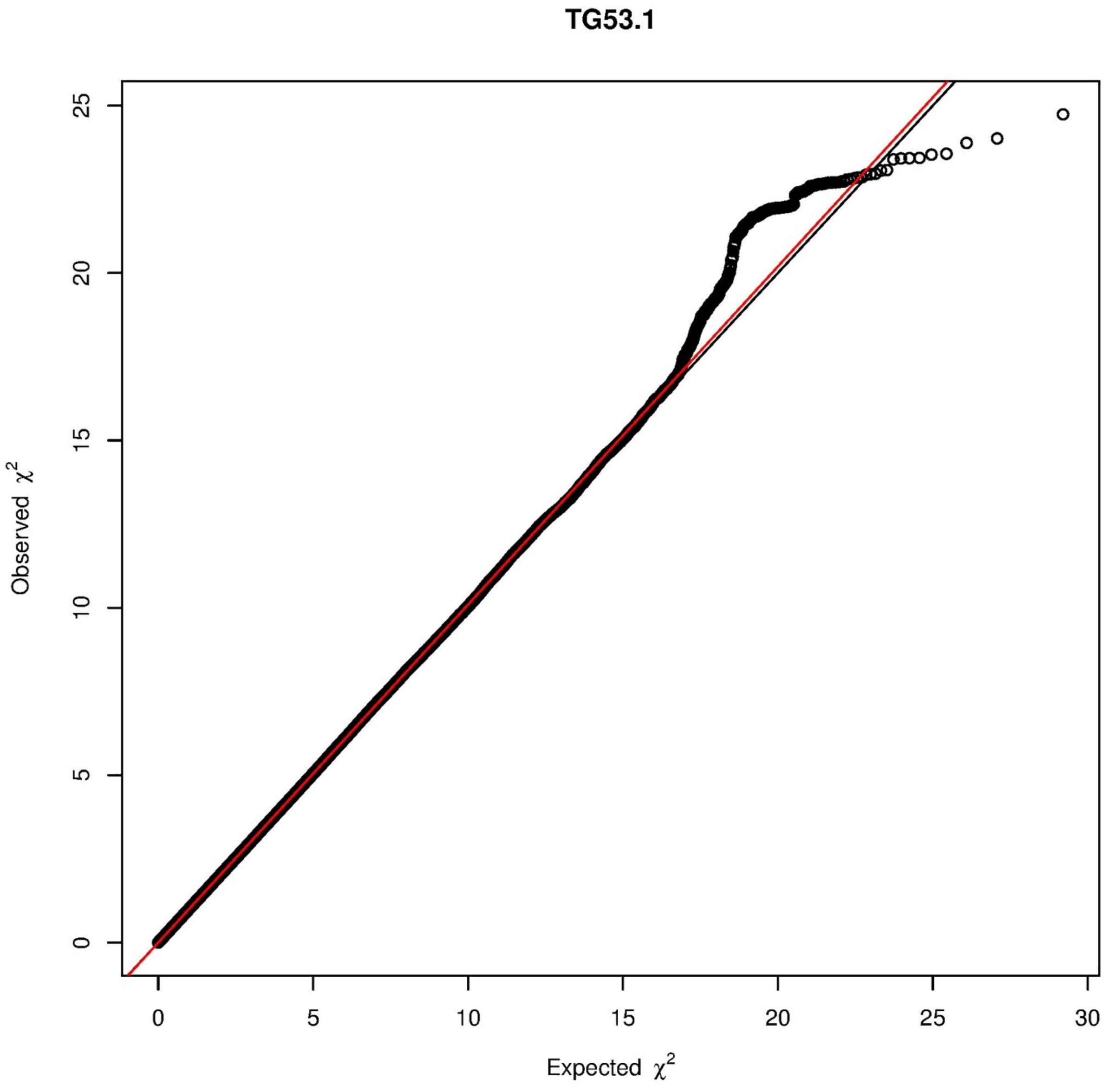
TG53.1
Observed χ²
Expected χ²
0
5
10
15
20
25
30

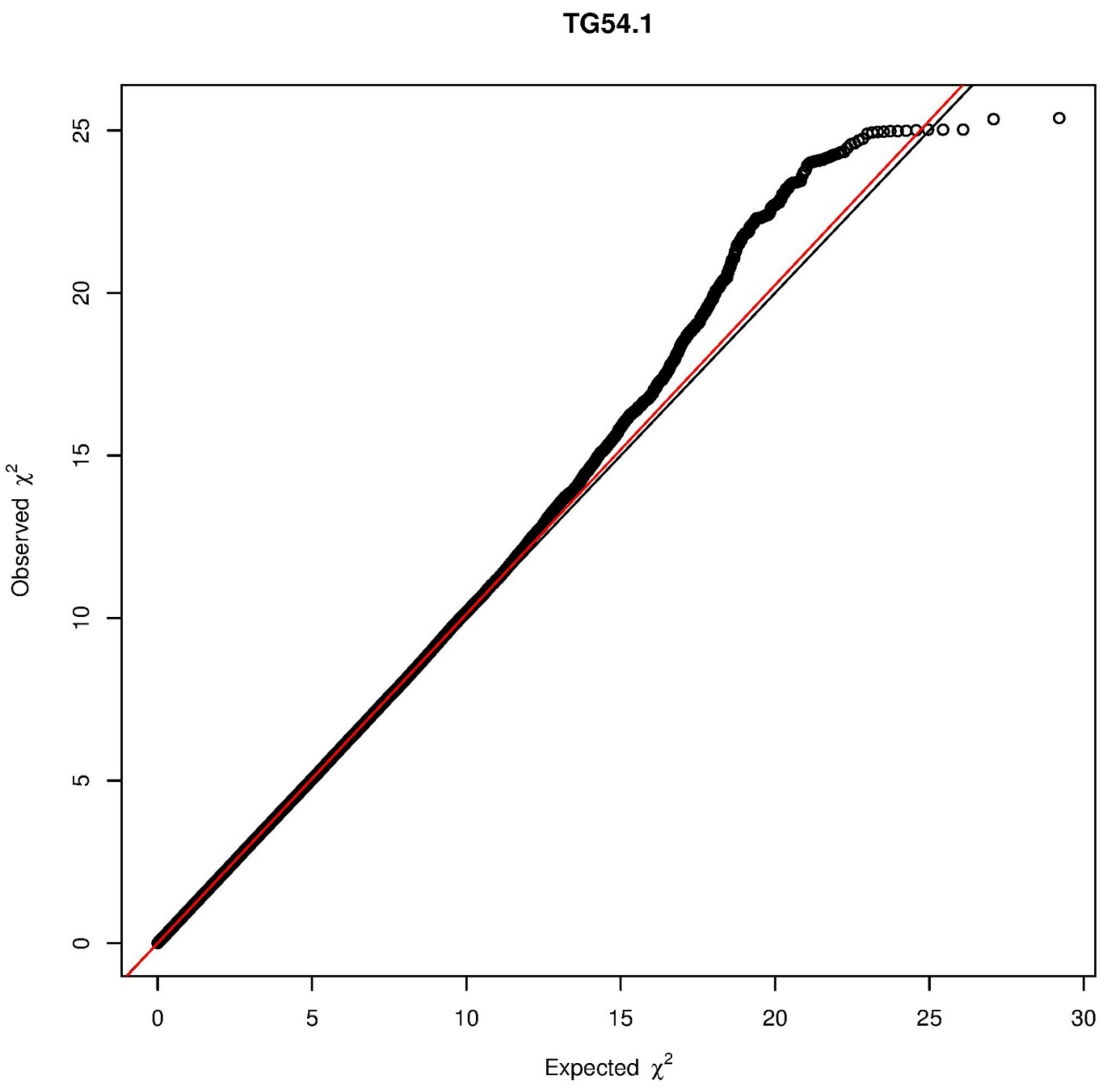

TG54.1
Observed χ²
Expected χ²
0
5
10
15
20
25
30

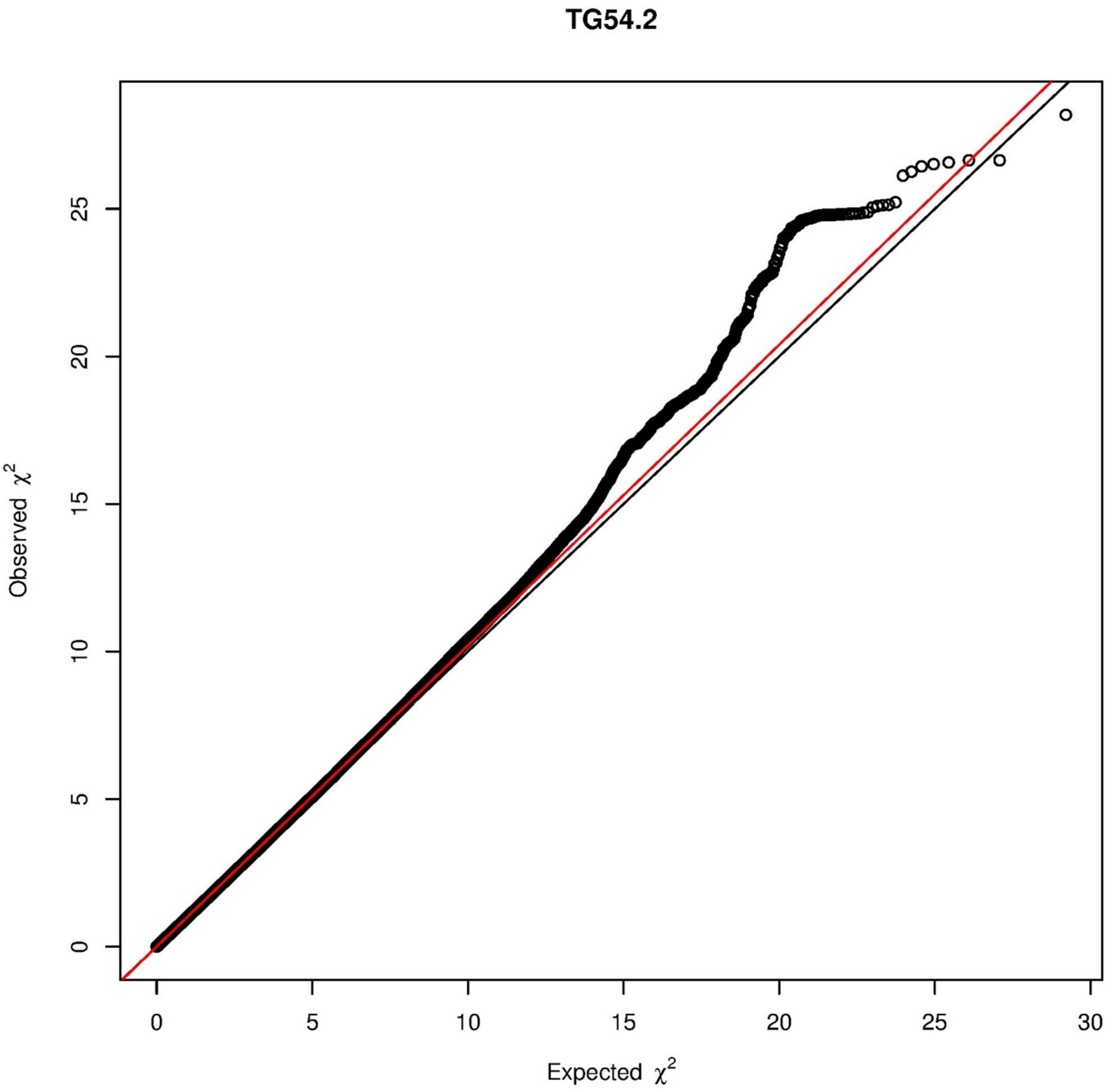

TG54.2
Observed $\chi^2$
Expected $\chi^2$
0
5
10
15
20
25
30

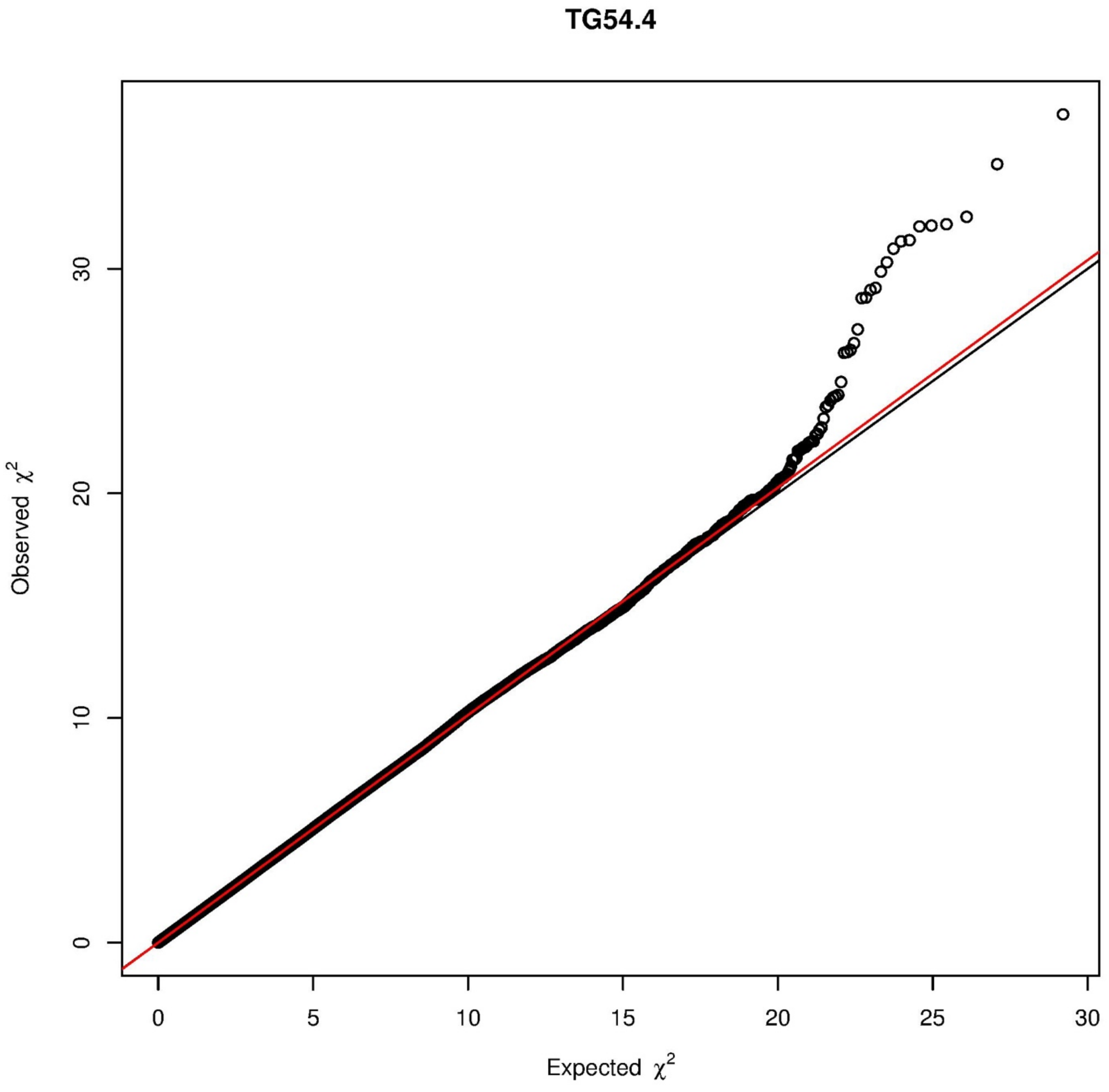

TG54.4
Observed χ²
Expected χ²
0
10
20
30
0
5
10
15
20
25
30

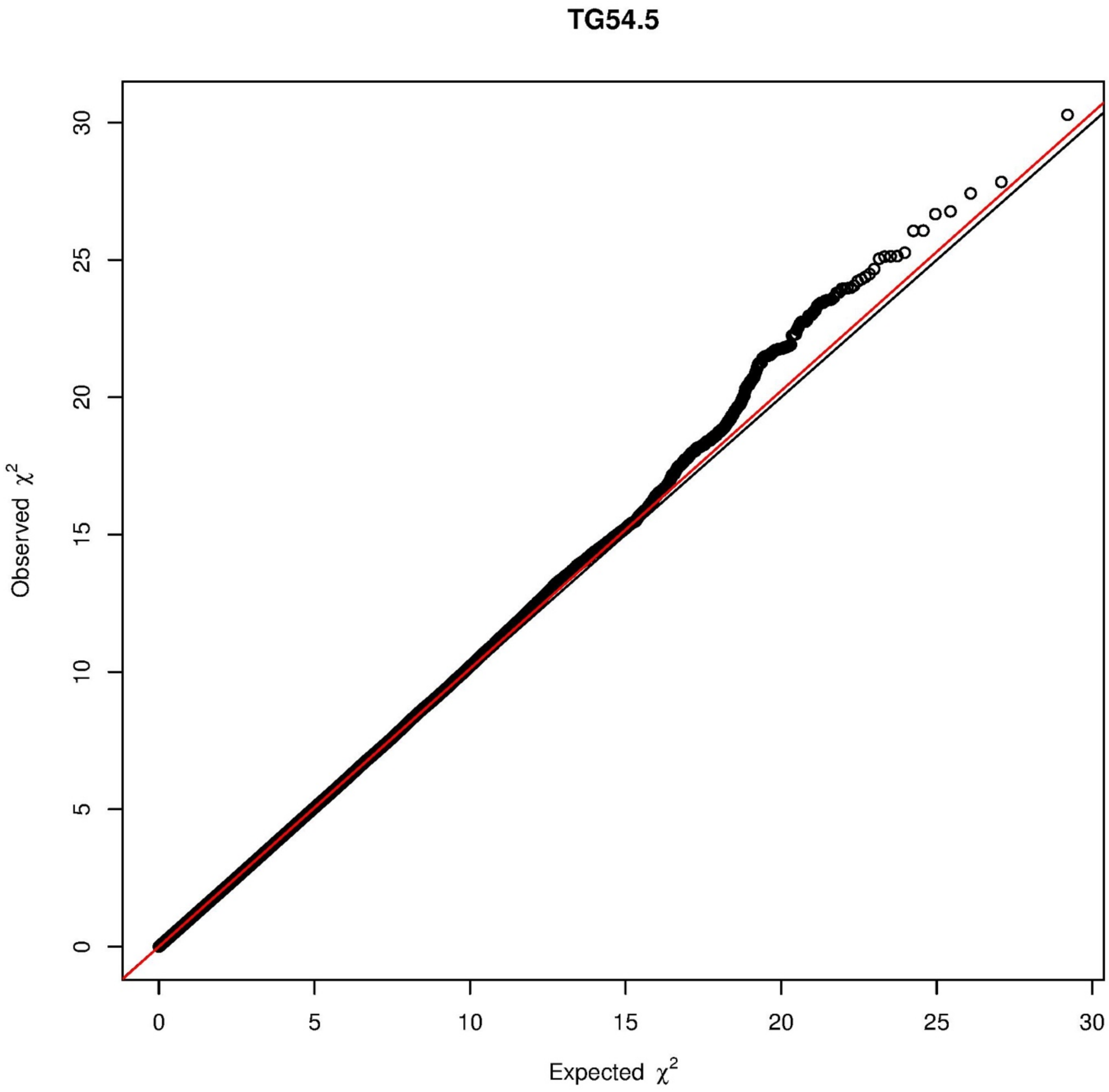
TG54.5
Observed $\chi^2$
Expected $\chi^2$
0
5
10
15
20
25
30

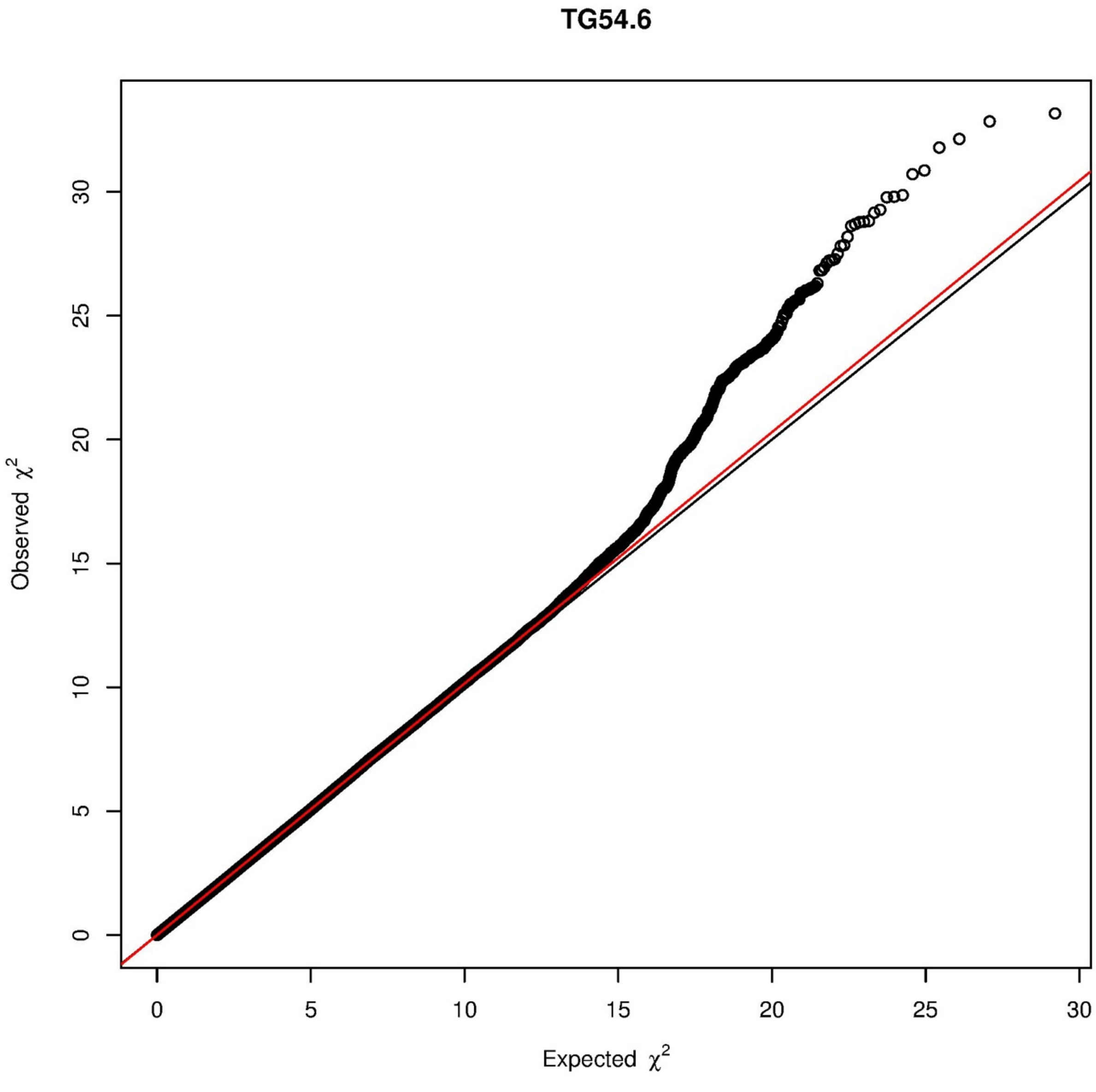

TG54.6
Observed $\chi^2$
Expected $\chi^2$
0
5
10
15
20
25
30
0
5
10
15
20
25
30

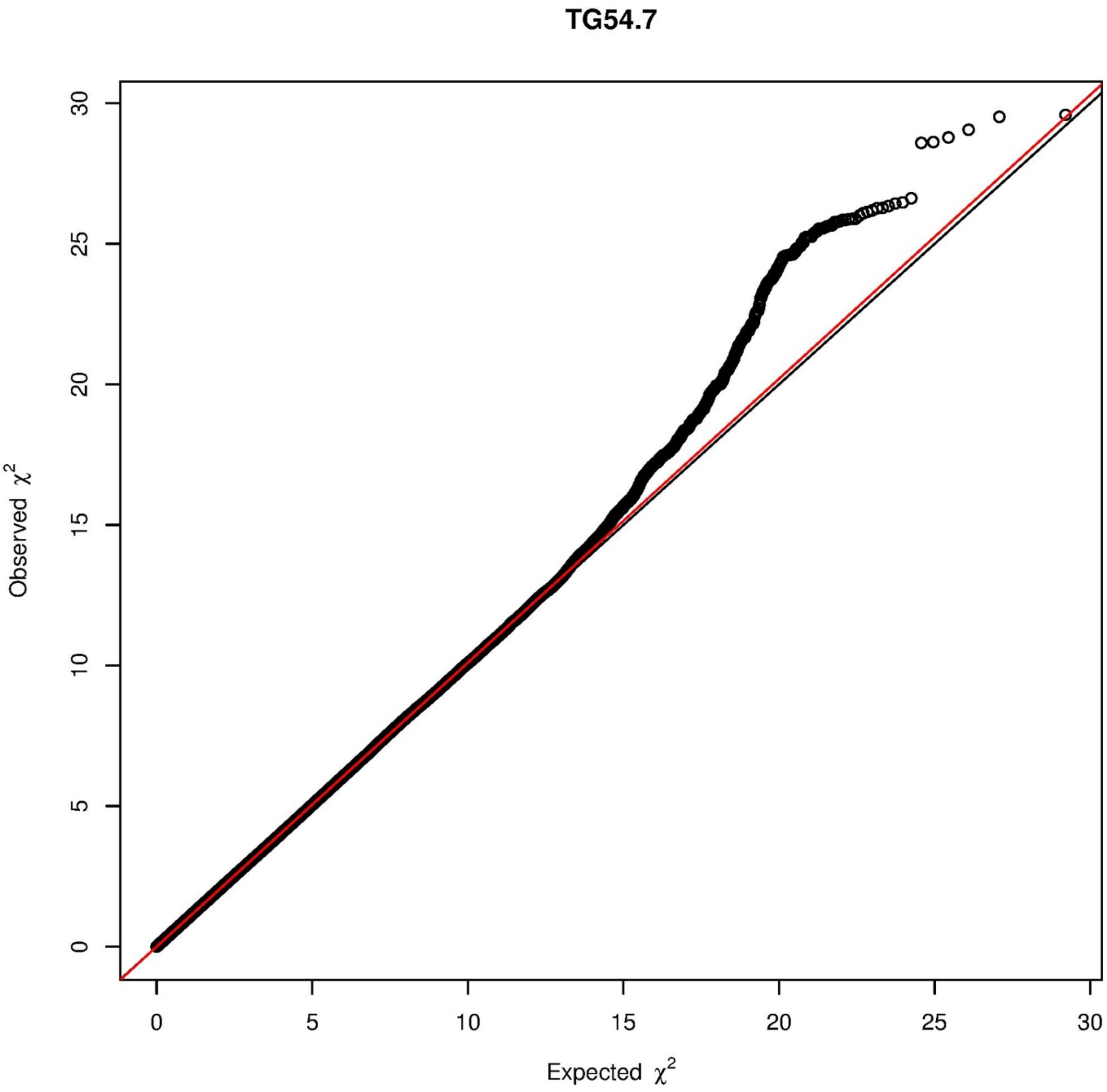
TG54.7
Observed $\chi^2$
Expected $\chi^2$
0
5
10
15
20
25
30
0
5
10
15
20
25
30

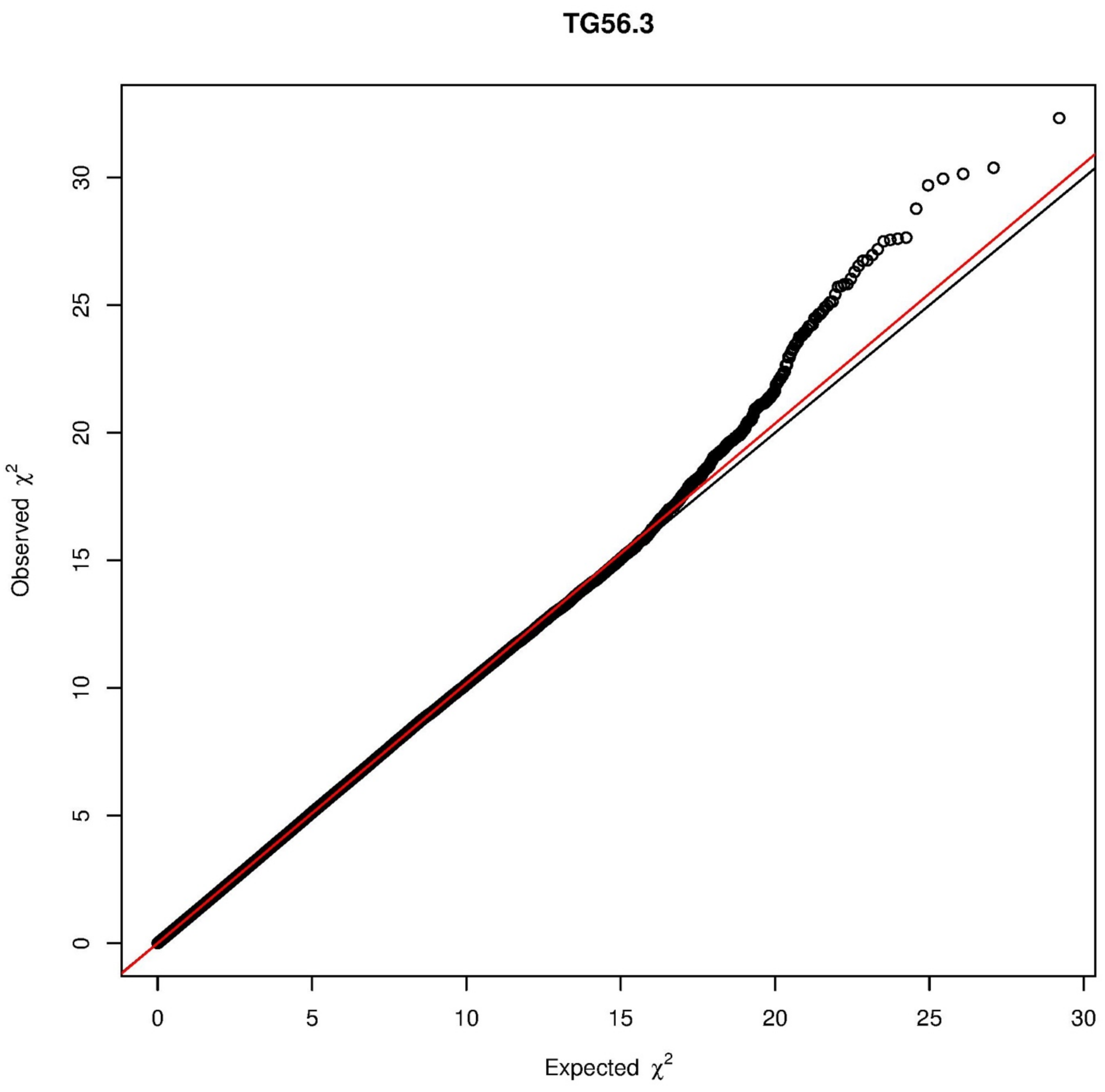
TG56.3
Observed $\chi^2$
Expected $\chi^2$
0
5
10
15
20
25
30

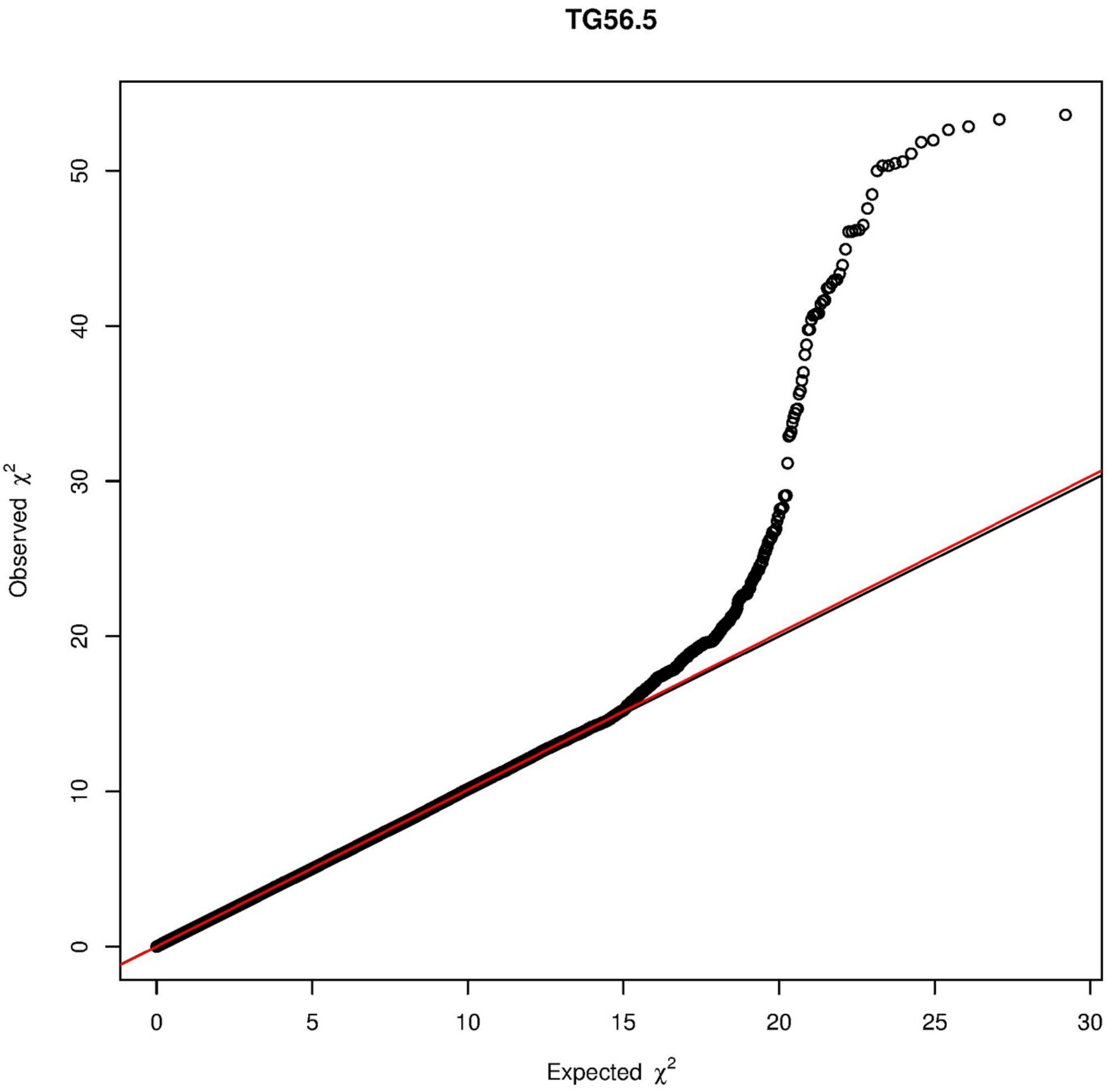
TG56.5
Observed $\chi^2$
Expected $\chi^2$
0
10
20
30
40
50
0
5
10
15
20
25
30

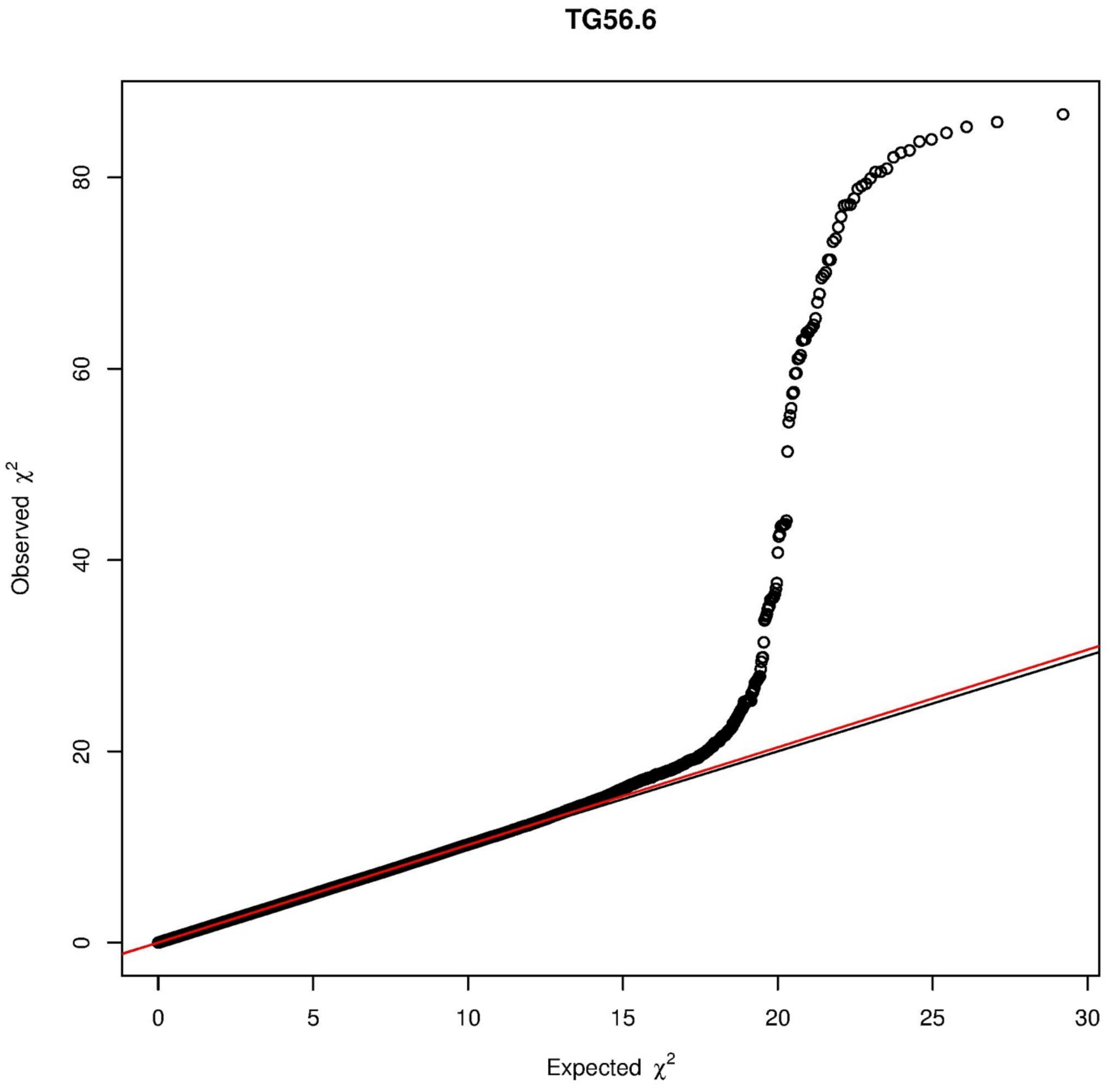
TG56.6
Observed $\chi^2$
Expected $\chi^2$
0
20
40
60
80
0
5
10
15
20
25
30

**TG56.7**

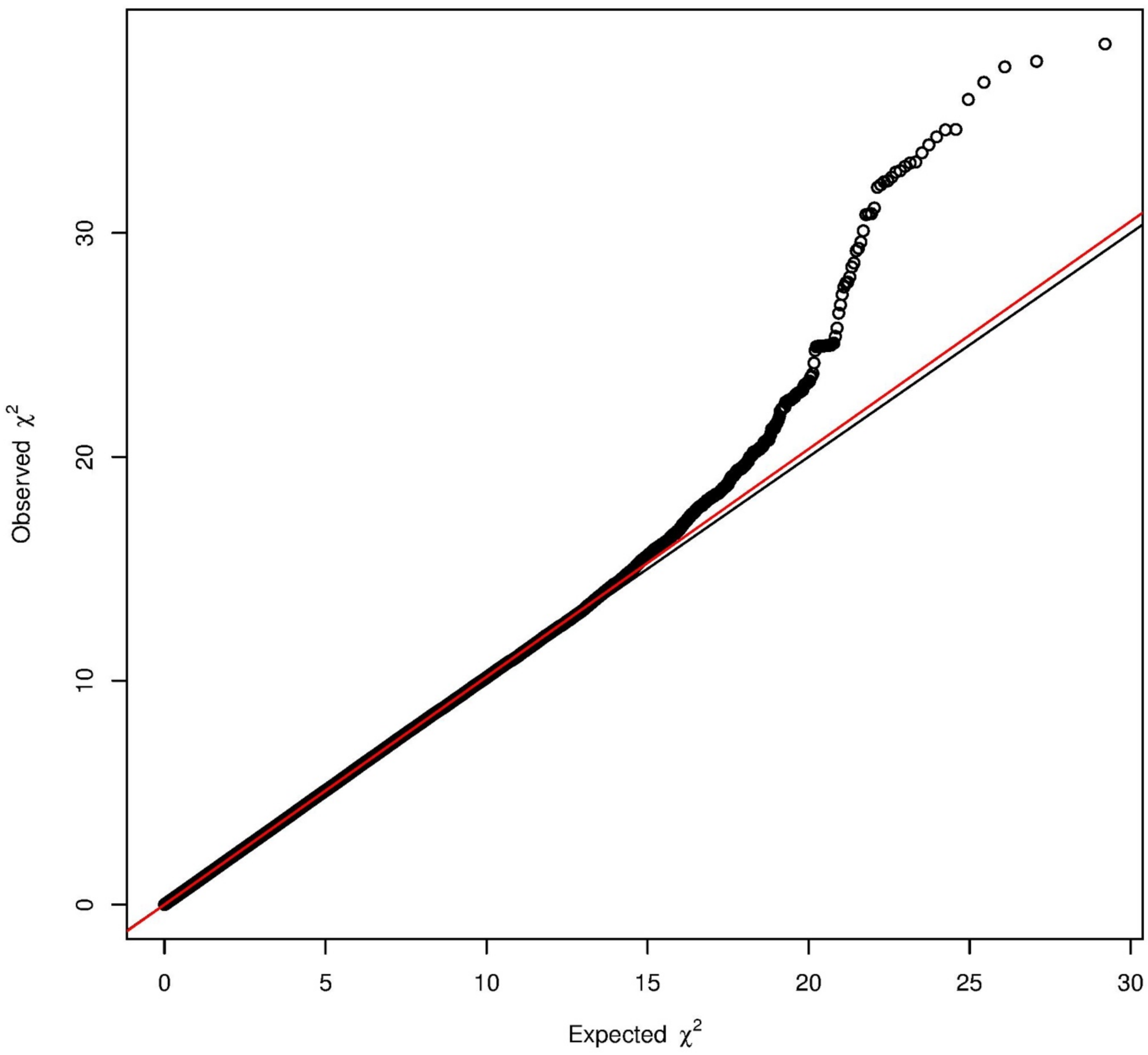

**Supplementary Figure 3** Manhattan plots of the metabolites studied

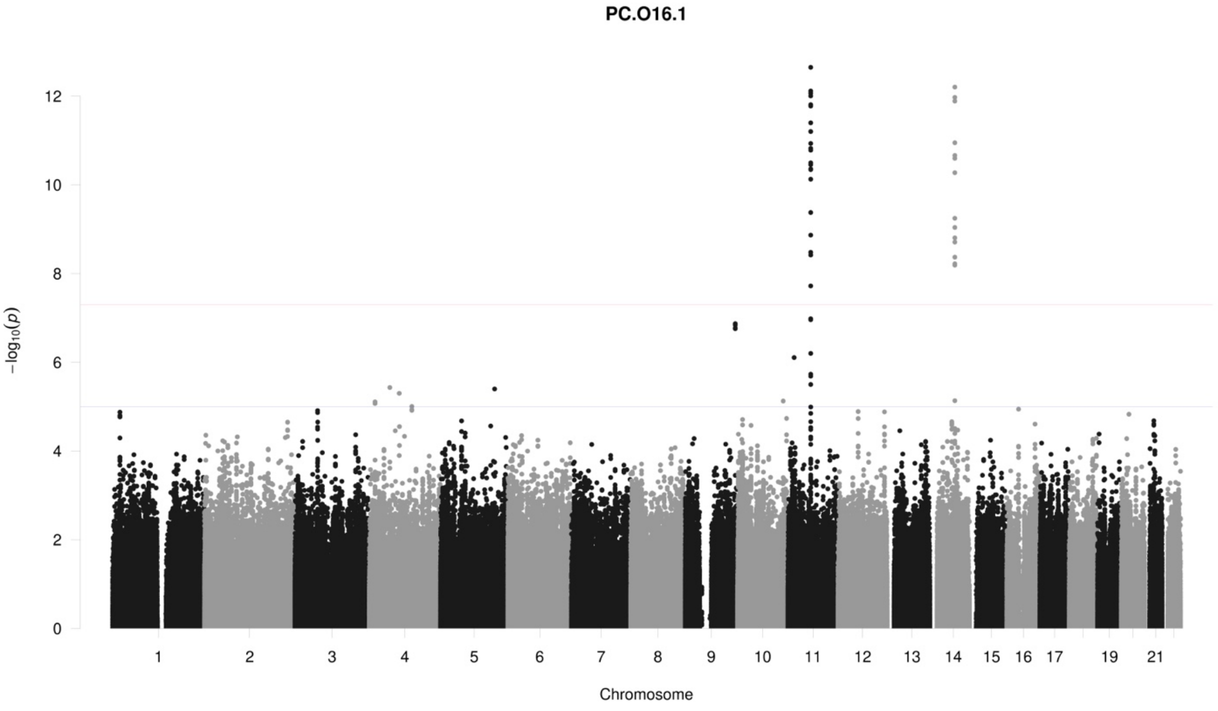

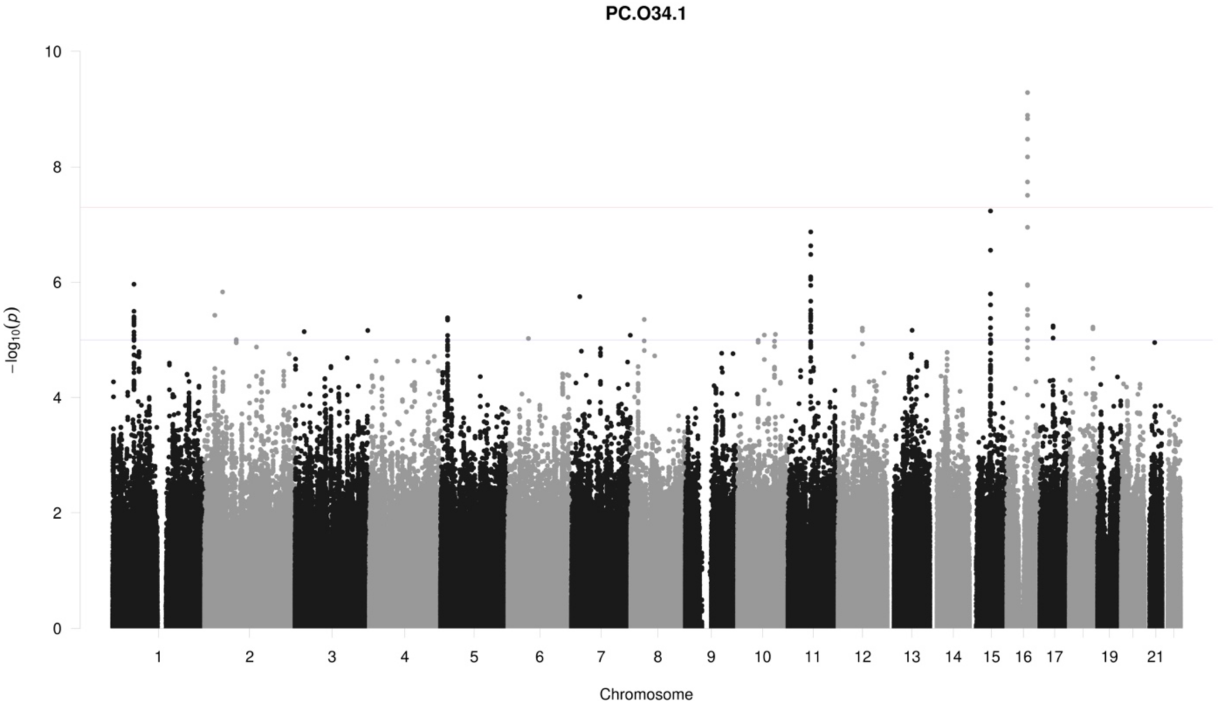

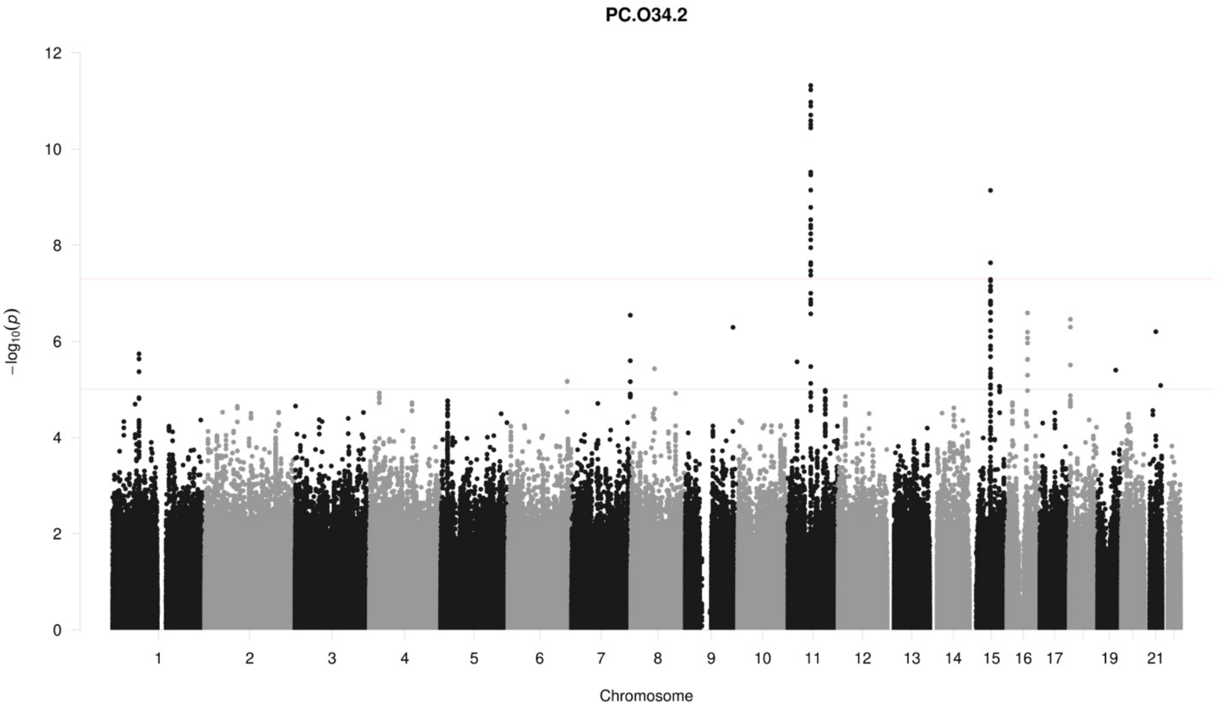
PC.O34.2
$-\log_{10}(p)$
Chromosome
0
2
4
6
8
10
12
19
21

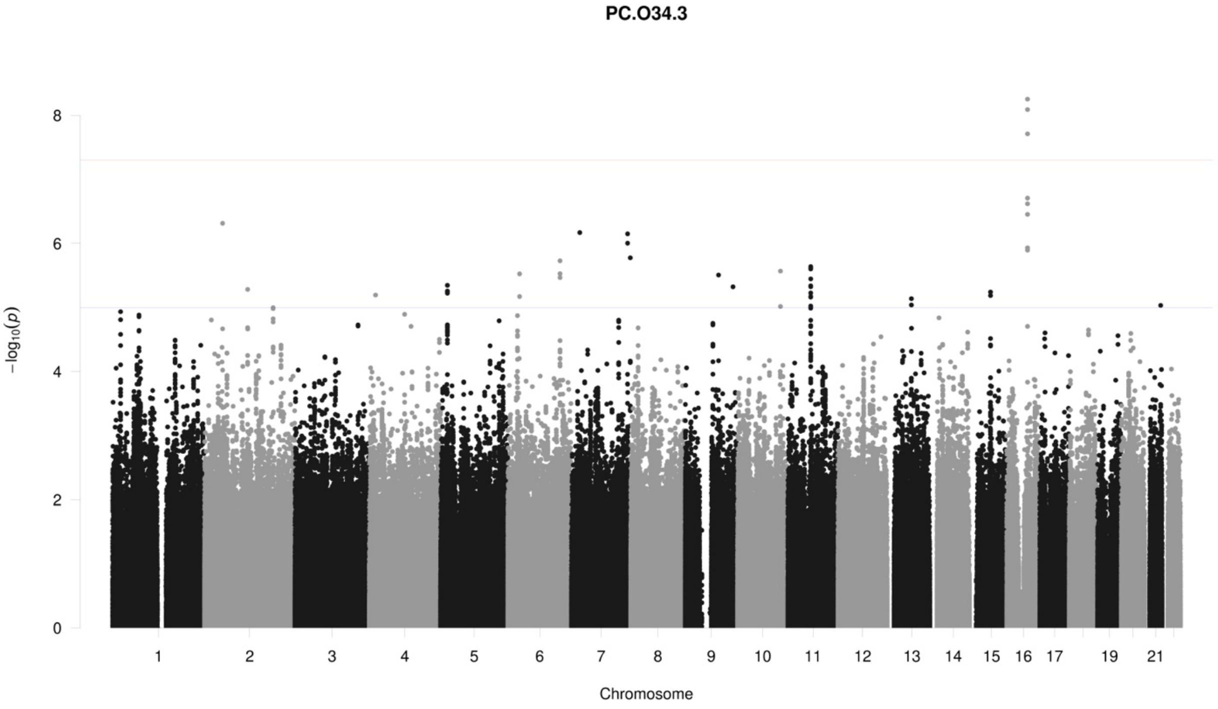
PC.O34.3
$-\log_{10}(p)$
Chromosome
0
2
4
6
8
19
21

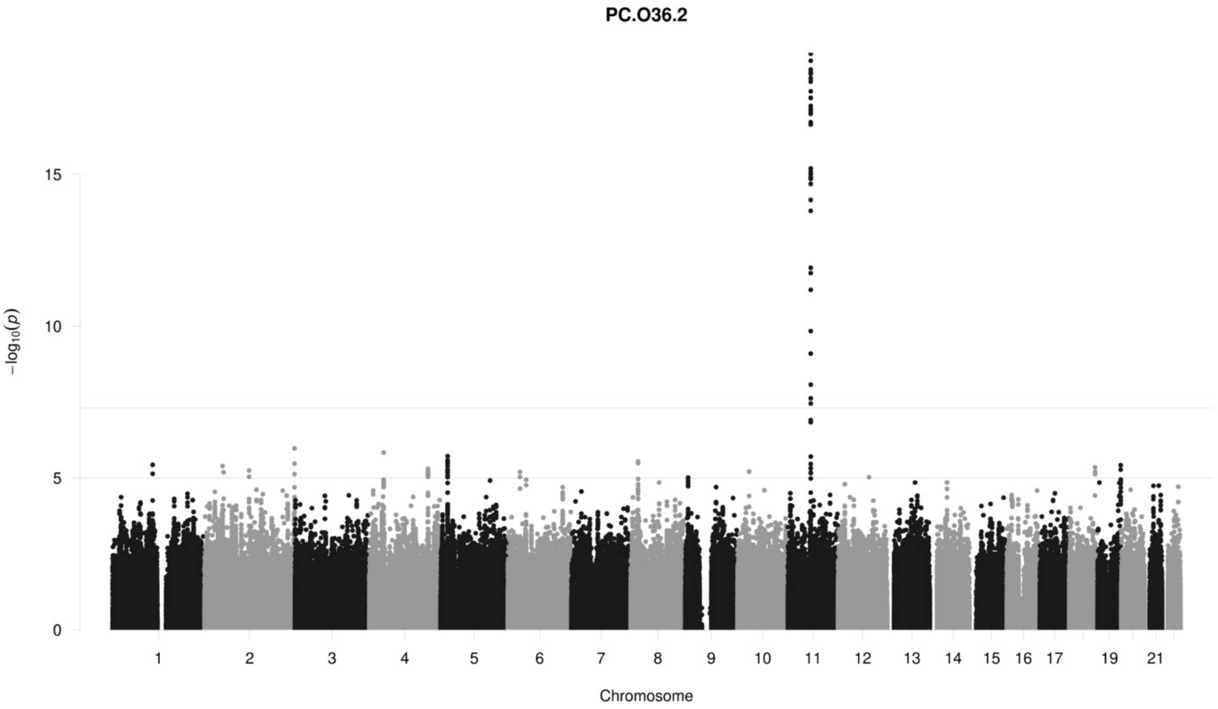
PC.O36.2
15
10
5
0
−log10(p)
1 2 3 4 5 6 7 8 9 10 11 12 13 14 15 16 17 19 21
Chromosome

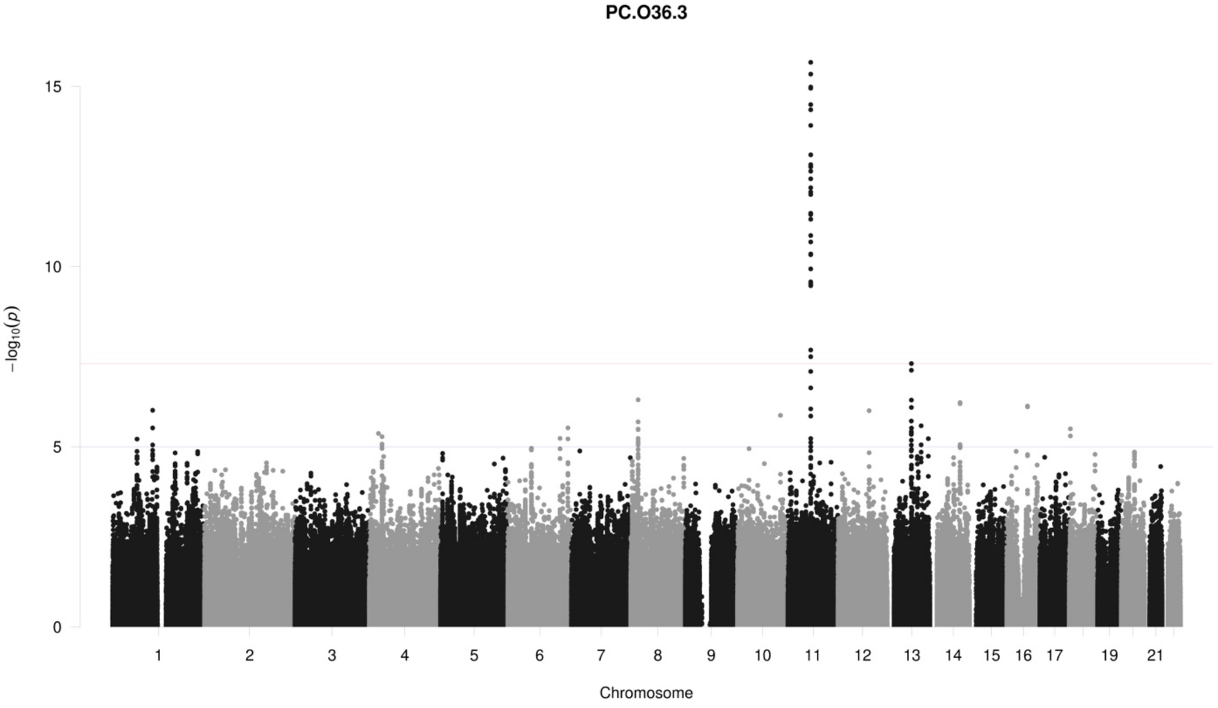
PC.O36.3
15
10
5
0
−log10(p)
1 2 3 4 5 6 7 8 9 10 11 12 13 14 15 16 17 19 21
Chromosome

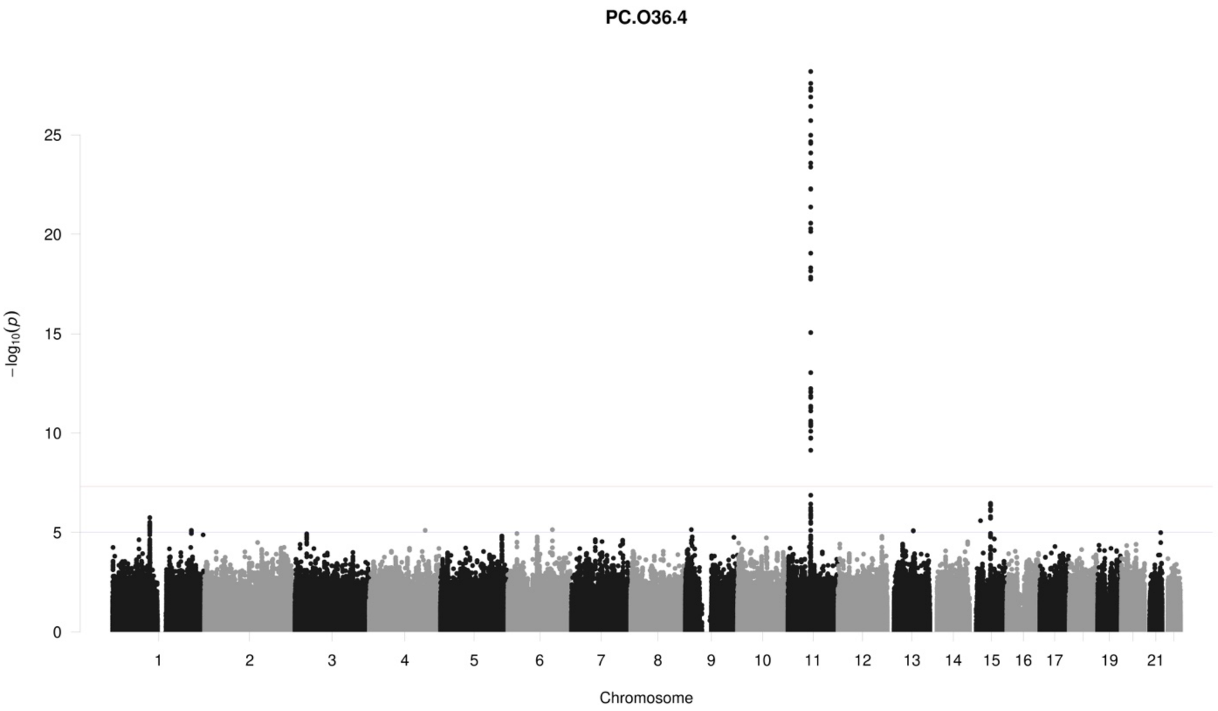
PC.O36.4
25
20
15
10
5
0
−log10(p)
19
21
Chromosome

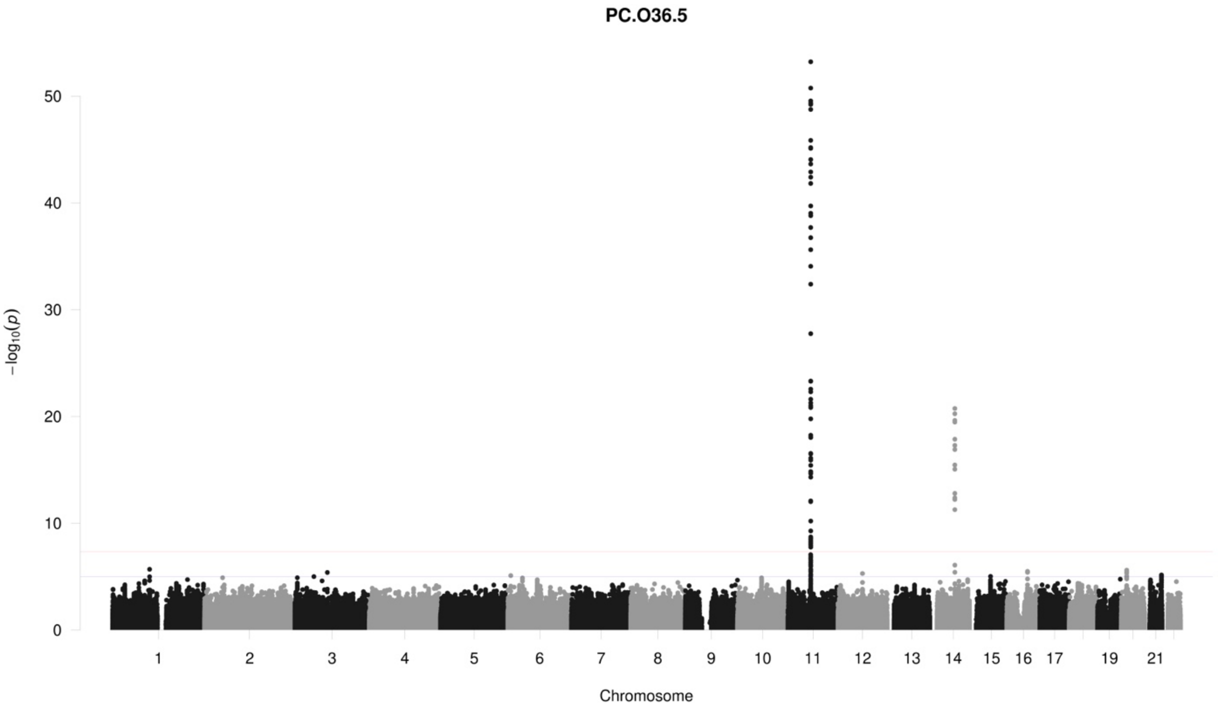
PC.O36.5
50
40
30
20
10
0
−log10(p)
19
21
Chromosome

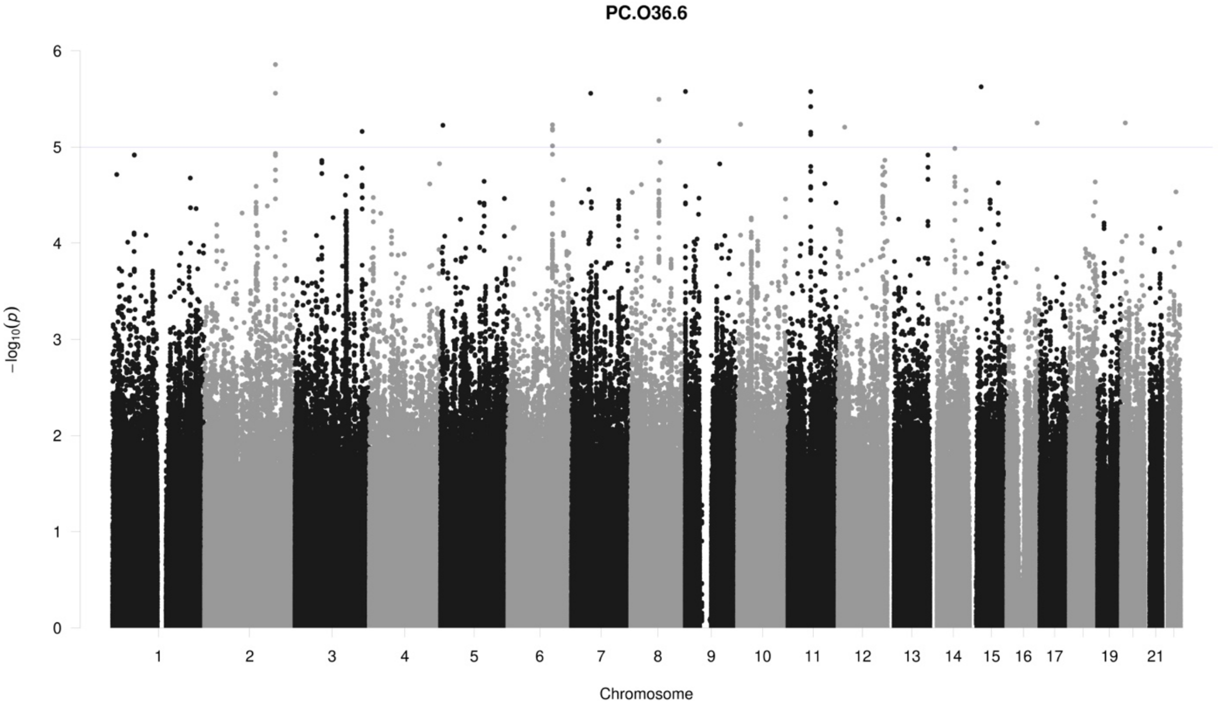
PC.O36.6
6
5
4
3
2
1
0
−log10(p)
19
21
Chromosome

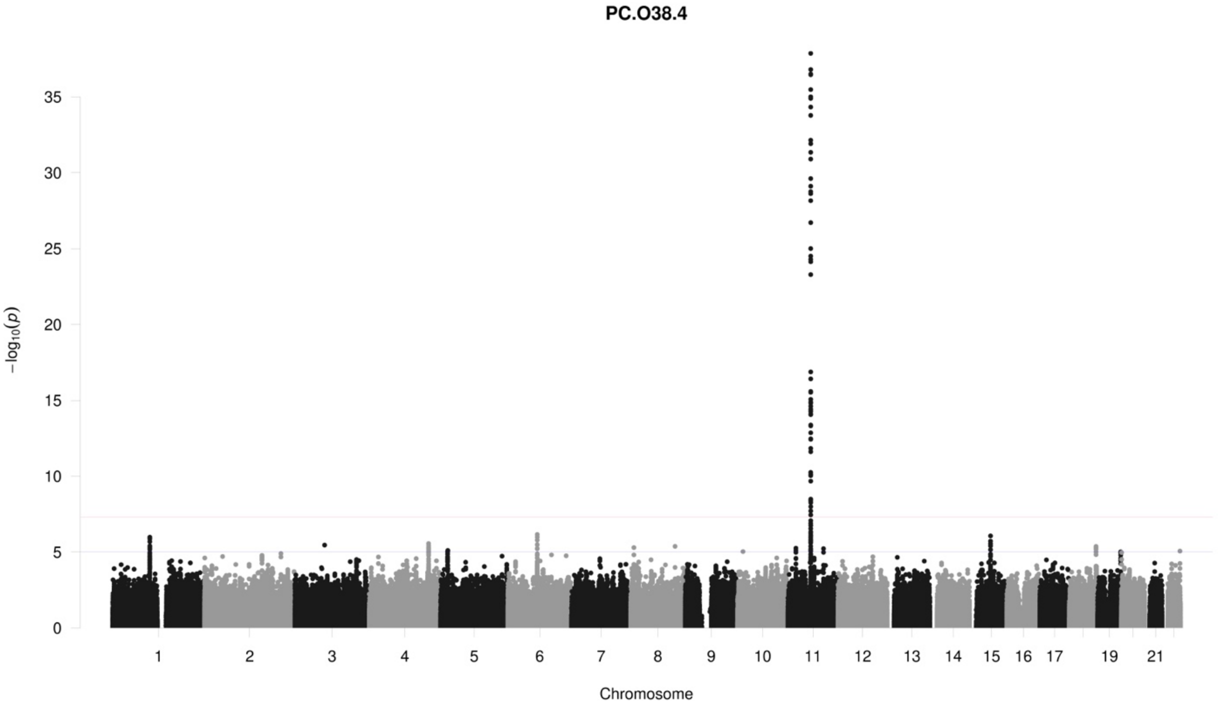
PC.O38.4
35
30
25
20
15
10
5
0
−log10(p)
19
21
Chromosome

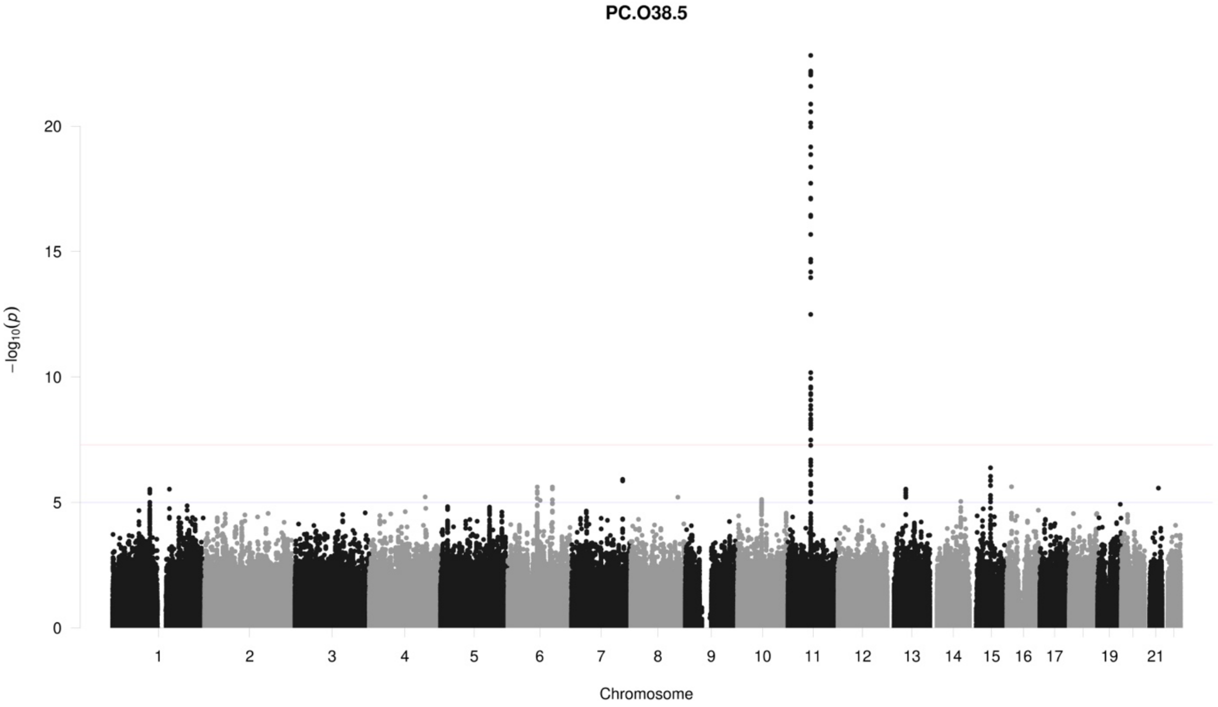
PC.O38.5
20
15
10
5
0
$-\log_{10}(p)$
19
21
Chromosome

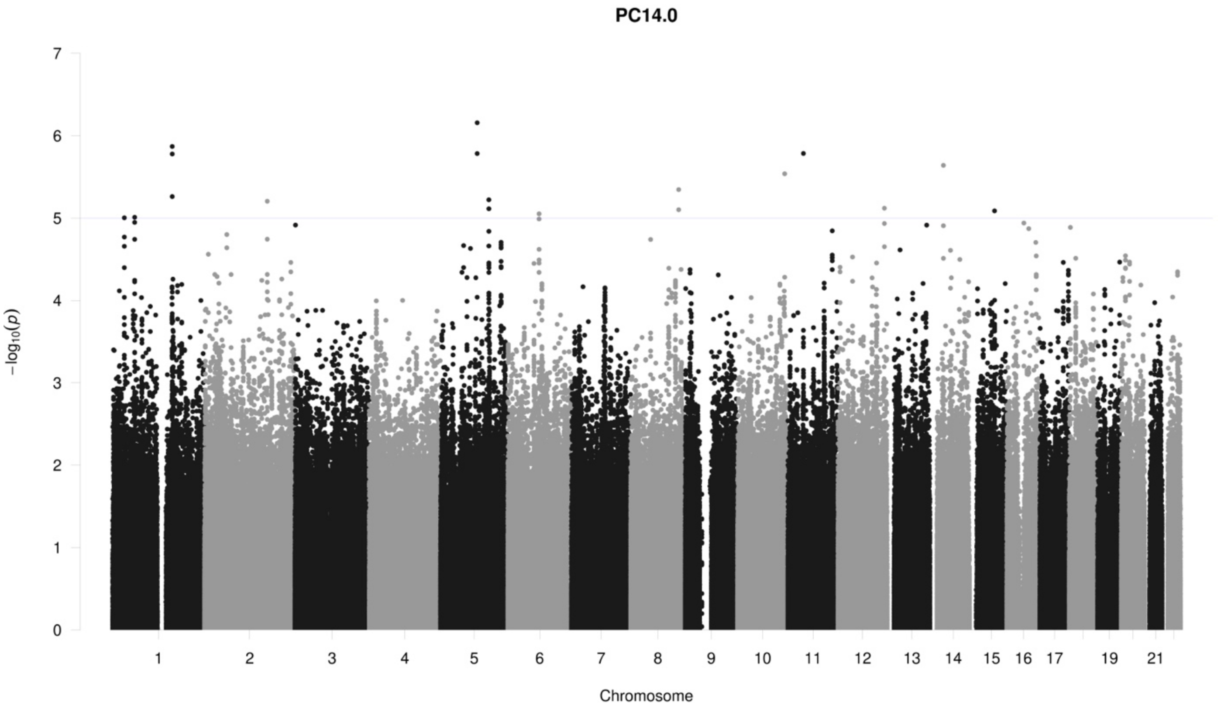
PC14.0
7
6
5
4
3
2
1
0
$-\log_{10}(p)$
19
21
Chromosome

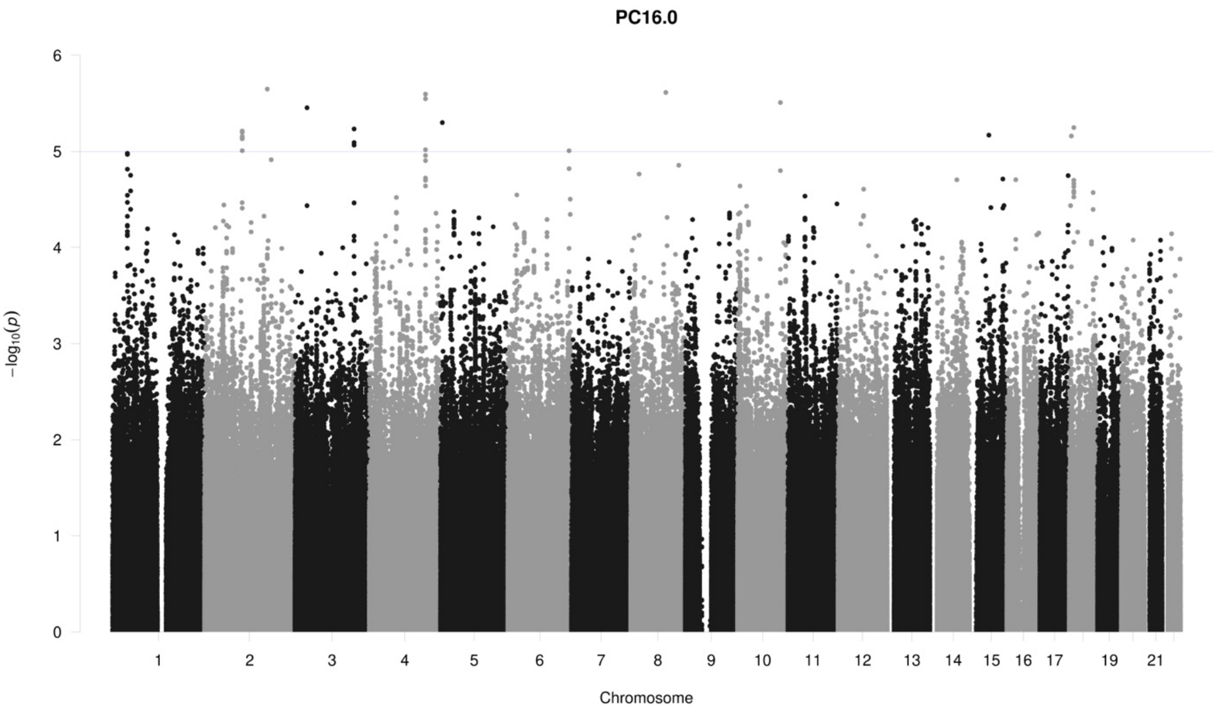

PC16.0
−log10(p)
0
1
2
3
4
5
6
19
21
Chromosome

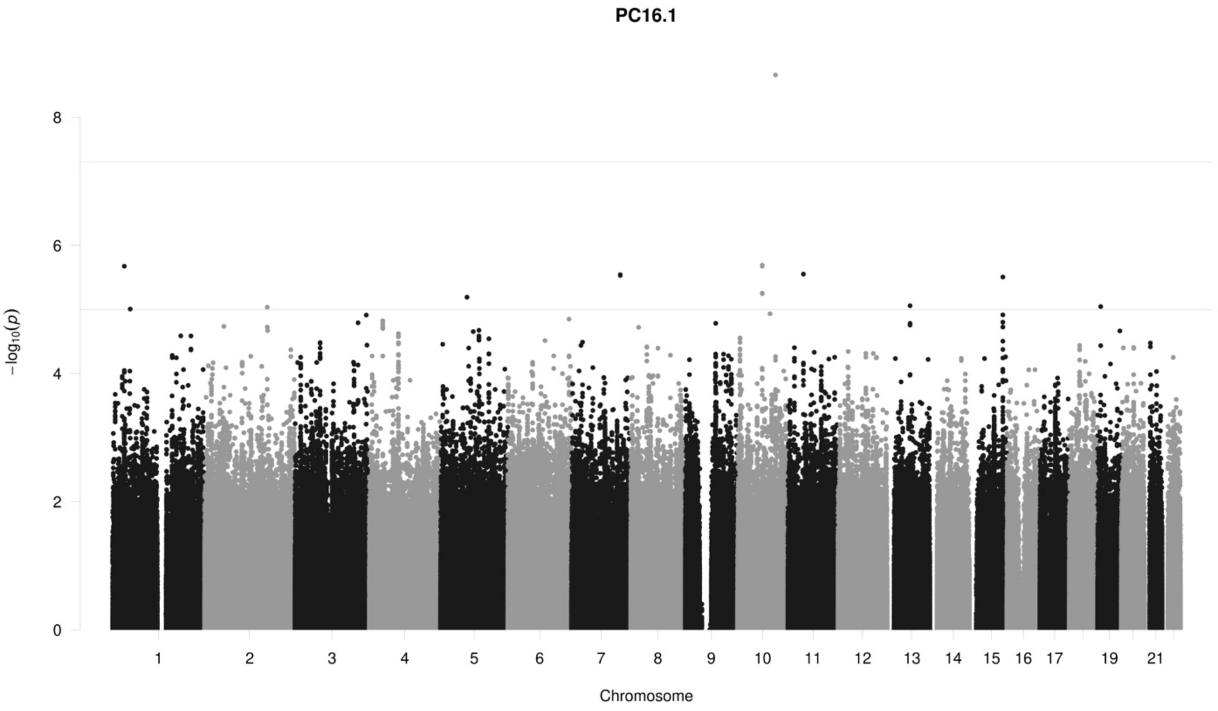

PC16.1
−log10(p)
0
2
4
6
8
19
21
Chromosome

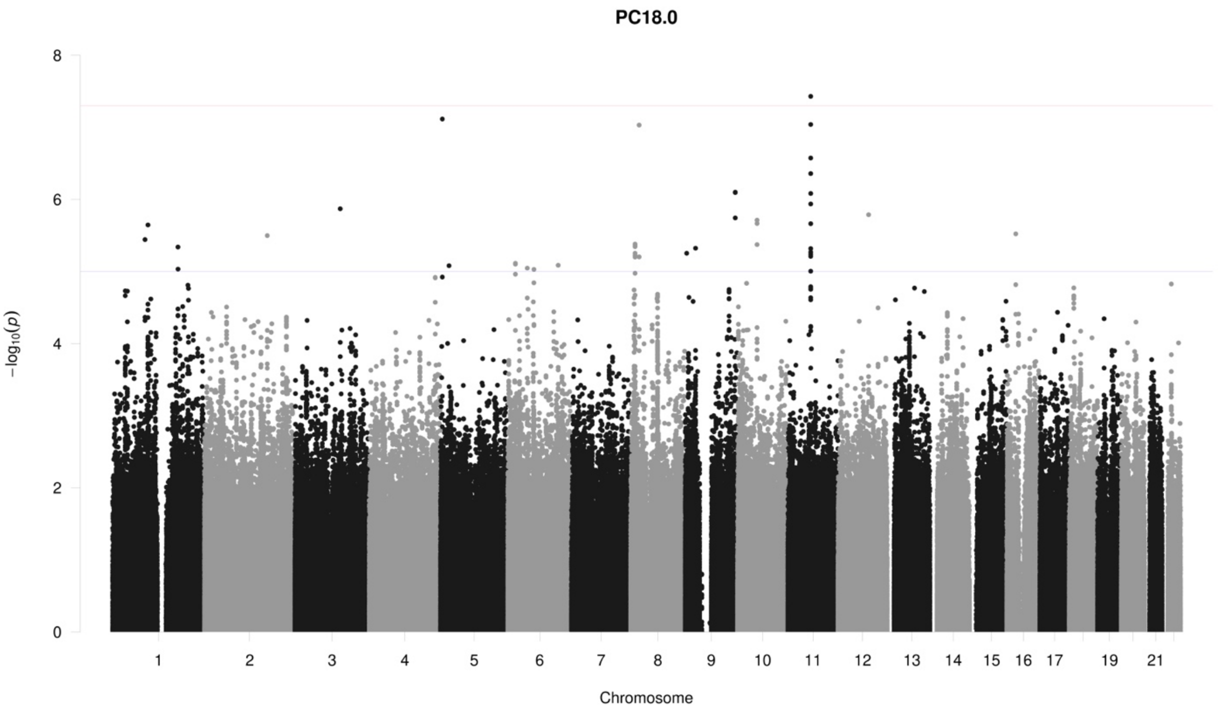
PC18.0
$-\log_{10}(p)$
0
2
4
6
8
19
21
Chromosome

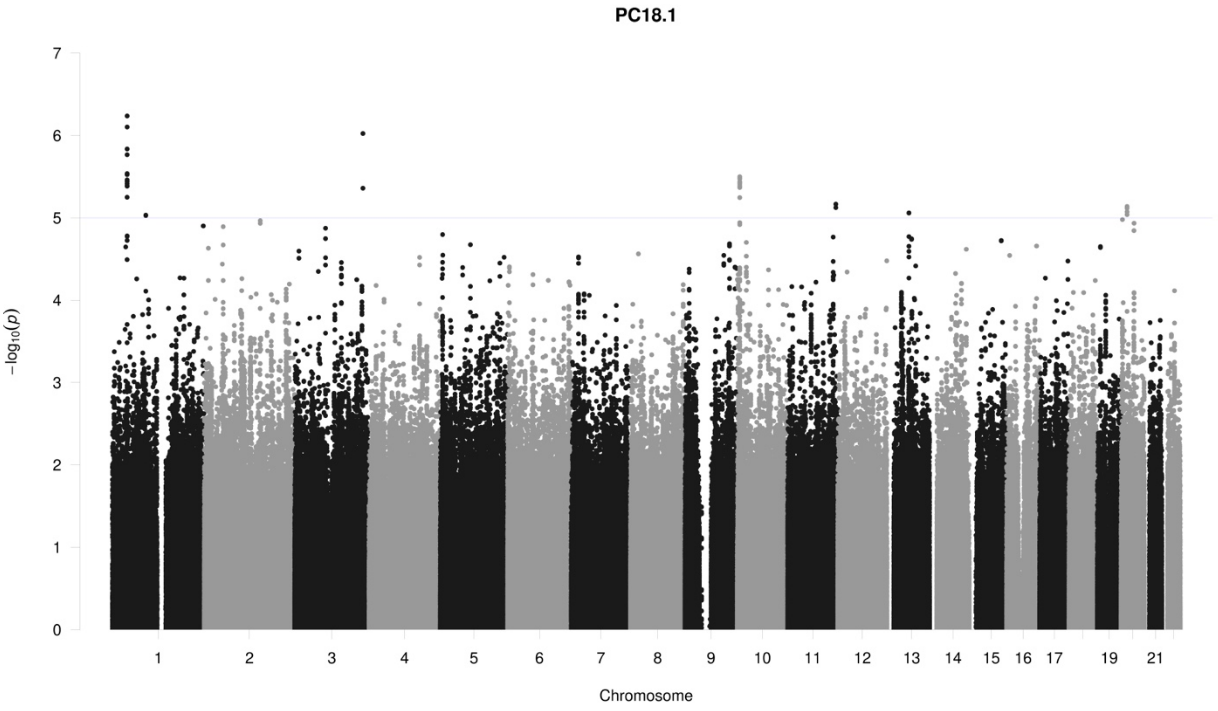
PC18.1
$-\log_{10}(p)$
0
1
2
3
4
5
6
7
19
21
Chromosome

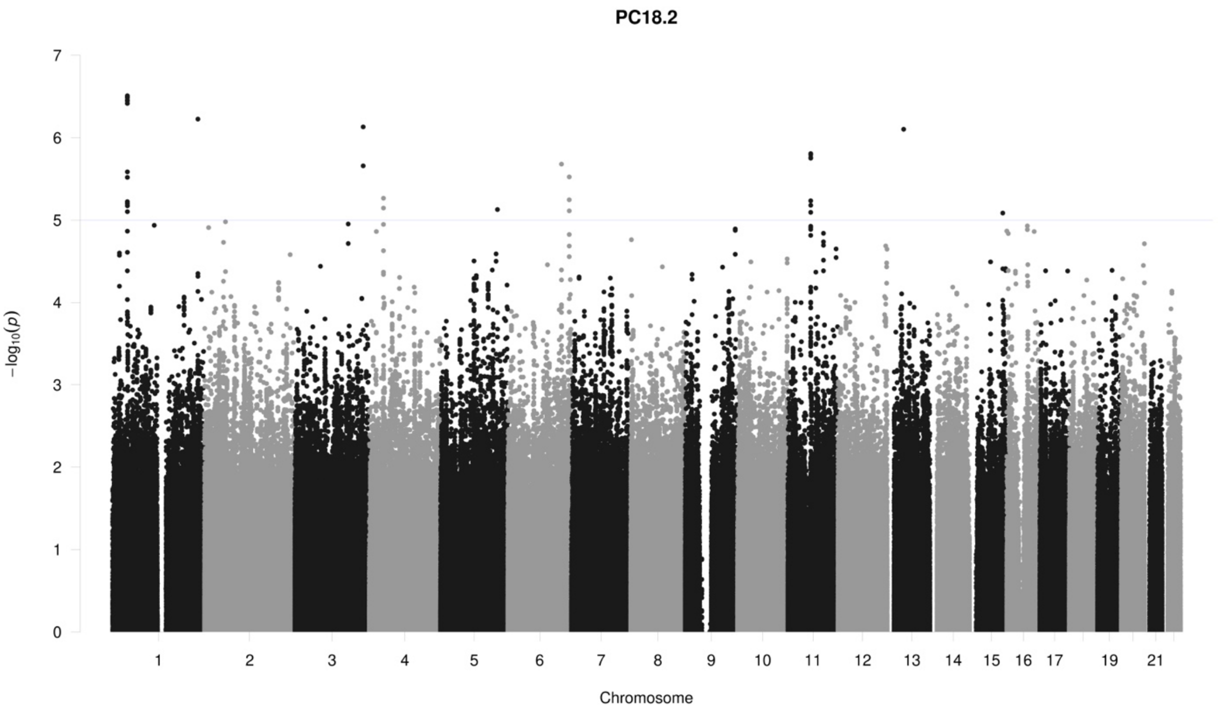
PC18.2
−log10(p)
Chromosome
19
21

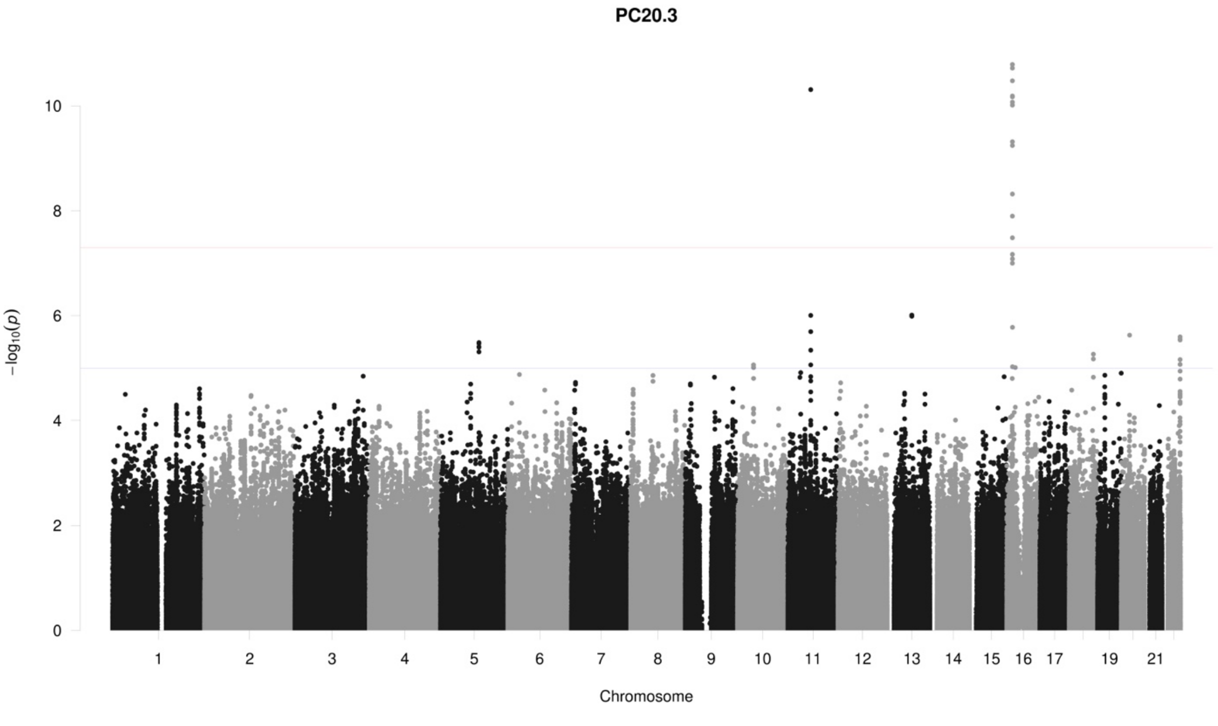
PC20.3
−log10(p)
Chromosome
19
21

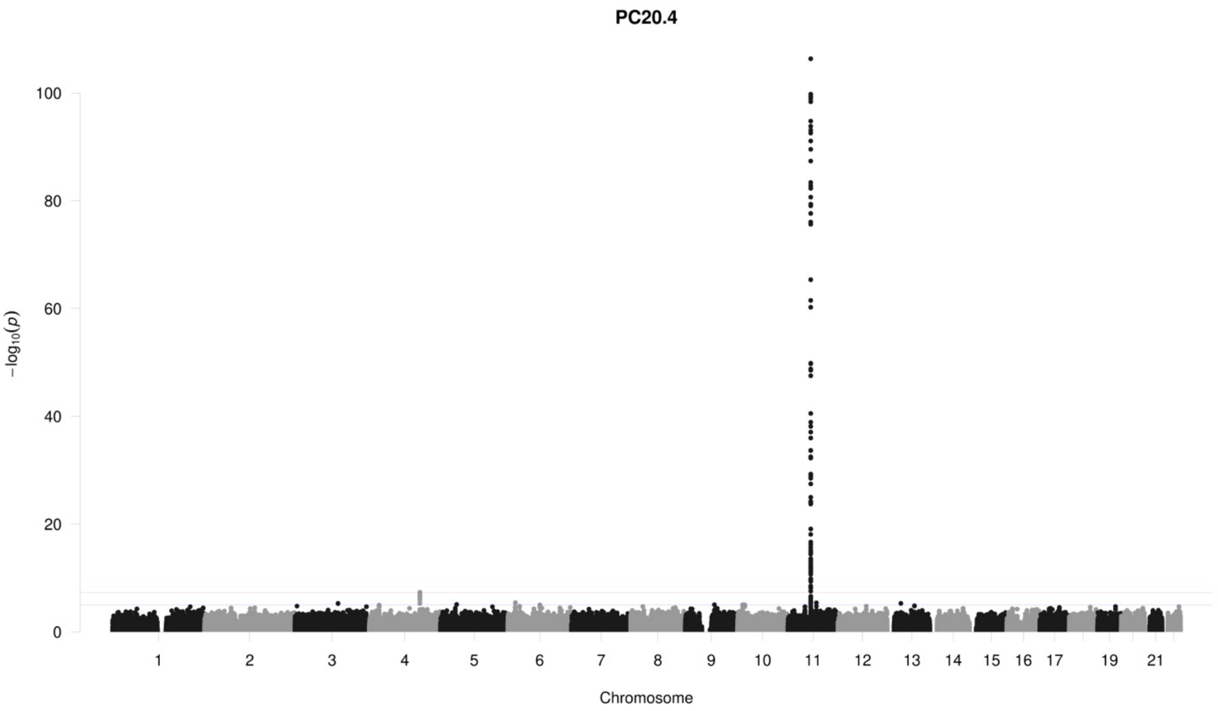
PC20.4
100
80
60
40
20
0
−log10(p)
1 2 3 4 5 6 7 8 9 10 11 12 13 14 15 16 17 19 21
Chromosome

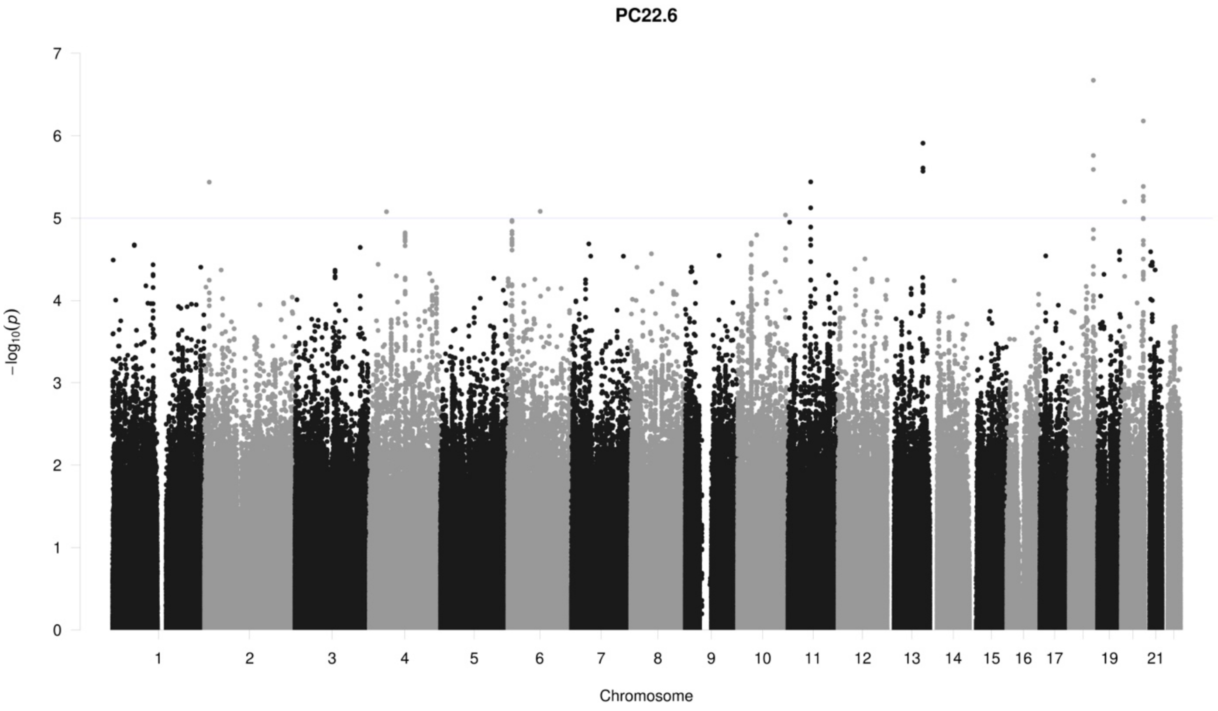
PC22.6
7
6
5
4
3
2
1
0
−log10(p)
1 2 3 4 5 6 7 8 9 10 11 12 13 14 15 16 17 19 21
Chromosome

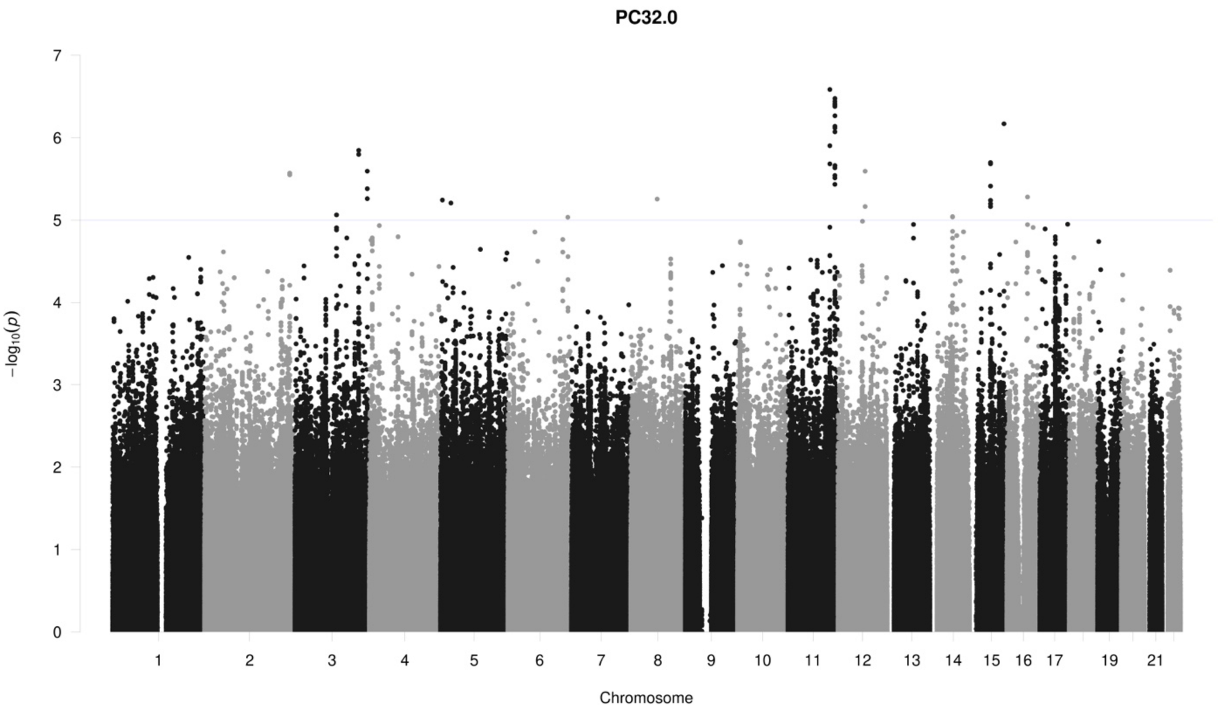

PC32.0
−log10(p)
Chromosome
19
21

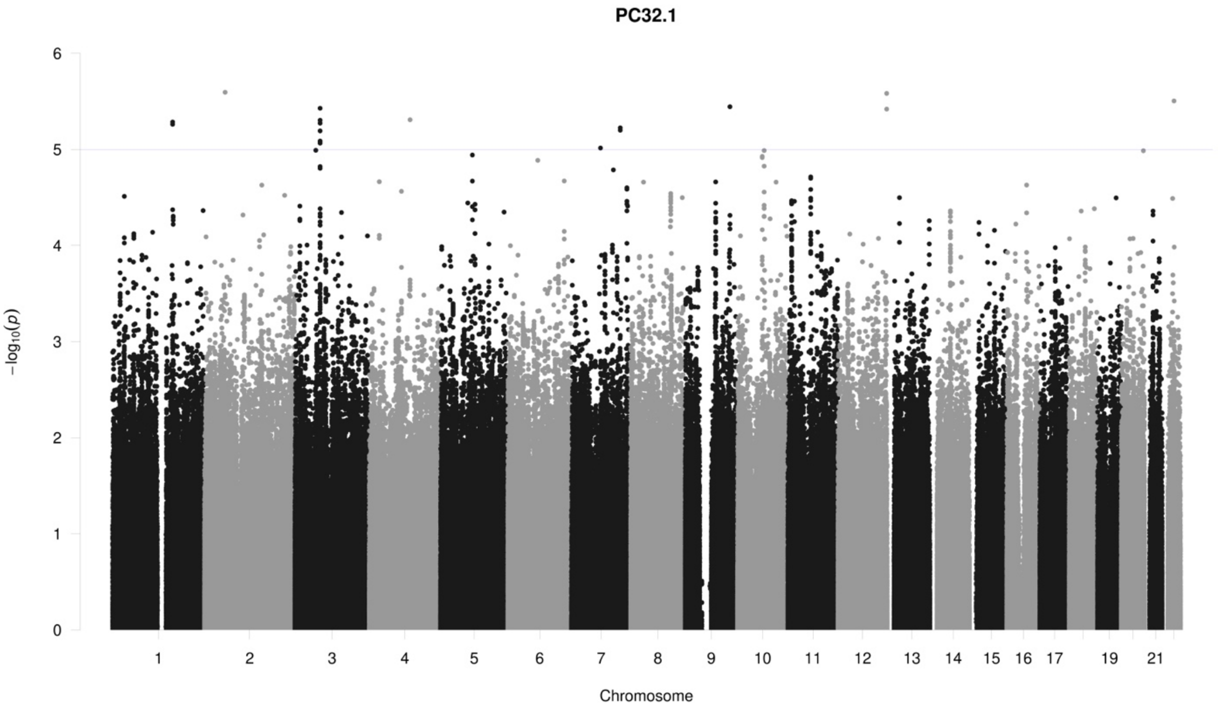

PC32.1
−log10(p)
Chromosome
19
21

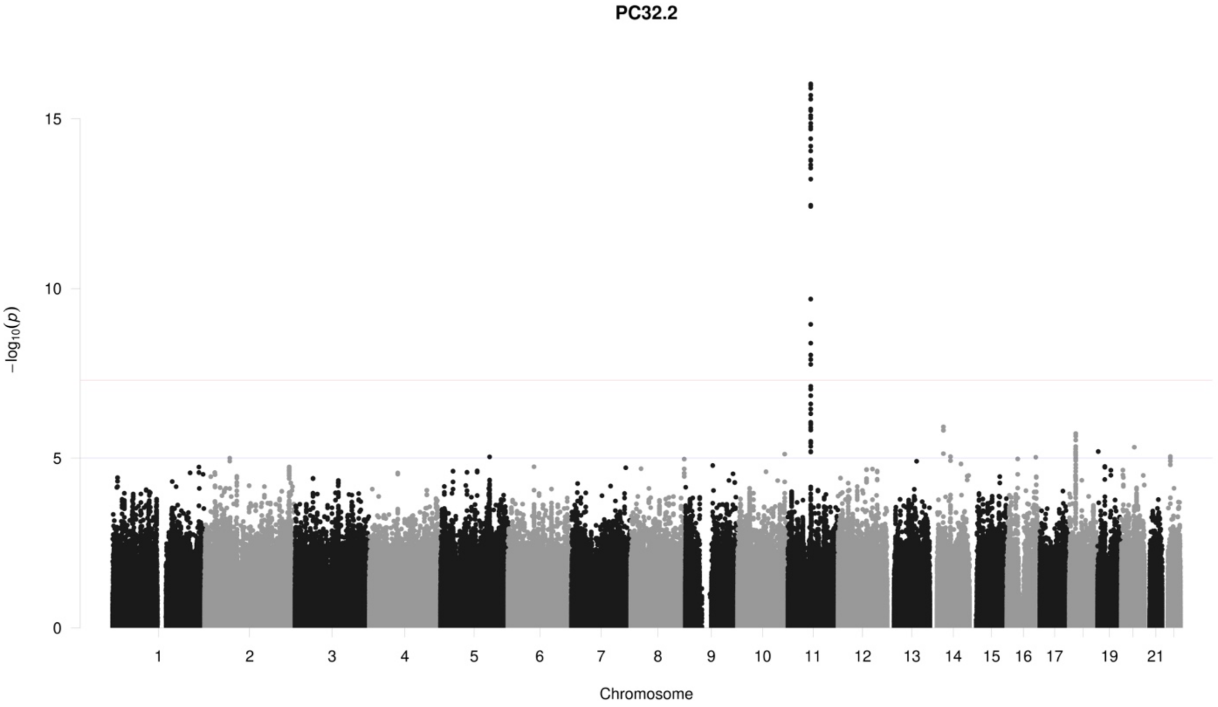
PC32.2
15
10
5
0
−log10(p)
19
21
Chromosome

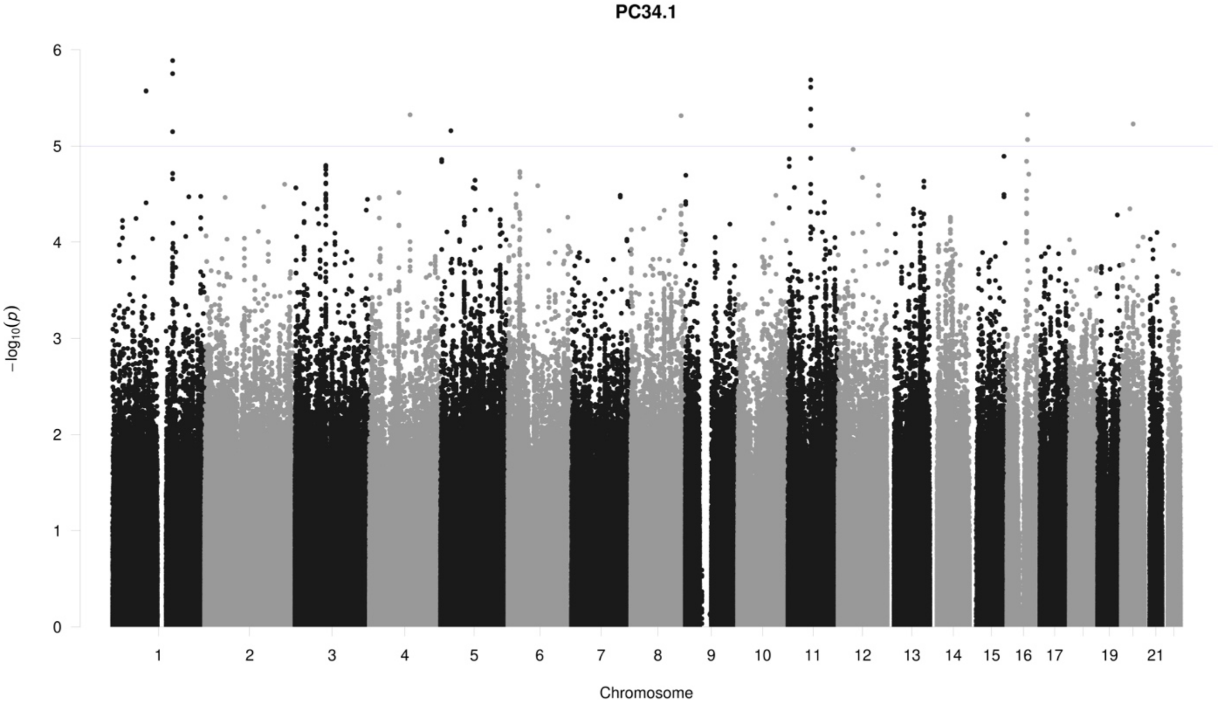
PC34.1
6
5
4
3
2
1
0
−log10(p)
19
21
Chromosome

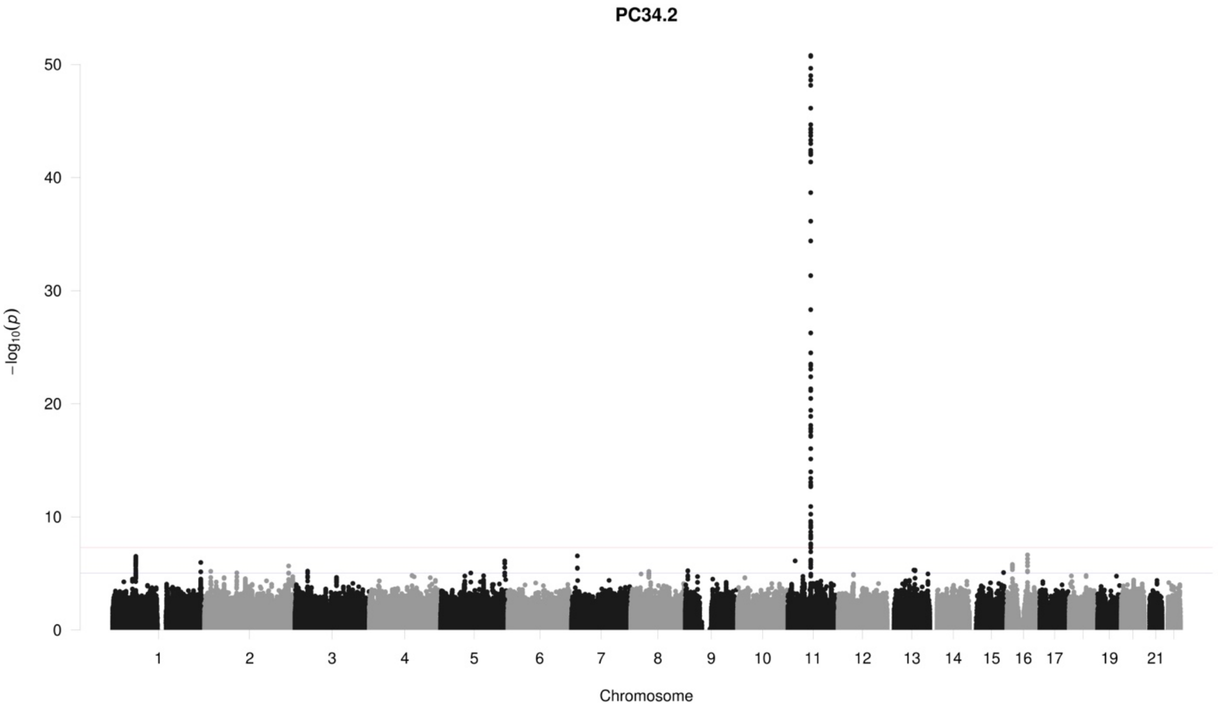
PC34.2
50
40
30
20
10
0
−log10(p)
19
21
Chromosome

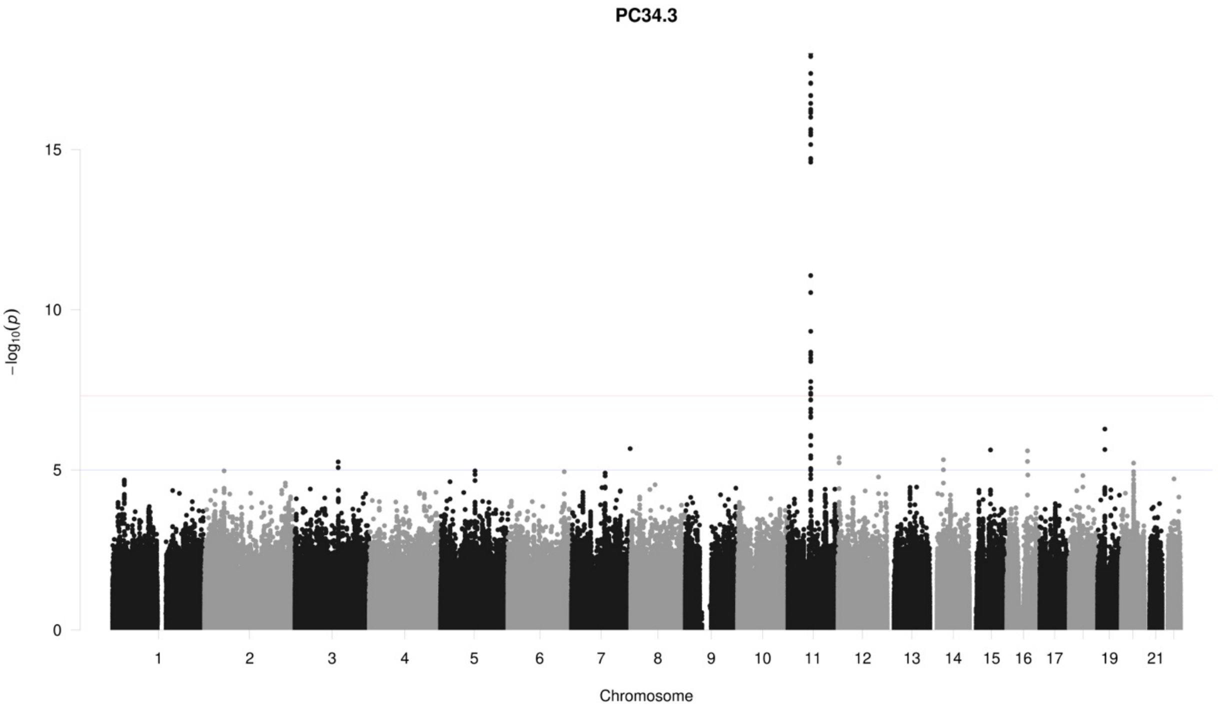
PC34.3
15
10
5
0
−log10(p)
19
21
Chromosome

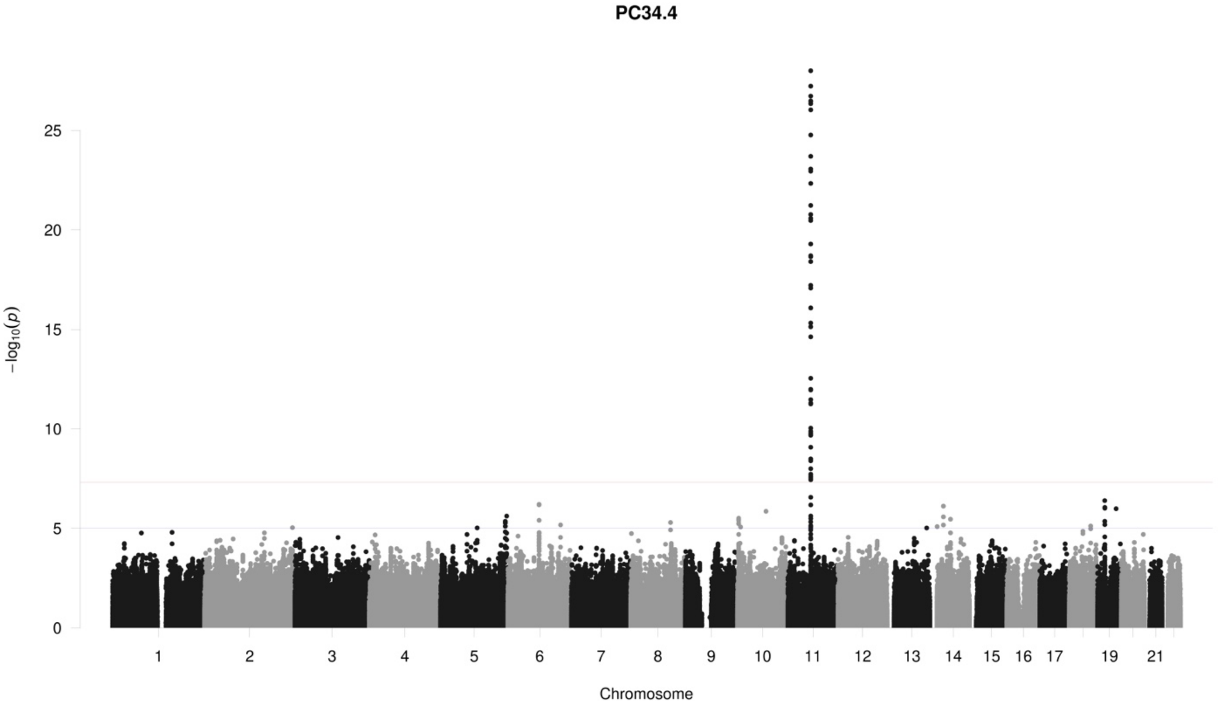
PC34.4
−log10(p)
25
20
15
10
5
19
21
Chromosome

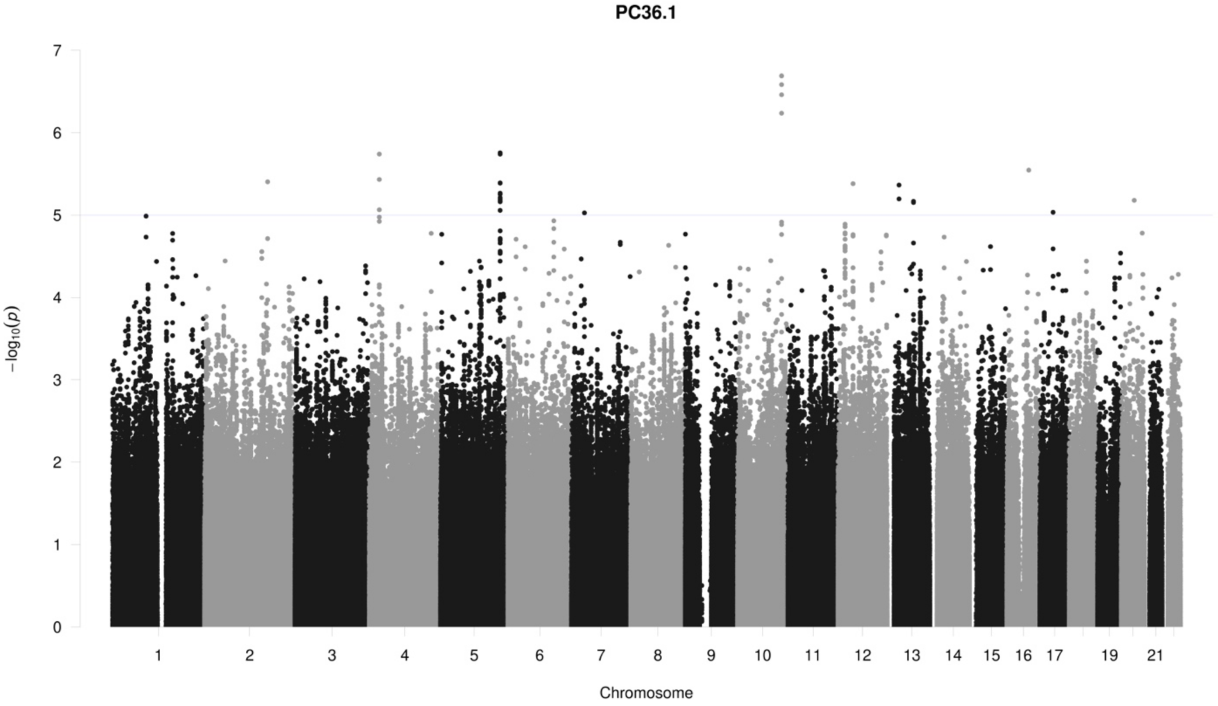
PC36.1
−log10(p)
7
6
5
4
3
2
1
19
21
Chromosome

**PC36.2**

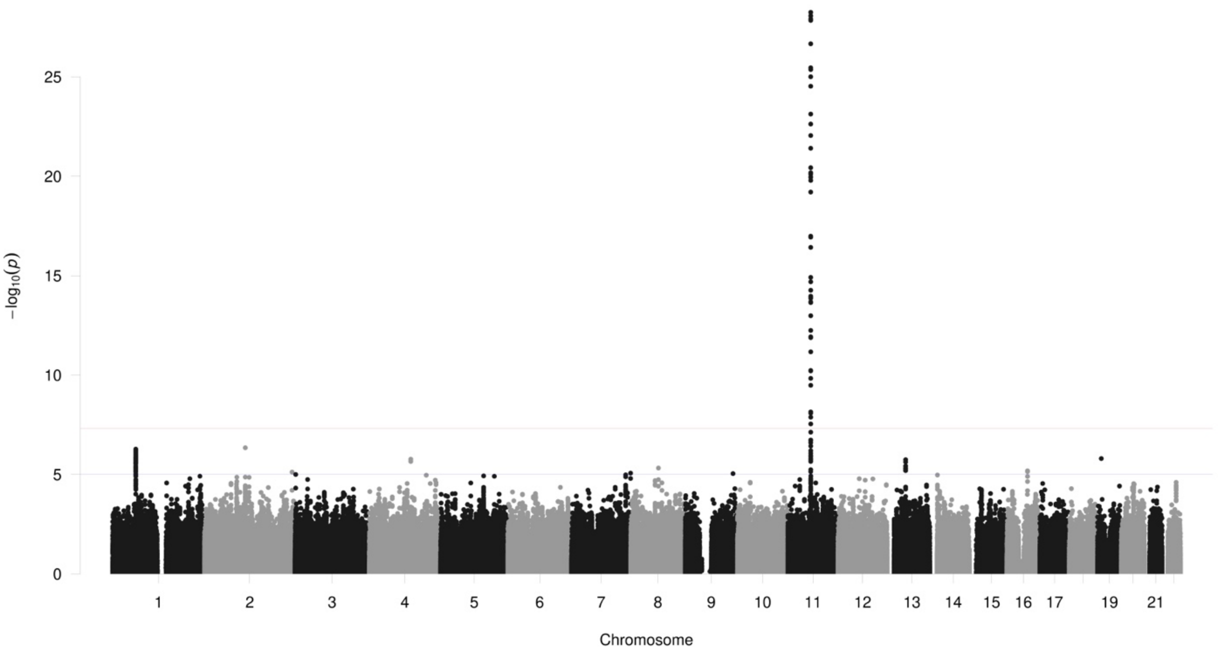

25
20
15
10
5
0
−log10(p)
19
21
Chromosome

**PC36.3**

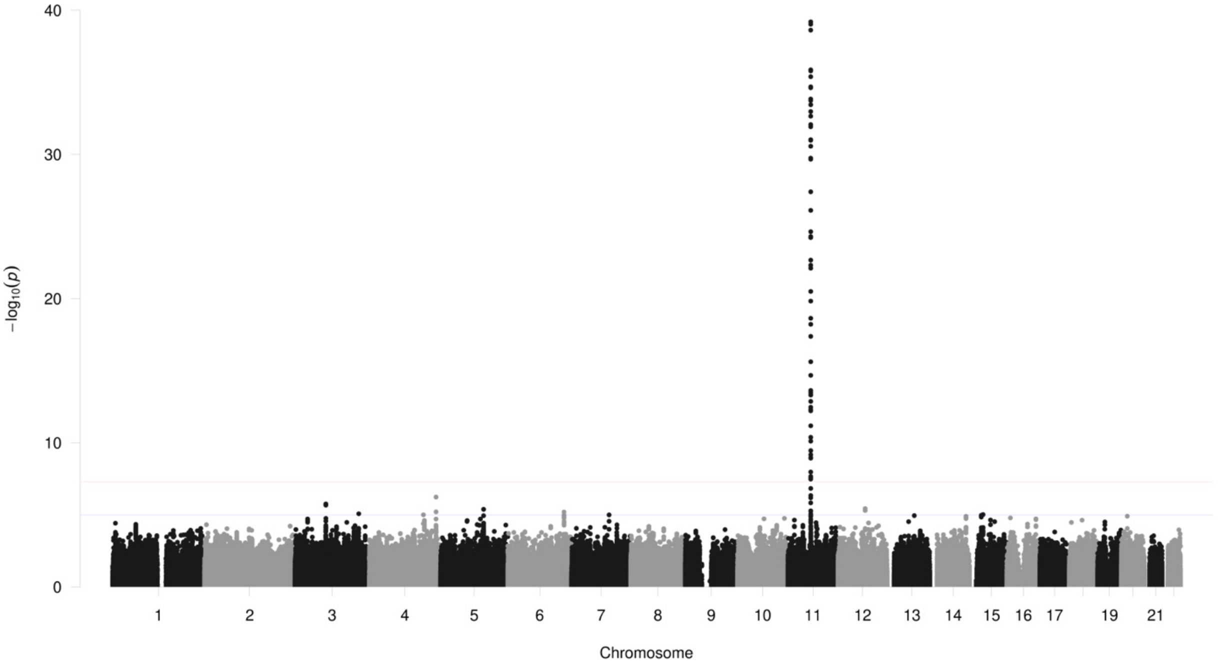

40
30
20
10
0
−log10(p)
19
21
Chromosome

PC36.4

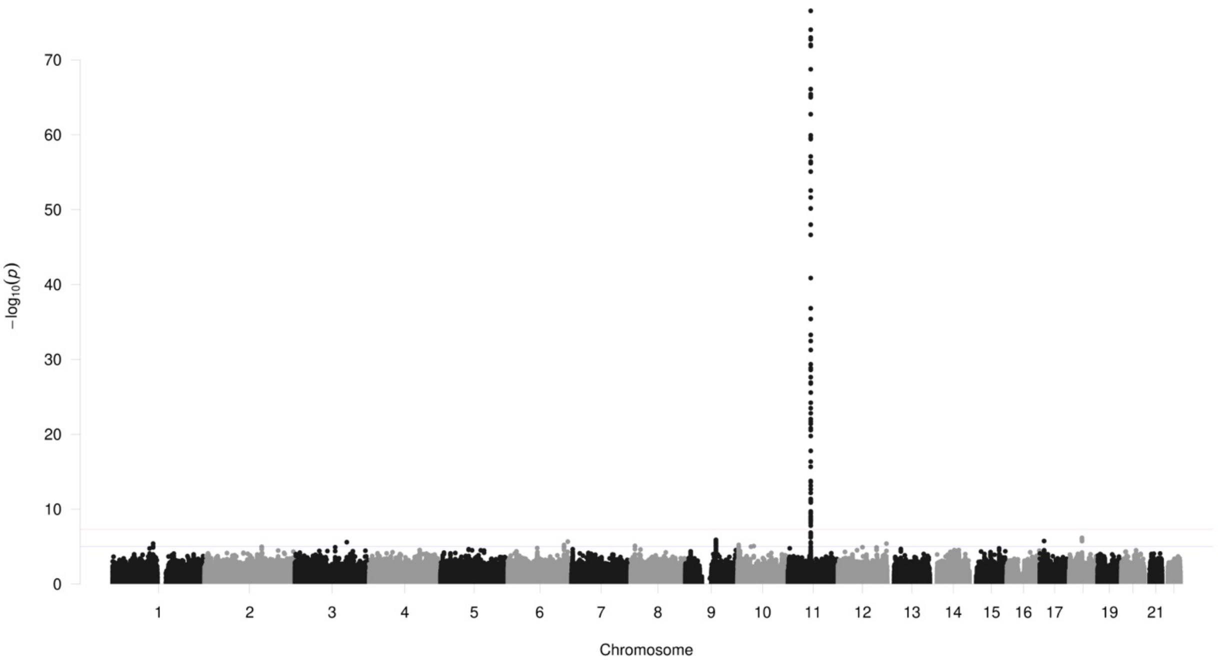

70
60
50
40
30
20
10
0
−log10(p)
19
21
Chromosome

PC36.5

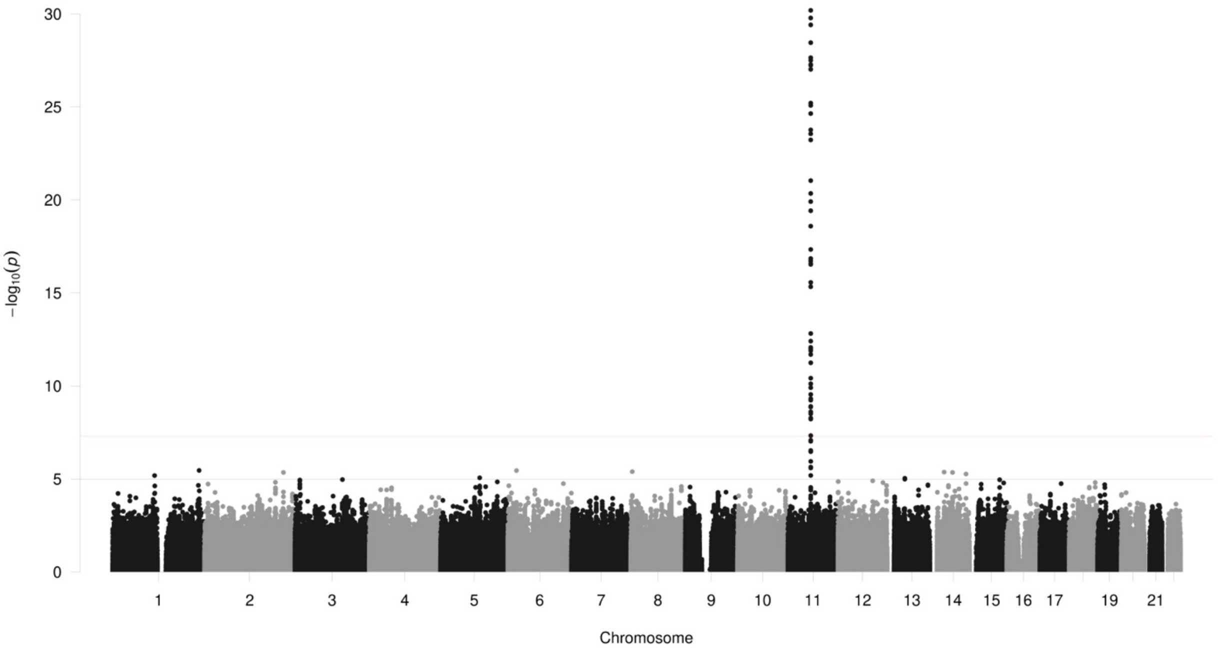

30
25
20
15
10
5
0
−log10(p)
19
21
Chromosome

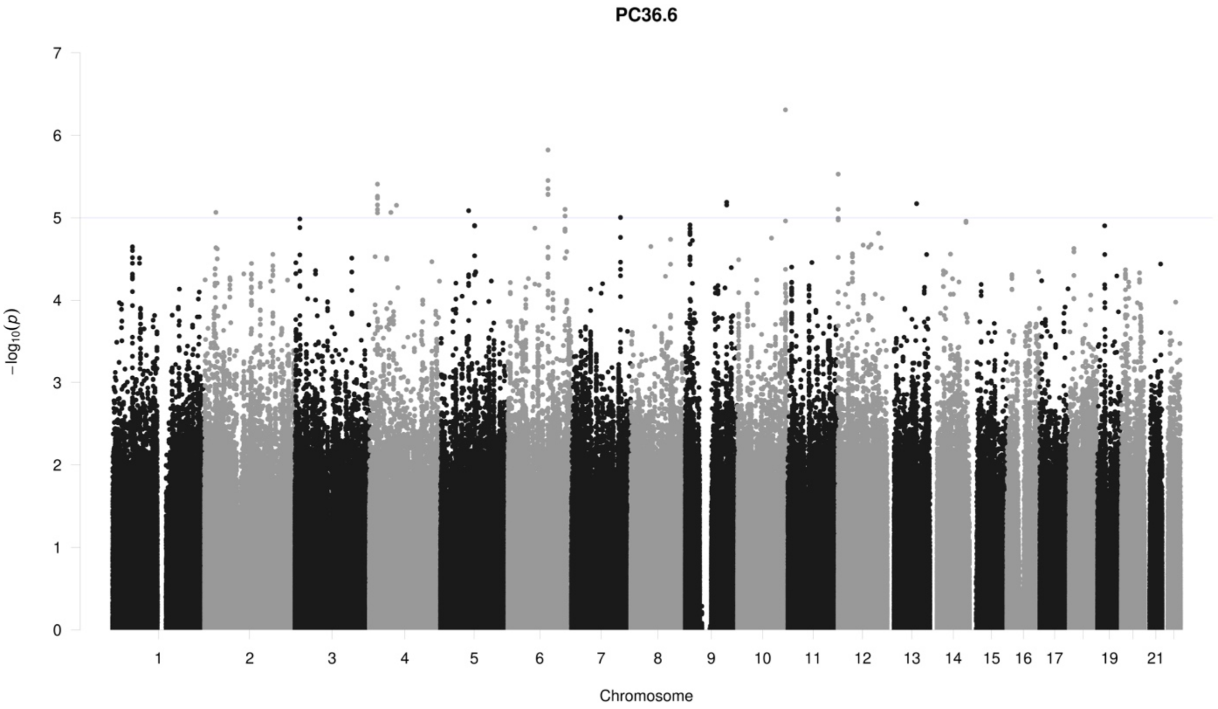
PC36.6
−log10(p)
Chromosome

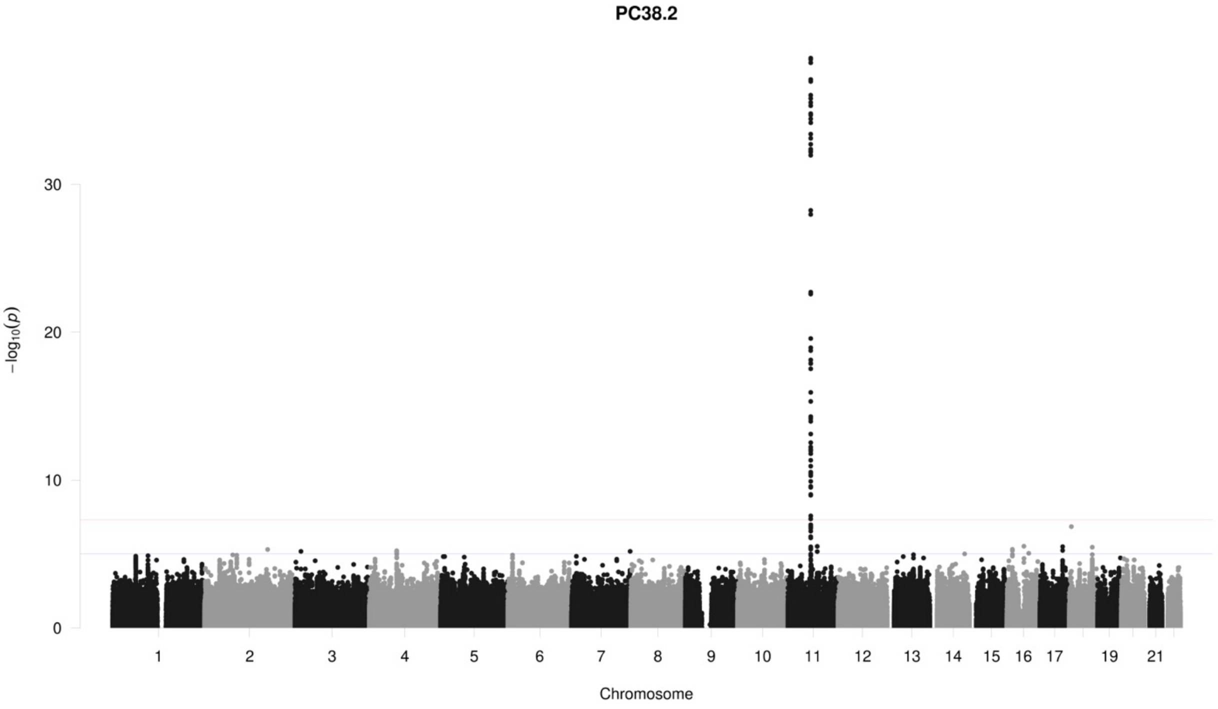
PC38.2
−log10(p)
Chromosome

**PC38.3**

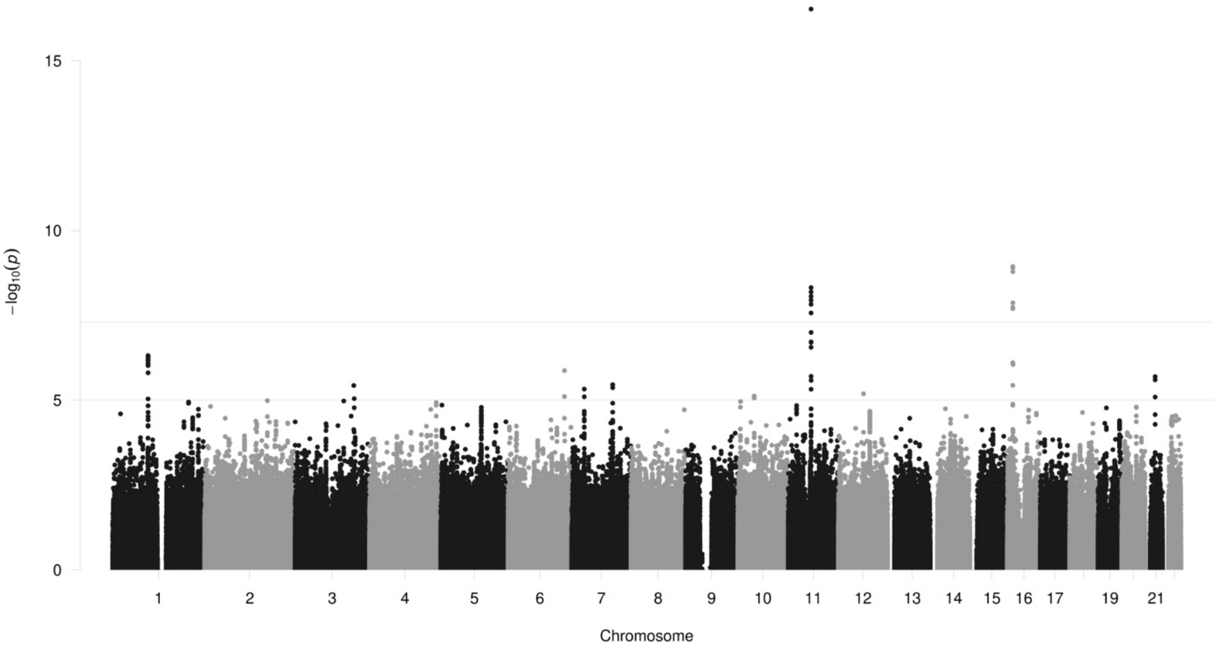

15
10
5
0
−log10(p)
19
21
Chromosome

**PC38.4**

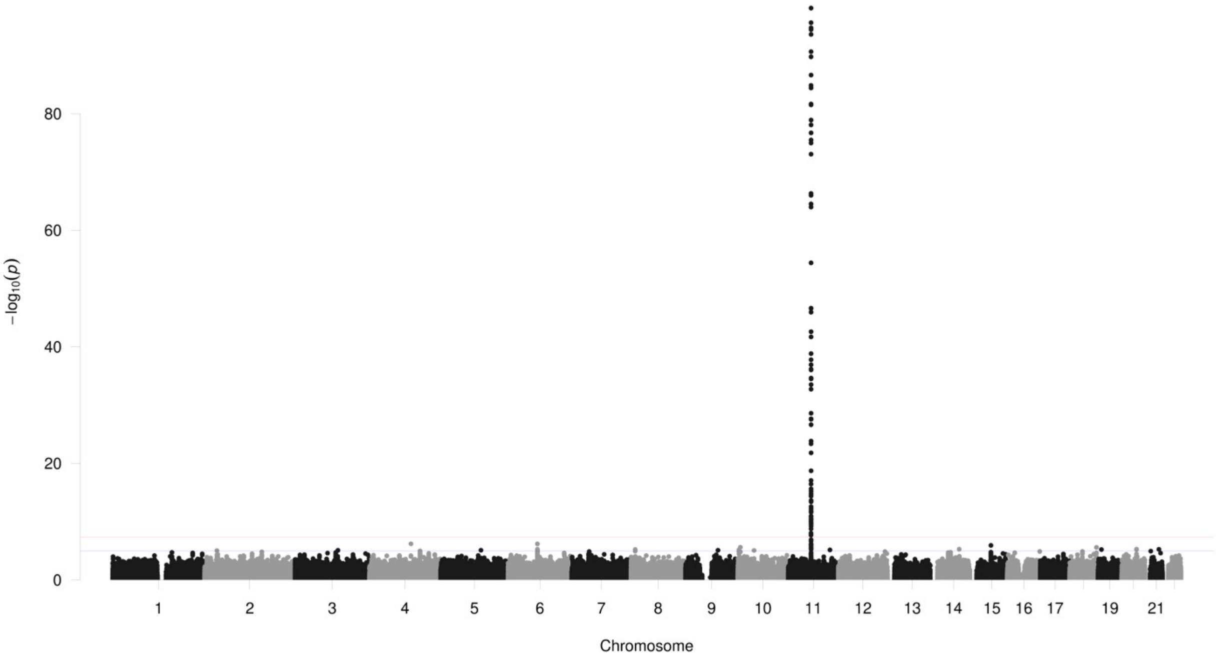

80
60
40
20
0
−log10(p)
19
21
Chromosome

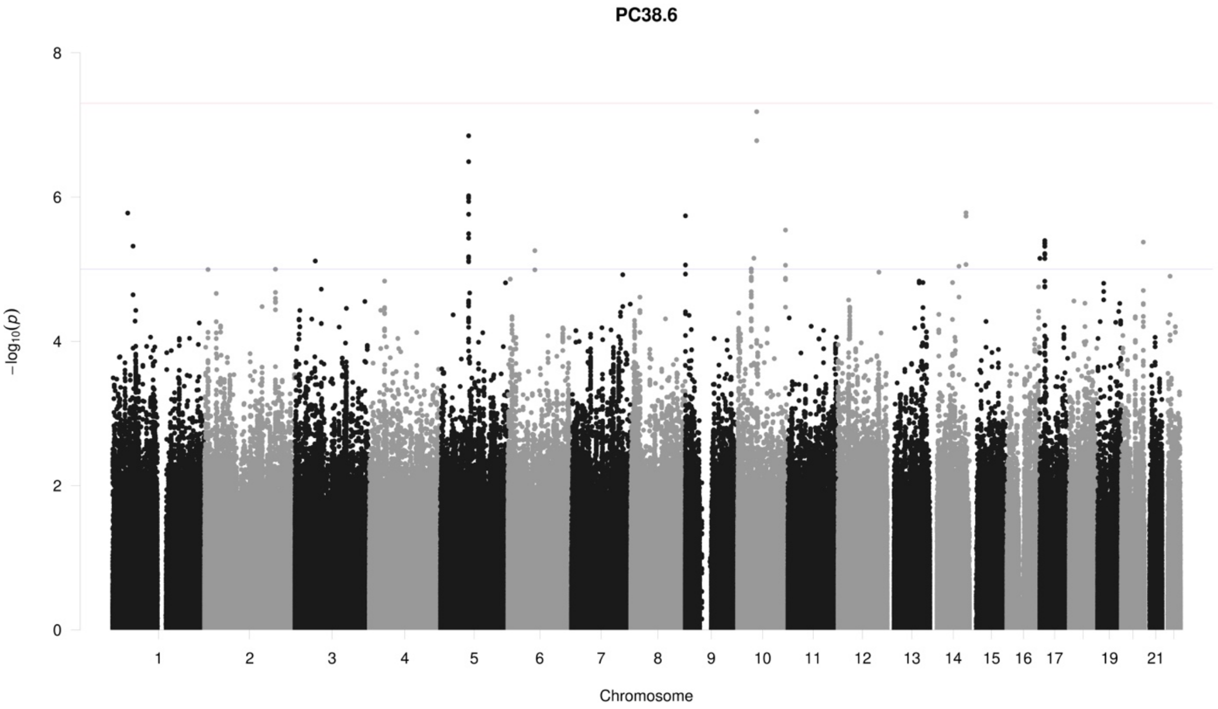
PC38.6
−log10(p)
Chromosome
0
2
4
6
8
19
21

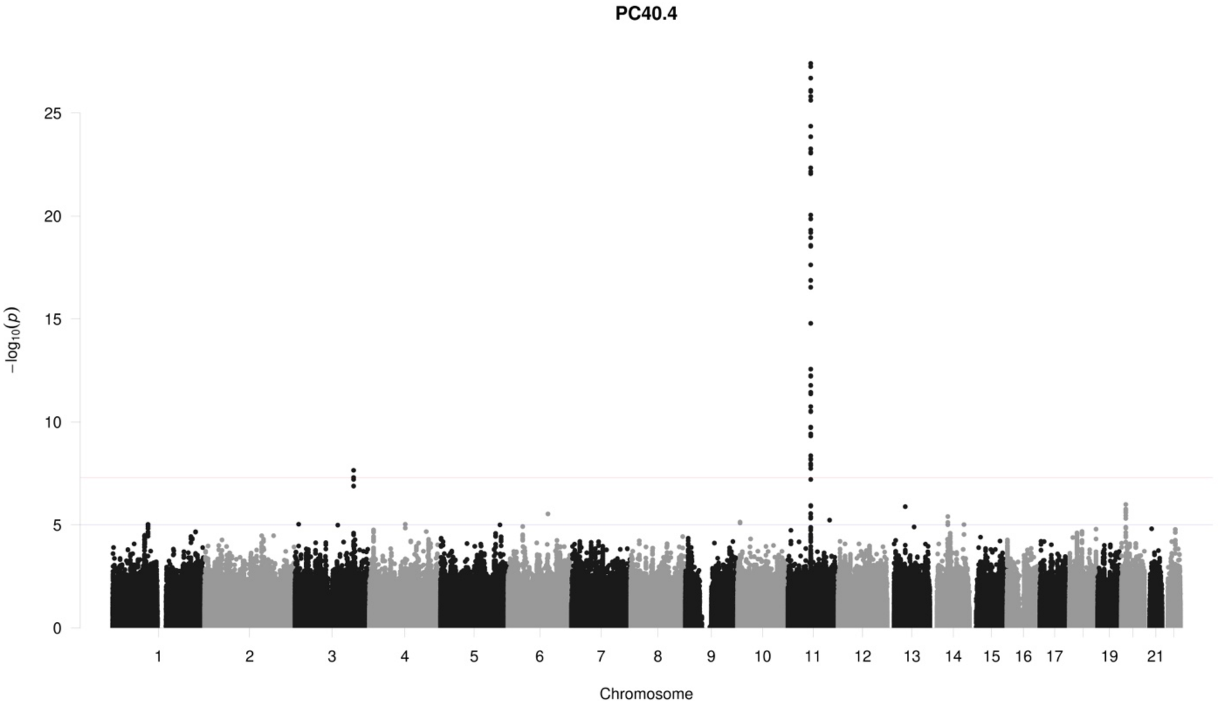
PC40.4
−log10(p)
Chromosome
0
5
10
15
20
25
19
21

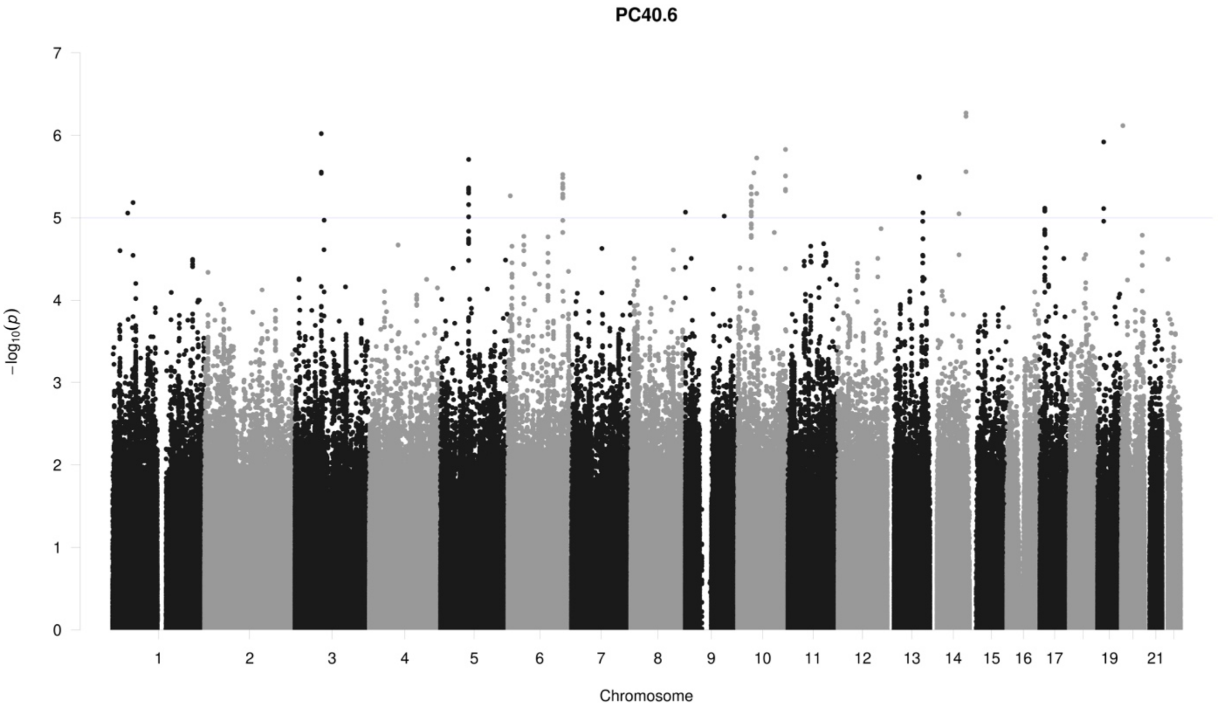
PC40.6
−log10(p)
0
1
2
3
4
5
6
7
19
21
Chromosome

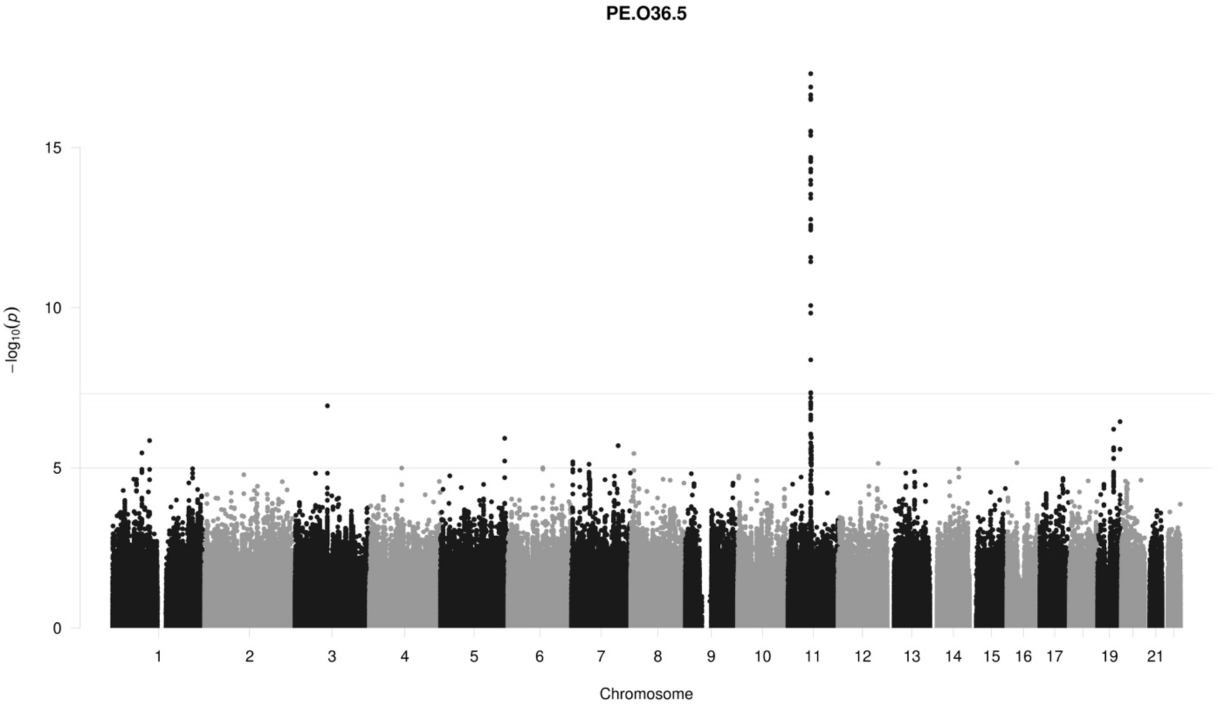
PE.O36.5
−log10(p)
0
5
10
15
19
21
Chromosome

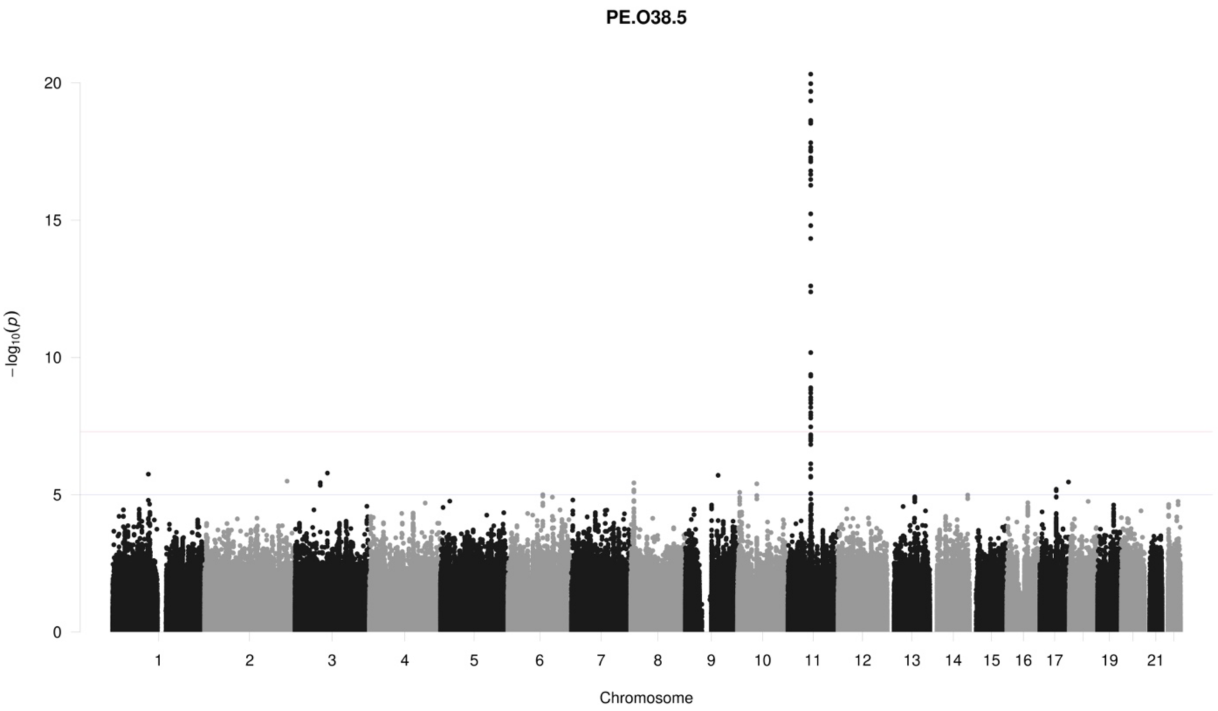
PE.O38.5
20
15
10
5
0
−log10(p)
19
21
Chromosome

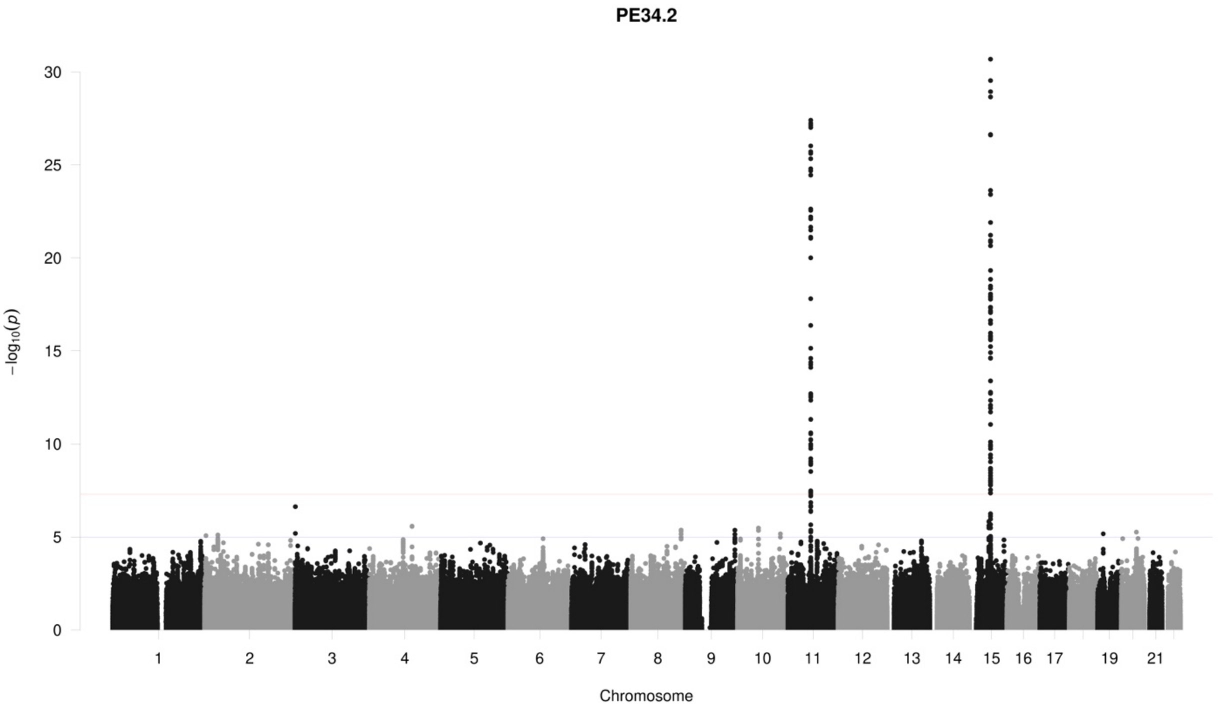
PE34.2
30
25
20
15
10
5
0
−log10(p)
19
21
Chromosome

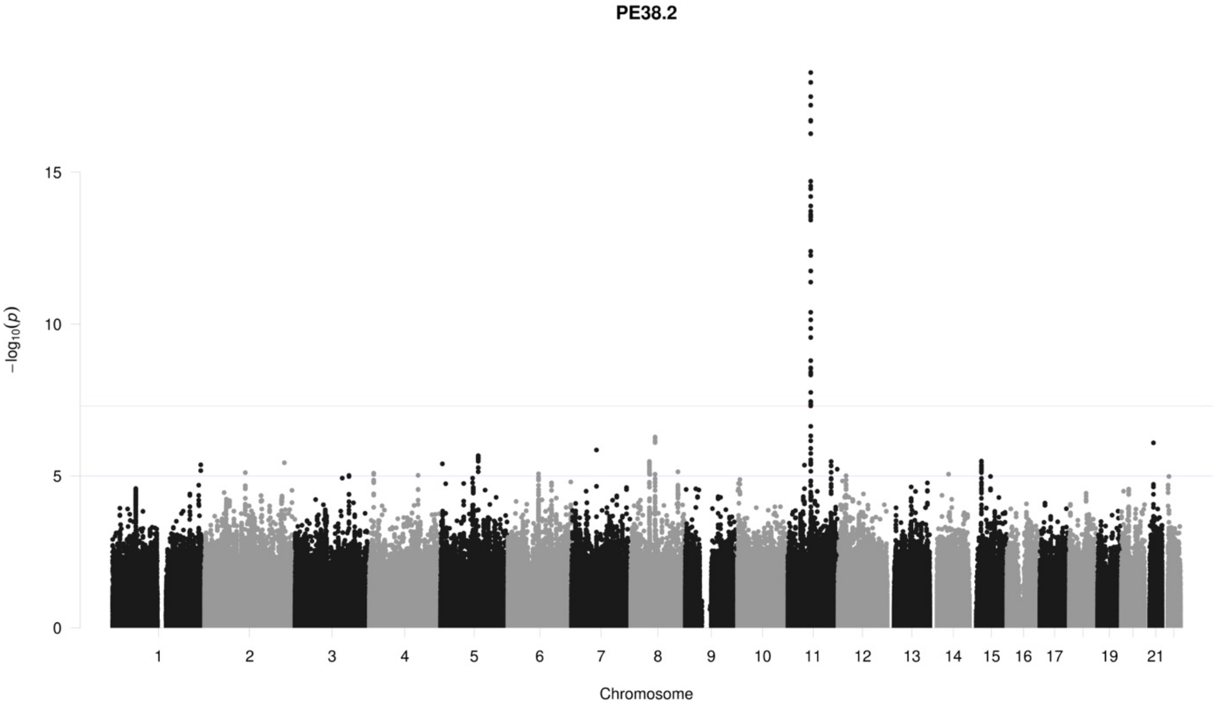
PE38.2
−log10(p)
15
10
5
19
21
Chromosome

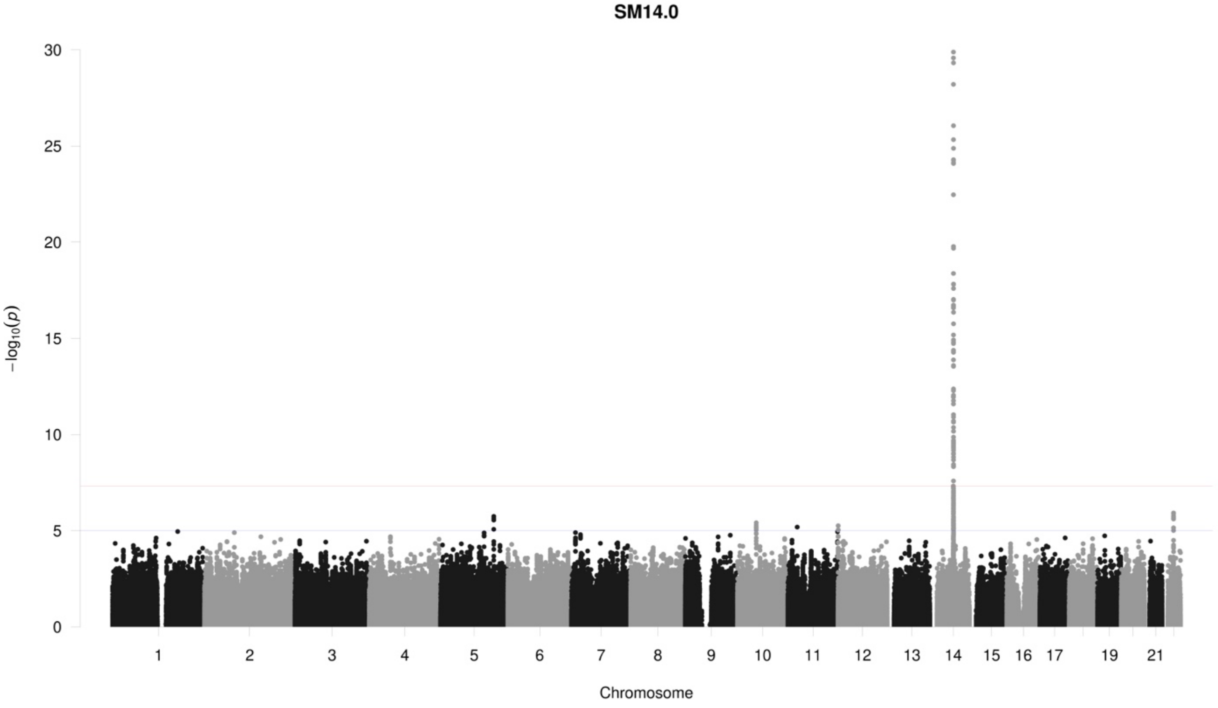
SM14.0
−log10(p)
30
25
20
15
10
5
19
21
Chromosome

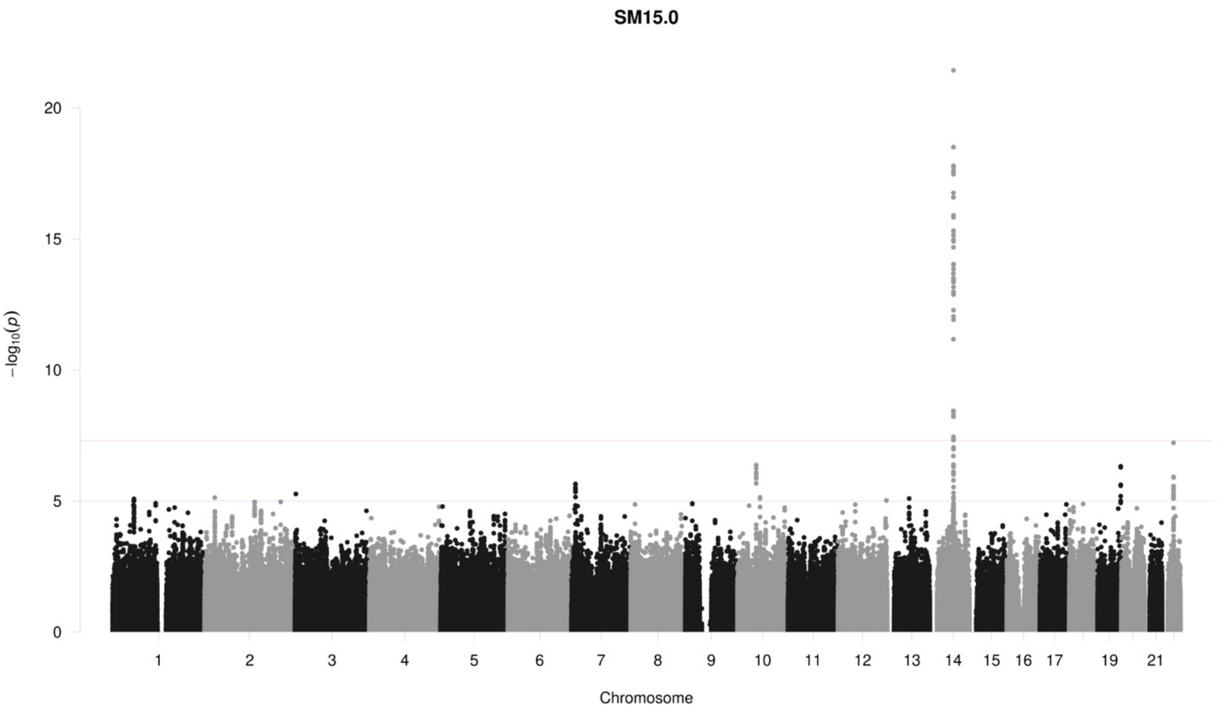

SM15.0
20
15
10
5
0
−log10(p)
19
21
Chromosome

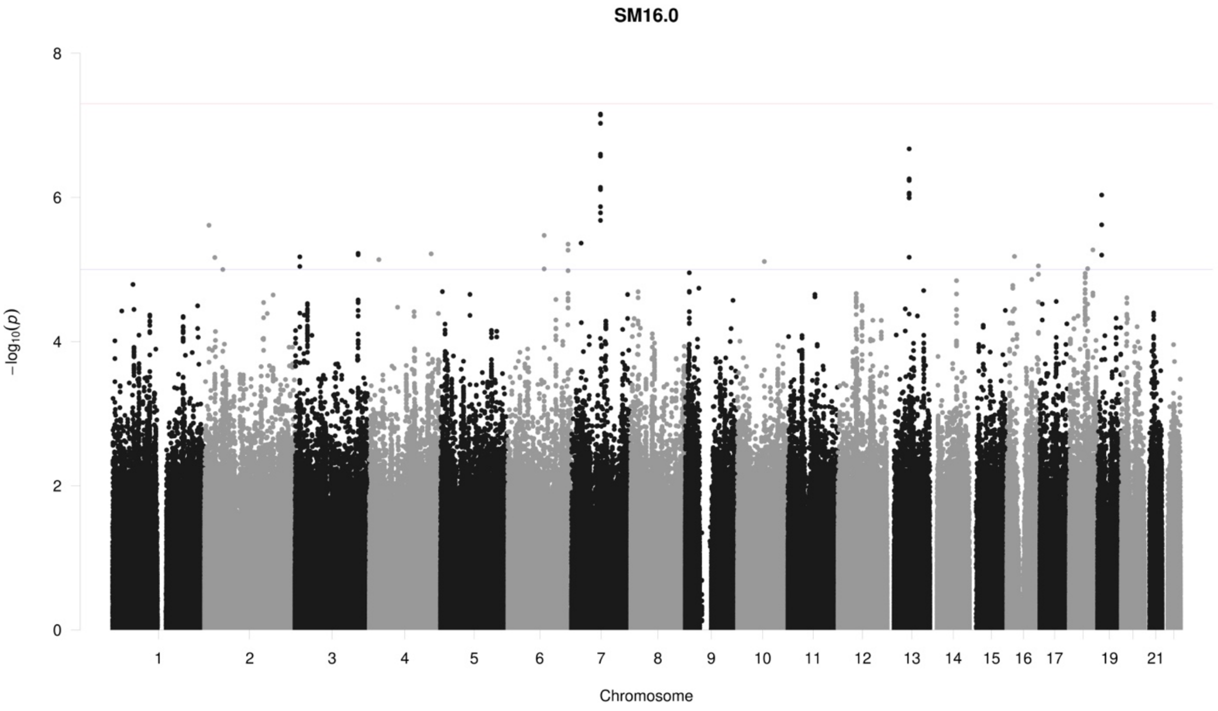

SM16.0
8
6
4
2
0
−log10(p)
19
21
Chromosome

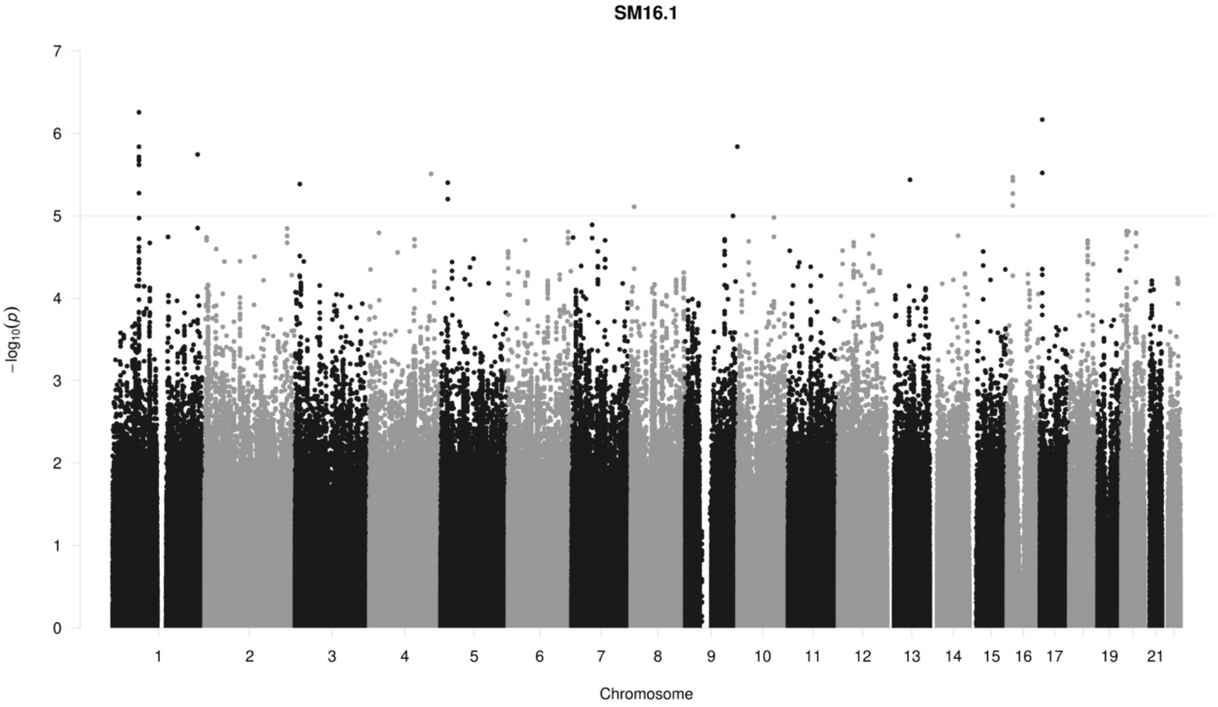
SM16.1
7
6
5
4
3
2
1
0
−log10(p)
19
21
Chromosome

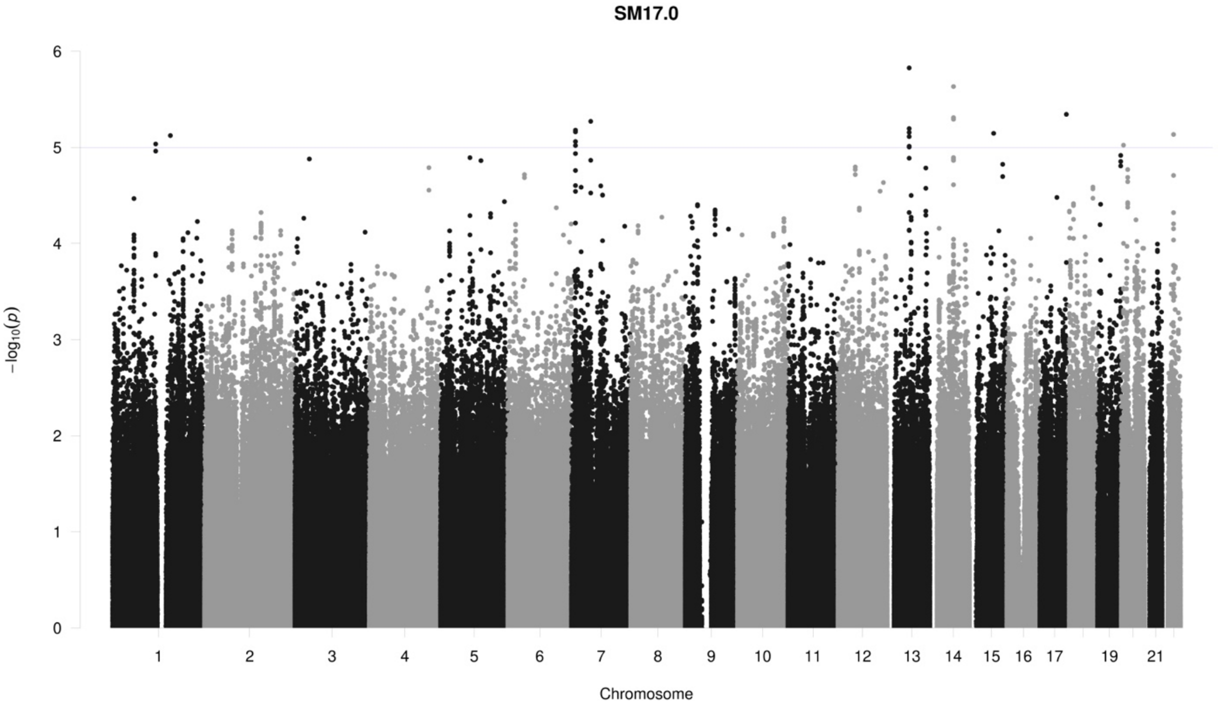
SM17.0
6
5
4
3
2
1
0
−log10(p)
19
21
Chromosome

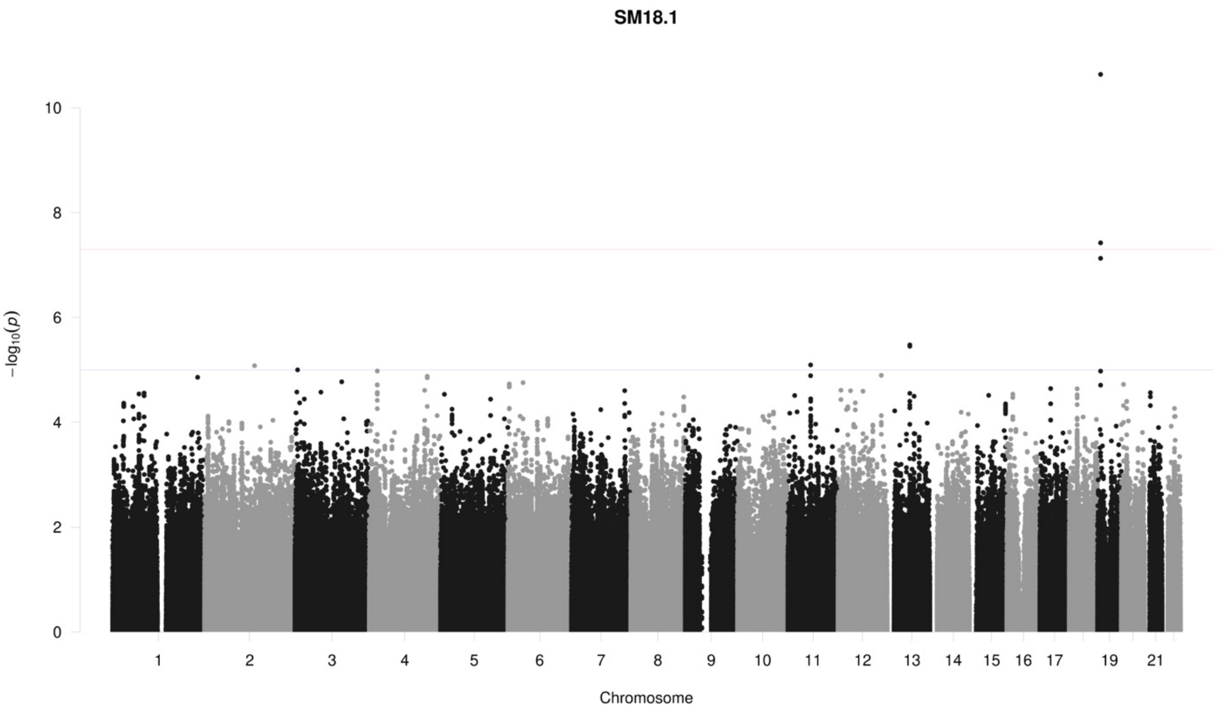
SM18.1
−log10(p)
0
2
4
6
8
10
19
21
Chromosome

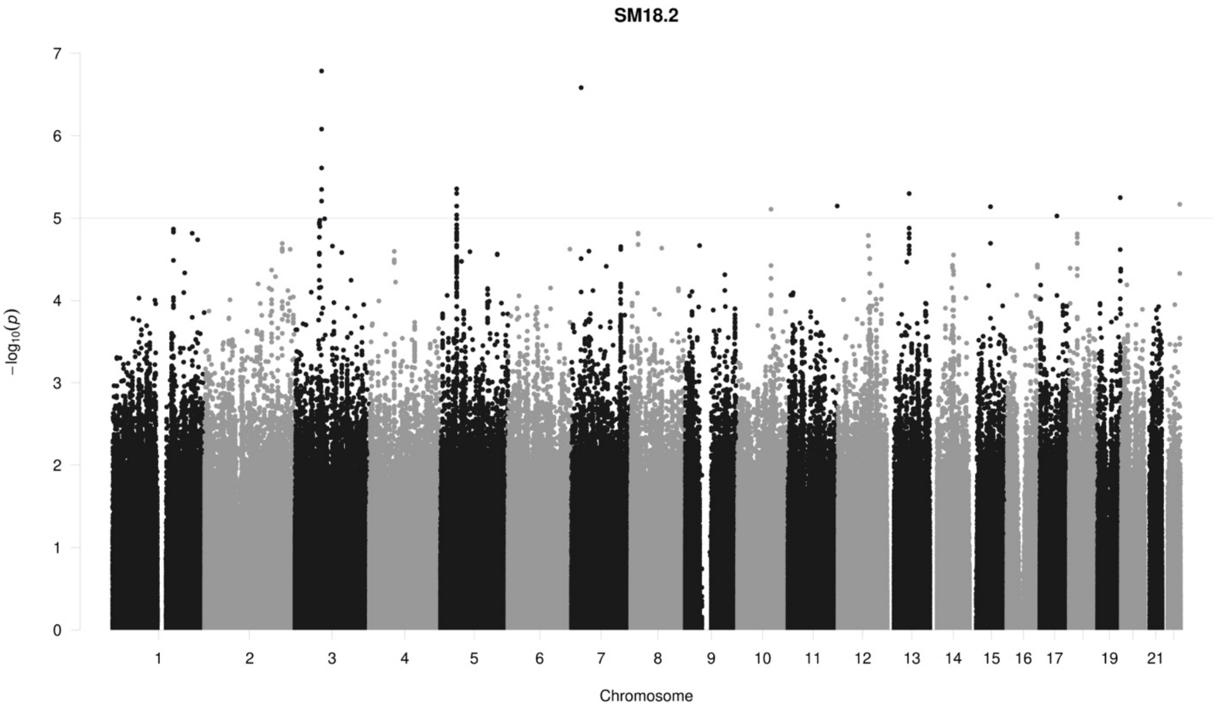
SM18.2
−log10(p)
0
1
2
3
4
5
6
7
19
21
Chromosome

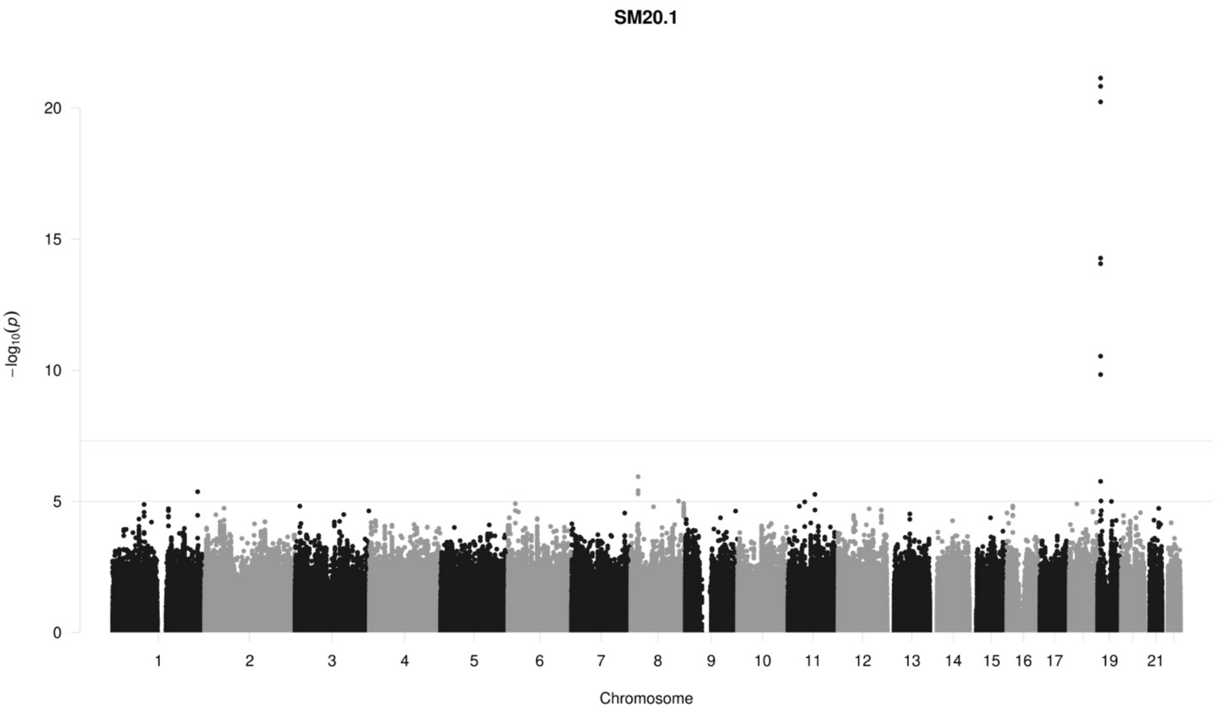
SM20.1
20
15
10
5
0
−log10(p)
19
21
Chromosome

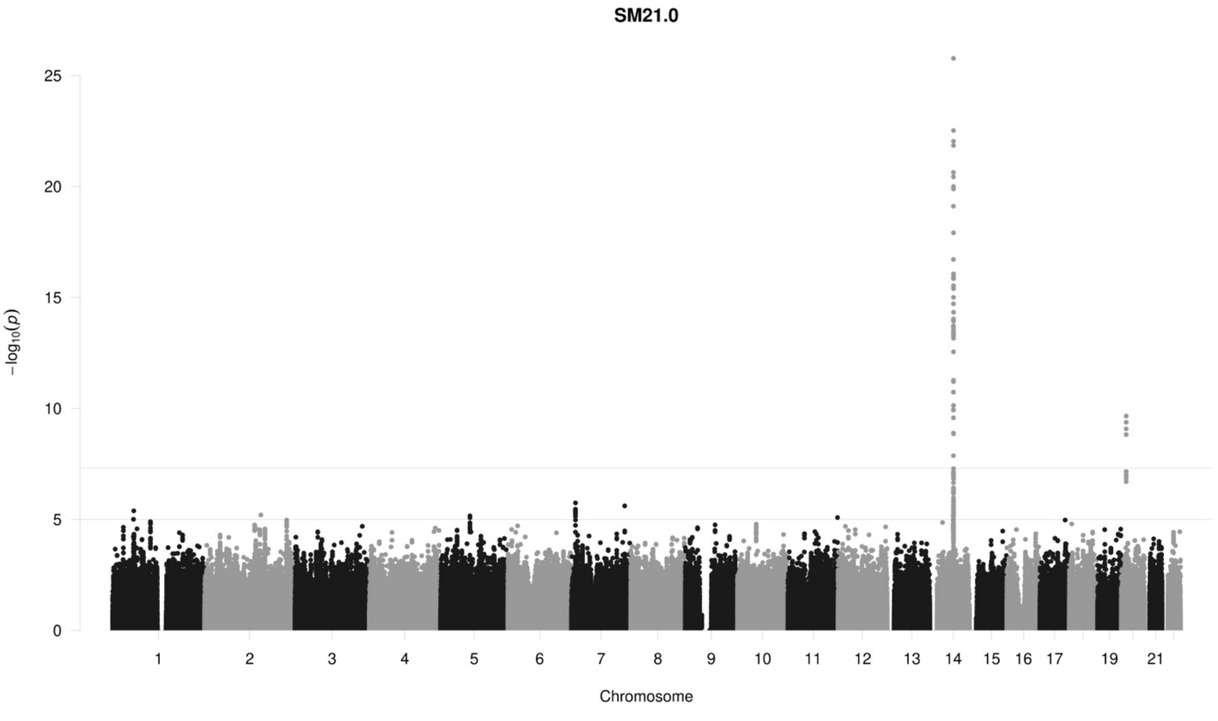
SM21.0
25
20
15
10
5
0
−log10(p)
19
21
Chromosome

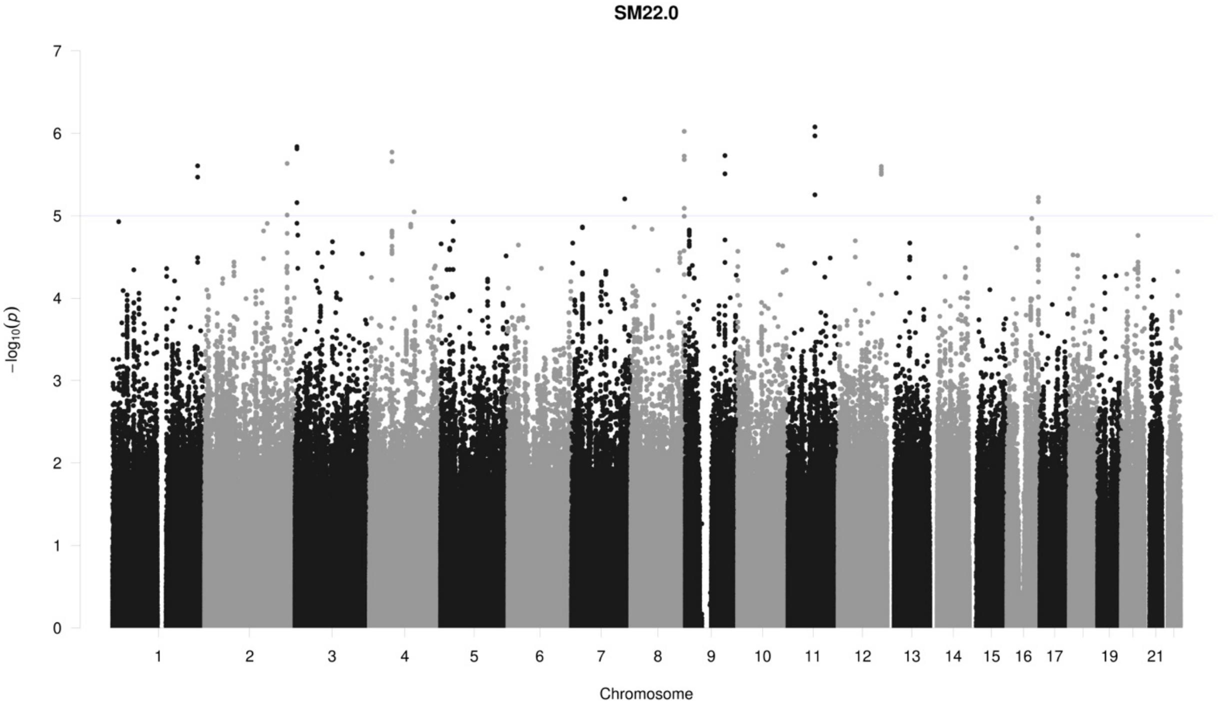
SM22.0
−log10(p)
Chromosome
1 2 3 4 5 6 7 8 9 10 11 12 13 14 15 16 17 19 21

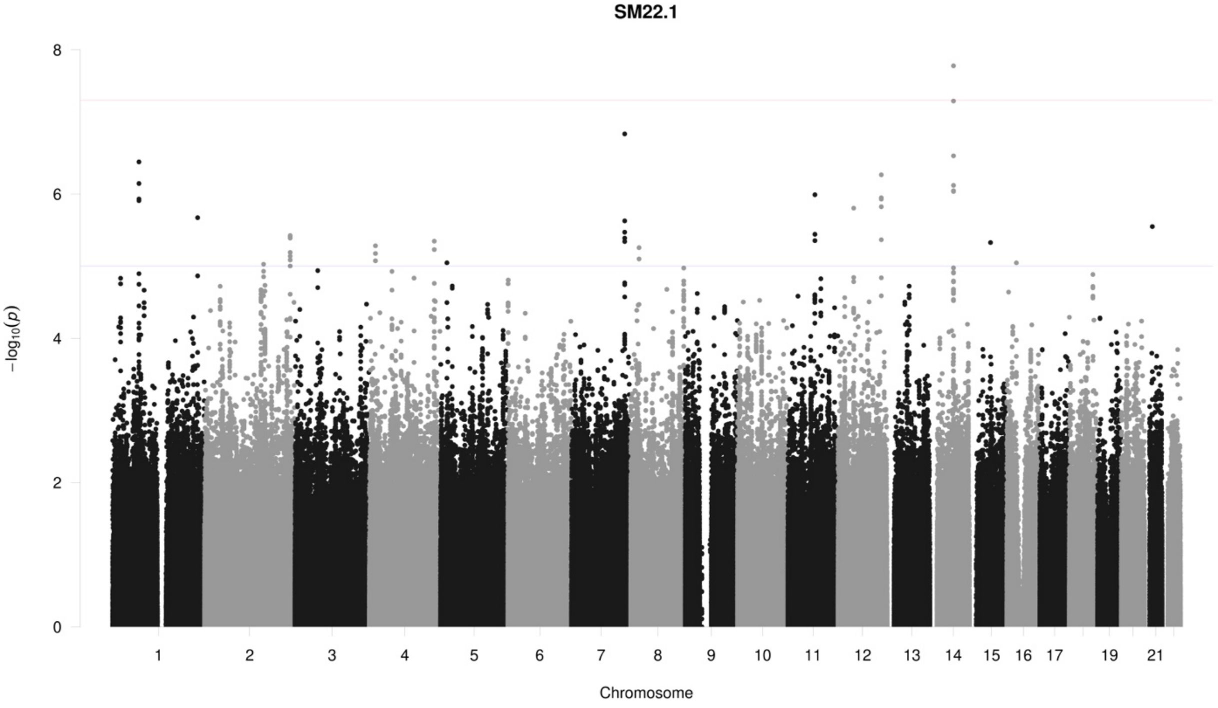
SM22.1
−log10(p)
Chromosome
1 2 3 4 5 6 7 8 9 10 11 12 13 14 15 16 17 19 21

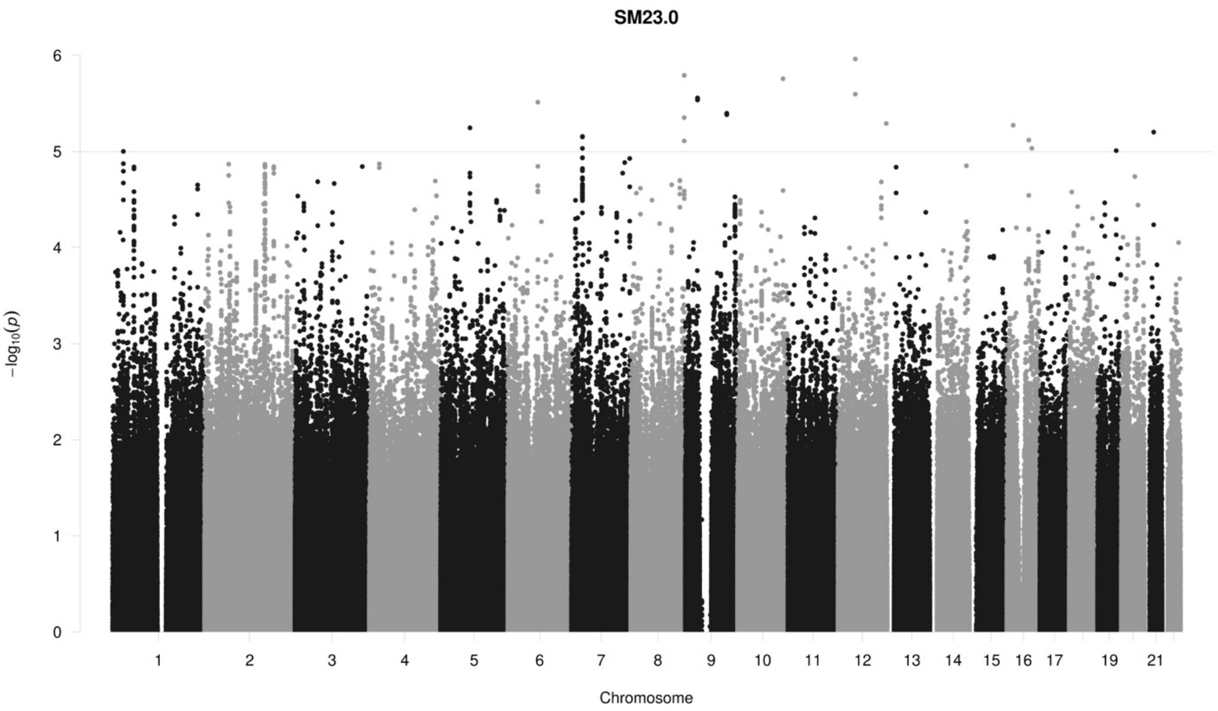
SM23.0
−log10(p)
0
1
2
3
4
5
6
19
21
Chromosome

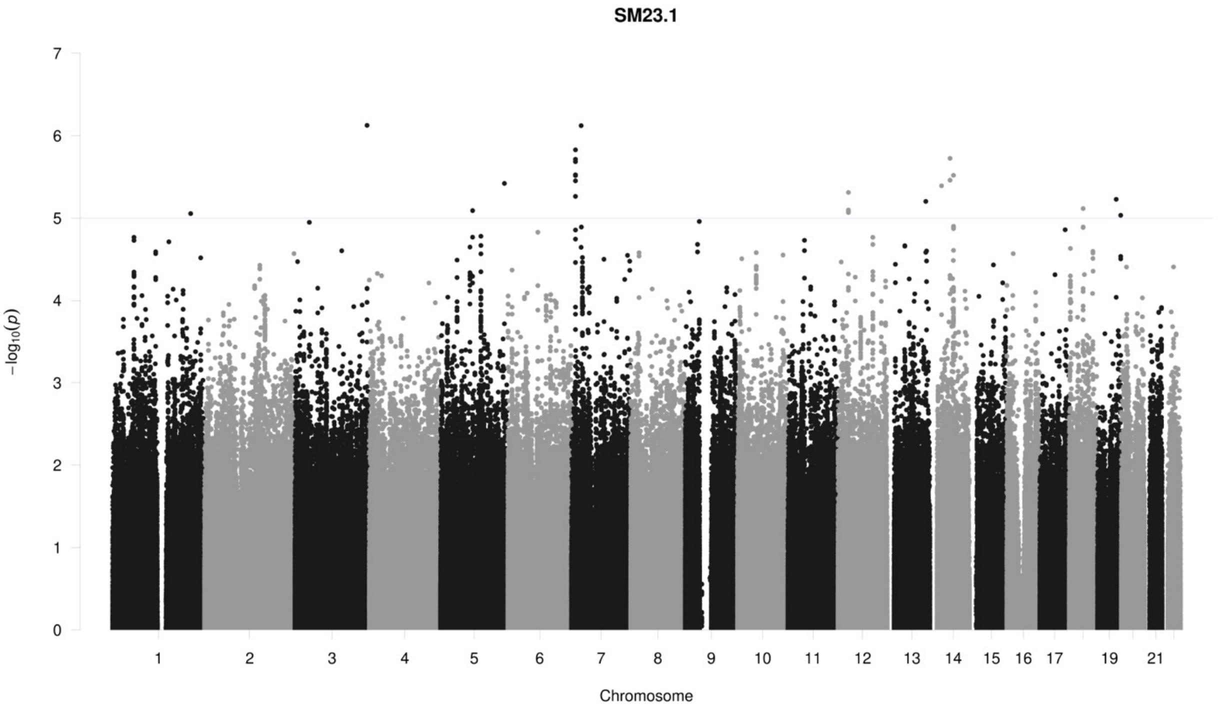
SM23.1
−log10(p)
0
1
2
3
4
5
6
7
19
21
Chromosome

## SM24.0

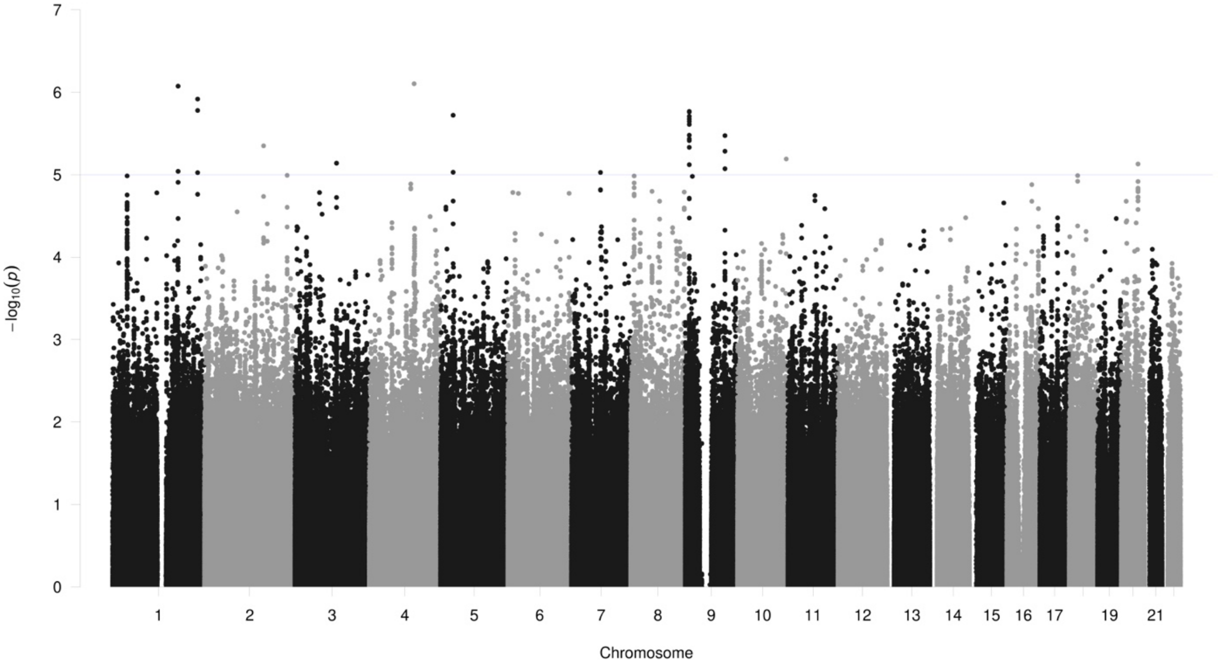

7
6
5
4
3
2
1
0
$-\log_{10}(p)$
19
21
Chromosome

## SM24.1

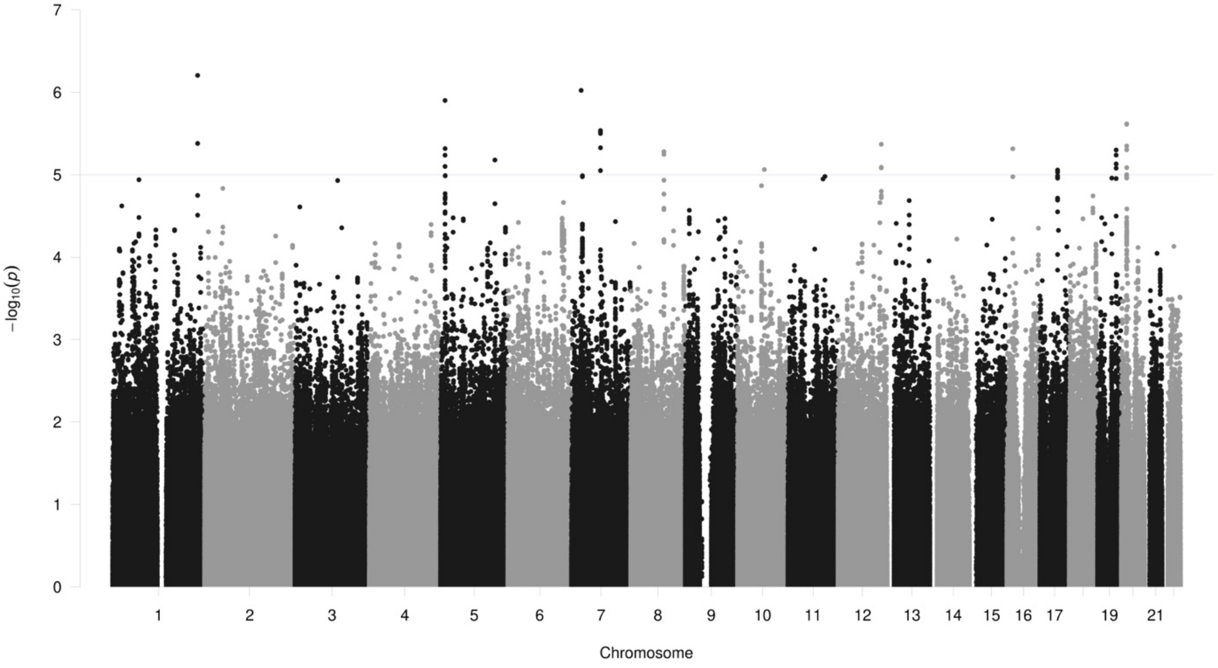

7
6
5
4
3
2
1
0
$-\log_{10}(p)$
19
21
Chromosome

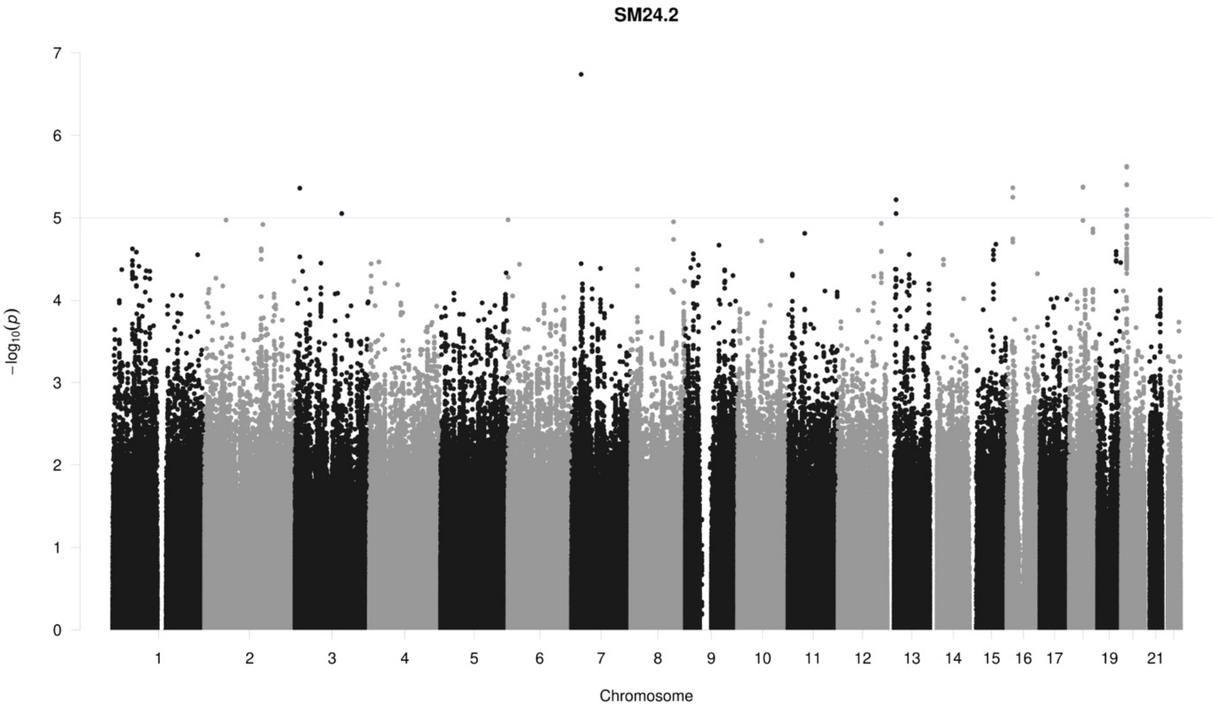
SM24.2
−log10(p)
Chromosome
0
1
2
3
4
5
6
7
19
21

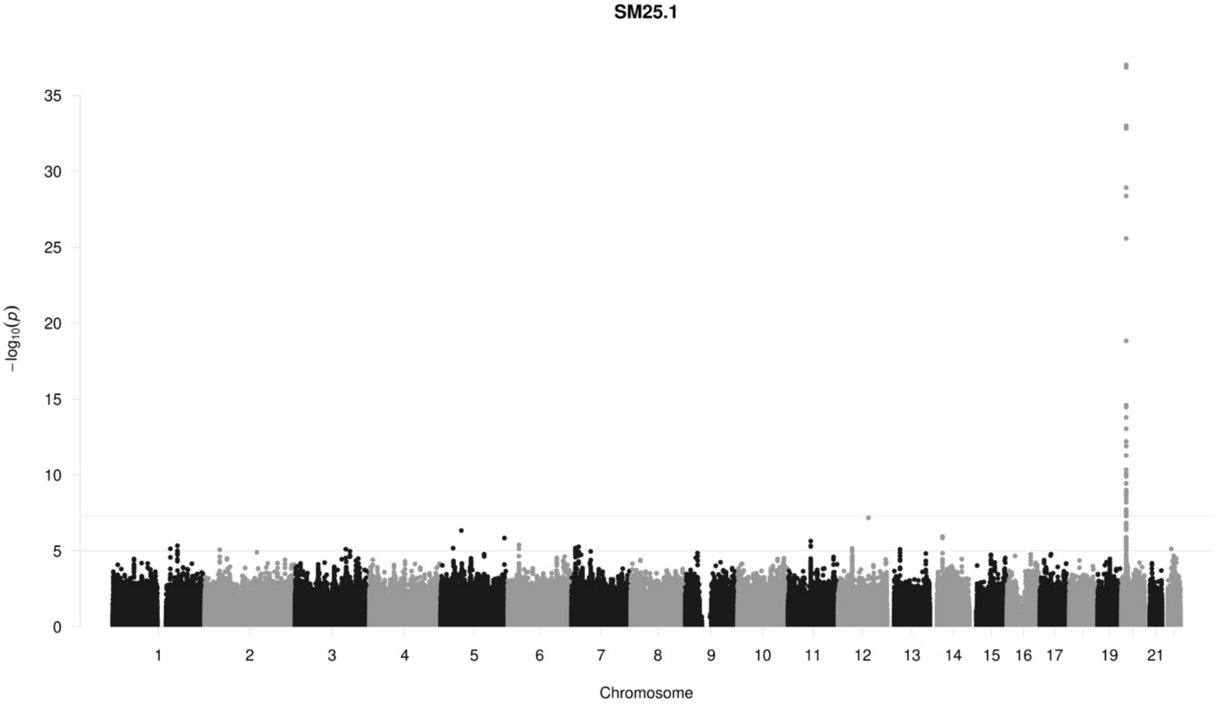
SM25.1
−log10(p)
Chromosome
0
5
10
15
20
25
30
35
19
21

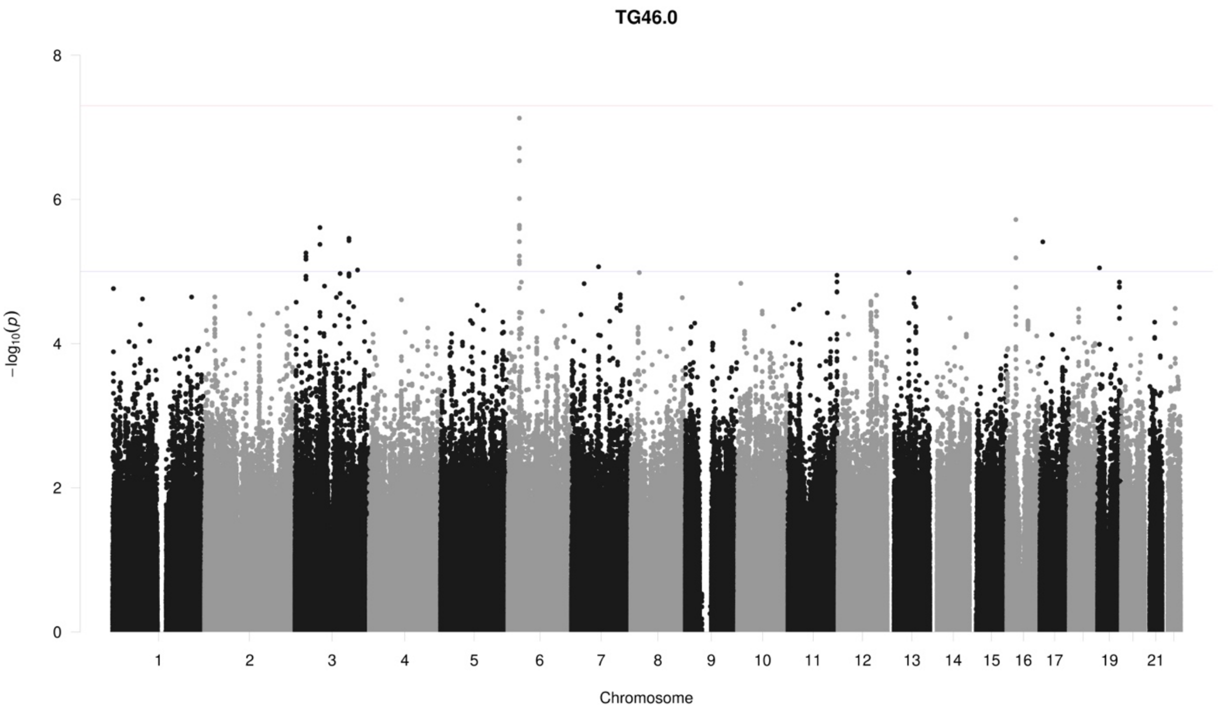
TG46.0
8
6
4
2
0
−log10(p)
19
21
Chromosome

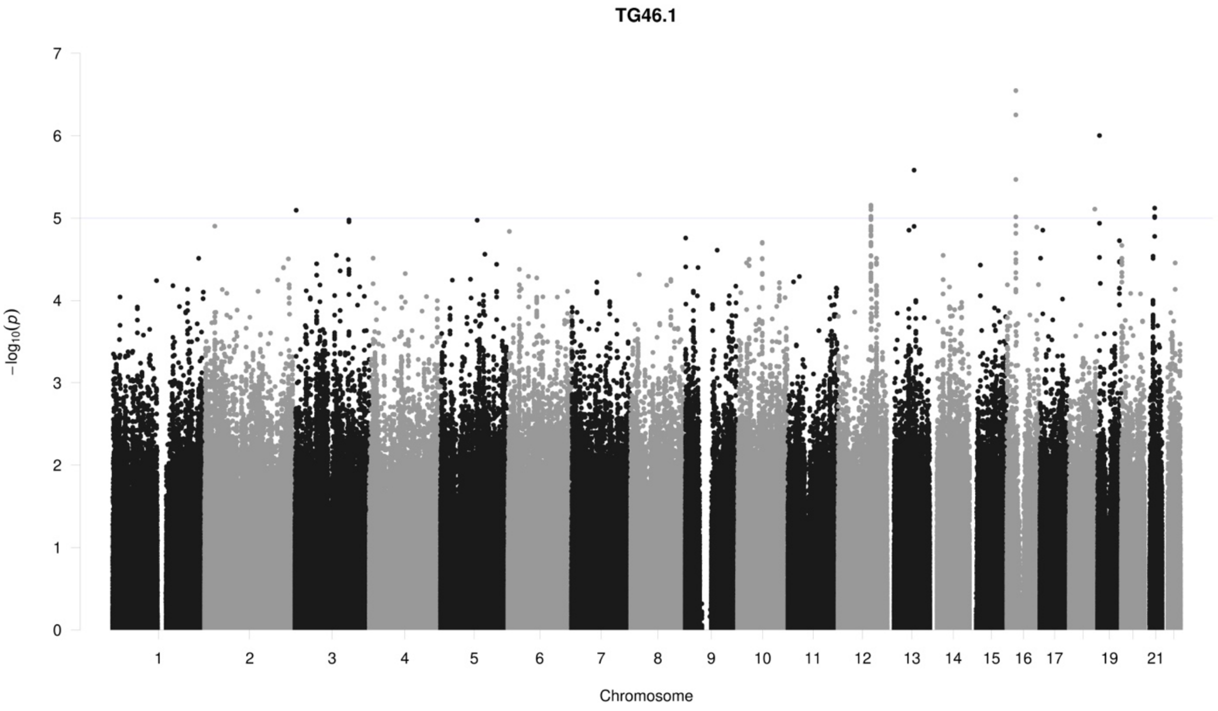
TG46.1
7
6
5
4
3
2
1
0
−log10(p)
19
21
Chromosome

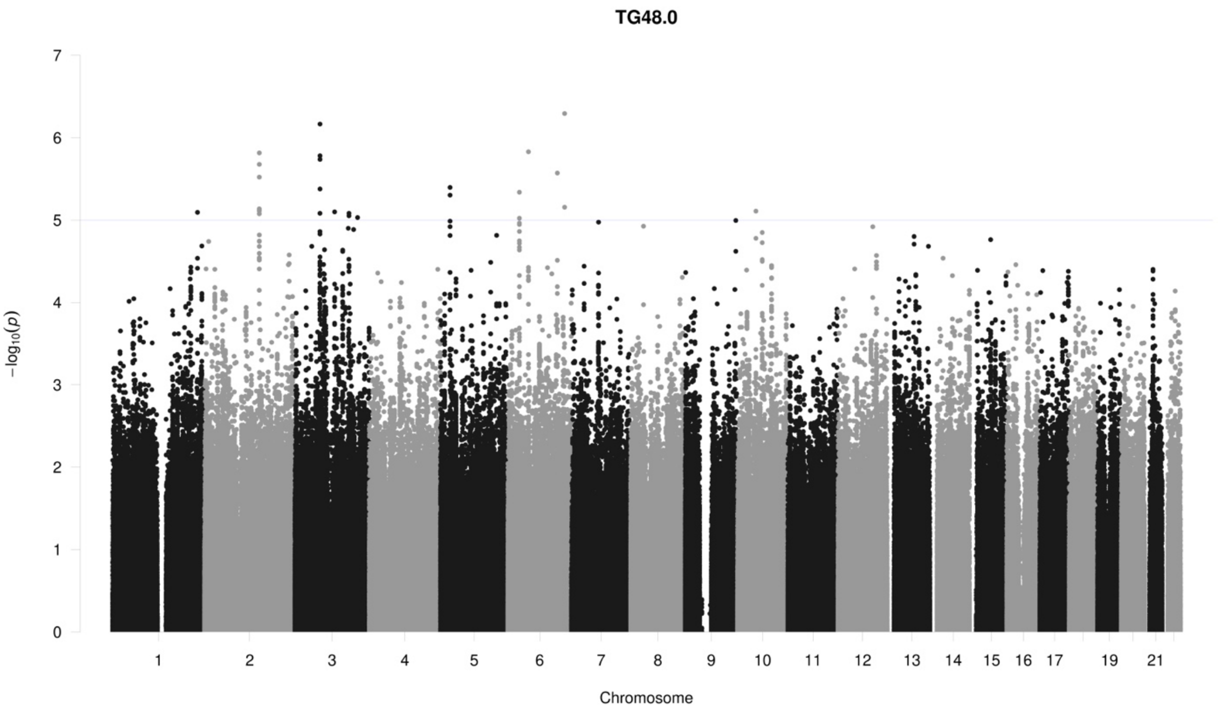
TG48.0
−log10(p)
0
1
2
3
4
5
6
7
1 2 3 4 5 6 7 8 9 10 11 12 13 14 15 16 17 19 21
Chromosome

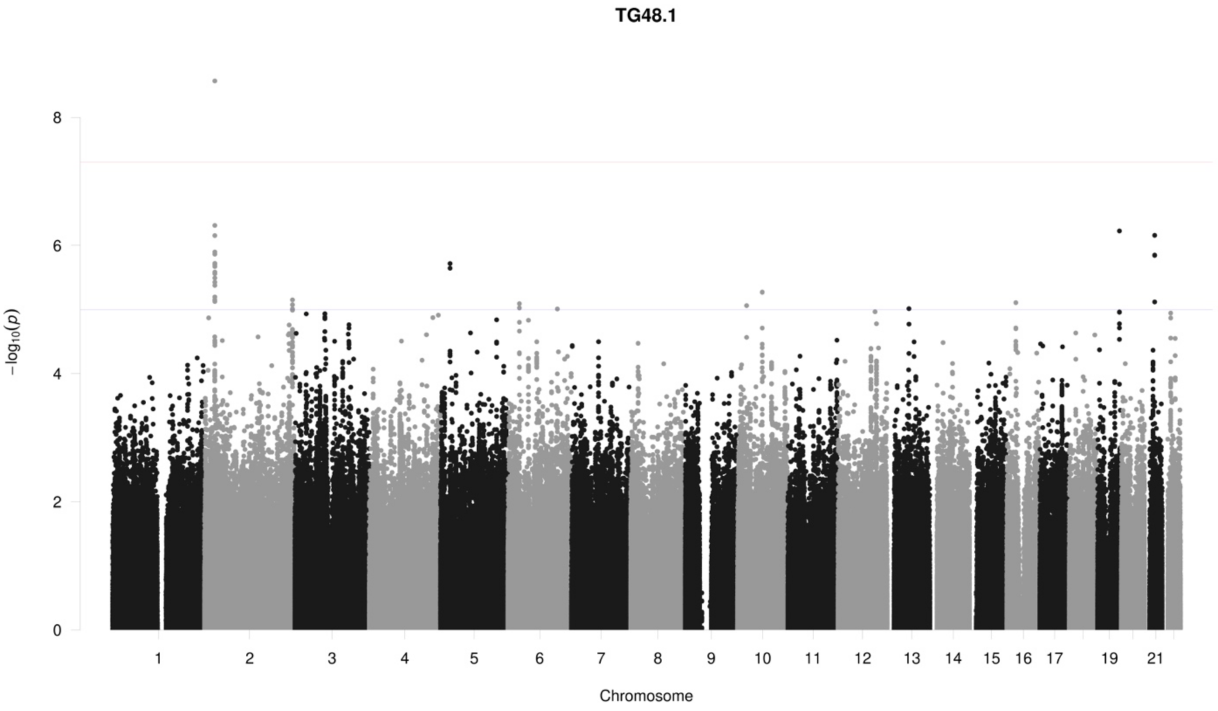
TG48.1
−log10(p)
0
2
4
6
8
1 2 3 4 5 6 7 8 9 10 11 12 13 14 15 16 17 19 21
Chromosome

## TG48.2

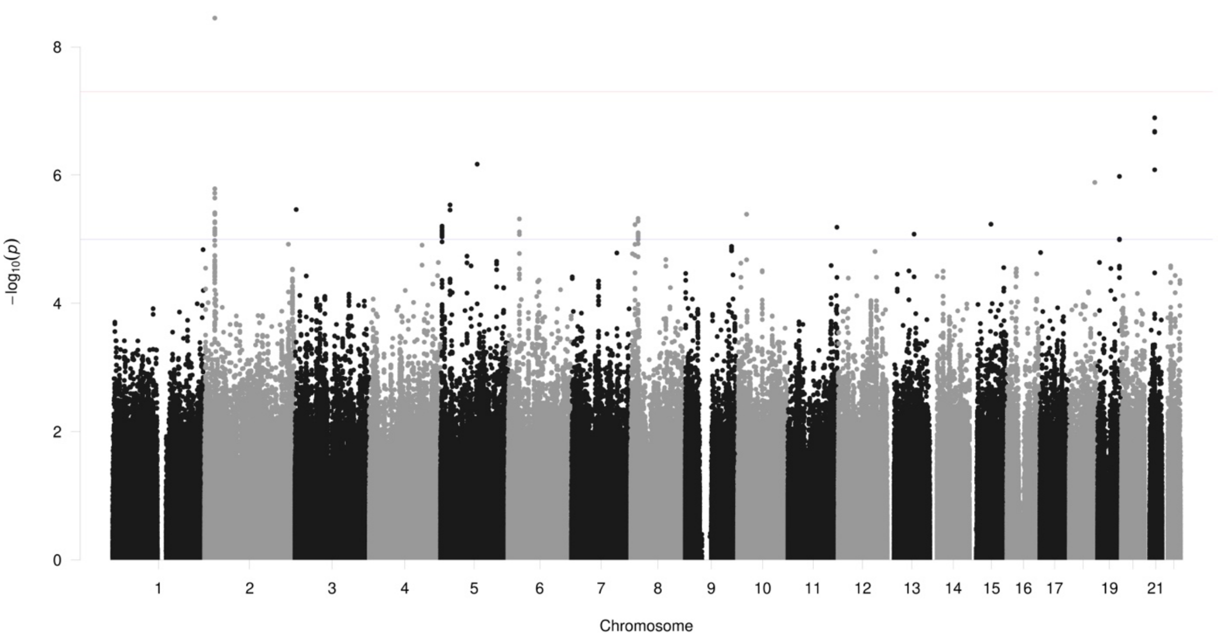
0
2
4
6
8
−log10(p)
19
21
Chromosome

## TG48.3

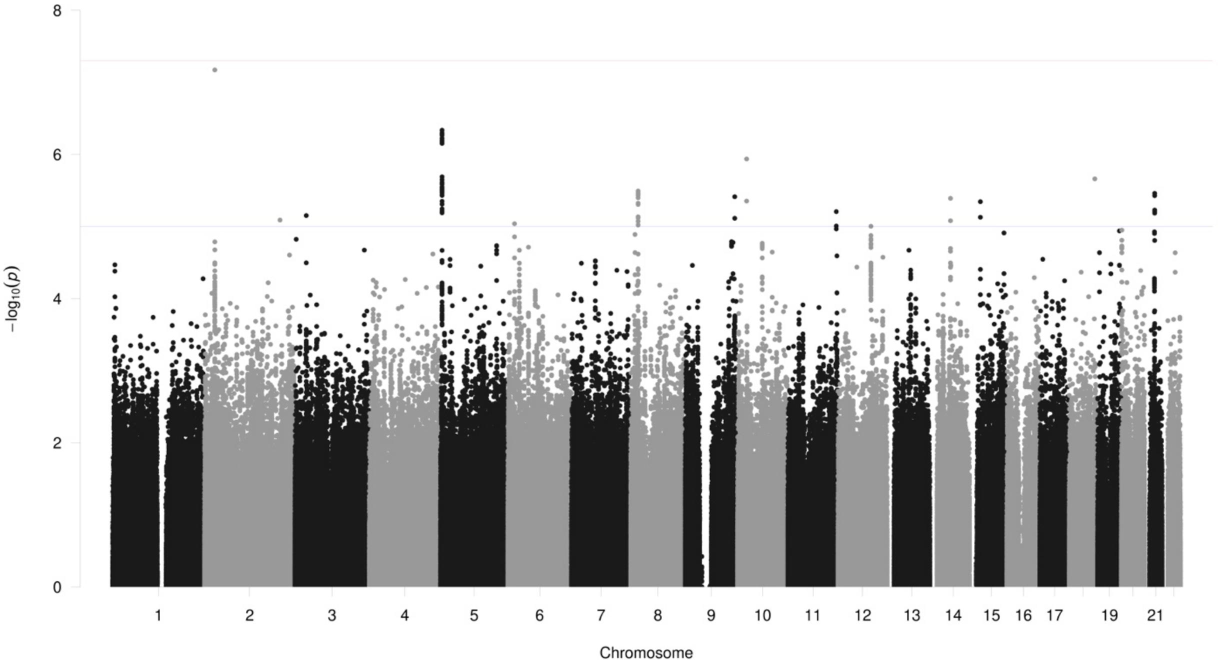
0
2
4
6
8
−log10(p)
19
21
Chromosome

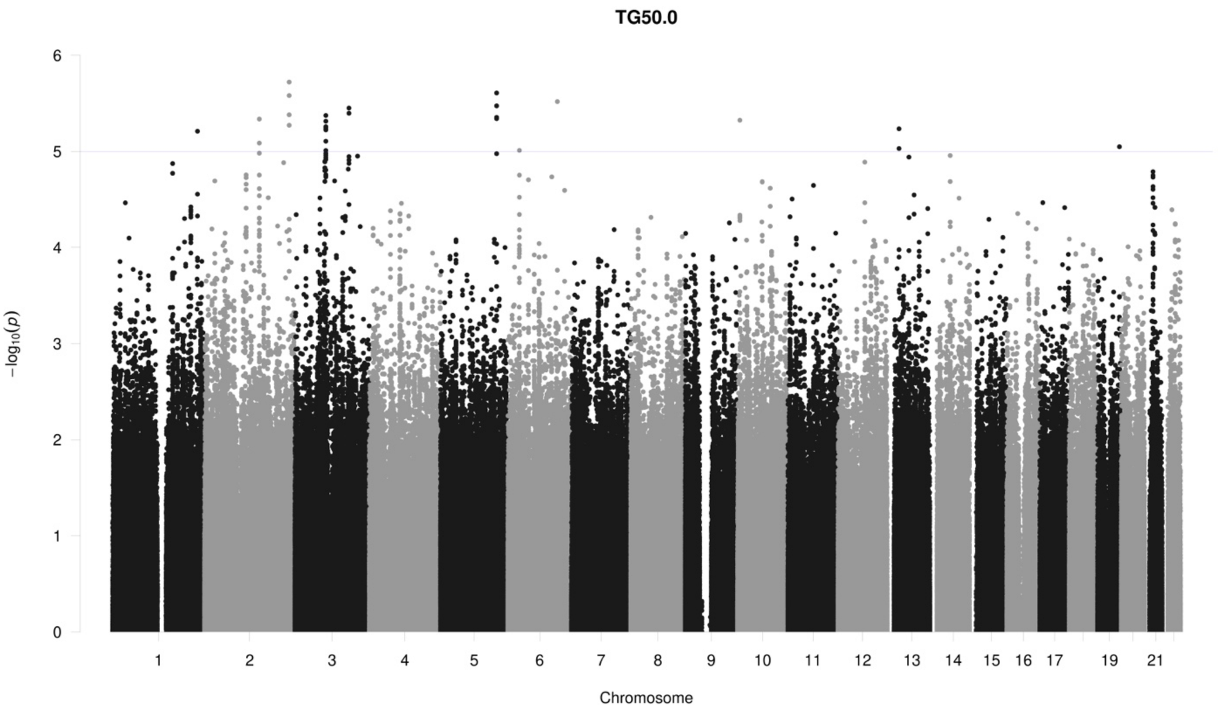
TG50.0
−log10(p)
0
1
2
3
4
5
6
19
21
Chromosome

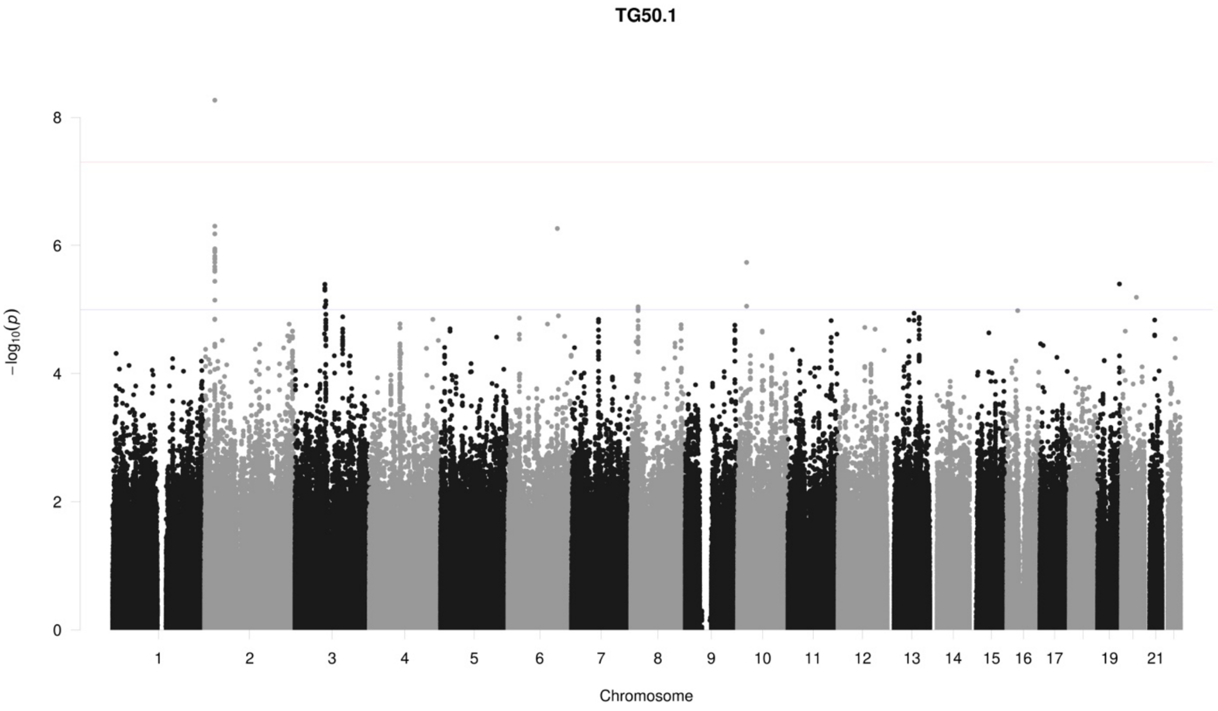
TG50.1
−log10(p)
0
2
4
6
8
19
21
Chromosome

**TG50.2**

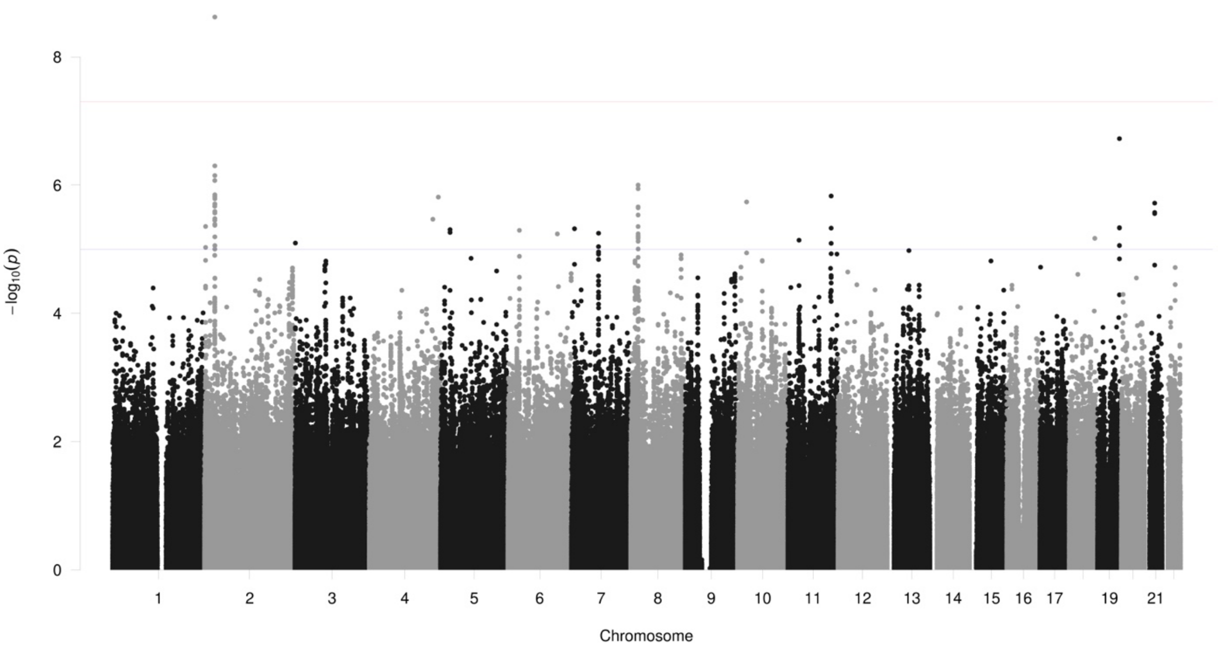

8
6
4
2
0
−log10(p)
19
21
Chromosome

**TG50.3**

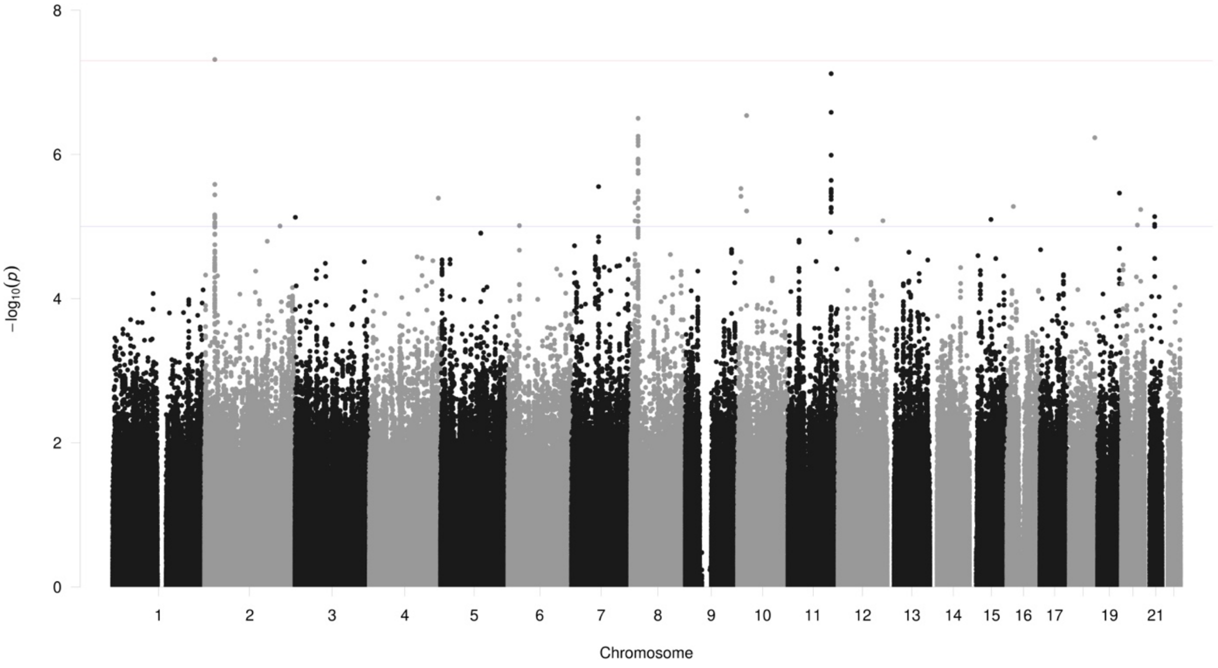

8
6
4
2
0
−log10(p)
19
21
Chromosome

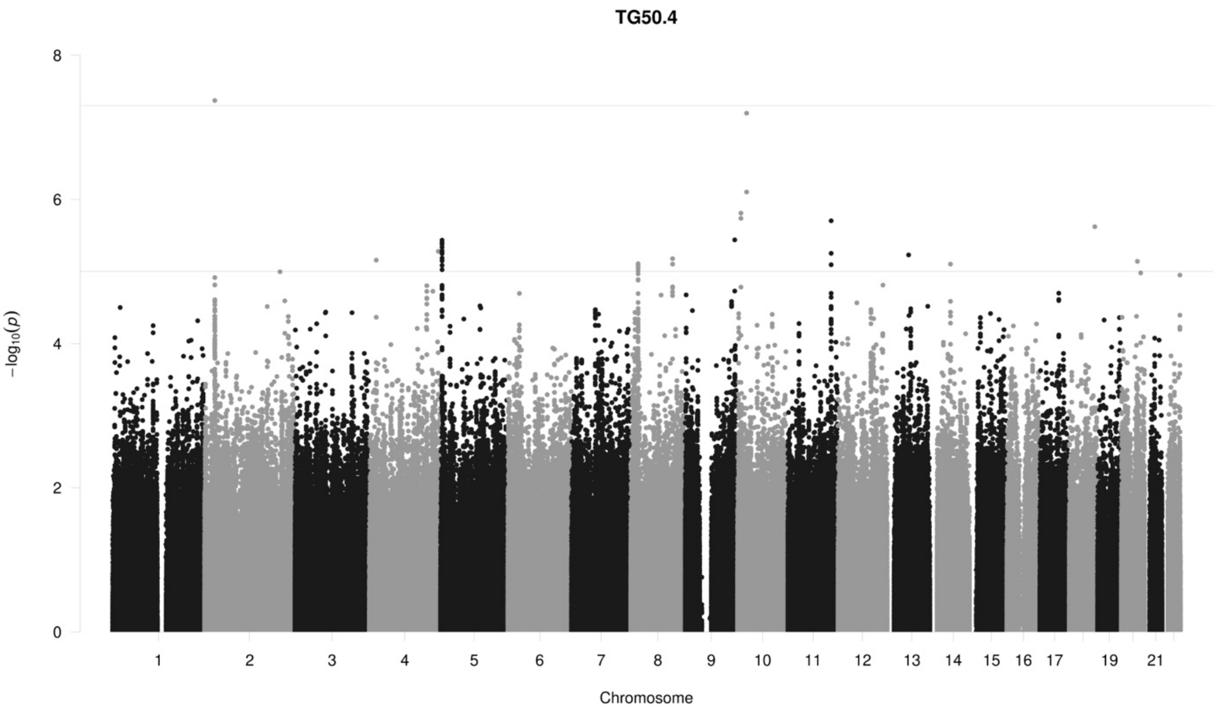
TG50.4
−log10(p)
0
2
4
6
8
19
21
Chromosome

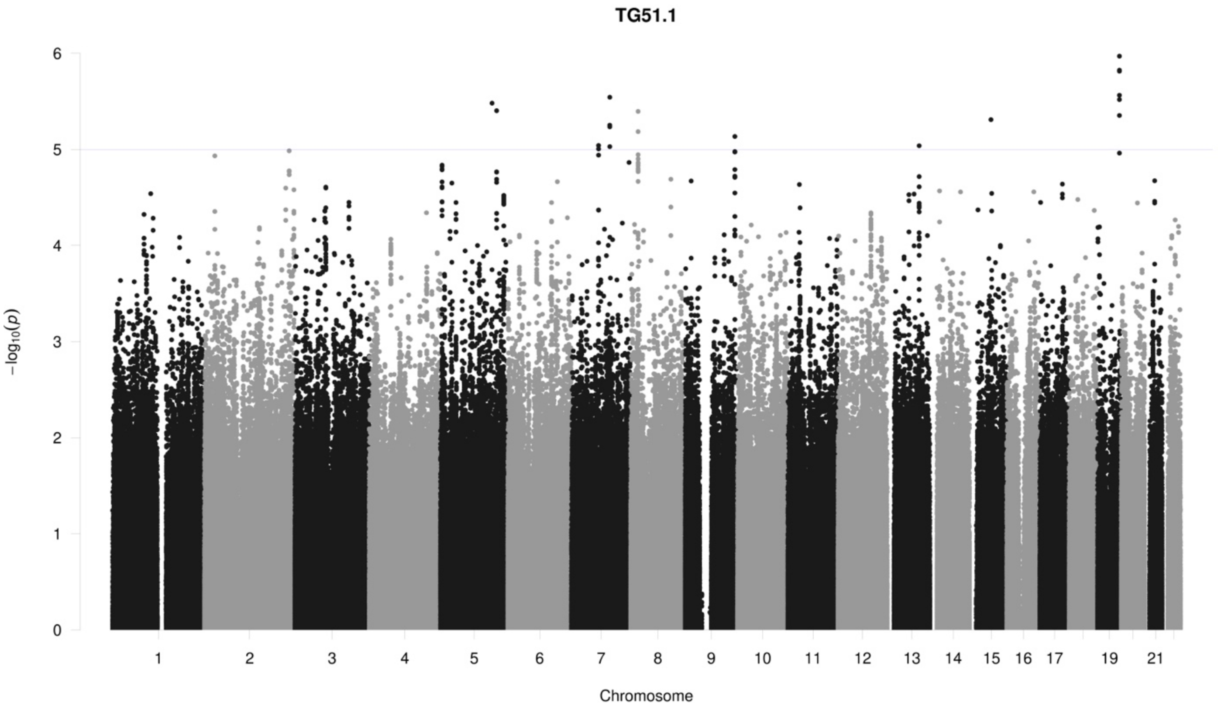
TG51.1
−log10(p)
0
1
2
3
4
5
6
19
21
Chromosome

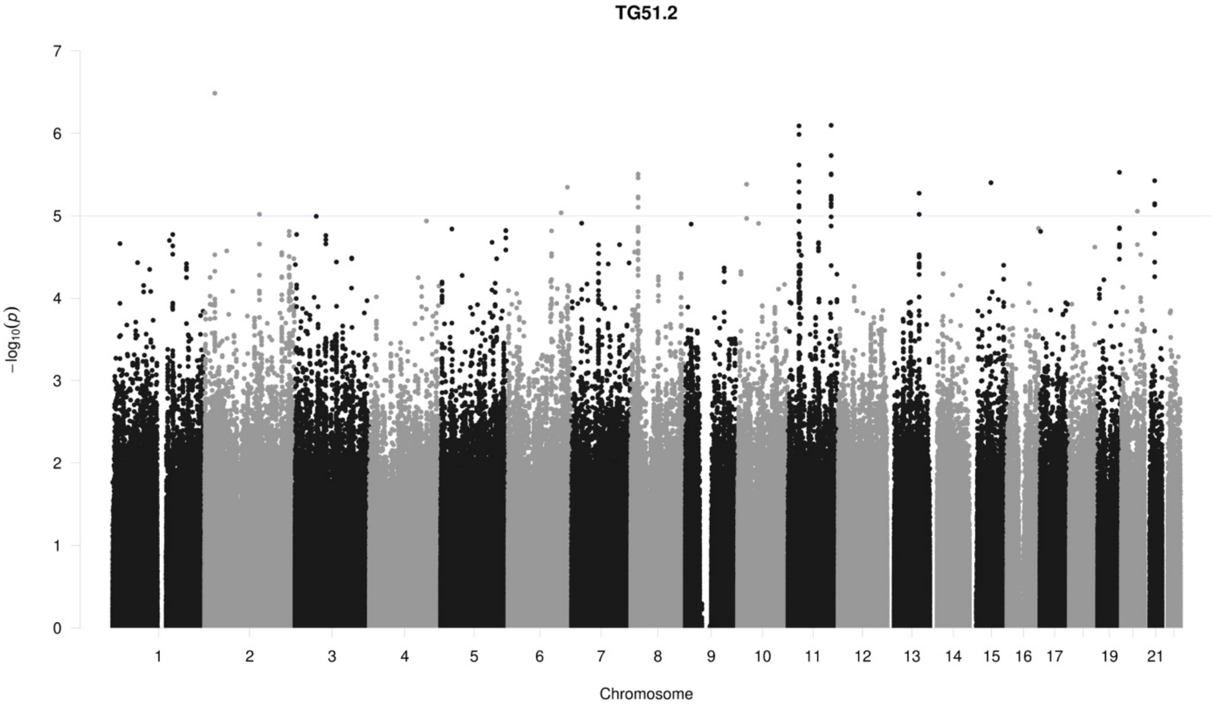
TG51.2
0
1
2
3
4
5
6
7
−log10(p)
19
21
Chromosome

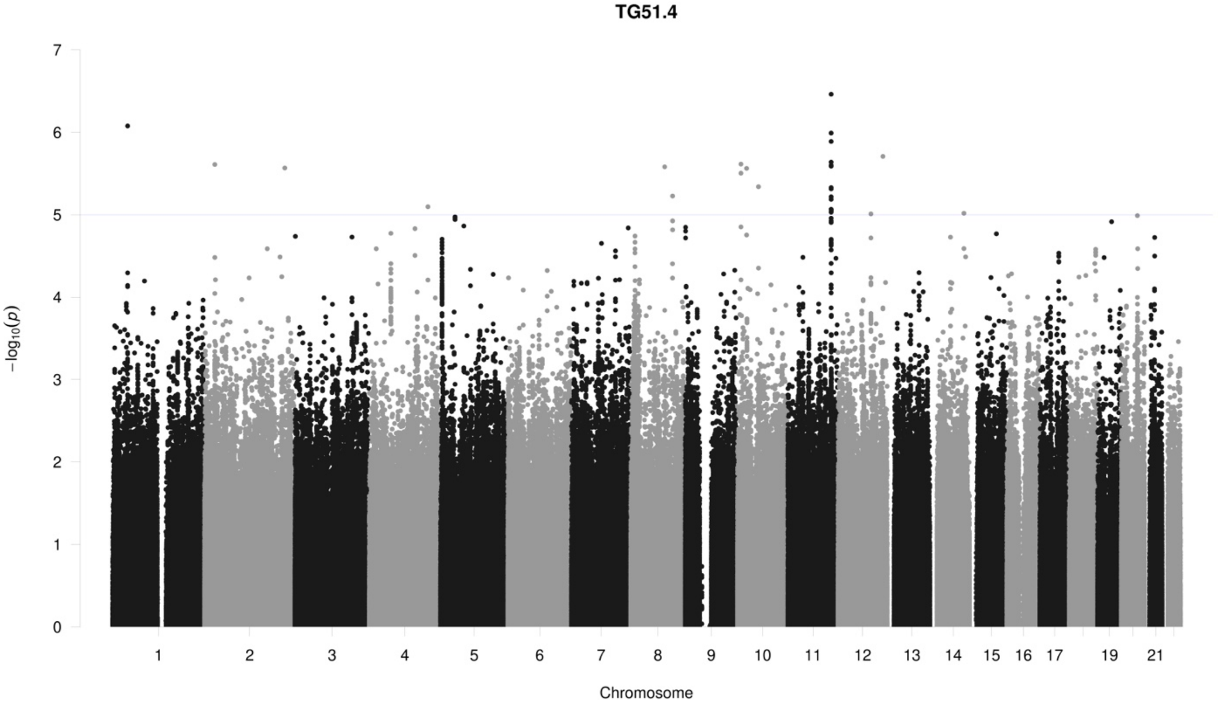
TG51.4
0
1
2
3
4
5
6
7
−log10(p)
19
21
Chromosome

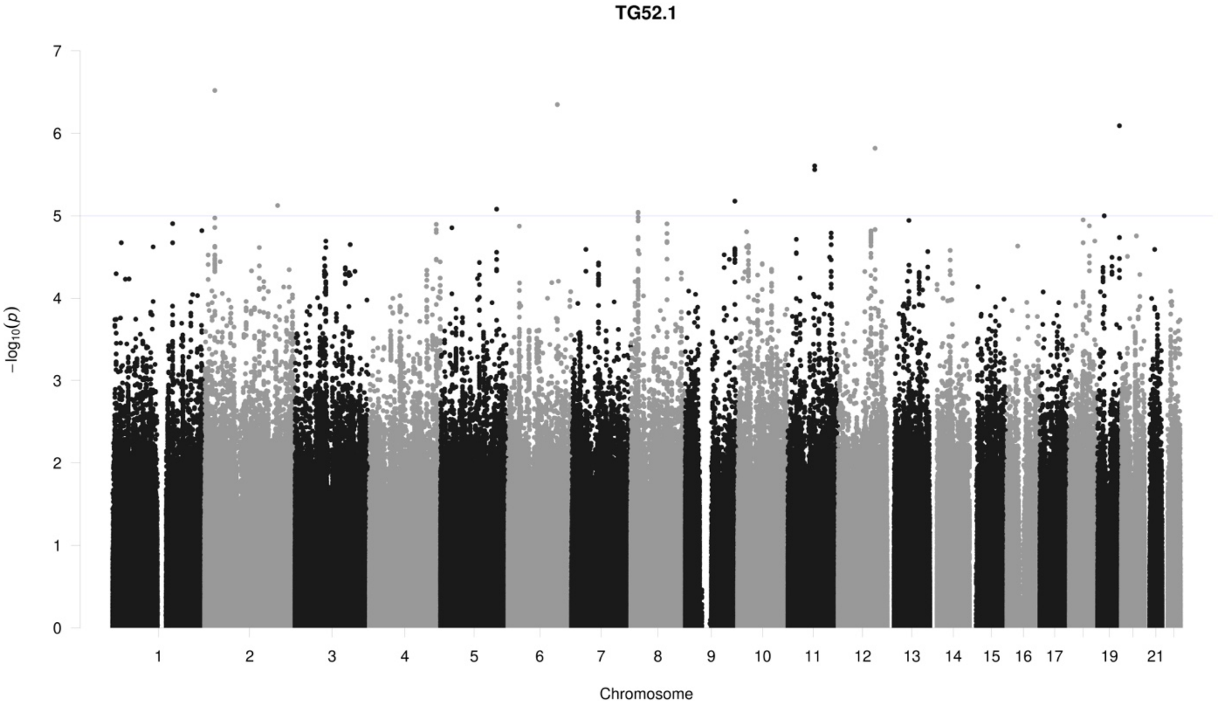
TG52.1
7
6
5
4
3
2
1
0
−log10(p)
19
21
Chromosome

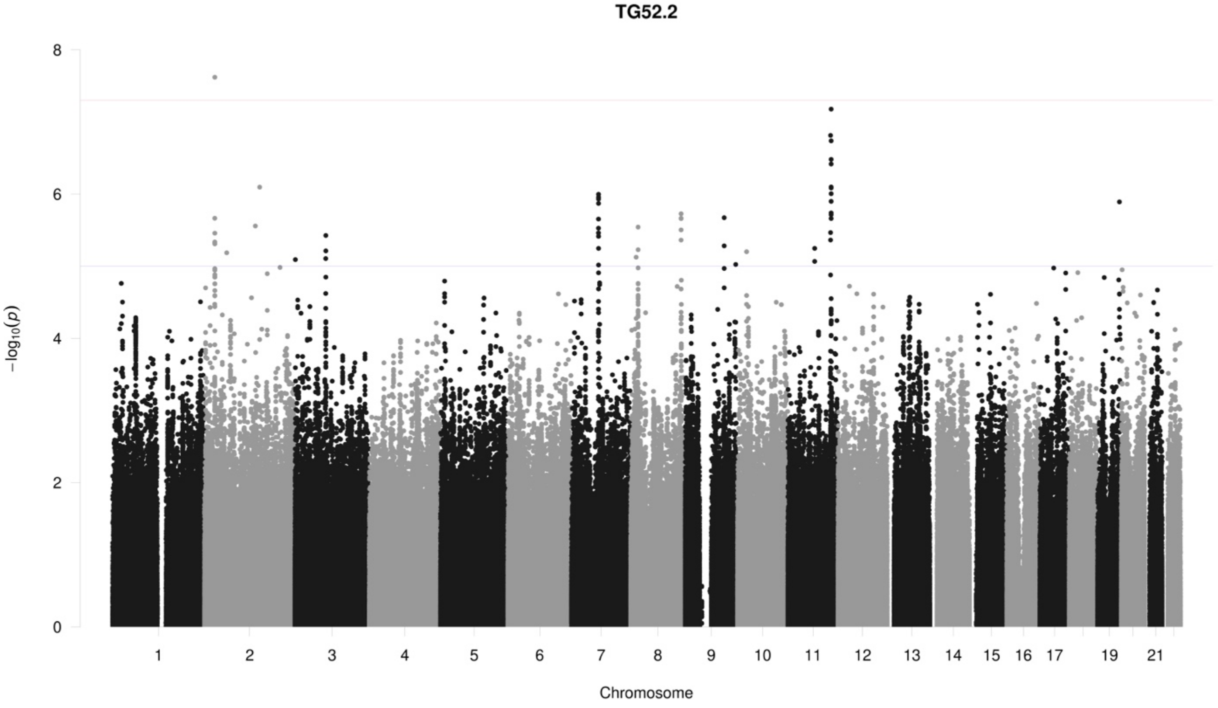
TG52.2
8
6
4
2
0
−log10(p)
19
21
Chromosome

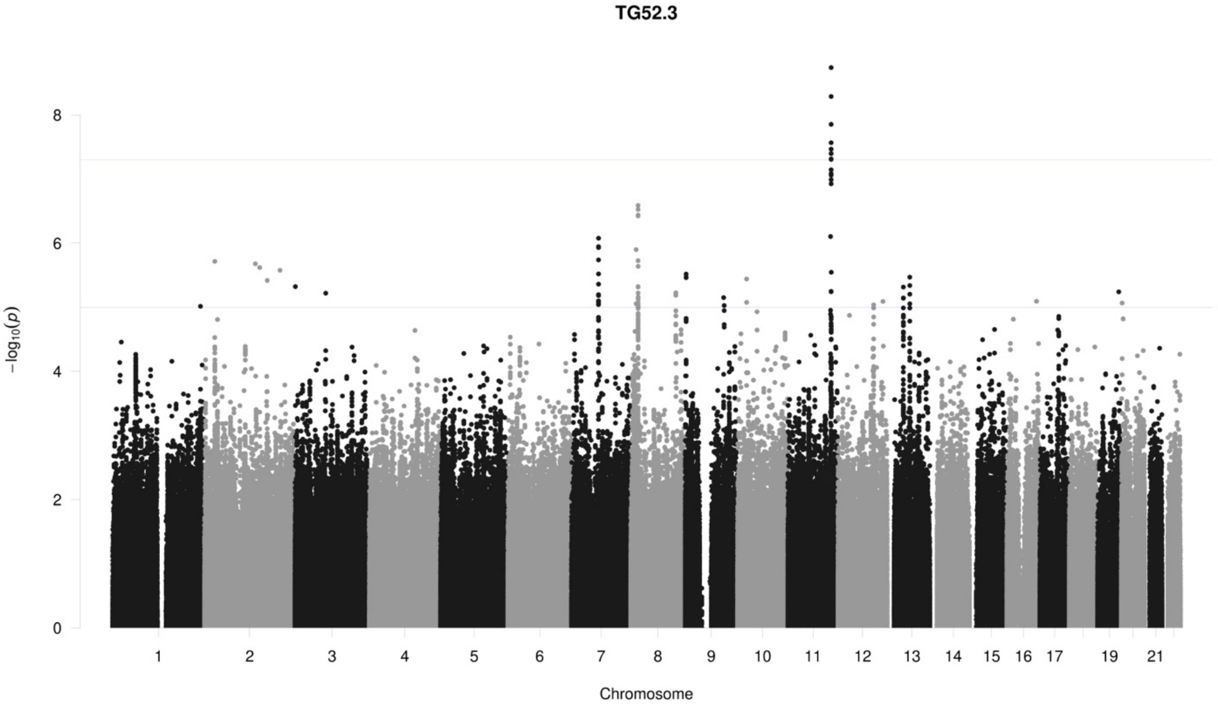
TG52.3
8
6
4
2
0
−log10(p)
19
21
Chromosome

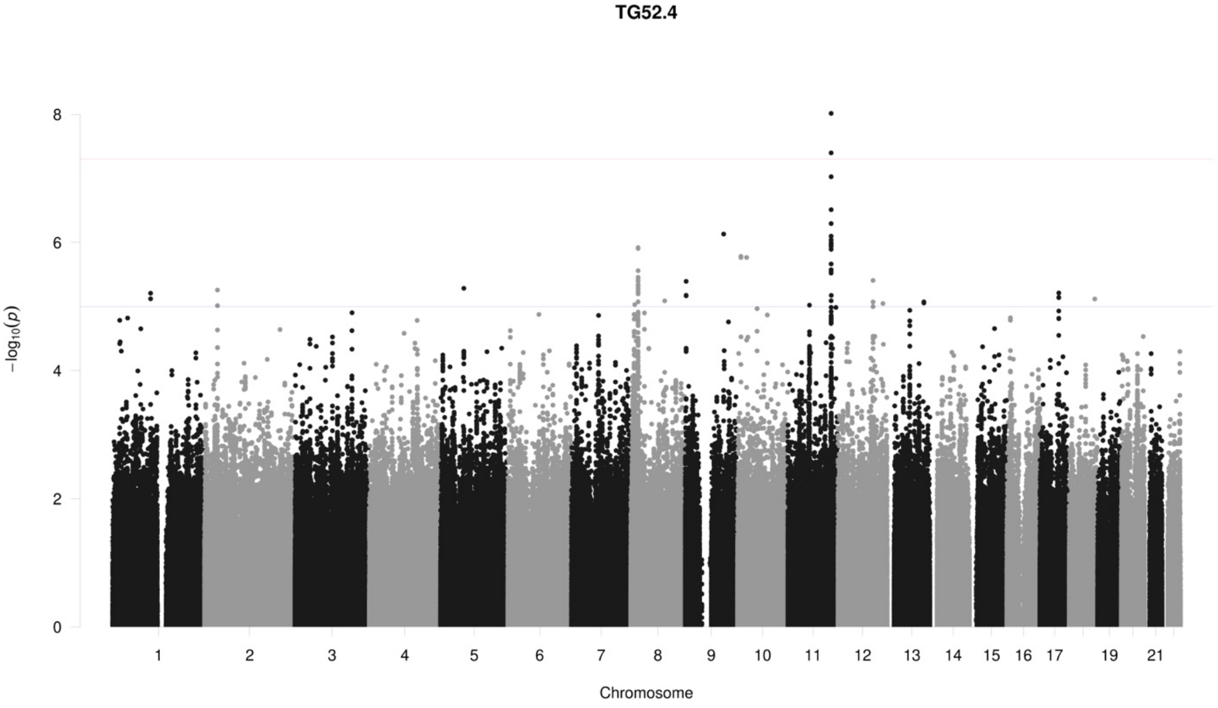
TG52.4
8
6
4
2
0
−log10(p)
19
21
Chromosome

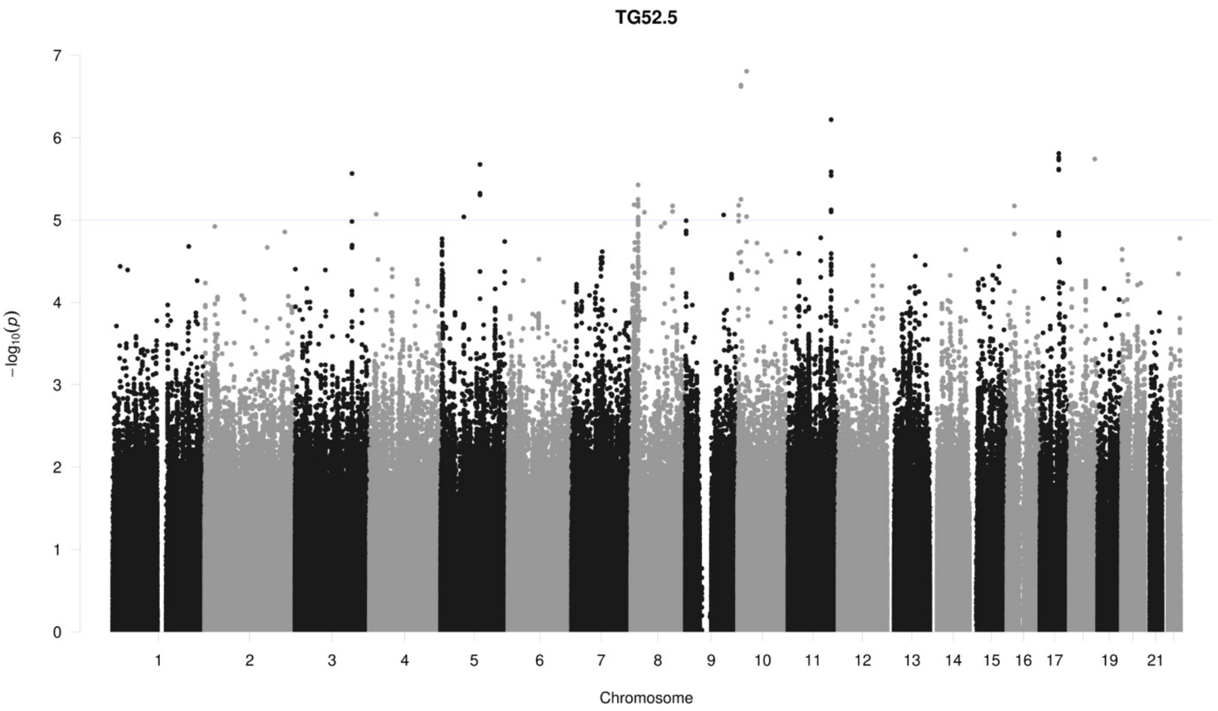

TG52.5
−log10(p)
Chromosome

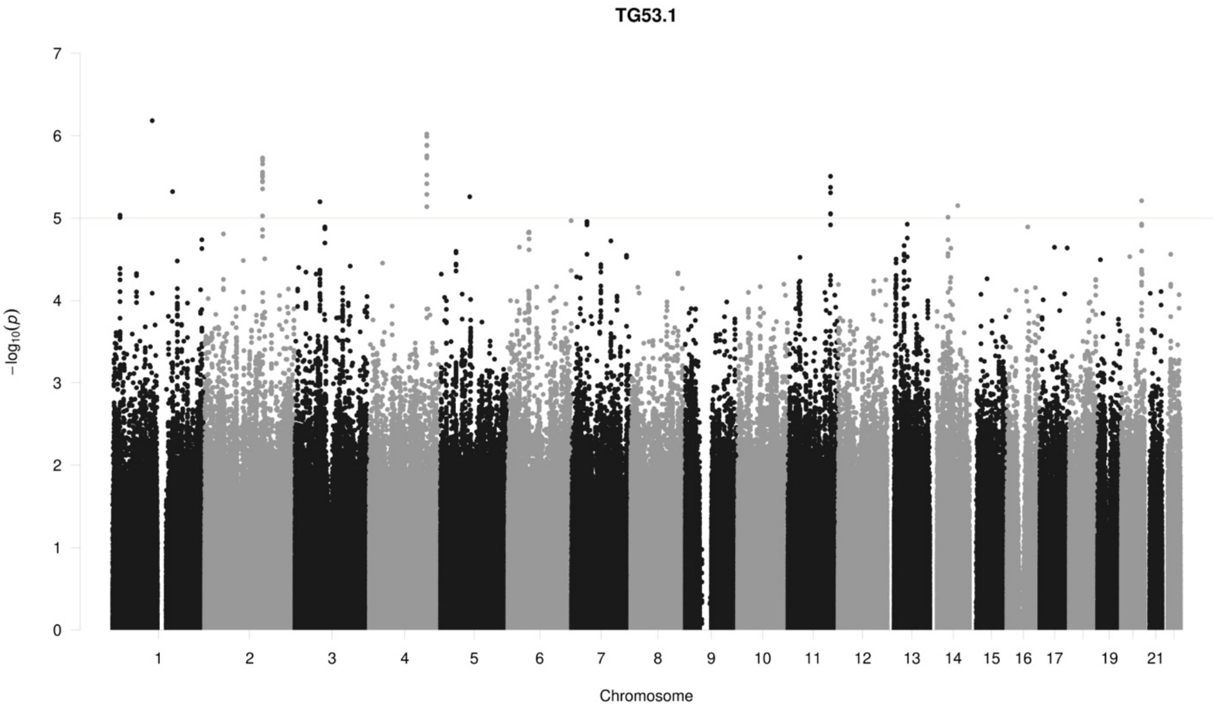

TG53.1
−log10(p)
Chromosome

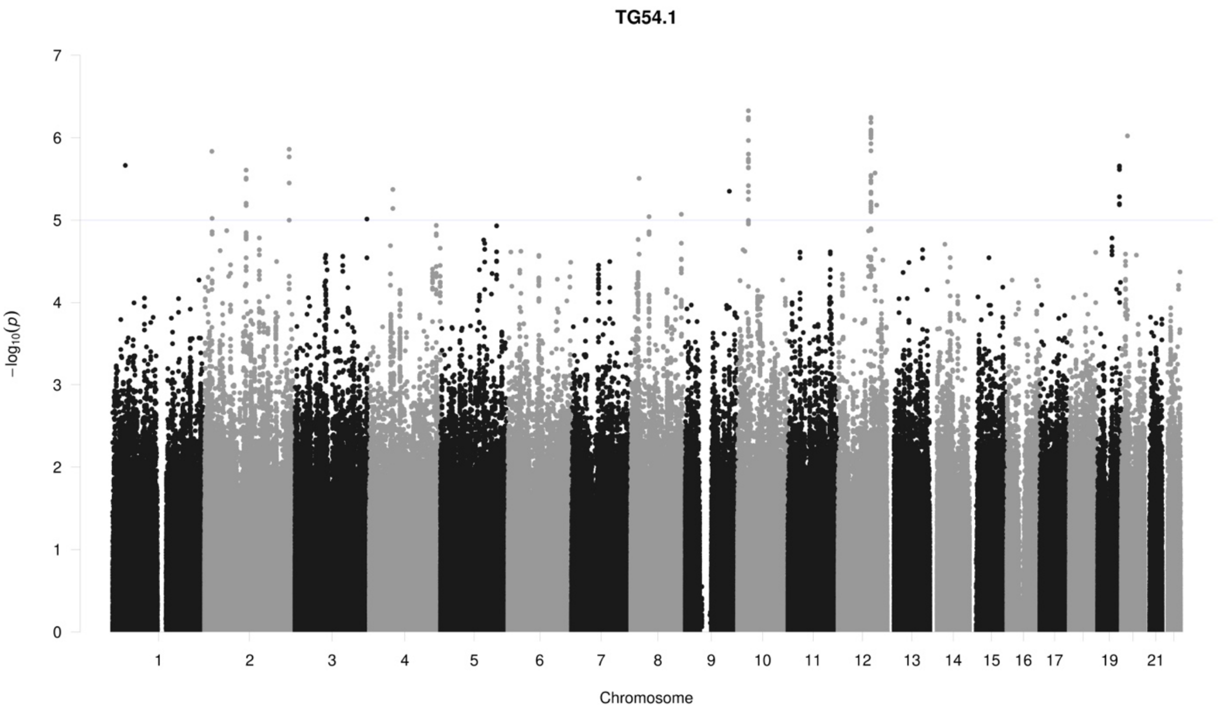
TG54.1
0
1
2
3
4
5
6
7
−log10(p)
19
21
Chromosome

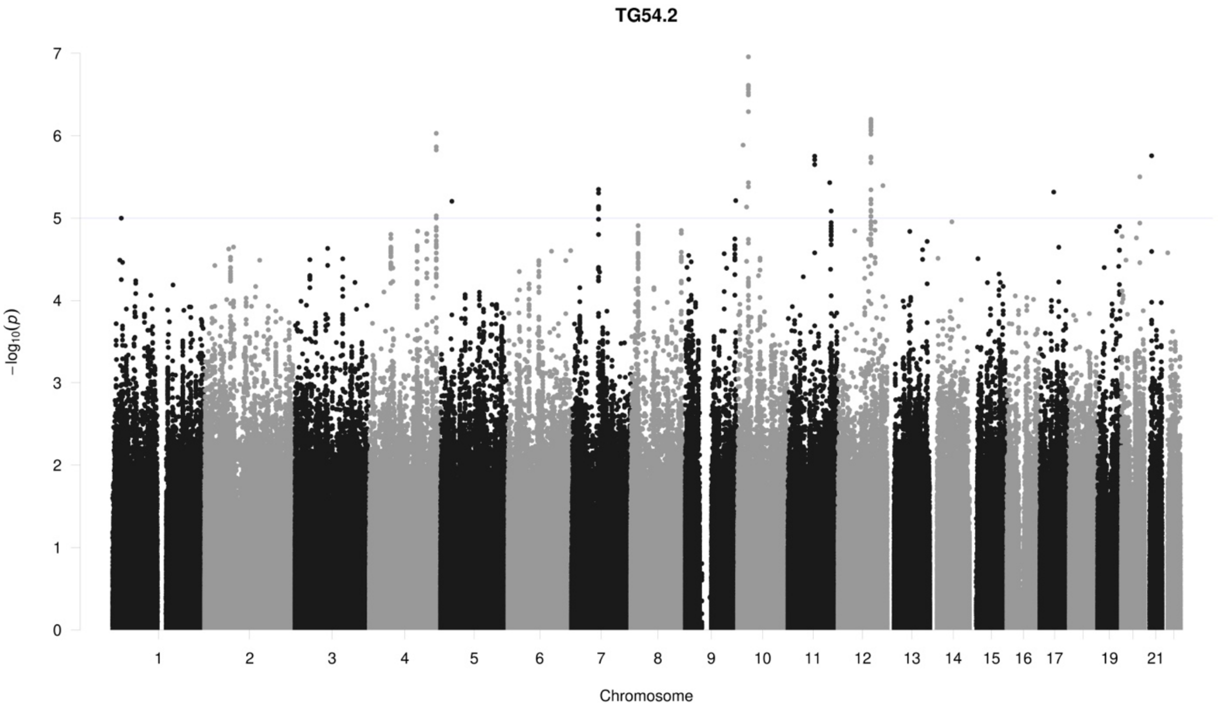
TG54.2
0
1
2
3
4
5
6
7
−log10(p)
19
21
Chromosome

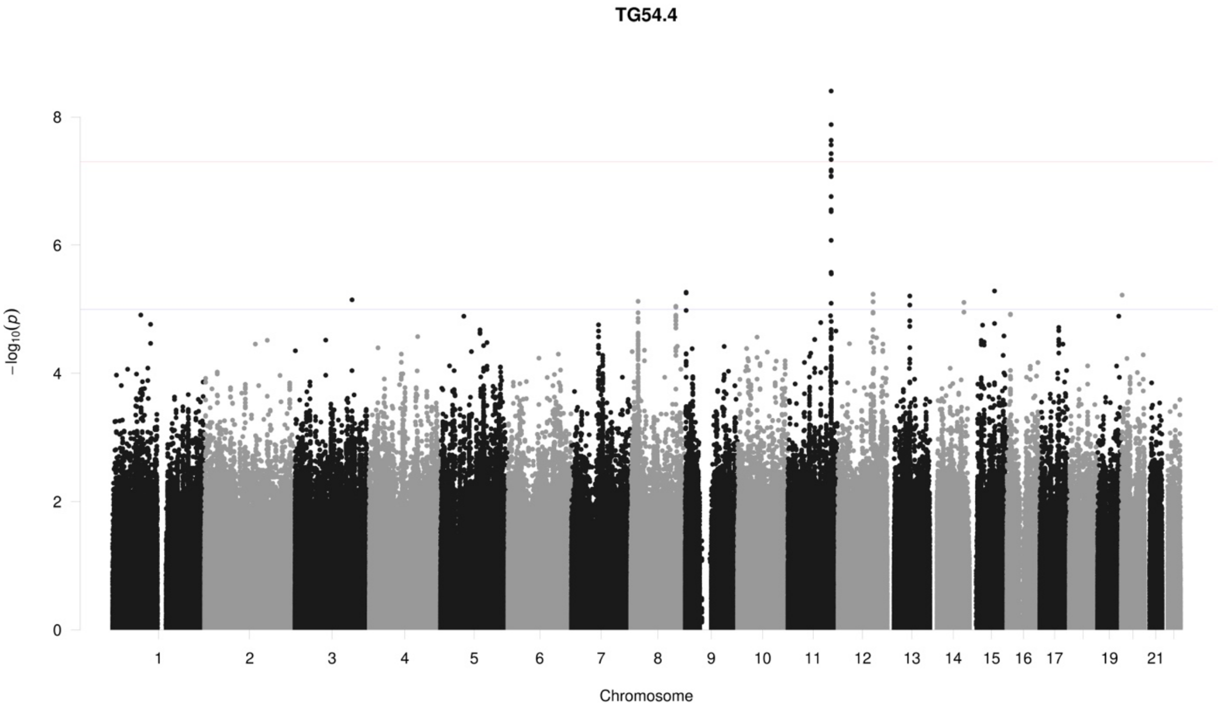
TG54.4
−log10(p)
8
6
4
2
19
21
Chromosome

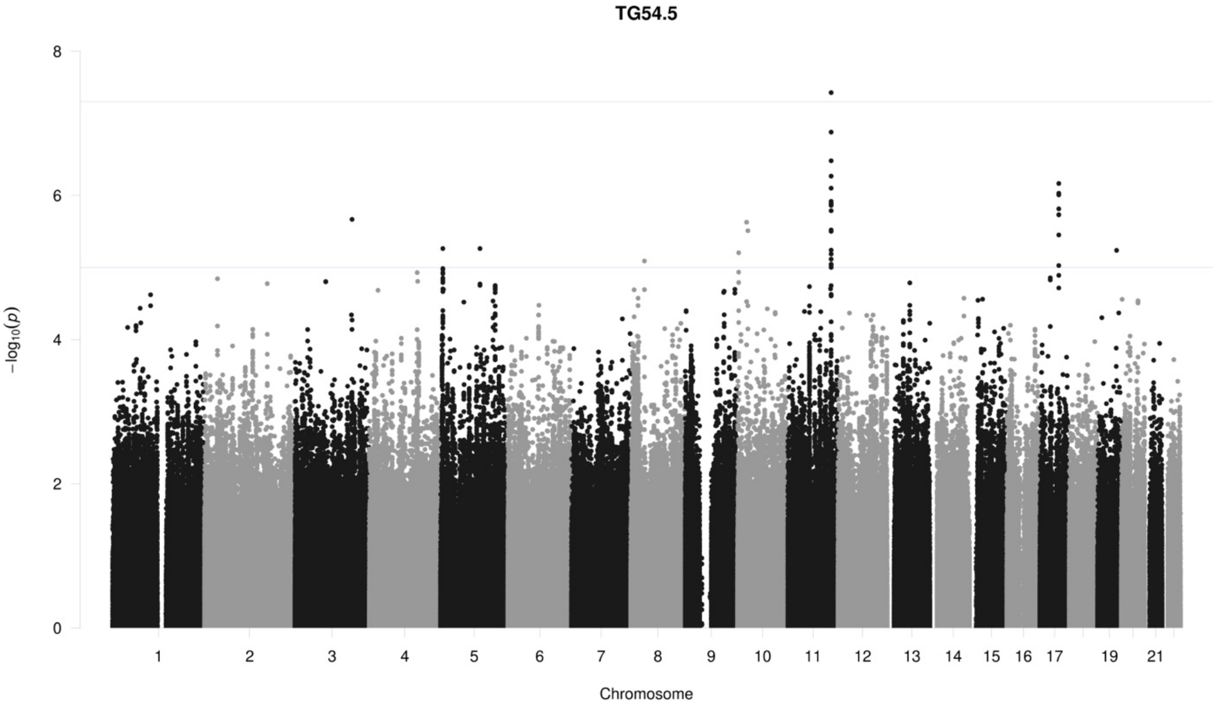
TG54.5
−log10(p)
8
6
4
2
19
21
Chromosome

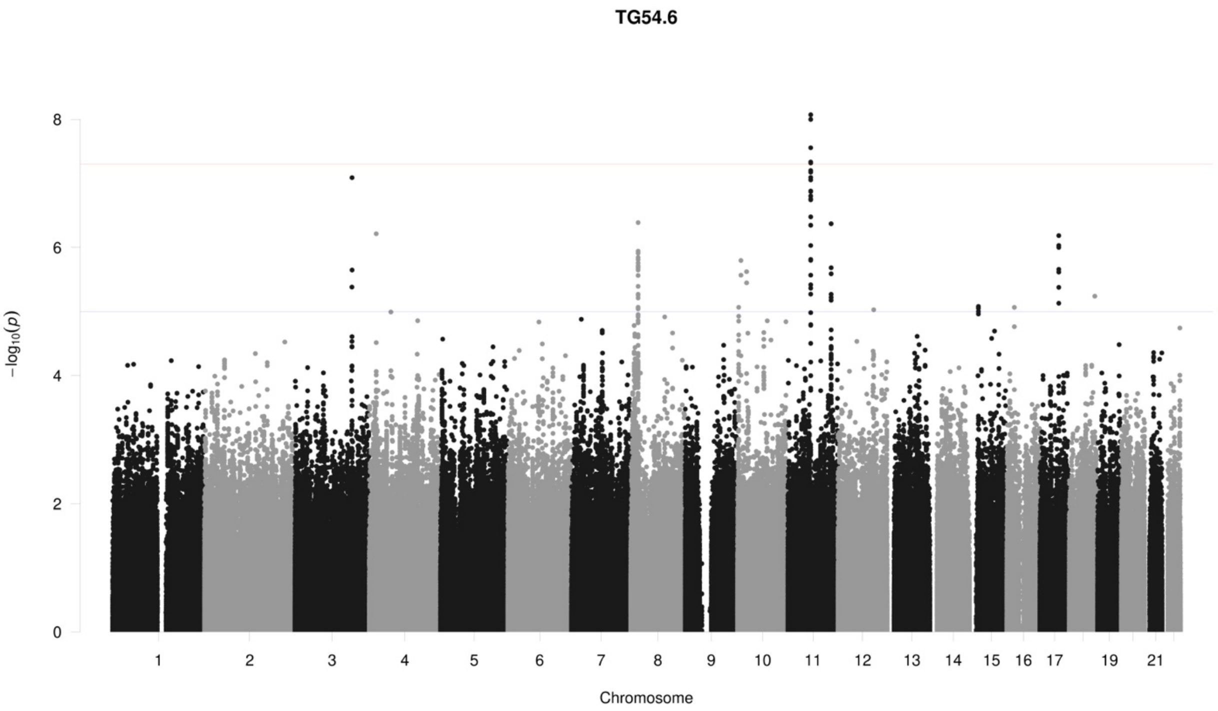
TG54.6
−log10(p)
Chromosome
0
2
4
6
8
19
21

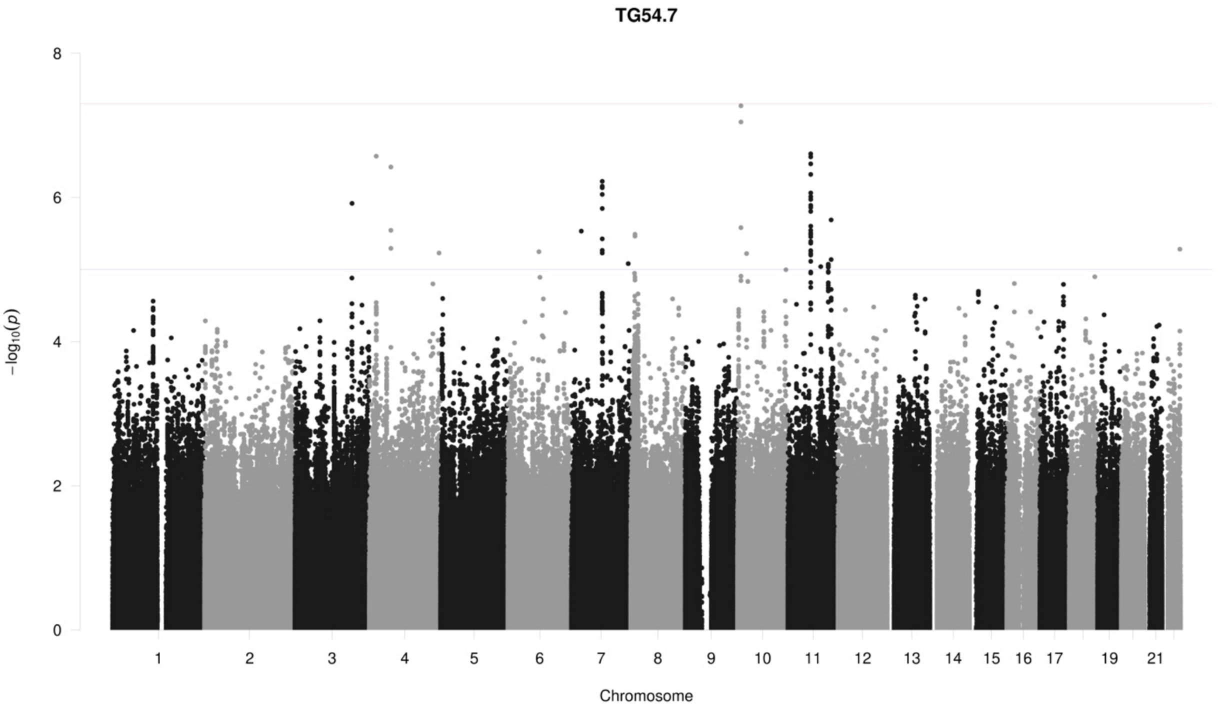
TG54.7
−log10(p)
Chromosome
0
2
4
6
8
19
21

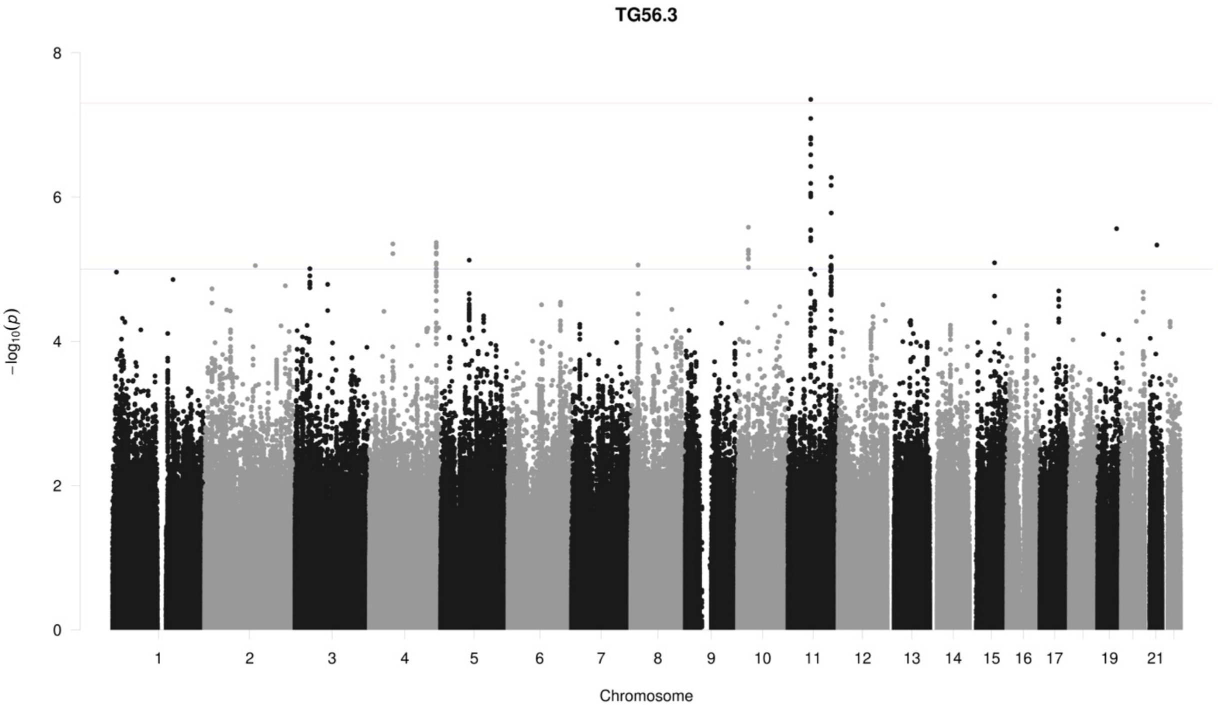
TG56.3
−log10(p)
0
2
4
6
8
19
21
Chromosome

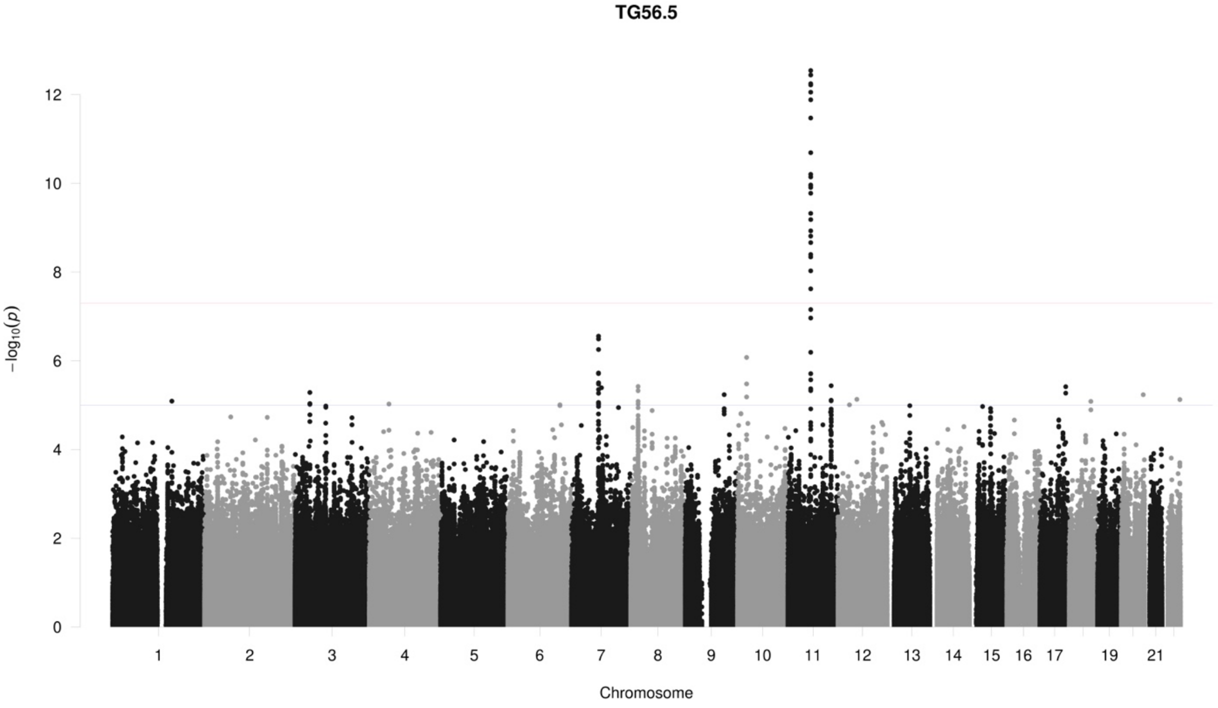
TG56.5
−log10(p)
0
2
4
6
8
10
12
19
21
Chromosome

**TG56.6**

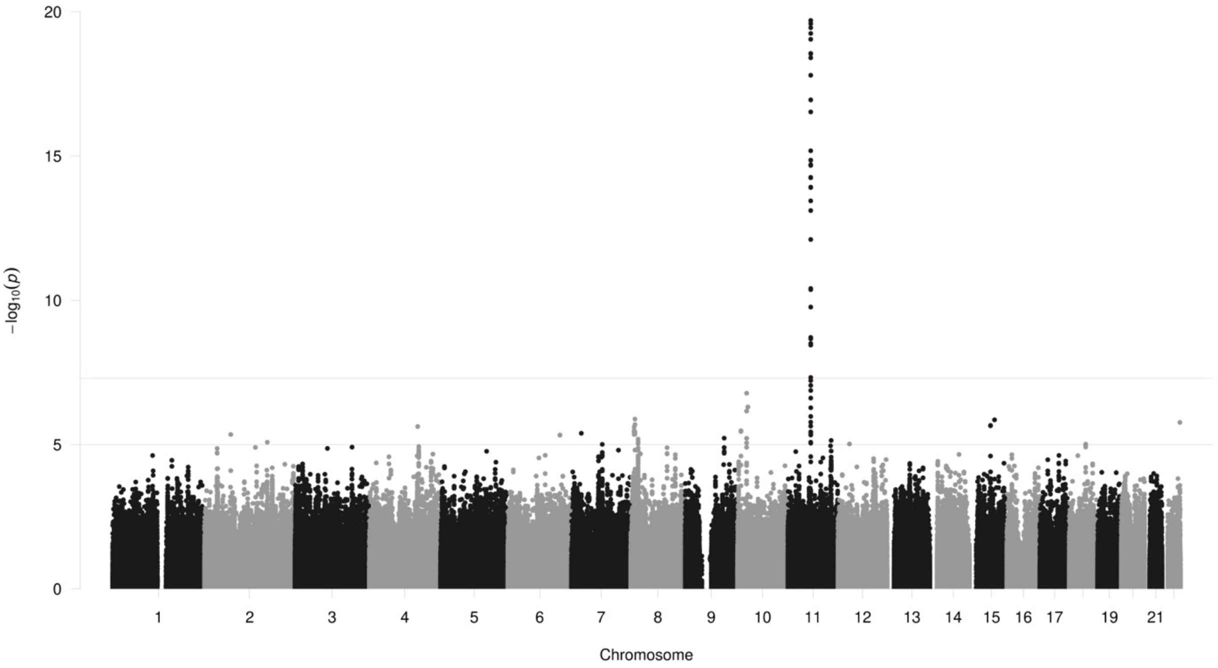

20
15
10
5
0
−log10(p)
19
21
Chromosome

**TG56.7**

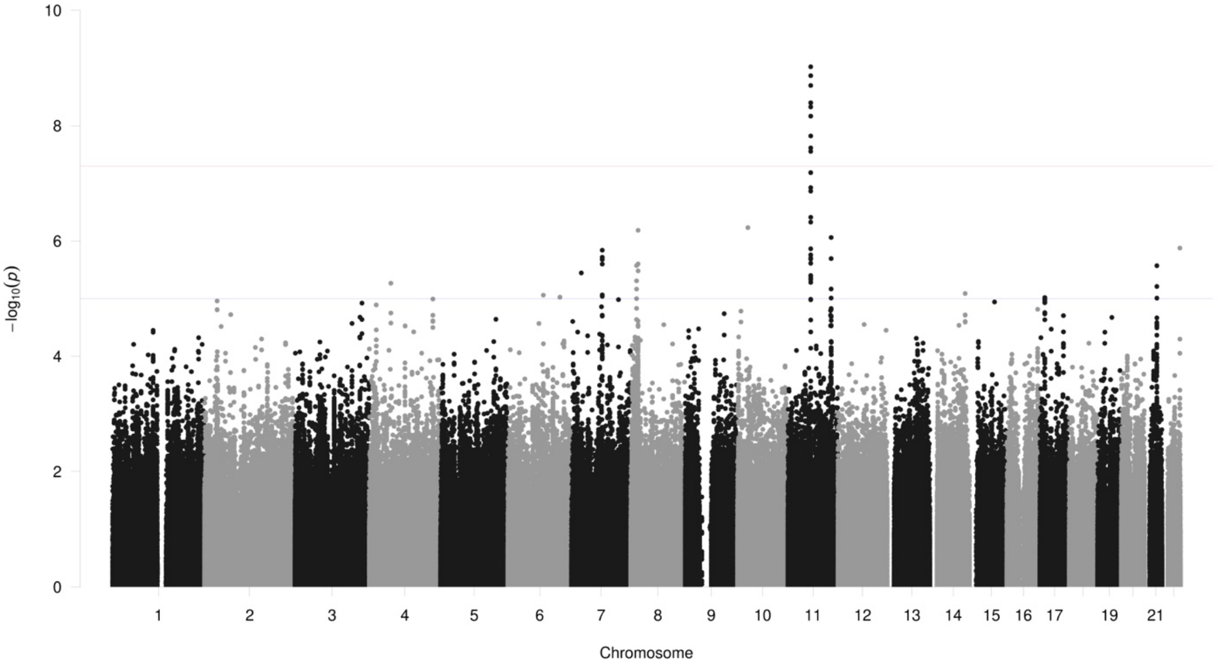

10
8
6
4
2
0
−log10(p)
19
21
Chromosome
